# A novel family of nonribosomal peptides modulate collective behavior in *Pseudovibrio* bacteria isolated from marine sponges

**DOI:** 10.1101/2020.12.15.422899

**Authors:** Laura P. Ióca, Yitao Dai, Sylvia Kunakom, Jennifer Diaz-Espinosa, Aleksej Krunic, Camila M. Crnkovic, Jimmy Orjala, Laura M. Sanchez, Antonio G. Ferreira, Roberto G. S. Berlinck, Alessandra S. Eustáquio

## Abstract

Collective behavior is a common feature of life. Although swarming motility and biofilms are opposed collective behaviors, both contribute to bacterial survival and host colonization. We have identified a link between motility/biofilms and a nonribosomal peptide synthetase-polyketide synthase gene cluster family (*ppp*) conserved in *Pseudovibrio* and *Pseudomonas* Proteobacteria known to interact with diverse eukaryotes. After developing reverse genetics for *Pseudovibrio*, we discovered two pseudovibriamide families, heptapeptides with a reversal in chain polarity via an ureido linkage **1**-**6** and related nonadepsipeptides **7**-**12**. Imaging mass spectrometry showed that **1** was excreted whereas **7** was colony-associated. Deletion of *pppA* abolished production of **1**-**12** leading to reduced motility and increased biofilm production. *pppD* mutants that produced only **1**-**6** showed motility comparable to the wild-type and reduced biofilm formation, indicating that the excreted heptapeptides play a role in promoting motility. In contrast to lipopeptides widely known to affect swarming and biofilms, pseudovibriamides are not surfactants. Our results expand current knowledge on metabolites mediating bacterial collective behavior. Moreover, the establishment of reverse genetics will enable future exploration of the ecological and biotechnological potential of *Pseudovibrio* bacteria which have been proposed to contribute to marine sponge health.

**Significance:** Bacteria contribute to health and disease of plants and animals. Specialized metabolites produced by bacteria are important in mediating their behavior and the colonization of their hosts. We have identified a conserved gene cluster family in *Pseudovibrio* and *Pseudomonas* bacteria known to colonize marine animals and terrestrial plants, respectively. Using *Pseudovibrio* as a model, we show the encoded metabolites, which we termed pseudovibriamides, promote motility and decrease biofilms. In contrast to lipopeptides widely known to affect motility/biofilms, pseudovibriamides are not surfactants, but instead are linear peptides with a reversal in chain polarity. The discovery of pseudovibriamides expands current knowledge of bacteria collective behavior. The establishment of reverse genetics will enable exploration of the ecological and biotechnological potential of *Pseudovibrio* bacteria.

**Classification:** Biological Sciences, Microbiology

## Introduction

Collective behavior is a common feature of biological systems from schools of fish and flocks of birds to swarms of bacteria. Bacterial swarming is the collective motion of bacterial cells on a solid surface, which is powered by rotating flagella^1,2^. In contrast, biofilms are nonmotile, self-organized cellular aggregates bounded by extracellular polymers^3^. Although bacterial biofilms represent a concept opposite to motility, both motility and biofilm formation contribute to population survival and host colonization^4,5^. Identifying which specialized metabolites mediate these processes will bring us closer to understanding collective, coordinated behavior in bacteria.

Herein we report a link between swarming and biofilms, and a nonribosomal peptide synthetase (NRPS) polyketide synthase (PKS) gene cluster family that is conserved in both *Pseudovibrio* and *Pseudomonas* bacteria known to colonize animals and plants (Fig. 1). We termed this biosynthetic gene cluster (BGC) *ppp* for *<u>P</u>seudovibrio* and *<u>P</u>seudomonas* nonribosomal <u>p</u>eptide.

**Fig. 1.**
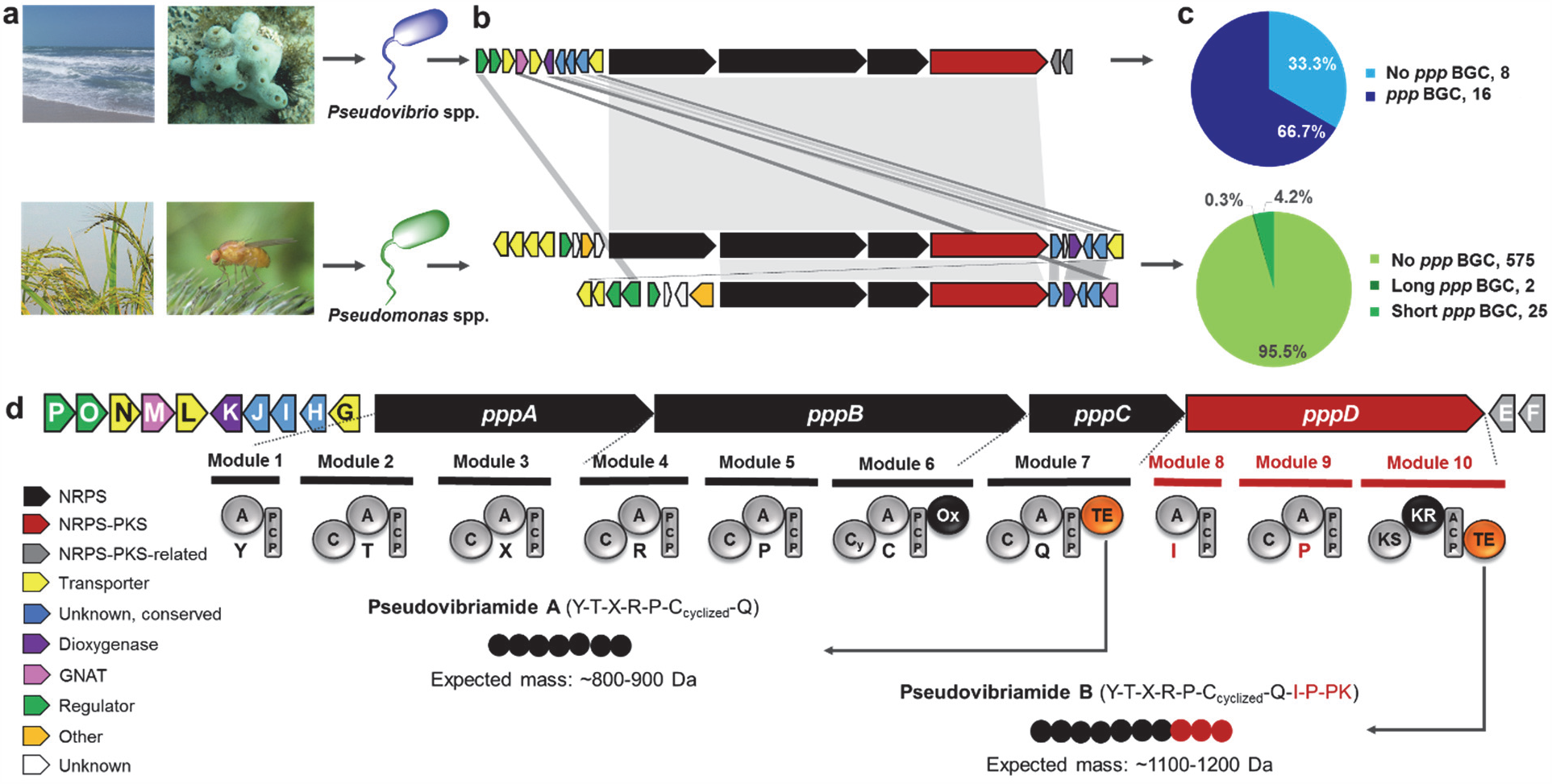
The *ppp* gene cluster family conserved in *Pseudovibrio* and *Pseudomonas* bacteria known to establish associations with eukaryotic hosts. (**a**) *Pseudovibrio* and *Pseudomonas* species isolated from marine invertebrates, and from terrestrial plants and animals, respectively, contain a member of a (**b**) hybrid nonribosomal peptide synthetase (NRPS) polyketide synthase (PKS) biosynthetic gene cluster (BGC) family we named *ppp* for *<u>P</u>seudovibrio* and *<u>P</u>seudomonas* nonribosomal <u>p</u>eptide. (**c**) Prevalence of the *ppp* BGC in available *Pseudovibrio* (24) and *Pseudomonas* (602) genomes. For *Pseudovibrio*, all available genomes were analyzed, whereas for *Pseudomonas*, only complete genomes. (**d**) The *ppp* BGC from *Pseudovibrio brasiliensis* Ab134 and its predicted products. Product structures were predicted based on the NRPS-PKS modular architecture and the specificity code of adenylation (A) domains. The single letter amino acid code is indicated below each A domain. If the amino acid could not be predicted, it is indicated with “X”. The polyketide unit is indicated as PK and predicted to be acetate. Products are represented as beads on a string. The string containing seven black beads represents heptapeptides that would be formed if assembly stopped at the first thioesterase (TE) on module 7, which we later termed pseudovibriamides A; whereas the string also including the three red beads represent the full-length products pseudovibriamides B. Genes in panels **b** and **d** are colored based on predicted function and as shown in panel **d**: NRPS, black; NRPS-PKS, red; NRPS-PKS-related (4’-phosphopantetheinyl transferase and type II thioesterase, respectively), grey; transporter, yellow; dioxygenase, purple; three unknown genes that are conserved in all BGCs, blue; other unknown, white; regulatory gene, green; GCN5-related *N*-acetyltransferase (GNAT), pink; other function; orange. Domain key: A, adenylation; ACP/PCP, acyl/peptidyl carrier protein; C, condensation; Cy, condensation and heterocyclization; KR, ketoreductase; KS, ketosynthase; Ox, oxidase; TE, thioesterase.

*Pseudomonas* is a genus of γ-Proteobacteria discovered in the 19^th^ century. *Pseudomonas* species are ubiquitous in both terrestrial and aquatic environments. *Pseudomonas* species can be free-living or associated with eukaryotic hosts including fungi, plants, and animals, with some species being pathogenic to their hosts^6^. In contrast, the genus *Pseudovibrio* of marine α-Proteobacteria was only recently described in 2004^7^. *Pseudovibrio* bacteria have been predominantly isolated from healthy marine sponges^8^. The current hypothesis is that *Pseudovibrio* spp. form a mutualistic relationship with marine sponges, although details of the interaction remain to be elucidated^8,9^. The identification of specialized metabolites produced by *Pseudovibrio* is an important step towards understanding *Pseudovibrio*’s ecological roles. For instance, *Pseudovibrio* spp. produce the antibiotic tropodithietic acid^10^ which is active against pathogenic, marine *Vibrio* spp. Tropodithietic acid likely contributes to the mutualistic relationship between *Pseudovibrio* and marine invertebrates, by serving a protective role to the animal host^11^. Additional metabolites reported from *Pseudovibrio* include the cytotoxic 2-methylthio-1,4-naphthoquinone and indole alkaloid tetra(indol-3-yl)ethanone, bromotyrosine-derived alkaloids, the red pigment heptyl prodiginin, and anti-fouling diindol-3-ylmethanes^8,12–15^.

In this investigation we report the complete genome of *Pseudovibrio brasiliensis* Ab134, the establishment of reverse genetics methods for this marine sponge-derived isolate, the discovery and structures of a novel family of nonribosomal peptides which we named pseudovibriamides (**1**-**12**), and pseudovibriamides’ role in affecting flagellar motility and biofilm formation. Pseudovibriamides A1-A6 (**1**-**6**) are heptapeptides containing three modified amino acids: a thiazole, a dehydrated threonine (dehydroaminobutyric acid), and a reversal in chain polarity of the first amino acid (tyrosine) through an ureido linkage. Pseudovibriamides B1-B6 (**7**-**12**) are nonadepsipeptide derivatives containing a propionylated hydroxyproline. Using *Pseudovibrio* as a model, our results reveal novel specialized metabolites employed by bacteria to modulate motility and biofilms, ultimately expanding current knowledge of bacteria collective behavior.

## Results

### The *ppp* BGC from *Pseudovibrio* is located in a plasmid

*P. brasiliensis* strain Ab134 was isolated from the sponge *Arenosclera brasiliensis*^16^. A draft genome had previously been reported^17^. Here, we re-sequenced *P. brasiliensis* strain Ab134 using short-read Illumina sequencing and long-read Nanopore sequencing. Hybrid assembly followed by confirmation of plasmids via PCR (Fig. S1 and Tables S1-S2) led to a genome of 6.03 Mb divided into one chromosome and five plasmids (Fig. 2 and Table S3). Seven biosynthetic gene clusters were identified using antiSMASH (Table S4)^18^, including the NRPS-PKS *ppp* gene cluster located on plasmid 2 (Fig. 2). Using BLAST analyses we further identified a *tda* gene cluster putatively encoding the biosynthesis of the known antibiotic tropodithietic acid^19^ on plasmid 1 (Fig. 2).

**Fig. 2.**
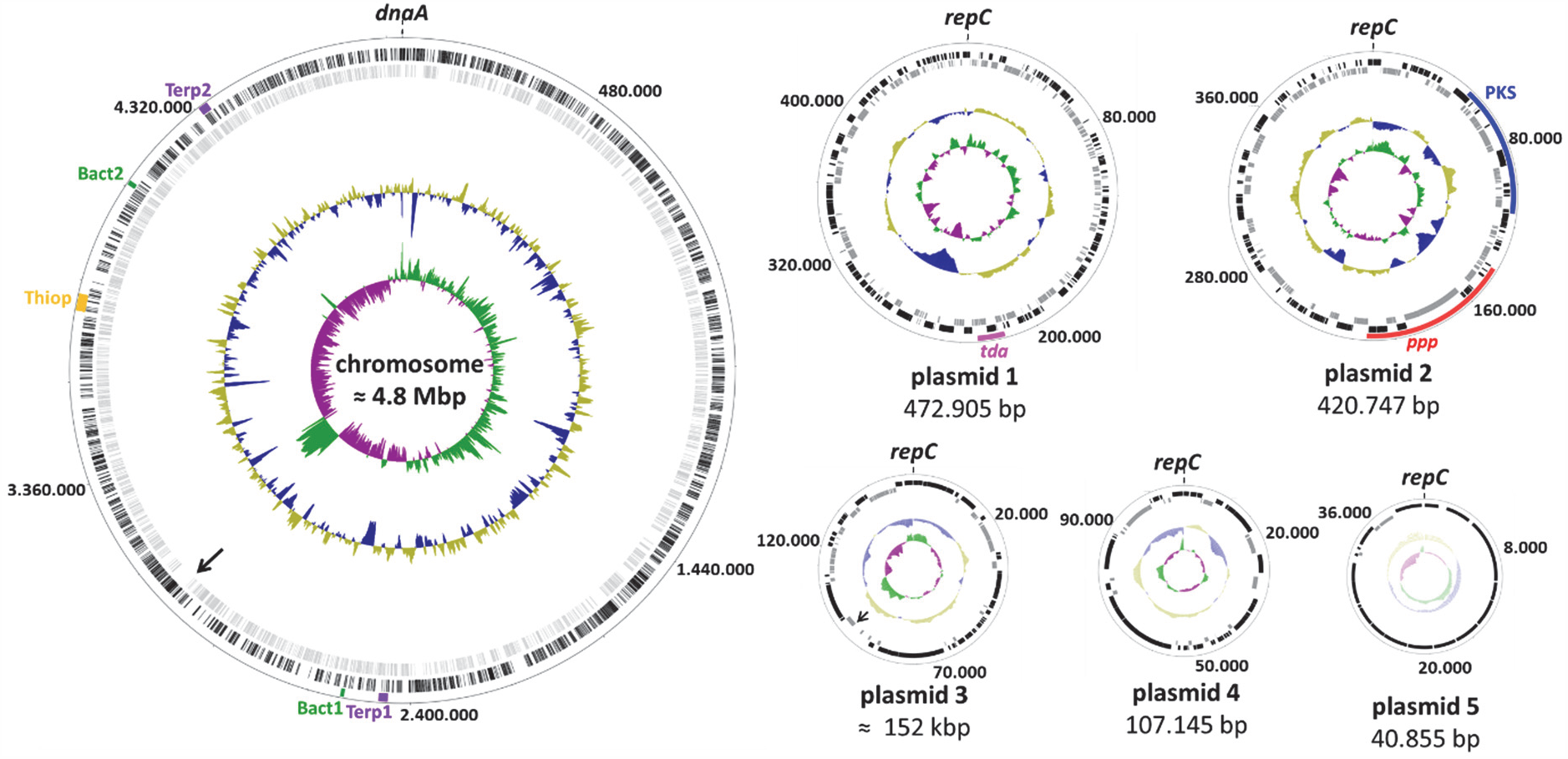
The genome of *Pseudovibrio brasiliensis* Ab134. The outer circle depicts the size in base pairs. Lane 1 (from the outside in) shows the eight predicted BGCs, color-coded based on biosynthetic class: bact, bacteriocin (green); terp, terpene (purple); thiop, thiopeptide (yellow); *tda*, tropodithietic acid (pink); PKS, polyketide synthase (blue). The *ppp* BGC (red) investigated in this study is located on plasmid 2. Lanes 2 and 3 show predicted open reading frames (ORFs) on the leading (black) and lagging (gray) strands, respectively. Lanes 4 and 5 depict normalized plot of GC content (yellow/blue) and normalized plot of GC skew (purple/green), respectively. The chromosome is oriented to *dnaA*, plasmids are oriented to *repC*. See Table S3 for further details. Replicons are represented not strictly to scale. Two remaining gaps (one in the chromosome and one in plasmid 3) are indicated with an arrow.

### The *ppp* BGC is conserved in eukaryote-interacting bacteria

Bioinformatics analyses using multigene blast^20^ revealed the *ppp* BGC to be conserved in various species of *Pseudovibrio* and *Pseudomonas* (Fig. 1c, Fig. S2, and Table S5). We were intrigued by the fact that the *ppp* BGC was conserved in different classes of Proteobacteria known to colonize eukaryotic hosts and decided to investigate it further. All members of the *ppp* BGC family include one NRPS-PKS gene (*pppD*) and either three or two NRPS genes (*pppABC* or *pppBC*, respectively, Fig. 1b). Other conserved genes encode a transporter (*pppG*), three proteins of unknown function (*pppHIJ*), and an iron/α-ketoglutarate-dependent dioxygenase (*pppK*). Additional transporter genes (e.g. *pppL* and *pppN*) and regulatory genes (e.g. *pppP* and *pppO* encoding a two-component system sensor histidine kinase and response regulator, respectively) are also found flanking the clusters (Fig. 1b). Moreover, the BGC present in *Pseudovibrio* also encodes a phosphopantheteinyl transferase (*pppE*), and a type II thioesterase (*pppF*) not present in *Pseudomonas* BGCs, and a *N*-acetyltransferase family protein (*pppM*) that is present only in the short version of the *Pseudomonas* BGC (Fig. 1b).

The larger version of the *ppp* BGC is found in 16 out of 24 *Pseudovibrio* genomes available in GenBank, 13 of which have been isolated from marine sponges, two from seawater, and one from a marine tunicate. In addition, this larger version of the *ppp* BGC was also found in at least two *Pseudomonas* species known to be plant pathogens, *Pseudomonas asplenii* ATCC23835 and *Pseudomonas fuscovaginae* LMG2158. The shorter version of the *ppp* BGC was more common in *Pseudomonas* than the longer one and was found in about 4% of *Pseudomonas* genomes (25 out of 602 complete genomes analyzed) including plant pathogen *P. syringae* pv. *Lapsa* ATCC10859, and insect pathogen *P. entomophila* L48. Based on the NRPS-PKS domain organization (Fig. 1d), we predicted the full-length product to contain nine amino acids and one acetate unit. The amino acid prediction for all *ppp* BGCs is highly conserved (Fig. S3). The domain organization found in the *ppp* assembly line is reminiscent of the shorter rimosamide NRPS-PKS of *Streptomyces rimosus* (Fig. S2, BGC 19-21), which likewise contains two thioesterase domains and a terminal, AT-less PKS module^21^.

### Establishment of reverse genetics for *P. brasiliensis*

In order to aid with compound identification, and to probe the function of the *ppp* BGC, we developed reverse genetics for *P. brasiliensis* Ab134. No methods for genetic engineering of *Pseudovibrio* had been previously reported. After establishing procedures based on protocols available for other members of the Rhodobacteraceae family^22,23^, mutants in which the *pppA* gene was replaced with a kanamycin-resistance marker via homologous recombination were obtained (Fig. S4a-b). Comparative metabolite analysis of the wild-type and three independent mutants was then employed to identify the encoded products.

### Identification of pseudovibriamides

Comparative metabolite analysis of the wild-type and *pppA* mutant strains using mass spectrometry led to the identification of five signals that were detected only in the wild-type Ab134 strain (Fig. 3a) suggesting they were the products of the *ppp* gene cluster. The two larger compounds of *m/z* 1156 and 1170 differ only by a methyl group (Δ14 Da), while tandem mass spectrometry (LC-MS/MS) analyses (Fig. S5) showed that a smaller *m/z* 844 was present as a fragment of the two larger compounds, indicating their relatedness. The other two smaller compounds, *m/z* 826 and *m/z* 858 differ from *m/z* 844 only by the removal of water (-Δ18 Da) and the addition of a methyl group (+Δ14 Da), respectively.

**Fig. 3.**
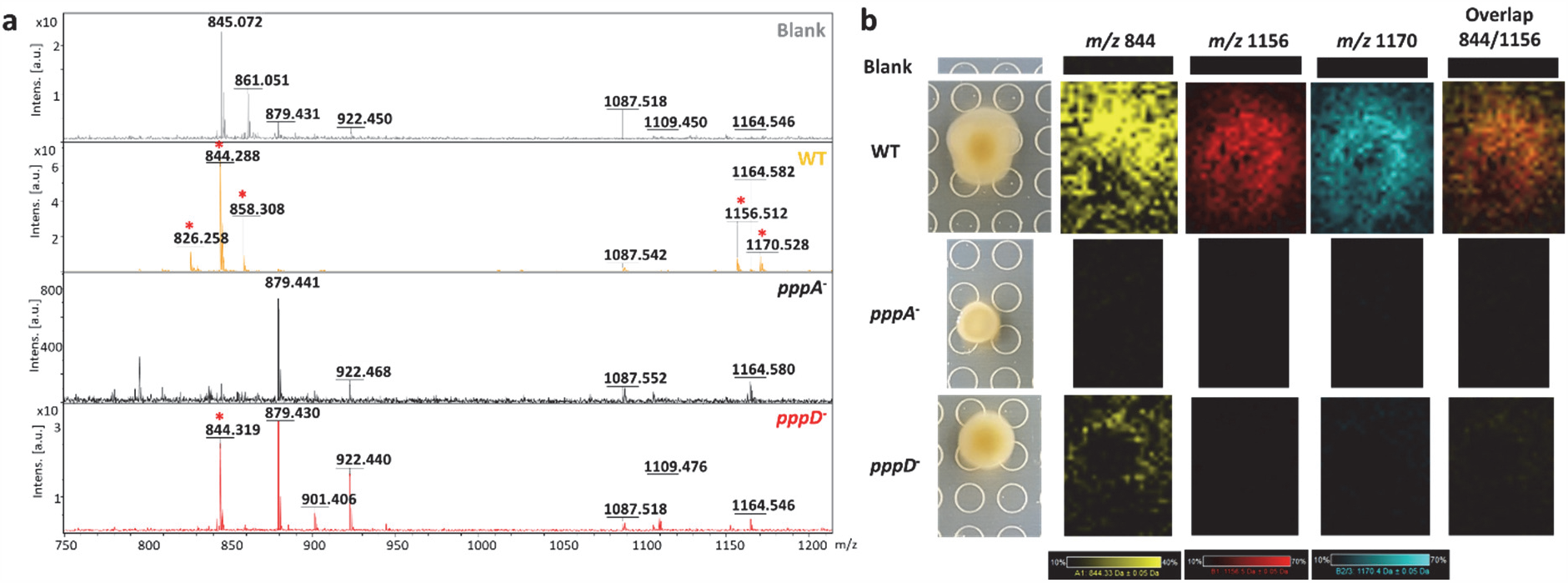
Comparative metabolite analysis of wild-type, *pppA* and *pppD* mutant strains of *P. brasiliensis* Ab134 using MALDI-TOF mass spectrometry. (**a**) dried-droplet analysis of culture extracts. Peaks of interest are highlighted with red asterisks. (**b**) Imaging mass spectrometry analysis showing the spatial distribution of selected compounds on and around microbial colonies. *P. brasiliensis* wild-type and mutant strains were grown on marine agar for 18 h before matrix application and analysis.

In order to test whether the smaller compounds with *m/z* ∼800 would be the product of modules **1-7** (*pppABC*, Fig. 1d) and to investigate the role of these smaller compounds, *pppD* mutants were generated by replacement with a kanamycin-resistance marker as done for *pppA* (Fig. S4c). Mass spectrometry confirmed that *pppD* mutants do not produce the full-length products and accumulate only the heptapeptides (Fig. 3a). We name the heptapeptides pseudovibriamides A and the full-length products pseudovibriamides B.

### Pseudovibriamide A is excreted and pseudovibriamide B is cell-associated

Investigation of the spatial distribution of pseudovibriamides in microbial colonies by MALDI-TOF imaging mass spectrometry (IMS) analyses (Fig. 3b) indicated that the full-length products (*m/z* 1156 and 1170) were colony-associated whereas the smaller heptapeptide *m/z* 844 appeared to be excreted. This observation was corroborated by our inability to identify the full-length products in liquid culture supernatants whereas the heptapeptides are readily identified.

### Isolation and structure elucidation of pseudovibriamides A

After testing four different growth conditions (Fig. S6), growth on solid media (swarming assay conditions) provided superior pseudovibriamide relative yields compared to liquid cultures (Fig. S6 and Online Methods). Three growth experiments and extraction procedures were carried out to obtain enough material for isolation and structure elucidation. We first extracted 400 swarming plates (8-L, extract F) with MeOH. After several rounds of fractionation and HPLC purification, we obtained 5.5 mg of **1** and 2.7 mg of **2**. The HRESIMS (Fig. S7) analysis of **1** displayed a [M + H]^+^ at *m/z* 844.3389, corresponding to a molecular formula C_36_H_50_N_11_O_11_S (calc. 844.3412, error -2.3 ppm). Inspection of 1D and 2D NMR data for **1** (Figs. S8-S15, Table S6) allowed the identification of five amino acids as Tyr, Ala, Arg, Pro and Gln. Two additional modified residues were identified. First, a dehydrated Thr (dehydroaminobutyric acid, *Z*-Dhb), based on a COSY correlation between the methyl group at δ_H_ 1.59 (d) and the vinylic methine hydrogen at δ_H_ 5.95 (q), as well as by observing a ROESY correlation between the methyl resonance at δ_H_ 1.59 with NH at δ_H_ 7.81, pointing to a *Z*-DhB residue. Second, a thiazole group (thz), based on HMBC coupling from H_β_ (thz, δ_H_ 8.16, s) to C_α_ (Pro, δ_C_ 58.5), C_α_ (thz, δ_C_ 149.1), C_δ_ (thz, δ_C_ 173.7) and the Gln amide carbonyl group at δ_C_ 173.2.

The connectivity between the amino acid residues was confirmed by observation of the key HMBC correlations as shown in Fig. S16 and by HRESIMS/MS analysis (Fig. S7, and Table S7). Inspection of the MS^2^ fragmentation pattern (Table S7) indicated the loss of a complete Tyr residue (*m/z* of 663.2673). A second ion at *m/z* 637.2866 indicated the loss of a Tyr-CO fragment. Such fragment losses would be possible due to the presence of an ureido linkage, leading to a reversal of chain polarity. HMBC correlations from the amine (NH-Tyr, δ_H_ 6.42) and C_α_H_2_ (Tyr, δ_H_ 4.30) to C_CO_ (δ_C_ 155.2), in addition to ^15^N-HMBC (Fig. S15) correlations from the NH_Tyr_ (δ_H_ 6.42) to the *N*-ureido nitrogen at δ_N_ 88.51, as well as from the NH_Dhb_ (δ_H_ 7.81) to the second N-ureido nitrogen at δ_N_ 92.17 (Table S6), confirmed our hypothesis. The absolute configurations of all amino acids was established as L by advanced Marfey’s analysis (Figs. S17-21)^24,25^, thus completing the structure of **1**, named pseudovibriamide A1 (Fig. 4).

**Fig. 4.**
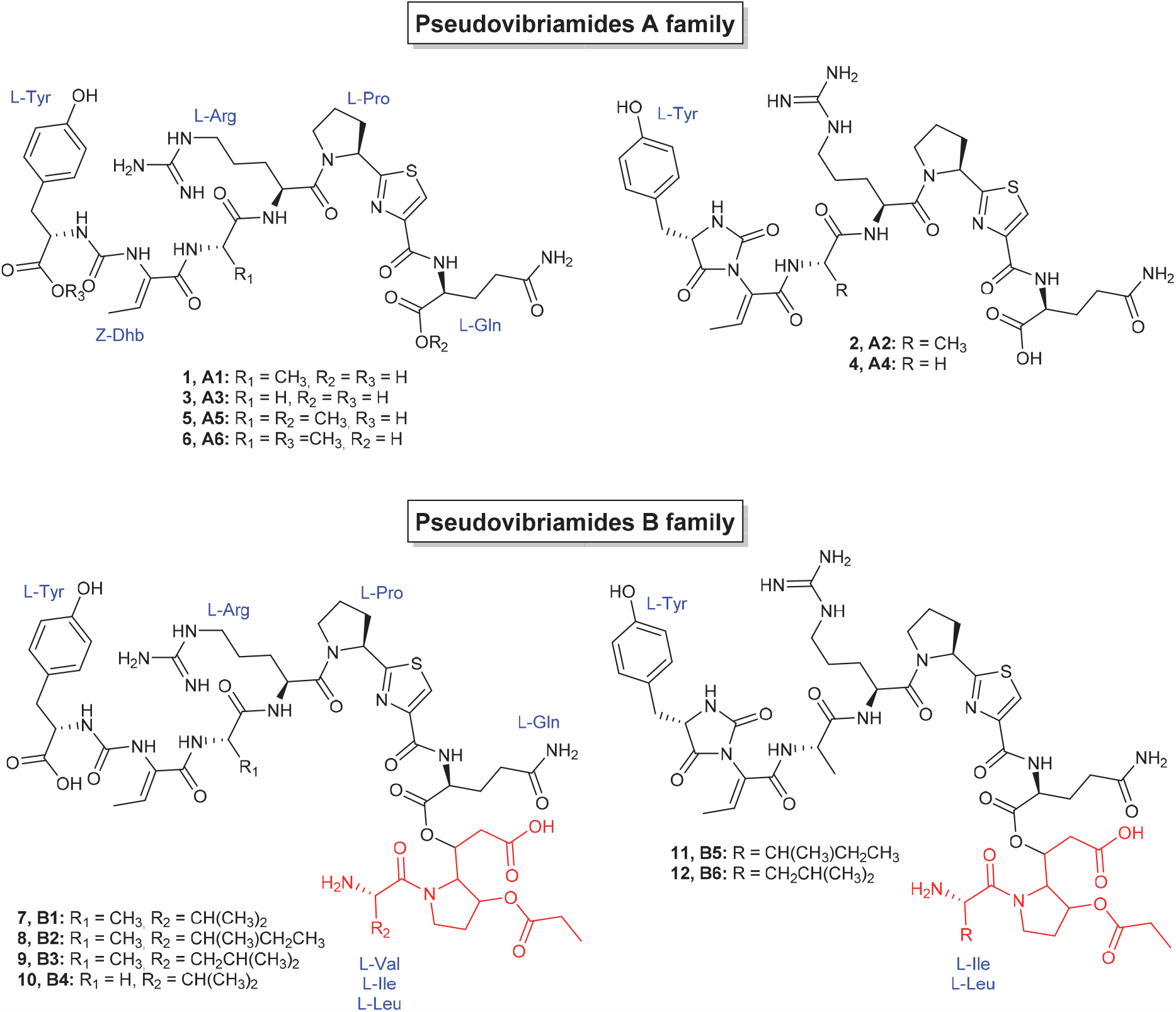
Structures of pseudovibriamides isolated from *P. brasiliensis* Ab134 and identified by analysis of spectroscopic data. Marfey’s analyses were performed for all pseudovibriamides, except **5** and **6**. The portion of the pseudovibriamide B structures originating from PppD is colored red.

A second growth experiment was performed after which we freeze-dried 600 swarming plates (12-L, extract FD) prior to extraction with MeOH. Fractionation and HPLC purification of the MeOH extract provided 2 mg of **2** and additional 8.8 mg of **1**. Compound **2** samples obtained from the two growth experiments were combined. The HRESIMS (Fig. S22) analysis of **2** displayed a [M + H]^+^ at *m/z* 826.3311, corresponding to a molecular formula C_36_H_48_N_11_O_10_S (calc. 826.3306, error 0.6 ppm), suggesting **1** might undergo a dehydration and/or cyclization leading to **2**. Inspection of the MS^2^ fragmentation pattern (Fig. S22, Table S8) indicated a cyclization between Tyr and *Z*-Dhb. Further analysis of 1D and 2D NMR data for **2** (Figs. S23-S29, Table S9) and advanced Marfey’s analysis allowed the confirmation of L-Tyr, *Z*-Dhb, L-Ala, L-Arg, L-Pro-thz and L-Gln (Figs. S30-S34). The HMBC NMR data of compound **2** did not point to the presence of a NH group for Dhb. Thus, we observed the formation of a unique imidazolidinyl-dione ring leading to **2**, which agrees with the complete spectroscopic data and that we named pseudovibriamide A2 (Fig. 4).

Further fractionation of the FD extract led to the isolation of 1 mg of **3** and 0.2 mg of **4** (Fig. 4). Analysis of the MS^2^ fragmentation pattern (Tables S10-S11, Figs. S35-S36) and advanced Marfey’s analysis (Figs. S37-S45) of **3** and **4** indicated the presence of a Gly instead of Ala (Fig. S37) as the only difference between **1** and **2**. Compounds **3** and **4** were named pseudovibriamide A3 and A4, respectively. We also obtained 0.5 mg of a mixture of **5** and **6** containing two chromatographic peaks with [M + H]^+^ at *m/z* 858.35 (Fig. S46) and molecular formula C_37_H_52_N_11_O_11_S, indicating the presence of a methylated form of **1**. The MS^2^ fragmentation data (Figs. S47-S48, Table S12) supports the structures **5** and **6** (Fig. 4), methylated at Gln and Tyr, respectively. Although these compounds are possibly artifacts of MeOH extraction, such hypothesis requires further experimental confirmation.

### Isolation and structure elucidation of pseudovibriamides B

Because we were unable to obtain enough pure, full-length product from the 8-L and 12-L cultivations described above, we performed a larger scale cultivation and modified the extraction procedure. Since the IMS experiments had indicated that the larger peptides were cell-associated, one thousand swarming plates were prepared (20-L), and the cells were harvested from the plates (Fig. S49). MS-guided purification provided 5.9 mg of **7**, 6.2 mg of a mixture of **8** and **9**, 2.7 mg of **10** and 1.4 mg of a mixture of **11** and **12**. Unfortunately, pseudovibriamides B degraded during the acquisition of NMR data. Degradation was also observed for pseudovibriamide A1, since the bond between NH_Dhb_ and C_α’-Dhb_ is easily hydrolyzed (Figs. S50-51, Table S13). The HRESIMS (Fig. S52) analysis of **7** displayed a [M + H]^+^ at *m/z* 1156.5106, corresponding to a molecular formula C_51_H_74_N_13_O_16_S (calc. 1156.5097, error 0.8 ppm). Its degradation product (Fig. S53) displayed [M + H]^+^ at *m/z* 950.4446, corresponding to the molecular formula C_41_H_64_N_11_O_13_S (calc. 950.4406, error 4.2 ppm). The MS^2^ data indicated that peptide **7** shares the carbon skeleton of **1** (Fig. S7); therefore, we could identify the presence of L-Tyr, *Z*-Dhb, L-Ala, L-Arg, L-Pro-thz and L-Gln by NMR analysis (Figs. S54-S60, Table S14) and confirm the presence of L-amino acids by Marfey’s analysis (Figs. S61-S66). Analysis of 1D and 2D NMR spectra of **7** and advanced Marfey’s analysis indicated the presence of a L-Val residue (Fig. S66). Based on homology between the *ppp* BCG and the rimosamide BGC, we hypothesized that **7** also presented a modified hydroxyproline (Hyp) residue. An HMBC correlation between the CH of δ_H_ at 5.10 (δ_C_ 70.4) and CO (173.0) indicated a propionylated Hyp fragment. The correlation between CH_α’-Hyp_ at δ_H_ 5.33 (δ_C_ 69.3) and CO_Gln_ at δ_C_ 170.8 confirmed the final nonadepsipeptide structure of **7** (Fig. 4), named as pseudovibriamide B1. Additional confirmation of the amino acid sequence was established by HRESIMS/MS analysis (Fig. S52 and Table S15).

Pseudovibriamides B2 (**8**) and B3 (**9**) differ from **7** by a methyl group, due to the presence of either L-Ile or L-Leu instead of L-Val, as shown by the respective MS (Fig. S67 and Table S16) and advanced Marfey’s analyses (Fig. S68-S76). Comprehensive and detailed analysis of MS and MS^2^ data of pseudovibriamides B4 to B6 (Fig. S77-S78, Table S17-S18) together with advanced Marfey’s analysis (Fig. S79-S92) led to the proposed structures for **10** to **12**, respectively (Fig. 4).

### Pseudovibriamides modulate collective behavior

During the process of generating *pppA* mutants, we noticed that the diameter of *pppA* mutant colonies was significantly smaller than wild-type colonies grown on solid agar (Fig. S93). Based on this observation, we hypothesized the products of the *ppp* BGC may be involved in surface motility. The only surface motility mode^1,2^ that has been previously described for both α- and γ-Proteobacteria is swarming. Yet, *Pseudovibrio*’s ability to swarm had not been previously reported. *Pseudovibrio* spp. do contain one to several lateral or subpolar flagella^7,26^ so that the minimum requirement for swarming motility is met. Indeed, we were able to show that the wild-type Ab134 is capable of swarming (Fig. 5a and Fig. S94). Moreover, we found that Eiken agar is required to allow observation of the swarming phenotype, as swarming could not be observed on more commonly used agars such as Bacto agar, most likely due to Eiken agar’s lower surface tension (Fig. S94). Agars of inherently low surface tension have been shown to be required for swarming of bacteria that do not produce surfactants^1^. Finally, Ab134 swarming motility decreases with increasing agar concentrations, as has been observed for other temperate swarmers (Fig. S95)^2,27^.

**Fig. 5.**
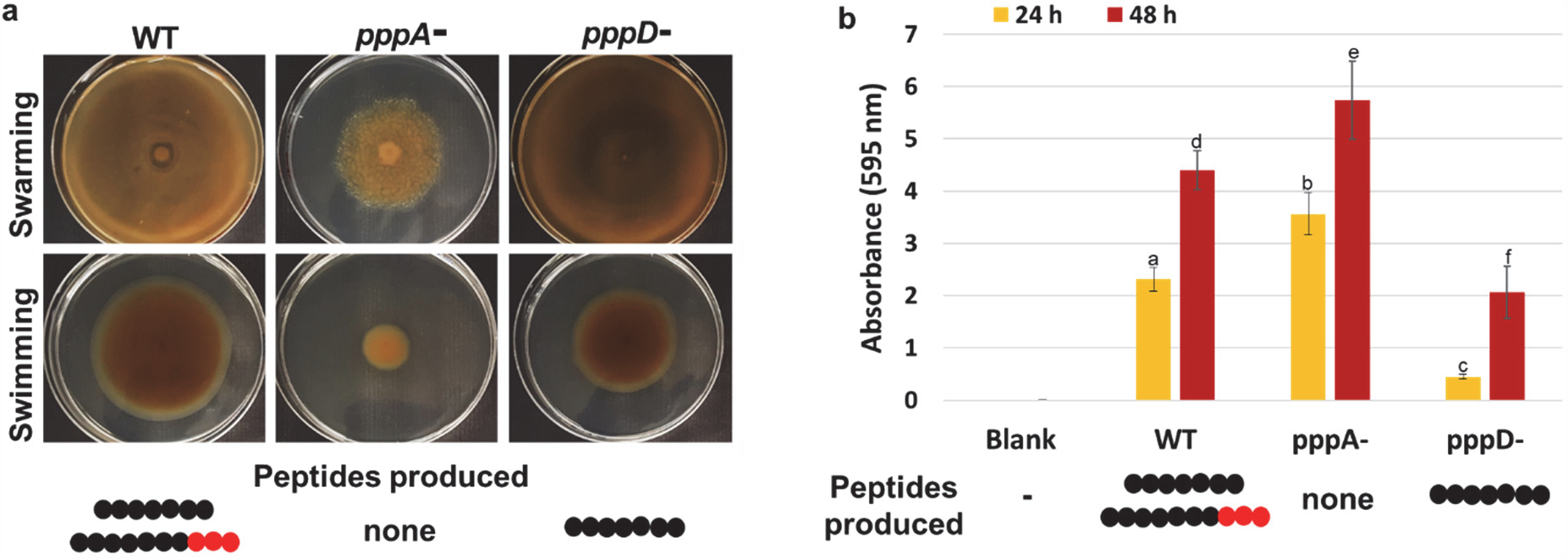
Effect of *pppA* and *pppD* deletion on flagellar motility and biofilm formation. (**a**) Swarming assay (top) and swimming assay (bottom) performed on marine agar containing either 0.5% or 0.3% Eiken agar, respectively. Plates were inoculated with 5 µL of cryo-preserved cultures of Ab134 wild-type (WT) or *pppA*^-^/*pppD*^-^ mutant strains normalized to OD_600_ of 1.0. Pictures shown were taken at 48 hours after inoculation. The assay was performed multiple times, each time in triplicates, and similar results were obtained each time. Representative results are shown. (**b**) Biofilm assay using 0.1% crystal-violet solution to quantify biofilm adherence and microcolony formation. Bars are average of six replicates (Table S19), and error bars represent standard deviation. Lowercase letters above bar indicate statistically different values according to the Tukey test 95% confidence level (Table S19). The seven black beads on a string represent heptapeptides **1**-**4** produced by the wild-type and *pppD*^-^ mutant strains; whereas the string also including the three red beads represent the full-length nonadepsipeptides **7**-**12** produced by the wild-type.

Having established swarming assays using 0.5% (w/v) Eiken agar, we were then able to show that the *pppA* mutant has an impaired swarming phenotype (Fig. 5a). Because swimming motility is, like swarming, powered by flagella, we investigated whether swimming was also affected by the *pppA* deletion. A swimming assay using low agar concentration (0.3%) showed that loss of pseudovibriamides A and B also impaired swimming motility (Fig. 5a). All three independent *pppA* mutants analyzed showed the same impaired motility phenotype (Fig. S96). To make sure that no unintended mutations were introduced during mutant generation, we sequenced the genome of one of the mutants using Illumina sequencing. Mapping of Illumina reads to our reference genome showed that no further mutations were present besides the *pppA* deletion. To provide further evidence for the link between the *ppp* gene cluster, pseudovibriamides, and the motility phenotype, a genetic complementation mutant was generated in which *pppA* was knocked back in the *pppA*^*-*^ mutant, i.e. the kanamycin resistance marker was replaced with *pppA* via homologous recombination leading to a reversal to the original wild-type genotype (Fig. S97). Pseudovibriamide production and the swarming phenotype were restored in the *pppA* knock-in mutant (Fig. S97).

Motility and biofilm development are considered opposite behaviors^28,29^. Thus, we employed an established biofilm assay^30^ to evaluate the effect of the *pppA* mutation on the ability of *Pseudovibrio* to form biofilms. We observed biofilm adherence and microcolony formation to be increased in the *pppA* mutant compared to the wild-type (Fig. 5b and Table S19). In contrast, flagellar motility of *pppD* mutants that still produce heptapeptides pseudovibriamides A was comparable to the wild-type (Fig. 5a), whereas biofilm adherence and microcolony formation was reduced in *pppD* mutants (Fig. 5b and Table S19).

### No surfactant activity was detected for *P. brasiliensis*

Surfactants are amphiphilic compounds that have been shown to lower surface tension and enable swarming. Biosurfactant examples include lipopeptides, glycolipids, steroid acids, and phospholipids. Based on their structures, no surfactant activity is expected for pseudovibriamides. Moreover, the requirement for the intrinsically low surface tension provided by Eiken agar (Fig. S94) suggests that *P. brasiliensis* Ab134 does not produce surfactants. To test that assumption, two accepted surfactant assays^31^ – drop collapse and oil spreading – were carried out. Indeed, no surfactant activity was observed for strain Ab134 under the assay conditions tested (Fig. S98).

## Discussion

Microbes play crucial roles in the function and health of plants and animals^32,33^. Marine sponges are the most ancient animals and are a dominant component of our oceans from reef systems to polar sea floors^34^. Microorganisms can comprise up to one-third of sponge biomass^34^. One of the main motivations to study the microbiome of sponges has been the discovery of bioactive molecules. However, ecological questions such as the importance of microbes to sponge health and the role of specialized metabolites produced by sponge microbes are especially relevant since sponge disease negatively affects the ecology of reef systems^35^. Extensive investigations of sponges’ microbiomes demonstrated that only α- and γ-Proteobacteria were dominant in most sponge species^36^. *Pseudovibrio* is a genus of α-Proteobacteria found in healthy sponges but absent from diseased ones, and it has been hypothesized that *Pseudovibrio* spp. contribute to marine sponge health^8^.

Lack of reverse genetics methods have hindered full exploration of the ecological and biotechnological potential of *Pseudovibrio* bacteria^8^. The reverse genetics procedures described here not only enabled identification of pseudovibriamides but also allowed us to identify their role in motility and biofilm formation. As *Pseudovibrio* bacteria are gaining increasing attention due to their apparently beneficial association with marine invertebrates and the bioactive compounds that have been identified to date, we expect these methods will be of broad interest to the scientific community^8^.

The widespread nature of the *ppp* gene cluster in *Pseudovibrio* spp. suggests that pseudovibriamides play an important ecological role in this genus. Our results show that the *ppp* gene cluster is located in a plasmid of *P. brasiliensis* Ab134 (Fig. 2). Although most *Pseudovibrio* genomes are draft genomes preventing determination of replicon count, the *ppp* BGC also appears to be present in plasmids of at least five other *Pseudovibrio* spp. (DSM17465, JCM12308, W64, Ad13 and Tun.PSC04-5.I4), due to the presence of replication genes *repB* or *repC* in the same contig (Table S20). In contrast, the *ppp* BGC is located in the chromosome of *Pseudomonas* strains (Fig. S99). Plasmid localization facilitates horizontal gene transfer and may help explain the wider distribution of the *ppp* BGC in *Pseudovibrio* compared to *Pseudomonas* (Fig. 1c). Moreover, it is interesting to note that the *ppp* BGC is located next to NRPS genes encoding lipopeptides in the two *Pseudomonas* species that contain the full-length *ppp* cluster (Fig. S99). *Pseudomonas* lipopeptides are known to influence swarming and biofilm formation^37^. Physical clustering of the *ppp* BGC with lipopeptide-encoding NRPS BGCs may facilitate co-regulation of different metabolites that are involved in the same processes.

The sequence of amino acids observed in pseudovibriamide’s structures is in agreement with the NRPS module organization (Fig. 6). The amino acid that could not be predicted was identified as L-Ala (module 3), for which the specificity code shows little conservation^38^. L-Gly can also be substituted at this position as in **3, 4**, and **10**. Moreover, three amino acid modifications are present in pseudovibriamides, i.e. heterocyclization of cysteine followed by oxidation leading to a thiazole ring formation, reversal of tyrosine’s chain polarity via an ureido linkage, and dehydration of Thr to yield Dhb acid.

**Fig. 6.**
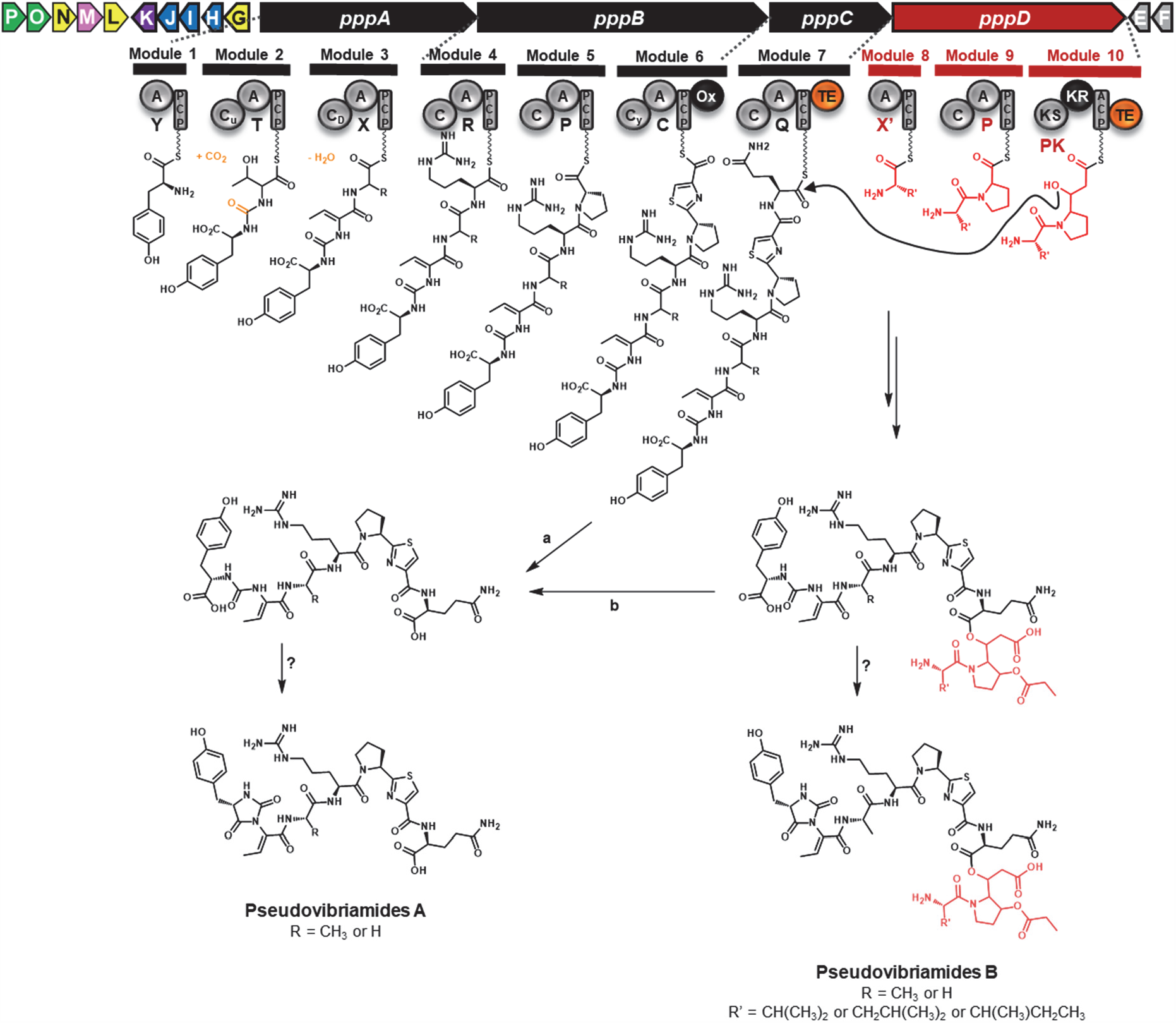
The *ppp* gene cluster of *P. brasiliensis* Ab134 and its products. Organization of the *ppp* gene cluster and proposed biosynthesis of pseudovibriamides. Codes are as in Fig. 1, except for C_u_, ureido-linkage formation condensation domain, and C_D_, dehydration condensation domain. Two potential pathways for accumulation of pseudovibriamides A are indicated (**a**) shunt products of module 7 or (**b**) ester hydrolysis of pseudovibriamides B.

Thiazole biosynthesis in nonribosomal peptides is well documented and involves a heterocyclization (Cy) domain and an oxidoreductase domain^39^, both of which are present in the respective module encoded in *pppB* (Fig. 6). Several peptides harbor a reversal in chain polarity through an ureido linkage (Table S21). The formation of this unusual connection has been investigated *in vitro*, indicating that the C domain catalyzes the reaction using bicarbonate/CO_2_ ^40,41^. Phylogenetic analysis of domains encoded in *pppA* from *Pseudovibrio* and *Pseudomonas* revealed that the first C domain (at module 2) clades together with condensation domains known to lead to ureido linkages (Fig. S100), supporting the ureido functionality determined from spectroscopic data. The implications of this structural feature for the phenotypes associated with the pseudovibriamides are, as of yet, unknown.

Finally, for all full-length *ppp* BGCs, the second A domain present in PppA is predicted to load Thr (Fig. S3). Our spectroscopic analyses of pseudovibriamides supports dehydration of Thr to yield Dhb acid. None of the *ppp* standalone genes are predicted to catalyze dehydration. However, phylogenetic analysis of the corresponding C domain of module 3 shows that it clades together with C domains that likewise process dehydroamino acids in the biosynthesis of mycrocystin and nodularin from cyanobacteria (Fig. S101, Table S22). The C domain of MycA (module 2) has been hypothesized to play a role in dehydration of Ser and Thr leading to microcystins containing dehydroalanine (Dha) and Dhb, respectively^42–45^. Thus, although experimental evidence is outstanding, the *ppp* NRPS appears to contain three C domains that putatively catalyze reactions other than simple amino acid condensation, i.e. reversal of chain polarity via an ureido linkage, dehydration, and cyclization (Fig. 6).

Another intriguing and unique feature of pseudovibriamides **2, 4, 11**, and **12** is the presence of an imidazolidinyl-dione ring. While this heterocycle is commonly found in marine sponge and other alkaloids (Table S23), to the best of our knowledge, it has not been reported as a bacterial peptide residue. Additional studies are required to unveil how this cyclization proceeds.

The structure of pseudovibriamide B is reminiscent of depsipeptide rimosamides from *Streptomyces rimosus*^21^. The rimosamide BGC likewise contains a NRPS-PKS system with two TE domains in which RmoI encodes a trimodular NRPS-PKS that loads Val/Ile, Pro and malonate, respectively, analogously to PppD (Fig. 6). Both clusters also encode putative iron/α-ketoglutarate-dependent dioxygenases (*pppK* and *rmoL*) which may be responsible for hydroxylation of Pro. The enzyme responsible for alkylation of Hyp has not been identified in either case. The *ppp* gene cluster of *P. brasiliensis* Ab134 and the short version of the *Pseudomonas ppp* BGC both contain a gene with sequence similarity to *N*-acetyltransferase family proteins as candidate to catalyze alkyl transfer.

Our results show that heptapeptides such as **1** (*m/z* 844 in Fig. 3b) are excreted to promote flagellar motility while reducing biofilm formation (Fig. 5), whereas nonadepsipeptides such as **7** and **8**/**9** (*m/z* 1156 and 1170 in Fig. 3b) are cell-associated. It is unclear whether the heptapeptides are shunt products of module 7 or hydrolysis products of the nonadepsipeptides (Fig. 6).

In conclusion, collective behaviors such as swarming motility and biofilm formation are beneficial for bacterial survival and for the colonization of eukaryotic hosts^2,5^. The majority of nonribosomal peptides shown to be implicated in swarming motility and biofilms are lipopeptides having surfactant activity ^e.g. 46,47^. In contrast, pseudovibriamides have distinct structures and are not surfactants. Identification of pseudovibriamides and their link to motility and biofilm formation opens the door to future studies on how they mediate these processes, ultimately expanding current knowledge of bacterial collective behavior.

## Materials and Methods

### Chemicals and general experimental procedures

All chemicals were acquired from Sigma-Aldrich, Alfa Aesar, VWR, and Thermo Fisher Scientific, unless otherwise noted. Solvents were of HPLC grade or higher. Restriction enzymes were purchased from New England Biolabs. Oligonucleotide primers were synthesized by Sigma-Aldrich (see Table S1 for sequences). Molecular biology procedures were carried out according to the manufactures’ instructions (New England Biolabs, Thermo Scientific, Promega, Sigma-Aldrich, Qiagen, Zymo Research, Illumina, Oxford Nanopore) unless otherwise noted.

### Cultivation conditions

*P. brasiliensis* Ab134 was routinely cultivated at 30 ^°^C on Difco™ Marine Broth 2216 (MB) or on Difco™ Marine Agar 2216 (MA). For selection of *pppA::neo* and *pppD::neo* mutants, the medium was supplemented with kanamycin at 200 μg/mL. Motility and biofilm assays were performed without addition of kanamycin. Cultures for metabolite analyses were also obtained without addition of kanamycin. *E. coli* DH5α(λpir) used for propagation of and cloning with pDS132-based vectors^22^ and *E. coli* SM10(λpir) used for conjugation were routinely cultured in LB medium at 37 ^°^C unless otherwise stated. Chloramphenicol (25 μg/mL) and kanamycin (50 μg/mL) were used for selection of *E. coli*. All strains were cryo-preserved in 20% glycerol [*v/v*] at -80 ^°^C.

### Genome sequencing, assembly, and PCR confirmation of plasmids

*P. brasiliensis* Ab134 was isolated from specimens of the marine sponge *A. brasiliensis* collected at approximately 10 m depth by scuba diving, at João Fernandinho Beach, Búzios, state of Rio de Janeiro (22^°^24’49’’ S 41^°^52’54’’ W) in January 2011^16^. A draft genome of strain Ab134 obtained using Illumina sequencing was previously reported (GenBank accession number: MIEL00000000)^17^. In order to generate a complete genome, we repeated short-read sequencing with Illumina and also obtained long-read Nanopore data. Genomic DNA was isolated from cells grown in Difco MB 2216 overnight at 30 °C using Qiagen’s Blood & Cell Culture DNA Midi Kit. Illumina and Oxford Nanopore sequencing was performed by the Sequencing Core of the University of Illinois at Chicago as previously described^48^. Hybrid assembly performed with Spades^49^ as previously described^48^ led to seven contigs.

The presence of four plasmids was confirmed by PCR (Table S2 and Fig. S1). The PCR reactions (25 μL) were carried out according to manufacturer’s instructions using Phusion High-Fidelity DNA Polymerase (Thermo Scientific) and primers described in Tables S1-S2. Genomic DNA isolated from a 5-mL marine broth overnight culture was extracted with the GenElute™ Bacterial Genomic DNA Kit (Sigma-Aldrich) and used as template. Each reaction was placed using 50 ng of gDNA, 0.2 mM of each dNTPs, 0.2 μM of each primer, 5 μL of 5× HF Phusion buffer, 1U of Phusion™ DNA polymerase and 3% of DMSO; water was added up to the total volume. The thermal cycling conditions were: 60 s at 98 ^°^C; 30 cycles of 98 ^°^C for 10 s, T_a_ ^°^C for 30 s and 72 ^°^C for 60 s; and a terminal hold for 5 min at 72 ^°^C. Primer pairs and T_a_ used in each reaction are described in Table S2. From the three remaining contigs, NODE1 and NODE4, presented a 104 bp overlap in a repetitive region of the genome (containing CTATCGAC repeats), and were considered as part of the chromosome. Different sets of primers were tested (Table S2) in order to close two remaining gaps (one in the chromosome and one in plasmid 3) but with no success. The fifth plasmid was proposed based on the presence of a replication gene, leading to one chromosome and five plasmids, and a total genome size of 5.8 Mb (Fig. 2 and Table S3). The genome sequence has been deposited in GenBank under accession code NNNN [place holder].

### Bioinformatics analyses

The genome was annotated using RAST^50^. Biosynthetic gene clusters were predicted with antiSMASH^18^ followed by manual curation and further annotation using BLAST^51^. Although the *tda* gene cluster was not predicted by antiSMASH, we were able to identify it by Blasting the genome against known genes (NCBI GenBank accession number CP002977.1, Table S4). Identification of homologous gene clusters was performed using MultiGeneBlast^20^. To construct the C domain phylogenetic tree (Fig. S99-S100), the antiSMASH results were uploaded into Geneious Prime and the condensation domains were extracted. The C-domain amino acid sequences were aligned using MUSCLE 3.8.425 (plugin inside Geneious software) with the standard parameters. The tree (Fig. S99-S100) was built using the neighbor-joining algorithm implemented in the Geneious Prime software. The *ppp* BGC was deposited in MiBIG under accession code NNNN [place holder].

### Plasmid construction

*P. brasiliensis* Ab134 genomic DNA was isolated using GenElute™ Bacterial Genomic DNA Kit (Sigma-Aldrich) and used as template for amplification of ∼700-bp homology arms (construction of pJD01 and pLI01) and a 9850-bp fragment containing *pppA* (pYD02). Phusion high-fidelity DNA polymerase (Thermo Scientific) was used for PCR amplification.

*Construction of pJD01 used to inactivate pppA*. Primer pairs P1_XbaI_NRPSup_f / P2_NRPSup_R and P5_NRPSdown_f / P6_NRPSdown_XbaI_r were used to amplify homology arms upstream and downstream of *pppA*, respectively, from *P. brasiliensis* Ab134 genomic DNA. Primer pair P3NRPSup_neo_f / P4_NRPSdown_neo_r was used to amplify the kanamycin-resistance marker from pUCPneo^52^. The three obtained fragments were assembled using splicing by overlap extension-PCR (SOE-PCR) as follows. The extension reaction (50 μL) consisted of equimolar concentrations of each template (about 100 ng total), 0.2 mM of each dNTP, 3% of DMSO and 0.02 U/μL Phusion high-fidelity DNA polymerase (Thermo Fisher Scientific) in 1× HF Phusion buffer supplied with enzyme and using the following thermal cycling parameters: 60 s at 98 ^°^C, 5 cycles of 98 ^°^C for 10 s, 69 ^°^C for 30 s and 72 ^°^C for 90 s; and a final hold at 10 ^°^C. Before the amplification step, the primer pair P1_XbaI_NRPSup_f / P6_NRPSdown_XbaI_r was added to the PCR tubes and the following extension program was performed: 60 s at 98 ^°^C; 30 cycles of 98 ^°^C for 10 s, 62 ^°^C for 30 s and 72 ^°^C for 90 s; and a terminal hold for 7 min at 72 ^°^C. Primers P1_XbaI_NRPSup_f and P6_NRPSdown_XbaI_r introduced the restrictions sites XbaI into the final PCR product. The PCR product was digested with XbaI, ligated into the same site of pDS132^22^ and transformed into *E. coli* DH5α(λpir) to yield pJD01.

*Construction of pLI01 used to inactivate pppD*. pLI01 was constructed as described for pJD01 with minor modifications. Primers pairs oLI1/oLI2 and oLI3/oLI4 were used to amplify the homology arms upstream and downstream of *pppD*, respectively. Primer pair P_neo_f/P_neo_r was used to amplify *neo* from pUCPneo. The PCR reaction (50 μL) was as for pDJ01, except that the annealing temperatures were 70 ^°^C for the extension step, and 58 ^°^C for the amplification step. Primers oLI1 and oLI4 introduced the restrictions sites XbaI into the final PCR product. The PCR product was digested with XbaI, ligated into the same site of pDS132^22^ and transformed into *E. coli* DH5α(λpir) to yield pLI01.

*Construction of pYD02 used to genetically complement the pppA mutant (pppA knockin)*. The Primer pair P1_SpeI_NRPSup_f and P6_ NRPSdown_SpeI_r was used to amplify *pppA* and its upstream and downstream homology arms from *P. brasiliensis* Ab134 genomic DNA. The PCR amplification (50 μL) consisted of 1 μL *P. brasiliensis* Ab134 genomic DNA (∼100 ng), 0.2 mM of each dNTP, 3% of DMSO, 100 nM (each) primer, and 0.02 U Phusion high-fidelity DNA polymerase (Thermo Fisher Scientific) in 1× HF Phusion buffer supplied with enzyme. The following thermal cycling parameters were used: 1 min at 98 ^°^C for initial denaturation, 30 cycles of 10 s at 98 ^°^C for denaturation, 30 s at 61 ^°^C for annealing and 5 mins at 72 ^°^C for extension; 10 mins at 72 ^°^C for final extension; and a final hold at 10 ^°^C. Primers P1_SpeI_NRPSup_f and P6_NRPSdown_SpeI_r introduced the restriction site SpeI into the final PCR product. The PCR product was then digested with SpeI, and ligated into the XbaI site of pDS132^22^ to yield pYD002 after transformation into *E. coli* DH5α(λpir).

### Gene replacement in *P. brasiliensis* Ab134

The *pppA* and *pppD* genes were inactivated by replacement with a kanamycin-resistance marker via homologous recombination after introducing pJD01 or pLI01, respectively, into *P. brasilensis* Ab134 via conjugation from *E. coli* SM10(λpir). For conjugation, strain Ab134 was cultured in MB and *E. coli* SM10(λpir) with either pJD01 or pLI01 was cultured in LB containing kanamycin (50 μg/mL) and chloramphenicol (25 μg/mL). Both Ab134 and *E. coli* strains were cultured at 30°C, 200 rpm, to an OD_600_ of 0.4 - 0.6. Cells from 20 mL cultures were harvest by centrifugation at 4000 rpm and 4 ^°^C. Ab134 cells were resuspended into 2 mL of MB. *E. coli* cells were washed twice with 20 mL LB to remove antibiotics prior to resuspension in 2 ml LB. 0.50 mL of each culture (Ab134 and *E. coli*) were carefully mixed in a 1.5-mL tube and plated onto two MA plates (0.5 mL of the mixture per plate). Negative control plates were prepared by plating only *E. coli* (0.25 mL) or only Ab134 (0.25 mL) onto marine agar. Plates were incubated at 30 ^°^C for 16-20 hours. A loopfull (10 µL loop) of cells were then streaked onto MA containing kanamycin (200 μg/mL) to select for the incoming plasmid and carbenicillin (50 μg/mL) to selectively kill *E. coli*. Obtained clones were streaked again onto selective plates and single colonies were analyzed by PCR for single crossover (SCO) using primer pairs TP_ctg3-43_f/P_neo_r (*pppA* SCO up),TP_ctg3-43_r/P_neo_f (*pppA* SCO down); oLI23/P_neo_r (*pppD* SCO up) and oLI24/P_neo_f (*pppD* SCO down). Once single crossovers were confirmed, they were streaked onto non-selective MA plates and grown for two days at 30 ^°^C. A loopful of cells were then streaked for single colonies onto MA containing 5% sucrose (the pDS132 vector has a *sacB* counterselection marker) and incubated for two days at 30 ^°^C. Obtained colonies were replica plated onto MA containing either kanamycin 200 μg/mL or chloramphenicol 2 μg/mL. Kanamycin-resistant, chloramphenicol-sensitive clones were analyzed by colony PCR to confirm the gene replacement using primers pairs TP_f/TP_r for *pppA*::*neo* and oLI23/oLI24 for *pppD*::*neo* in reactions (25 μL) containing 0.2 mM each dNTP, 3% DMSO, 0.2 μM (each) primer, and 1.25 U DreamTaq Green DNA polymerase (Thermo Fisher Scientific) in the buffer supplied with enzyme. Thermocycling conditions were initial denaturation for 3 min at 94 ^°^C; amplification for 30 cycles (94 ^°^C for 30 s, 49 ^°^C (*pppA*::*neo*)/53 ^°^C (*pppD*::*neo*) for 30 s, 72 ^°^C for 10 min; and a final extension for 12 min at 72 ^°^C. For further confirmation of mutants, the PCR reaction described above was repeated using genomic DNA isolated using GenElute™ Bacterial Genomic DNA Kit (Sigma-Aldrich).

Genetic complementation of the *pppA* mutant was achieved by replacing the *neo* marker with *pppA* via homologous recombination. Plasmid pYD002 was introduced into the *pppA*^*-*^mutant via conjugation from *E. coli* SM10(λpir). Conjugation was carried out as described above, after which cells were streaked onto MA containing chloramphenicol (8 μg/mL) to select for the incoming plasmid and carbenicillin (50 μg/mL) to selectively kill *E. coli*. Single colonies were analyzed by PCR for single crossover events using primer pairs TP_ctg3-43_f/ TP_SCO_pppA_r and TP_ctg3-43_r/ TP_SCO_pppA_f. Once single crossover events were confirmed, clones were streaked onto non-selective MA plates and grown for two days at 30 ^°^C. A loopful of cells were then streaked for single colonies onto MA containing 5% sucrose and incubated for two days at 30 ^°^C. Obtained colonies were replica plated onto MA containing either kanamycin 100 μg/mL, chloramphenicol 8 μg/mL, or no antibiotics. The genomic DNA of kanamycin-sensitive and chloramphenicol-sensitive clones was isolated using the GenElute™ Bacterial Genomic DNA Kit (Sigma-Aldrich), and analyzed by PCR to confirm the gene replacement using primers pairs TP_ctg3-43_f / TP_ctg3-43_r in reactions (20 μL) containing 0.25 mM each dNTP, 5% DMSO, 250 nM (each) primer, and 0.02 U Phusion high-fidelity DNA polymerase (Thermo Fisher Scientific) in 1× HF buffer supplied with enzyme. Thermocycling conditions were initial denaturation for 30 s at 98 ^°^C; amplification for 30 cycles: 98 ^°^C for 10 s, 62.6 ^°^C for 30 s, 72 ^°^C for 5 mins; a final extension for 10 min at 72 ^°^C; and a final hold at 10 ^°^C.

### Flagellar motility assays

Flagellar motility assays were established following previously reported, general guidelines^1^.

#### Swarming assay

MB was prepared according to manufacturer’s instructions and complemented with 0.5% of Eiken agar (Eiken Chemical CO., Japan). Ab134 does not swarm on other agars such as Difco because it appears to require the lower surface tension provided by Eiken agar (Fig. S94). Moreover, a concentration of 0.5% Eiken agar seems ideal for observation of the swarming phenotype, as the rate of swarming decreases with increasing agar concentrations (Fig. S95). After autoclaving, the agar medium was equilibrated in a water bath set to 50 ^°^C for 2h before pouring the plates (20 mL), which were then let dry open under a biosafety laminar flow hood for 20 min immediately before inoculation. Cryo stocks of Ab134 wild-type and mutant cultures normalized to OD_600_ of 1.0 were prepared. To perform the swarming assays, 5 μL of cryo stock was carefully dropped on the center of the plate, let dry for 15 min and incubated at 30 °C. A tray containing water was placed at the bottom of the incubator to control humidity. Pictures were taken daily. The swarming phenotype is apparent after two days when using cryo stocks for inoculation.

#### Swimming assay

MB was prepared as above with the exception that Eiken agar was added to a final concentration of 0.3% (w/v). 1 μL of a cryo stock normalized to OD_600_ of 1.0 was carefully inoculated inside the agar by avoiding touching the bottom of the plate. Plates were incubated as above.

### Biofilm assay

Biofilm assays were carried out using a microtiter plate method as previously described^30^ with minor modifications. 1 mL MB contained in 48-well plates, was inoculated with 20 μL seed culture. Seed cultures were prepared by growing each strain to OD_600_ of 1.0 on MB at 30 °C. Cells were harvested by centrifugation and washed twice with artificial seawater (Instant Ocean). Artificial seawater was used as seed for blanks. Biofilm formation was monitored after 24h and 48h using 0.1% crystal violet (Sigma-Aldrich) as described^30^. Statistical analysis were performed using OriginPro 8.5 (Tukey test, 95% confidence level).

### Drop-collapsing assay

Drop-collapsing assays were carried out as described before^53^, with minor modifications. A 20 μL water droplet was placed on a hydrophobic surface (Parafilm ‘M’), followed by the addition of 5 μL of bacterial cells suspension from a 5 mL MB overnight culture. The OD_600_ of the cells suspension was set to 0.25. Water was used as negative control and a solution of 0.1% Tween 20 as positive control (Fig. S97a). The assay was performed in triplicate. A second assay was performed to compare the surface tension of Difco and Eiken agar. For this assay, a 5-μL water droplet was placed on either MA (containing Difco agar) or MB + 1.2% Eiken agar (Fig. S94). For visualization purposes 0.002% of crystal violet were added to the droplet (no influence was observed on the shape of the droplet).

### Oil spreading assay

The oil spreading assay was performed according to Youssef et al.^31^ with some modifications. Petri dishes were rinsed with DI water and filled with 25 mL DI water. 10 µL of crude oil was added to the surface of the water. 10 µL of each sample was then dropped on the surface of the oil, and the diameter of the cleared zone was measured with a ruler. Samples consisted of supernatants from overnight bacterial cultures. *P. brasiliensis* Ab134 wild-type and *pppA*^-^mutant were cultured in marine broth. *Bacillus subtilis* strains used as control were cultivated in LB medium at 37 °C (Fig. S97b).

### MALDI-ToF MS analyses

Measurements were performed using an Autoflex Speed LRF mass spectrometer (Bruker Daltonics) equipped with a smartbeam™-II laser (355 nm). Dried-droplet analyses were performed routinely in positive reflectron mode (1000 shots; RepRate: 2000 Hz; delay: 9297 ns; ion source 1 voltage: 19 kV; ion source 2 voltage: 16.55 kV; lens voltage: 8.3 kV; mass range: 400 to 3,500 Da). External calibration was made with a Bruker Daltonics peptide calibration standard. Data was acquired by flexControl software v. 3.4.135.0 (Bruker Daltonics) and analyzed using flexAnalysis software v. 3.4 or flexImaging software v. 4.0.

#### Dried-droplet (DD) analysis

Samples were diluted in MeOH and mixed in 1:1 or 1:2 ratio with universal matrix and 1 μL of the mixture was applied as a thin film onto a MALDI ground-steel target plate (Bruker Daltonics, Billerica, MA). Universal matrix composition: mixture of 1:1 of α-cyano-4-hydroxycinnamic acid (CHCA) and 2,5-dihydroxybenzoic acid (2,5-DHB) (Sigma-Aldrich solubilized in 78:22 MeCN in H_2_O + 0.1% TFA.

#### Imaging mass spectrometry (IMS) analysis

Imaging mass spectrometry analyses were carried out as described before^54^. *P. brasiliensis* Ab134 was cultured on a thin layer of MA (10 mL in 9 cm plate) at 30^°^C, for 2 days. The region of interest was excised from the plate and transferred to a MALDI MSP 96 anchor plate (Bruker Daltonics, Billerica, MA). A photograph was taken. A 1:1 mixture of recrystallized CHCA/DHB was applied using a 53-µm stainless steel sieve (Hogentogler & Co., Inc., Columbia, MD). The plate was then transferred to an oven and desiccated at 37°C over 5h. Following adherence to the MALDI plate, another photographic image was taken and the sample was subjected to MALDI-TOF mass spectrometry (Autoflex; Bruker Daltonics, Billerica, MA) for IMS acquisition. Data were acquired in positive reflectron mode, with a 500-µm laser interval in the XY and a mass range of 400 to 3,500 Da.

### HPLC-PDA-ELSD-ESIMS analysis

Routine HPLC-PDA-ELSD-ESIMS analyses were performed on a Waters chromatography system consisting of a Waters 2695 Alliance control system coupled to a Waters 2696 UV-Visible spectrophotometric detector with photodiode array detector (PDA, λ_máx_ 200 – 800 nm), connected to a Waters 2424 evaporative light scattering detector (ELSD) and Waters Micromass ZQ2000 mass spectrometry (MS) detector, operated by an Empower 2.0 platform. The ELSD detector was optimized to the following parameters: gain of 100, gas pression (N_2_) of 50 psi, drift tube temperature of 80 ^°^C and nebulizer temperature of 36 ^°^C, 60 %. The MS detector was optimized to the following parameters: capillary voltage of 3.0 kV, source temperature of 100 ^°^C, desolvation temperature of 350 ^°^C, operating on electrospray ionization (ESI) in positive and negative mode, (300 – 1500 Da), cone gas (N_2_) flow of 50 L/h and desolvation flow of 350 L/h. Quick fractionation analysis were performed using a Water C_18_ X-Terra reversed phase column (2.1 x 50 mm, 3.5 μm), with H_2_O (A) and MeOH/MeCN 1:1 (B), both acidified with 0.1% formic acid, as mobile phases, eluted with a gradient of 5-50% of B in 5 min, followed 50-100% in 2 min. Analyses of final fractions were performed using a Water C_18_ X-Terra reversed phase column (4.6 x 250 mm, 5 μm), using the same solvent system as above, in a gradient of 5-50% of B in 22 min, followed by 50-100% in 7 min.

### UPLC-HRESIMS and MS/MS analysis

UPLC-HRESIMS and MS/MS analyses were performed on a Waters Acquity UPLC H-class system coupled to a Waters Xevo G2-XS QToF mass spectrometer (MS) with electrospray ionization (ESI), using data-dependent acquisition in positive mode with the following parameters: capillary voltage of 1.2 kV, cone voltage of 30 V, source temperature of 100 °C, desolvation temperature of 450 °C, cone gas (N_2_) flow of 50 L/h, desolvation gas (N_2_) flow of 750 L/h, scanning range from 200 to 1500 Da, scan time of 0.2 s, collision energy ramp of 30-50 V. Internal calibration was accomplished with a solution of leucine enkephalin (20 μg/ml, Sigma-Aldrich^®^) infused by the lock-mass probe at a flow rate of 10 μL/min. UPLC was performed at 0.5 mL/min through a BEH C_18_ (2.1 × 100 mm × 1.7 μm, Waters) column at 40 °C. Samples were kept at 15 °C. Mobile phases were H_2_O (A) and CH_3_CN (B), both acidified with 0.1% formic acid in a gradient of 5-30% of B in 6 min, following 30-100% B in 2 min.

### Evaluation of culture conditions for isolation

Four culture conditions and extraction procedures were tested in order to obtain the products of the *ppp* BGC. *P. brasiliensis* Ab134 was cultivated on: 1) 50 mL of MB in a 250-ml Erlenmeyer flask, and the whole culture (cells and broth) was extracted with AcOEt; 2) 50 mL MB + 10% (w/v) of a 1:1:1 mixture of macroporous adsorptive resins XAD-2, -4 and -7HP, followed by a MeOH extraction of the combined resins; 3) 50 mL of MB + two cotton balls (cells grow attached to the cotton), followed by MeOH extraction of the cotton balls as described before^55^; and 4) agar plates with 20 mL MB + 0.5% Eiken agar, followed by extraction of the complete content of the plates by sonication with MeOH. All cultures were grown for 4 days, at 30 ^°^C. Growth experiments on liquid media were performed under 160 rpm. All extracts were evaluated by MALDI-TOF MS (Fig. S6). According to the TIC for pseudovibriamide signals, cultivation on solid media (swarming assay conditions) led by far to the highest amounts of pseudovibriamides, followed by cultivation using cotton, liquid cultivations without resin, and resin addition at the time of inoculation.

### Isolation of pseudovibriamides and NMR analysis

*P. brasiliensis* Ab134 was cultivated on a total of 2,000 agar plates each containing 20 mL MB with 0.5% Eiken agar (total 40 L) for 3 days at 30 ^°^C. A 4-μL drop of a cryo stock (normalized to OD_600_ 1.0) was inoculated in the middle of each plate as done for a typical swarming assay. Three different extracts were generated: F, FD and C. Extract F (“F” stands for filtrate extract) was obtained from the total contents (cells and media) of 400 plates. Agar medium and cells were first frozen at -80 °C. After defrosting (twice), the mixture was centrifuged at 4000 rpm, 17 ^°^C for 30 min. The liquid was separated by filtration and a mixture of resins (XAD-2, -4, -7HP and -16 1:1:1:1) was added. Following agitation overnight, the mixture of resins was recovered, washed twice with H_2_O and extracted with MeOH (twice) and acetone (once). The organic solvents were pooled and evaporated, yielding the crude extract F (4.9 g). Extract FD (“FD” stands for freeze-<u>d</u>ried extract) was obtained from the total contents of 600 plates (cells and media). Agar medium and cells were freeze-dried and then extracted twice with MeOH by sonication at 40 kHz for 30 min. The MeOH extract was recovered by filtration and dried under reduced pressure yielding the crude extract FD (50.9 g). Extract C (“C” stands for <u>c</u>ells only) was obtained from 1000 plates, from which only the cells were recovered using a spatula (Fig. S49). The cell mass was frozen overnight and after defrosting, extracted twice with MeOH by sonication at 40 kHz for 30 min. The extract was recovery by filtration and evaporated, yielding 11.2 g of C extract. All the following fractionations steps were MS guided. Extract F (4.9 g, fractionation steps are depicted on Fig. S102) was solubilized in MeOH and centrifuged to remove insoluble parts, and the liquid was adsorbed into 2 g of Diaion HP-20SS and subjected to chromatography on Diaion HP-20SS (10 g) column, eluted with a gradient of IPA in H_2_O (details on Fig. S102, step 1), to give four fractions. Fraction F2 (556.8 mg), was subjected to chromatography on a C18 column (Supelco Discovery, 5g), eluted with a gradient of MeOH in H_2_O (details on Fig. S102, step 2), to give three fractions. Fraction F2B (415.7 mg) was further fractionated by reversed-phase semi preparative HPLC using a Phenomenex Onyx Monolithic C18 column (100 x 4.6 mm), 4 mL/min flow rate, and automatic fraction collection every 30s (from 0.5-12.5 min). The method consisted of a gradient of 10-30% MeOH/MeCN 1:1 in H_2_O with 0.1% formic acid (FA) over 10 min, a 3 min wash with 100% MeOH/MeCN 1:1, and re-equilibration for 2 min (Fig. S102, step 3). The 24 fractions were combined into 12 fractions after HPLC analysis. Fraction F2B67 (48.8 mg) was fractionated by HPLC using a reversed-phase Phenomenex Kinetex C18 column (250 x 10 mm, 100 Å), 4 mL/min flow rate, eluted with a isocratic of 19% MeOH/MeCN 1:1 in H_2_O + 0.1% FA over 30 min, to give five fractions (Fig. S102, step 4). Fraction F2B67D (14.2 mg) was further fractionated by HPLC using a Phenomenex Luna HILIC column (150 x 4.6 mm, 5 μm), 1 mL/min flow rate, eluted with a isocratic of 9:1 MeCN/H_2_O (both + 0.1% FA) over 15 min, yielding 5.5 mg of **1** and 2.7 mg of **2** (Fig. S102, step 5). 1D and 2D NMR spectra of **1** were obtained at 25^°^C on a Bruker Avance AVII 900 MHz spectrometer equipped with a 5 mm CPTCI probe, using chemical shifts of DMSO-*d*_6_ as reference. The NMR sample was prepared in a 3 mm Shigemi microtube assembly matched for DMSO.

Extract FD (50.5 g, fractionation steps are depicted on Fig. S103) was solubilized in MeOH and centrifuged to remove insoluble parts (Fig. S103, step 1). The solvent was evaporated, and the extract was subjected to chromatography on a C18 column (Stracta C18-E, 55 μm, 70Å, 10 g), eluted with a gradient of MeOH in H_2_O (both + 0.1% FA), to give five fractions (details on Fig. S103, step 2). Fraction FD2 (2.9 g) was subjected to chromatography on a Phenyl column (Stracta Ph, 55 μm, 70Å, 10 g), eluted with a gradient of MeOH in H_2_O (both + 0.1% FA), to give nine fractions (details on Fig. S103, step 3). Fraction FD2B (690.0 mg) was further fractionated by size-exclusion chromatography on a column of Biogel-P2 (103 x2 cm), eluted with 4:1 H_2_O/MeOH (both + 0.5% FA), coupled with an automatic fraction collector. 200 fractions (4.5 mL each) were collected and combined into 5 fractions after HPLC analysis (Fig. S103, step 4). Fraction FD2B3 (177.3 mg) was fractionated by HPLC using a reversed-phase Phenomenex Kinetex PFP column (250 x 4.6 mm, 5 μm, 100 Å), 1 mL/min flow rate, eluted with a isocratic of 25% MeOH/MeCN 11:14 in H_2_O + 0.1% FA over 40 min, to give five fractions (Fig. S103, step 5). Fraction FD3B3A (4.4 mg) was further fractionated by HPLC using a reversed-phase Inertsil Ph-3 column (250 x 4.6 mm, 5 μm, GL Sciences), eluted with a isocratic of 25% MeOH/MeCN 3:2 in H_2_O (all + 0.1% TFA), to afford 1.0 mg of **3** (Fig. S103, step 6). Fraction FD3B3D (6.5 mg) was fractionated using the same column and elution to afford extra 2.0 mg of **2**; while fractionation of FD3B3C (2.8 mg) afforded 0.2 mg of **4** (Fig. S103, step 6). Fraction FD2B2 (66.2 mg) was partially (Fig. S103, step 7) fractionated by HPLC using a reversed-phase Inertsil Ph-3 column (250 x 10 mm, 5 μm, GL Sciences), eluted with a isocratic of 26% MeOH/MeCN 15:11 in H_2_O (all + 0.1% TFA) to give 0.5 mg of **5** and **6**.

After combining samples of **2** from both growth experiments, 1D and 2D NMR spectra were obtained at 25^°^C on a Bruker Advance III 600 MHz spectrometer.

Extract C (11.2 g, fractionation steps are depicted on Fig. S104) was fractionated as extract FD for steps 2 to 4 (2: C18 → 3: Ph → 4: Biogel-P2). In step 5, fraction C23C1 (33.1 mg) was fractionated by HPLC using a reversed-phase Inertsil Ph-3 column (250 x 10 mm, 5 μm, GL Sciences), eluted with isocratic 29% MeOH/MeCN 17:12 in H_2_O (all + 0.1% TFA), to afford 5.9 mg of **7**, together with 6.2 mg mixture of **8** and **9**, as well as, 2.7 mg of **10** and 1.4 mg mixture of **11** and **12** (Fig. S104). 1D and 2D NMR of **7** spectra were obtained at 25^°^C on a Bruker Advance III 600 MHz spectrometer.

### Advanced Marfey’s analysis

Briefly, *c*.*a*. 100 μg of each compound were solubilized into 1 mL of 4 mol/L HCl and heated at 100 ^°^C for 6h, with stirring. The solution was cooled to room temperature and dried under vacuum. The hydrolysate was resuspended in 1 mL of H_2_O and dried three times. Marfey’s derivatization was performed using 1% 5-fluoro-2,4-dinitrophenyl-Nα-L-tryptophanamide (FDTA) in acetone, as described before^24,25^. The hydrolysate was resuspended in H_2_O, followed by the addition of 20 μL of 1 mol/L NaHCO_3_, and L-FDTA at 1.4x of the amino acid molar concentration of the compound hydrolysate. The mixture was stirred and heated at 40 ^°^C during 1h. The reaction was then cooled to room temperature and neutralized with 10 μL of 2 mol/L HCl. The amino acid standard were derivatized using both L- and D-FDTA. Samples were analyzed by UPLC-HRESIMS using column and conditions as described before^25^.

## Acknowledgements

We thank Dr. Fabiano Thompson (Federal University of Rio de Janeiro) for the *Pseudovibrio brasiliensis* strain Ab134 and for sharing genome data prior to publication, Dr. D. Schneider (Grenoble Alpes University) for the pDS132 vector, UIC’s Sequencing Core for genome sequencing, Dr. G. Pauli (UIC) for access to qTOF-MS used in preliminary studies, and Dr. T. Molinski (University of California San Diego) for the FDTA Marfey’s reagent. Genome sequencing and assembly was performed by the UIC Research Resources Center, supported in part by NCATS through Grant UL1TR002003. Financial support for this work was provided by the Division of Integrative Organismal Systems of the National Science Foundation under grant IOS-1917492 (to ASE), by the São Paulo Research Foundation FAPESP grants 2013/50228-8 and 2015/01017-0 (to RGSB), by FAPESP scholarships 2016/05133-7 and 2018/10742-8 (to LPI), National Institutes of Health National Institute of General Medical Sciences R01GM125943 (to LMS) and by startup funds from the Department of Pharmaceutical Sciences (to ASE and LMS) and the Center for Biomolecular Sciences, University of Illinois at Chicago (to ASE).

## Author contributions

JDE established reverse genetics for *P. brasiliensis* Ab134 under ASE mentorship and obtained *pppA* mutants. ASE isolated high MW DNA for sequencing, and performed initial motility assays. SK performed initial mass spectrometry and imaging analyses. CMC performed initial MS/MS analysis and identified the two related compound groups. JO advised CMC. LPI generated *pppD* mutants, performed motility and biofilm assays, performed imaging mass spectrometry experiments, and closed the genome. LPI also isolated pseudovibriamides and determined their structures with help from AK and RGSB. YD performed genetic complementation experiments. AK and AGF performed NMR experiments. AK helped analyze NMR data and proposed the ureido functionality. LS advised imaging mass spectrometry experiments. RGSB contributed with isolation and structure elucidation, mentoring of LPI, laboratory facilities and funding. ASE devised the project, contributed with overall experimental design, advised LPI, JDE and SK, and contributed with laboratory facilities and funding. LPI and ASE wrote the paper draft. All authors commented on, edited and approved the manuscript.

