## Supporting Information for "A novel family of nonribosomal peptides modulate collective behavior in *Pseudovibrio* bacteria isolated from marine sponges"

### Table of contents

**List of Supplementary Tables:**

**Table S1.** Oligonucleotide primers used in this study. 8
**Table S2.** Primer pairs used to amplify *P. brasiliensis* Ab134 genome gaps or to confirm the presence of plasmids.. 9
**Table S3.** Details of final genome assembly of *P. brasiliensis* Ab134. 10
**Table S4.** Gene clusters found in *P. brasiliensis* Ab134's genome by antiSMASH and BLAST analyses. 10
**Table S5.** List of *Pseudovibrio* and *Pseudomonas* genomes which harbor a member of the *ppp* BGC family. 11
**Table S6.** $^{1}$H, DEPTQ $^{13}$C and $^{15}$N NMR data for **1**. 13
**Table S7.** Fragment ions observed by UPLC-QTOF-MS$^2$ analysis of **1**. 14
**Table S8.** Fragment ions observed by UPLC-QTOF-MS$^2$ analysis of **2**. 14
**Table S9.** $^{1}$H and $^{13}$C NMR data for **2**. 15
**Table S10.** Fragment ions observed by UPLC-QTOF-MS$^2$ analysis of **3**. 16
**Table S11.** Fragment ions observed by UPLC-QTOF-MS$^2$ analysis of **4**. 16
**Table S12.** Selected fragment ions observed by UPLC-QTOF-MS$^2$ analysis for **5**/**6**. 17
**Table S13.** Fragment ions observed by UPLC-QTOF-MS$^2$ analysis of sample pseudovibriamide A1 degradation (*m/z* 638.2). 18
**Table S14.** $^{1}$H and $^{13}$C NMR data for **7**. 19
**Table S15.** Fragment ions observed by UPLC-QTOF-MS$^2$ analysis of **7**. 20
**Table S16.** Fragment ions observed by UPLC-QTOF-MS$^2$ analysis of **8/9**. 21
**Table S17.** Fragment ions observed by UPLC-QTOF-MS$^2$ analysis of **10**. 21
**Table S18.** Fragment ions observed by UPLC-QTOF-MS$^2$ analysis of **11/12**. 22
**Table S19**. Absorbance measurements of biofilm assay using 0.1% crystal-violet solution. 23
**Table S20.** *Pseudovibrio* genomes which harbor a member of the *ppp* BGC family. 24
**Table S21.** Nonribosomal peptide bacterial products containing ureido-linkages. 25
**Table S22.** Bacterial products containing Dhb residues. 27
**Table S23.** Natural products containing an imidazolidinyl-dione ring. 28

**List of Supplementary Figures:**

**Figure S1.** Multireplicon confirmation by PCR. 29
**Figure S2.** Representative Multigeneblast results. 30
**Figure S3.** Domain organization of various *ppp* BGCs belonging to *Pseudovibrio* spp. and *Pseudomonas* spp.. 32
**Figure S4.** *pppA* and *pppD* gene knockout. 33
**Figure S5.** Mass spectra MS$^2$ of compounds encoded by the *ppp* BGC observed on QTOF-MS analysis of the crude extract of *P. brasiliensis* Ab134 34

**Table S1.** Oligonucleotide primers used in this study. SOE-PCR overhangs are shown in blue, restriction sites in red and underlined. Random bases inserted to allow restriction of PCR fragments are in italics.

| Primers | Sequence 5'→3' |
|---|---|
| **Closing genome gaps and testing for circular replicons** | |
| **oLI5** | GGTTGGAGTTGAGGCATTG |
| **oLI6** | GACCACATGCCTCATGAAC |
| **oLI7** | CCTTGGCATTTATCTCCTGC |
| **oLI8** | CTTCGACGGTGCTCTTTATATG |
| **oLI9** | GGAACAACTCGGGCAATC |
| **oLI10** | GCTATCGTTTACTGGAACACG |
| **oLI15** | GTAGGTTGGTTTGCTCAGCA |
| **oLI16** | GGTACAACGGATCAGTTCTG |
| **oLI17** | GCTGCTGCTTCTGGTCAG |
| **oLI18** | GTGTCTGCAATTGGGCTC |
| **oLI19** | GGAATAAGGGCACATCTGC |
| **oLI20** | CCTGAGCAAATGCTTCAAG |
| **oLI21** | GAAGAGCTGGAAGCGAAC |
| **oLI22** | CCATCAGCGCGATAAAATC |
| **oLI29** | GTCGGCTTCAGCAATGGCTC |
| **oLI30** | CAAATCTTGTTCTGGCCCCATC |
| **oLI31** | GGAGCTCGGTACTGGCATCTGG |
| ***pppA* gene inactivation** | |
| **P_neo_f** | GCAAGGGCTGCTAAAGGAAG |
| **P_neo_r** | ACGGAAATGTTGAATACTCATACTC |
| **TP_ctg3-43_f** | CAGCAAGAAAAGCTAGGTTTGAG |
| **TP_ctg3-43_r** | GTTCTGGTGCTCCGGATAG |
| **P1_XbaI_NRPSup_f** | *AACTGC*TCTAGACAAACCACACTCCGCAGAGAG |
| **P2_NRPSup_r** | CTACGTTGGAAAGACTAAACTCACC |
| **P3_NRPSup_neo_f** | TGTTTTATGGGTGAGTTTAGTCTTTCCAACGTAGGCAAGGGCTGCTAAAGGAAG |
| **P4_NRPSdown_neo_r** | GCGTCAGGAAAATTCTTCGGCTGACGGAAATGTTGAATACTCATACTC |
| **P5_NRPSdown_f** | CAGCCGAAGAATTTTCCTGAC |
| **P6_NRPSdown_XbaI_r** | *AACTGC*TCTAGAGCTGTTCCCAATAGGCCTTC |
| ***pppD* gene inactivation** | |
| **P_neo_f** | GCAAGGGCTGCTAAAGGAAG |
| **P_neo_r** | ACGGAAATGTTGAATACTCATACTC |
| **oLI1** | *AACTGC*TCTAGACACAAGACAACCTGATTACG |
| **oLI2** | GCTTCCTTTAGCAGCCCTTGCATGTGAGATAGTGCTGTTCATG |
| **oLI3** | AGGAAGAGTATGAGTATTCAACATTTCCGTGCTAGATGACGTGCTCAGT |
| **oLI4** | *AACTGC*TCTAGACTCAAGAATGGTTTCAAACTC |

| | |
|---|---|
| **oLI23** | GCTGGTGGGAGAGCTCTATATC |
| **oLI24** | CGATCAGTTAGCTCAAGAGATTG |
| ***pppA* gene complementation** | |
| **P1_SpeI_NRPSup_f** | *GG*ACTAGTCAAACCACACTCCGCAGAGAG |
| **P6_ NRPSdown_SpeI_r** | *GG*ACTAGTGCTGTTCCCAATAGGCCTTC |
| **TP_ctg3-43_f** | CAGCAAGAAAAGCTAGGTTTGAG |
| **TP_ctg3-43_r** | GTTCTGGTGCTCCGGATAG |
| **TP_SCO_pppA_r** | CCGACGTGTAGATGATATAGGC |
| **TP_SCO_pppA_f** | GGTCTTCTCAACTGGTTGCG |

**Table S2.** Primer pairs used to amplify *P. brasiliensis* Ab134 genome gaps or to confirm the presence of plasmids. For sequences see Table S1 above.

| **Amplicon (final assembly)** | **Primer pair** | **$T_a$ (°C)** | **Amplicon obtained? (bp)** |
|---|---|---|---|
| **NODE2 (plasmid 1)** | oLI15/oLI16 | 62 | ✓ (750) |
| **NODE3 (plasmid 2)** | oLI17/oLI18 | 62 | ✓ (750) |
| **NODE6 (plasmid 4)** | oLI19/oLI20 | 62 | ✓ (2000) |
| **NODE7 (plasmid 5)** | oLI21/oLI22 | 62 | ✓ (1000) |
| **NODE1+NODE4 (chromosome)** | oLI6/oLI7 | 62 | X |
| **NODE1+NODE5** | oLI6/oLI9 | 62 | X |
| **NODE4+NODE1 (chromosome)** | oLI5/oLI8 | 62 | X |
| **NODE4+NODE5** | oLI7/oLI10 | 62 | X |
| **NODE5+NODE1** | oLI8/oLI9 | 62 | X |
| **NODE5+NODE4** | oLI7/oLI10 | 62 | X |
| **NODE4** | oLI7/oLI8 | 62 | X |
| **NODE5 (plasmid 3)** | oLI9/oLI10<br>oLI29/oLI30<br>oLI9/oLI30<br>oLI29/oLI10 | 61<br>68<br>61<br>63 | X<br>X<br>X<br>X |
| **NODE1** | oLI5/oLI6 | 62 | X |

**Table S3.** Details of final genome assembly of *P. brasiliensis* Ab134.

| Replicon | Size (bp) | GC (%) | Gap | Replication protein Blastp hit (organism) | Query cover (%) | Identity (%) |
|---|---|---|---|---|---|---|
| **Chromosome** | 4,836,770 | 52.2 | 1 | *dnaA* (*Pseudovibrio* spp.) | 99.3 | 100.0 |
| **Plasmid 1** | 472,905 | 52.6 | 0 | replication protein C (*Pseudovibrio* sp. JE062) | 97.5 | 97.2 |
| **Plasmid 2** | 420,747 | 52.6 | 0 | replication protein C (*Pseudovibrio* sp. FO-BEG1) | 99.7 | 99.5 |
| **Plasmid 3** | 151,988 | 49.0 | 1 | replication protein RepCa2 (*Rhizobium leguminosarum*) | 99.7 | 43.9 |
| **Plasmid 4** | 107,145 | 49.0 | 0 | replication protein C (*P. axinellae*) | 98.6 | 35.6 |
| **Plasmid 5** | 40,855 | 48.5 | 0 | replication protein C (*Pseudovibrio* sp. Alg231) | 99.7 | 69.2 |

**Table S4.** Gene clusters found in *P. brasiliensis* Ab134's genome by antiSMASH and BLAST analyses.

| BGC | Class | Location | Size (kb) |
|---|---|---|---|
| **1** | Terpene | Chromossome | 20.8 |
| **2** | Bacteriocin | Chromossome | 9.1 |
| **3** | Thiopeptide | Chromossome | 39.9 |
| **4** | Bacteriocin | Chromossome | 10.9 |
| **5** | Terpene | Chromossome | 22.5 |
| **6** | PKS | Plasmid 2 | 63.5 |
| **7** | NRPS-PKS | Plasmid 2 | 71.8 |
| **8** | TDA | Plasmid 1 | 14.0 |

**Table S5.** List of *Pseudovibrio* and *Pseudomonas* genomes which harbor a member of the *ppp* BGC family.

| NCBI accession | Isolated from | Organism |
|---|---|---|
| **[place holder]** | marine sponge *Arenosclera brasiliensis* | *Pseudovibrio brasiliensis* Ab134 |
| **NZ_FPBD01000007** | sea water | *Pseudovibrio denitrificans* DSM17465 |
| **BAZK01000008** | sea water | *Pseudovibrio denitrificans* JCM12308 |
| **NZ_LMCB01000043** | marine sponge *Axinella dissimilis* | *Pseudovibrio axinellae* Ad2 |
| **NZ_FOFM01000016** | marine sponge *Axinella dissimilis* | *Pseudovibrio axinellae* DSM24994 |
| **NZ_FREX01000017** | marine sponge *Spongia officinalis* | *Pseudovibrio* sp. Alg231-02 |
| **NZ_LCWZ01000112** | marine sponge *Polymastia penicillus* | *Pseudovibrio* sp. POLY-S9 |
| **NZ_LMCI01000029** | marine sponge *Axinella dissimilis* | *Pseudovibrio* sp. W64 |
| **NZ_LMCJ01000042** | marine sponge *Axinella dissimilis* | *Pseudovibrio* sp. W74 |
| **NZ_LMCK01000096** | marine sponge *Axinella dissimilis* | *Pseudovibrio* sp. WM33 |
| **NZ_LMCH01000003** | marine sponge *Axinella dissimilis* | *Pseudovibrio* sp. Ad5 |
| **NZ_LMCC01000034** | marine sponge *Axinella dissimilis* | *Pseudovibrio* sp. Ad13 |
| **NZ_LMCD01000015** | marine sponge *Axinella dissimilis* | *Pseudovibrio* sp. Ad14 |
| **NZ_LMCE01000064** | marine sponge *Axinella dissimilis* | *Pseudovibrio* sp. Ad26 |
| **NZ_LMCF01000190** | marine sponge *Axinella dissimilis* | *Pseudovibrio* sp. Ad37 |
| **NZ_LMCFG01000068** | marine sponge *Axinella dissimilis* | *Pseudovibrio* sp. Ad46 |
| **NZ_FNLB01000004** | marine tunicate *Synoicum adareanum* | *Pseudovibrio* sp. Tun.PSC04-5.I4 |
| **NC_008027** | fruit fly *Drosophila melanogaster* | *Pseudomonas entomophila* str L48 |
| **LT629972** | plant pathogen of rice *Oryza sativa* | *Pseudomonas fuscovaginae* str LMG2158 |
| **LT629777** | bird's nest fern *Asplenium nidus* | *Pseudomonas asplenii* str ATCC 23835 |
| **NZ_CP029608** | garden soil | *Pseudomonas kribbensis* str 46-2 |
| **NZ_CP028826** | rhizosphere of a soybean plant | *Pseudomonas fluorescens* str MS82 |
| **NZ_LN847264** | water sample | *Pseudomonas* sp. str CCOS 191 |
| **CP022313** | plant *Leonotis nepetifolia* | *Pseudomonas fluorescens* str NEP1 |
| **NZ_CP011566** | plant root *Conyza canadensis* | *Pseudomonas* sp. str DR 5-09 |

| | | |
|---|---|---|
| **LT629788** | outdoor gate at OSU | *Pseudomonas moraviensis* str BS3668 |
| **LT629709** | aerobic zone of Elbe sediment | *Pseudomonas reinekei* str BS3776 |
| **NZ_LT963395** | plant pathogen of cherry trees | *Pseudomonas cerasi* str PL963 |
| **NZ_LT962480** | plant associated *Prunus* sp. | *Pseudomonas syringae pv. syringae* str CFBP4215 |
| **LT629769** | corn lead surface | *Pseudomonas syringae* str 31R1 |
| **NZ_CP026568** | plant associated *Prunus avium* | *Pseudomonas syringae pv. syringae* str Pss9097 |
| **NZ_LT855380** | plant pathogen of cherry trees | *Pseudomonas viridiflava* str CFBP1590 |
| **NZ_CP026558** | plant associated *Prunus avium* | *Pseudomonas amygdali pv. morsprunorum* str R15244 |
| **NC_005773** | plant pathogen *Phaseolus vulgaris* | *Pseudomonas savastanoi pv. phaseolicola* str 1448A |
| **NZ_CP020351** | plant pathogen cucumber | *Pseudomonas amygdali pv. lachrymans* str NM002 |
| **NZ_CP008742** | plant pathogen *Olea europaea* | *Pseudomonas savastanoi pv. savastanoi* str NCPPB 3335 |
| **CM001986** | plant pathogen *Lycopersicon esculentum* | *Pseudomonas syringae pv. syringae* str SM |
| **NZ_LT962481** | plant pathogen *Citrus reticulata* | *Pseudomonas syringae pv. syringae* str CFB2118 |
| **NZ_LT963391** | plant pathogen *Prumus sp.* | *Pseudomonas syringae pv. cerasicola* str CFB6109 |
| **NZ_CP013183** | plant pathogen *Triticum aestivum* | *Pseudomonas syringae pv. lapsa* str ATCC 10859 |
| **NC_007005** | plant pathogen bean | *Pseudomonas synringae pv. syringae* str B728a |
| **NZ_CP028490** | plant pathogen *Triticum aestivum* | *Pseudomonas syringae pv. atrofaciens* str LMG5095 |
| **CM001763** | plant pathogen *Panicum miliaceum* | *Pseudomonas syringae pv. syringae* str B64 |
| **NZ_CP006256** | plan pathogen *Panicum miliaceum* | *Pseudomonas syringae pv. syringae* str HS191 |
| **NZ_CP005969** | plan pathogen *Pyrus communis* | *Pseudomonas syringar pv. syringae* str B301D |

**Table S6.** $^1$H, DEPTQ $^{13}$C and $^{15}$N NMR data for **1**, fraction F2B67D3 in DMSO-$d_6$ + TFA ($^1$H: 900 MHz, $^{13}$C: 226 MHz, $^{15}$N: 91 MHz).

| Residue | Position | $\delta_C$, type | $\delta_H$ (J in Hz) | $\delta_N$, type |
|---|---|---|---|---|
| **L-Tyr** | $CO_2H$ | 173.51, CO | | |
| | NH | | 6.42, s (NH) | 88.51, NH |
| | CO | 155.16, CO | | |
| | α | 54.09, CH | 4.30 | |
| | β | 36.68, $CH_2$ | 2.83, dd (6.7, 6.5); 2.89, dd (5.2, 4.9) | |
| | aromatic | 1: 127.00, C | | |
| | | 2, 2': 130.36, CH | 6.97, d (8.0) | |
| | | 3, 3': 115.06, CH | 6.67, d (8.0) | |
| | | 4: 156.11, C | | |
| **Z-Dhb** | NH | | 7.81, s | 92.17, NH |
| | CO | 164.97, CO | | |
| | α | 131.95, C | | |
| | β | 121.88, CH | 5.95, q (6.6) | |
| | γ | 12.62, $CH_3$ | 1.59, d (6.9) | |
| **L-Ala** | NH | | 7.72, s | |
| | CO | 172.37, CO | | |
| | α | 48.32, CH | 4.29, t (7.1) | |
| | β | 17.99, $CH_3$ | 1.23, d (6.9) | |
| **L-Arg** | NH | | 8.15, d (7.1) | |
| | CO | 170.65, CO | | |
| | α | 50.37, CH | 4.50, dt (7.7, 5.5) | |
| | β | 28.10, $CH_2$ | 1.53, m; 1.76, m | |
| | γ | 24.93, $CH_2$ | 1.53, m | |
| | δ | 40.43, $CH_2$ | 3.09, m; 3.12, m | |
| | guanidine | 156.8, C | 2.53, s | |
| | | | 7.35, s | |
| | | | 7.63, s; 7.69, s | |
| **L-Pro-Thz** | α | 58.47, CH | 5.32, dd (8.0, 2.1) | |
| | β | 31.47, $CH_2$ | 2.18, m; 2.24, m | |
| | γ | 24.10, $CH_2$ | 1.97, m; 2.00, m | |
| | δ | 46.76, $CH_2$ | 3.73, m | |
| | CO | 160.53, CO | | |
| | α' | 149.12, C | | |
| | β' | 123.98, CH | 8.16, m | |
| | δ' | 173.67, C | | |
| **L-Gln** | NH | | 8.43, d (7.9) | |
| | CO | 173.22, CO | | |
| | α | 51.86, CH | 4.38, m | |
| | β | 26.36, $CH_2$ | 1.97, m; 2.11, m | |
| | δ | 31.42, $CH_2$ | 2.15, m | |
| | $CONH_2$ | 173.8, CO | 6.79, s | 110.42, $NH_2$ |

**Table S7.** Fragment ions observed by UPLC-QTOF-MS$^2$ analysis of **1**, fraction F2B67D3 (*m/z* 844.3). The fragmentation scheme based on Roepstorff-Fohlmann-Biemann fragmentation nomenclature is shown.

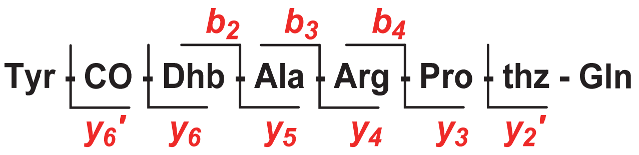

| | | *m/z* acurat. | *m/z* calc. | Error (ppm) |
|---|---|---|---|---|
| ***b*$_2$** | $C_{14}H_{15}N_2O_5^+$ | 291.0978 | 291.0981 | -1.0 |
| ***b*$_3$** | $C_{17}H_{20}N_3O_6^+$ | 362.1354 | 362.1352 | 0.6 |
| ***b*$_4$** | $C_{23}H_{32}N_7O_7^+$ | 518.2342 | 518.2363 | -4.1 |
| ***y*$_2$'** | $C_9H_{12}N_3O_4S^+$ | 258.0548 | 258.0549 | -0.4 |
| ***y*$_3$** | $C_{13}H_{19}N_4O_4S^+$ | 327.1130 | 327.1127 | 0.9 |
| ***y*$_4$** | $C_{19}H_{31}N_8O_5S^+$ | 483.2151 | 483.2138 | 2.7 |
| ***y*$_5$** | $C_{22}H_{36}N_9O_6S^+$ | 554.2515 | 554.2509 | 1.1 |
| ***y*$_6$** | $C_{26}H_{41}N_{10}O_7S^+$ | 637.2866 | 637.2880 | -2.2 |
| ***y*$_6$'** | $C_{27}H_{39}N_{10}O_8S^+$ | 663.2667 | 663.2673 | -0.9 |

**Table S8.** Fragment ions observed by UPLC-QTOF-MS$^2$ analysis of **2**, fraction F2B67D1/FD2B3D1 (*m/z* 826.3). The fragmentation scheme based on Roepstorff-Fohlmann-Biemann fragmentation nomenclature is shown.

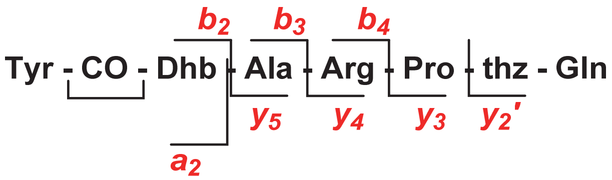

| | | *m/z* acurat. | *m/z* calc. | Error (ppm) |
|---|---|---|---|---|
| ***b*$_2$** | $C_{14}H_{13}N_2O_4^+$ | 273.0883 | 273.0875 | 2.9 |
| ***b*$_3$** | $C_{17}H_{18}N_3O_5^+$ | 344.1255 | 344.1246 | 2.6 |
| ***b*$_4$** | $C_{23}H_{30}N_7O_6^+$ | 500.2276 | 500.2258 | 3.6 |
| ***y*$_2$'** | $C_9H_{12}N_3O_4S^+$ | 258.0555 | 258.0549 | 2.3 |
| ***y*$_3$** | $C_{13}H_{19}N_4O_4S^+$ | 327.1135 | 327.1127 | 2.4 |
| ***y*$_4$** | $C_{19}H_{31}N_8O_5S^+$ | 483.2134 | 483.2138 | -0.8 |
| ***y*$_5$** | $C_{22}H_{36}N_9O_6S^+$ | 554.2501 | 554.2509 | -1.4 |
| ***a*$_2$** | $C_{13}H_{13}N_2O_3^+$ | 245.0927 | 245.0926 | 0.4 |
| ***a*$_3$** | $C_{16}H_{18}N_3O_4^+$ | 316.1306 | 316.1297 | 2.8 |
| ***a*$_4$** | $C_{22}H_{30}N_7O_5^+$ | 472.2324 | 472.2308 | 3.4 |
| ***a*$_6$** | $C_{30}H_{40}N_9O_6S^+$ | 654.2847 | 654.2822 | 3.8 |

**Table S9.** $^1$H and $^{13}$C NMR data for **2**, fraction F2B67D1/FD2B3D1 in DMSO-$d_6$ + TFA ($^1$H: 600 MHz, $^{13}$C: 150 MHz, $^1$H*: 900 MHz).

| | **Position** | **$\delta_C$ (type)** | **$\delta_H$ (J, Hz)** | **$\delta_H^*$ (J, Hz)** |
|---|---|---|---|---|
| L-Tyr | NH | | 8.44 | 8.50 |
| | CO | 154.64 (CO) | | |
| | α | 57.8 (CH) | 4.41 (t, 4.6) | 4.41 (t, 4.6) |
| | β | 35.2 ($CH_2$) | 2.90 | 2.89 |
| | aromatic | 1: 124.75 (C) | | |
| | | 2.2’: 131.00 (CH) | 6.95 (d, 7.6) | 6.94 (d, 7.9) |
| | | 3.3’: 114.9 (CH) | 6.63 (d, (7.6) | 6.63 (d, 7.8) |
| | | 4: 156.41 (COH) | | |
| | | 172.47 (CO) | | |
| z-Dhb | CO | 161.87 (CO) | | |
| | α | 126.04 (C) | | |
| | β | 135.5 (CH) | 6.68 (d, 6.7) | 6.68 (q, 6.7) |
| | γ | 12.62 ($CH_3$) | 0.94 (d, 6.7) | 0.93 (d, 6.7) |
| L-Ala | NH | | 8.09 | 8.14 (d, 6.9) |
| | CO | 172.07 (CO) | | |
| | α | 48.45 (CH) | 4.29 (q, 7.1) | 4.29 (q, 6.9) |
| | β | 17.88 (CH3) | 1.19 (d, 7.1) | 1.19 (d, 6.9) |
| L-Arg | NH | | 7.84 | 7.84 (d, 7.9) |
| | CO | 169.30 (CO) | | |
| | α | 50.6 (CH) | 4.48 (bs) | 4.48 (bs) |
| | β | 28.3 ($CH_2$) | 1.53 (m) | 1.52 (m) |
| | γ | 25.5 ($CH_2$) | 1.42 (m); 1.57 (m) | 1.40 (m); 1.57 (m) |
| | δ | 40.40 ($CH_2$) | 3.11 (m) | 3.10 (m) |
| | guanidine | 157.22 (C) | | |
| | | | 6.65 | 6.65 (s) |
| | | | 7.31 | 7.33 (s) |
| L-Pro-thz | α | 57.78 (CH) | 5.29 (dd, 7.8 and 3.7) | 5.29 (dd, 7.9 and 3.8) |
| | β | 32.83 ($CH_2$) | 1.86 (m); 2.27 (m) | 1.85 (m); 2.27 (m) |
| | γ | 23.89 ($CH_2$) | 1.98 (m) | 1.97 (m) |
| | δ | 46.71 ($CH_2$) | 3.75 (bs) | 3.75 (bq, 7.9) |
| | CO | 159.48 (CO) | | |
| | α' | 148.98 (C) | | |
| | β' | 122.86 (CH) | 8.10 (s) | 8.10 (s) |
| | δ' | 171.81 (C) | | |
| L-Gln | NH | | 8.38 (d, 6.1) | 8.38 (d, 6.0) |
| | CO | 174.9 (CO) | | |
| | α | 53.59 (CH) | 3.95 (q, 5.0) | 3.92 (q, 6.0) |
| | β | 29.07 ($CH_2$) | 1.83 (m); 2.00 (m) | 1.82 (m); 1.97 (m) |
| | δ | 31.70 ($CH_2$) | 2.02 (m); 2.11 (m) | 1.99 (m). 2.10 (m) |
| | $CONH_2$ | 174.24 (CO) | 7.15 | |

**Table S10.** Fragment ions observed by UPLC-QTOF-MS$^2$ analysis of **3**, fraction FD2B3A1 (*m/z* 830.3).

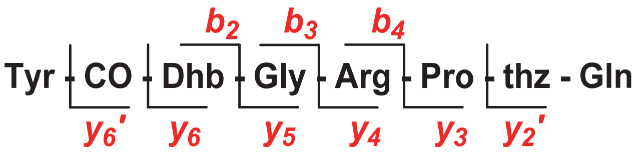

| | | *m/z* acurat. | *m/z* calc. | Error (ppm) |
|---|---|---|---|---|
| $b_2$ | $C_{14}H_{15}N_2O_5^+$ | 291.0987 | 291.0981 | 2.1 |
| $b_3$ | $C_{16}H_{18}N_3O_6^+$ | 348.1203 | 348.1196 | 2.0 |
| $b_4$ | $C_{22}H_{30}N_7O_7^+$ | 504.2211 | 504.2207 | 1.0 |
| $y_2'$ | $C_9H_{12}N_3O_4S^+$ | 258.0552 | 258.0549 | 1.2 |
| $y_3$ | $C_{13}H_{19}N_4O_4S^+$ | 327.1126 | 327.1127 | -0.3 |
| $y_4$ | $C_{19}H_{31}N_8O_5S^+$ | 483.2161 | 483.2138 | 4.8 |
| $y_5$ | $C_{21}H_{34}N_9O_6S^+$ | 540.2338 | 540.2353 | -2.8 |
| $y_6$ | $C_{25}H_{39}N_{10}O_7S^+$ | 623.2738 | 623.2724 | 2.2 |
| $y_6'$ | $C_{26}H_{37}N_{10}O_8S^+$ | 649.2517 | 649.2531 | 2.2 |
| $z_4$ | $C_{19}H_{28}N_7O_5S^+$ | 466.1867 | 466.1873 | -1.3 |
| $z_5$ | $C_{21}H_{29}N_8O_6S^+$ | 521.1940 | 521.1931 | 1.7 |

**Table S11.** Fragment ions observed by UPLC-QTOF-MS$^2$ analysis of **4**, fraction FD2B3C1 (*m/z* 812.3).

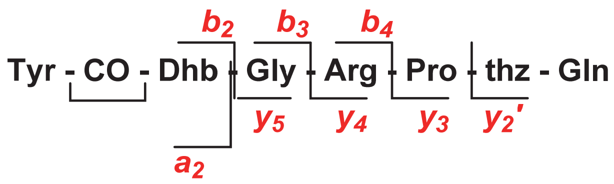

| | | *m/z* acurat. | *m/z* calc. | Error (ppm) |
|---|---|---|---|---|
| $b_2$ | $C_{14}H_{13}N_2O_4^+$ | 273.0877 | 273.0875 | 0.7 |
| $b_3$ | $C_{17}H_{18}N_3O_5^+$ | 330.1092 | 330.1090 | 0.6 |
| $b_4$ | $C_{23}H_{30}N_7O_6^+$ | 486.2105 | 486.2101 | 0.8 |
| $y_2'$ | $C_9H_{12}N_3O_4S^+$ | 258.0550 | 258.0549 | 0.4 |
| $y_3$ | $C_{13}H_{19}N_4O_4S^+$ | 327.1127 | 327.1127 | 0.0 |
| $y_4$ | $C_{19}H_{31}N_8O_5S^+$ | 483.2116 | 483.2138 | -4.6 |
| $y_5$ | $C_{22}H_{36}N_9O_6S^+$ | 540.2369 | 540.2353 | 3.0 |
| $a_2$ | $C_{13}H_{13}N_2O_3^+$ | 245.0926 | 245.0926 | 0.0 |
| $a_3$ | $C_{16}H_{18}N_3O_4^+$ | 302.1146 | 302.1141 | 1.7 |
| $a_4$ | $C_{22}H_{30}N_7O_5^+$ | 458.2151 | 458.2152 | -0.2 |
| $a_6$ | $C_{30}H_{40}N_9O_6S^+$ | 640.2667 | 640.2666 | 0.2 |

**Table S12.** Selected fragment ions observed by UPLC-QTOF-MS$^2$ analysis for **5**/**6** on fraction FD2B2A (*m/z* 858.3).

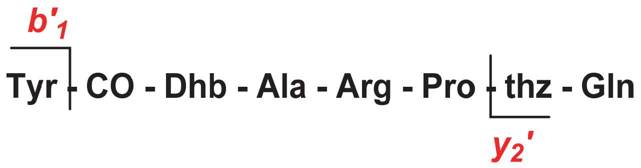

| $t_r$ (min) | | | *m/z* acurat. | *m/z* calc. | Error (ppm) |
|---|---|---|---|---|---|
| 4.7 | **[M + H]$^+$** | $C_{37}H_{52}N_{11}O_{11}S^+$ | 858.3560 | 858.3568 | -0.9 |
| | **[M - H]$^-$** | $C_{37}H_{50}N_{11}O_{11}S^-$ | 856.3398 | 856.3412 | -1.6 |
| | ***y$_2$'*-NH$_3$** | $C_{10}H_{11}N_2O_9S^+$ | 255.0441 | 155.0440 | 0.4 |
| 4.8 | **[M + H]$^+$** | $C_{37}H_{52}N_{11}O_{11}S^+$ | 858.3559 | 858.3568 | -1.0 |
| | **[M - H]$^-$** | $C_{37}H_{50}N_{11}O_{11}S^-$ | 856.3400 | 856.3412 | -1.4 |
| | ***b$_1$'*-COCH$_3$** | $C_8H_{10}NO^+$ | 136.0762 | 136.0762 | 0.0 |

**Table S13.** Fragment ions observed by UPLC-QTOF-MS$^2$ analysis of sample pseudovibriamide A1 degradation (*m/z* 638.2).

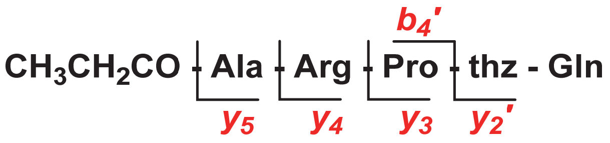

| | | *m/z* acurat. | *m/z* calc. | Error (ppm) |
|---|---|---|---|---|
| **$[M + H]^+$** | $C_{26}H_{40}N_9O_8S^+$ | 638.2740 | 638.2721 | 3.0 |
| **$[M - NH_3]^+$** | $C_{26}H_{37}N_8O_8S^+$ | 621.2459 | 621.2455 | 0.8 |
| **$[M - CONH_3]^+$** | $C_{25}H_{37}N_8O_7S^+$ | 593.2512 | 593.2506 | 1.0 |
| **$[M - CN_3H_5]^+$** | $C_{25}H_{35}N_6O_8S^+$ | 579.2249 | 579.2237 | 2.1 |
| **$[M - H]^-$** | $C_{26}H_{38}N_9O_8S^+$ | 636.2568 | 636.2564 | 0.6 |
| ***$b_4$'*** | $C_{13}H_{22}N_5O_4^+$ | 312.1683 | 312.1672 | 3.5 |
| ***$y_2$'*** | $C_9H_{12}N_3O_4S^+$ | 258.0563 | 258.0549 | 5.4 |
| ***$y_3$*** | $C_{13}H_{19}N_4O_4S^+$ | 327.1138 | 327.1127 | 3.4 |
| ***$y_4$*** | $C_{19}H_{31}N_8O_5S^+$ | 483.2137 | 483.2138 | -0.2 |
| ***$y_5$*** | $C_{22}H_{36}N_9O_6S^+$ | 554.2505 | 554.2509 | -0.7 |

**Table S14.** $^1H$ and $^{13}C$ NMR data for **7**, fraction C23C1B in DMSO-$d_6$ + TFA ($^1H$: 600 MHz, $^{13}C$: 150 MHz).

| Residue | Position | $\delta_C$, type | $\delta_H$ (J in Hz) |
|---|---|---|---|
| **L-Tyr** | $CO_2H$ | 174.04 | |
| | NH | | 6.11 |
| | CO | 155.19 | |
| | α | 54.09 | 4.25 |
| | β | 36.83 | 2.83, 2.72 |
| | aromatic | 1: 155.99 | |
| | | 2, 2': 114.99 | 6.66 |
| | | 3, 3': 130.14 | 6.95 |
| | | 4: 127.58 | |
| z-**Dhb** | NH | | 7.82 |
| | CO | 164.98 | |
| | α | 131.95 | |
| | β | 121.71 | 5.94 (q, 7.1) |
| | γ | 12.54 | 1.59 (d, 7.1) |
| **L-Ala** | NH | | 8.41 |
| | CO | 171.67 | |
| | α | 48.14 | 4.33 |
| | β | 17.99 | 1.26 (d, 7.0) |
| **L-Arg** | NH | | 8.25 |
| | CO | 170.27 | |
| | α | 50.28 | 4.56 |
| | β | 28.16 | 1.59, 1.78 |
| | γ | 24.89 | 1.56 |
| | δ | 40.37 | 3.11 |
| | guanidine | | 2.53 |
| | | 156.89 | |
| | | | 7.35 |
| | | | 7.53, 7.49 |
| **L-Pro-thz** | α | 58.52 | 5.31 |
| | β | 31.59 | 2.27 |
| | γ | 24.01 | 2.02, 1.97 |
| | δ | 46.80 | 3.78, 3.74 |
| | CO | 160.62 | |
| | α' | 149.03 | |
| | β' | 124.10 | 8.16 |
| | δ' | 173.87 | |
| **L-Gln** | NH | | 8.73 |
| | CO | 170.81 | |
| | α | 51.93 | 4.43 |
| | β | 31.14 | 2.14 |
| | δ | 25.62 | 2.15, 1.97 |
| | $CONH_2$ | 173.51 | |
| **Hyp*** | α | 57.12 | 4.77 |
| | β | 70.38 | 5.10 |
| | γ | 29.57 | 2.24 |

| | | | |
|---|---|---|---|
| | δ | 44.66 | 3.77, 3.52 |
| | α' (CH) | 69.28 | 5.33 |
| | α'' ($CH_2$) | 35.72 | 2.47 |
| | $CO_2H$ | 171.08 | |
| | $C_\beta O_2$ | 173.01 | |
| | β' ($CH_2$) | 26.81 | 2.30 |
| | β'' ($CH_3$) | 8.82 | 1.02 (t, 7.7) |
| **L-Val** | NH | | 8.10 |
| | CO | 169.26 | |
| | α | 55.26 | 4.08 |
| | β | 29.28 | 2.01 |
| | γ | 16.93<br>18.28 | 0.81<br>0.89 |

| *Key degradation signals* | | |
|---|---|---|
| CO | 199.24 | |
| $CH_2$ | 30.15 | 2.81 (ddd, 2.4, 7.1, 15.3) |
| $CH_3$ | 7.02 | 0.96 (t, 7.1) |

Hyp*, modified hydroxyproline.

**Table S15.** Fragment ions observed by UPLC-QTOF-MS[2] analysis of **7**, fraction C23C1B (*m/z* 1156.5).

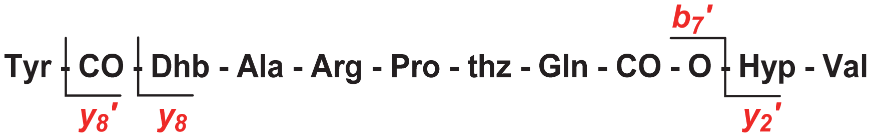

| | | ***m/z* acurat.** | ***m/z* calc.** | **Error (ppm)** |
|---|---|---|---|---|
| ***$b_7$'*** | $C_{36}H_{50}N_{11}O_8S^+$ | 844.3410 | 844.3412 | -0.2 |
| ***$y_2$'*** | $C_{15}H_{25}N_2O_5^+$ | 313.1786 | 313.1763 | 7.3 |
| ***$y_8$'*** | $C_{42}H_{63}N_{12}O_{13}S^+$ | 975.4393 | 975.4358 | 3.6 |
| ***$y_8$'-$y_2$'*** | $C_{27}H_{39}N_{10}O_8S^+$ | 663.2687 | 663.2673 | 2.1 |

**Table S16.** Fragment ions observed by UPLC-QTOF-MS$^2$ analysis of **8/9**, fraction C23C1C (*m/z* 1170.5).

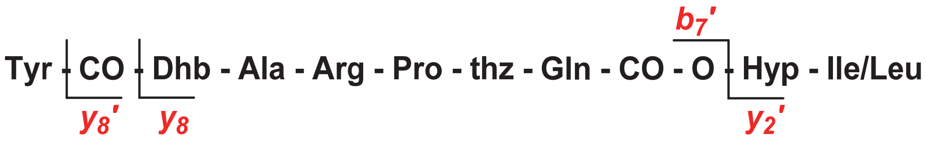

| | | *m/z* acurat. | *m/z* calc. | Error (ppm) |
|---|---|---|---|---|
| **[M+H]$^+$** | $C_{52}H_{76}N_{13}O_{16}S^+$ | 1170.5229 | 1170.5254 | -2.1 |
| **[M+2H]$^{++}$** | $C_{52}H_{77}N_{13}O_{16}S^{++}$ | 585.7676 | 585.7660 | 2.7 |
| **[M-H]$^-$** | $C_{52}H_{74}N_{13}O_{16}S^-$ | 1168.5101 | 1168.5097 | 0.3 |
| ***b$_7$'*** | $C_{36}H_{50}N_{11}O_8S^+$ | 844.3412 | 844.3412 | 0.0 |
| ***y$_2$'*** | $C_{16}H_{27}N_2O_5^+$ | 327.1922 | 327.1920 | 0.6 |
| ***y$_8$*** | $C_{42}H_{67}N_{12}O_{12}S^+$ | 963.4732 | 963.4722 | 1.0 |
| ***y$_8$'*** | $C_{43}H_{65}N_{12}O_{13}S^+$ | 989.4521 | 989.4515 | 0.6 |
| ***y$_8$'-y$_2$'*** | $C_{27}H_{35}N_{10}O_8S^+$ | 663.2673 | 663.2673 | 0.0 |

**Table S17.** Fragment ions observed by UPLC-QTOF-MS$^2$ analysis of **10**, fraction C23C1C (*m/z* 1142.5).

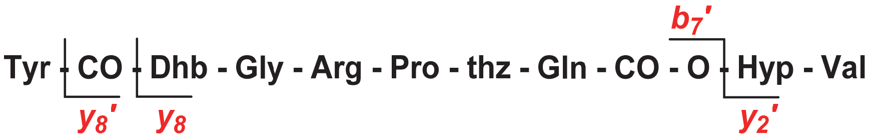

| | | *m/z* acurat. | *m/z* calc. | Error (ppm) |
|---|---|---|---|---|
| **[M+H]$^+$** | $C_{50}H_{72}N_{13}O_{16}S^+$ | 1142.4929 | 1142.4941 | -1.1 |
| **[M+2H]$^{++}$** | $C_{50}H_{73}N_{13}O_{16}S^{++}$ | 571.7523 | 571.7504 | 1,5 |
| **[M-H]$^-$** | $C_{50}H_{70}N_{13}O_{16}S^-$ | 1140.4781 | 1140.4684 | -0.3 |
| ***b$_7$'*** | $C_{36}H_{50}N_{11}O_8S^+$ | 830.3218 | 830.3255 | -4.5 |
| ***y$_2$'*** | $C_{15}H_{25}N_2O_5^+$ | 313.1763 | 313.1763 | 0.0 |
| ***y$_8$'*** | $C_{43}H_{65}N_{12}O_{13}S^+$ | 961.4154 | 961.4202 | -5.0 |
| ***y$_8$'-y$_2$'*** | $C_{27}H_{35}N_{10}O_8S^+$ | 649.2501 | 649.2517 | -2.5 |

**Table S18.** Fragment ions observed by UPLC-QTOF-MS$^2$ analysis of **11/12**, fraction C23C1D (*m/z* 1152.5).

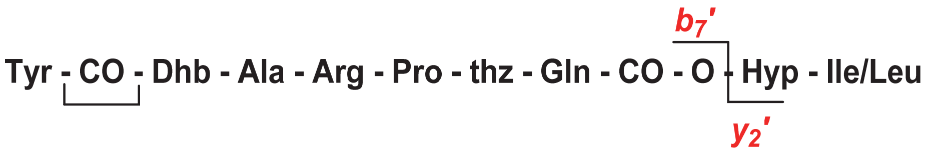

| | | *m/z* **acurat.** | *m/z* **calc.** | **Error (ppm)** |
|---|---|---|---|---|
| **[M+H]$^+$** | $C_{52}H_{74}N_{13}O_{15}S^+$ | 1152.5126 | 1152.5248 | -1.9 |
| **[M+2H]$^{++}$** | $C_{52}H_{75}N_{13}O_{15}S^{++}$ | 576.7609 | 576.7608 | 0.2 |
| **[M-H]$^-$** | $C_{52}H_{72}N_{13}O_{15}S^-$ | 1150.4996 | 1150.4992 | 0.3 |
| ***b$_7$'*** | $C_{36}H_{48}N_{11}O_{10}S^+$ | 826.3322 | 826.3306 | 1.9 |
| ***y$_2$'*** | $C_{16}H_{27}N_2O_5^+$ | 327.1912 | 327.1920 | -2.4 |

**Table S19**. Absorbance measurements of biofilm assay using 0.1% crystal-violet solution. Average value and standard deviation calculated for six replicates, considering all the biological replicates. If necessary, samples were diluted so that the measured absorbance values were always <1.0. The observed values using diluted samples were then multiplied accordingly to obtain the reported values below.

| | | | Absorbance (595 nm) | | | | | |
|---|---|---|---|---|---|---|---|---|
| **24h (replicates)** | *1* | *2* | *3* | *4* | *5* | *6* | *Average* | *Std. Dev.* |
| MB (blank) | 0.000 | 0.001 | -0.001 | 0.001 | 0.001 | 0.001 | 0.001 | 0.001 |
| Ab134-WT | 2.344 | 2.261 | 2.112 | 2.264 | 2.161 | 2.735 | 2.313 | 0.223 |
| *pppA*$^-$MT#01 | 4.264 | 3.542 | 3.224 | 3.442 | 3.432 | 4.174 | 3.689 | 0.431 |
| *pppA*$^-$MT#45 | 3.716 | 3.418 | 3.640 | 3.530 | 3.182 | 4.184 | 3.612 | 0.337 |
| *pppA*$^-$MT#47 | 3.994 | 3.054 | 3.200 | 3.052 | 3.246 | 3.890 | 3.406 | 0.424 |
| *pppD*$^-$MT#24 | 0.445 | 0.526 | 0.396 | 0.406 | 0.407 | 0.428 | 0.435 | 0.048 |
| *pppD*$^-$MT#67 | 0.512 | 0.520 | 0.464 | 0.431 | 0.428 | 0.449 | 0.467 | 0.040 |

| **48h (replicates)** | *1* | *2* | *3* | *4* | *5* | *6* | *Average* | *Std. Dev.* |
|---|---|---|---|---|---|---|---|---|
| MB (blank) | 0.000 | -0.001 | -0.003 | 0.000 | -0.002 | 0.004 | 0.004 | 0.002 |
| Ab134-WT | 4.708 | 4.604 | 3.976 | 3.938 | 4.396 | 4.788 | 4.402 | 0.369 |
| *pppA*$^-$MT#01 | 6.396 | 5.568 | 5.528 | 5.352 | 6.224 | 6.304 | 5.895 | 0.461 |
| *pppA*$^-$MT#45 | 7.252 | 5.560 | 4.772 | 5.308 | 6.124 | 5.924 | 5.823 | 0.846 |
| *pppA*$^-$MT#47 | 6.596 | 6.736 | 4.804 | 4.816 | 4.588 | 5.448 | 5.498 | 0.950 |
| *pppD*$^-$MT#24 | 1.240 | 1.050 | 1.456 | 1.256 | 1.312 | 2.902 | 2.597 | 0.682 |
| *pppD*$^-$MT#67 | 2.296 | 2.716 | 2.38 | 2.418 | 2.632 | 3.142 | 3.142 | 0.311 |

**Statistical analysis performed** according to the Tukey test with 95% confidence level on **O**riginPro 8.5.

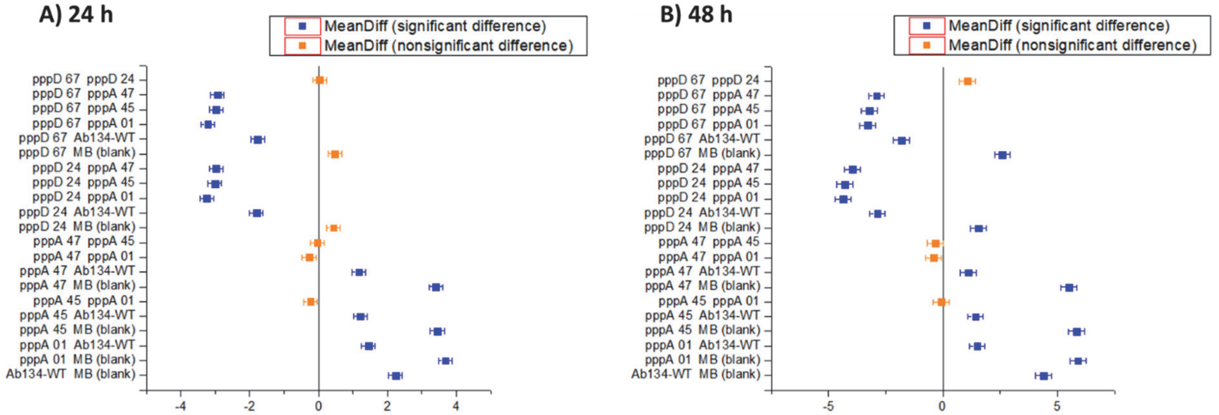

**Table S20.** *Pseudovibrio* genomes which harbor a member of the *ppp* BGC family, highlighting the presence of an origin of replication for plasmids at the same contig (ctg).

| Organism | Ctg # | NCBI accession | Ctg size (bp) | Replication protein (position) |
|---|---|---|---|---|
| ***P. denitrificans* DSM17465** | 36 | NZ_FPBD01000007 | 422,114 (95 bp overlapped ends) | *repC* (14,322-15,641) |
| ***P. denitrificans* JCM12308** | 94 | BAZK01000008 | 421,911 (no overlap) | *repC* (396,695-395,328) |
| ***P. axinellae* Ad2** | 162 | NZ_LMCB01000043 | 72,529 | - |
| ***P. axinellae* DSM24994** | 93 | NZ_FOFM01000016 | 72,693 | - |
| ***Pseudovibrio* sp. Alg231-02** | 23 | NZ_FREX01000017 | 61,264 | - |
| ***Pseudovibrio* sp. POLY-S9** | 271 | NZ_LCWZ01000112 | 50,837 | - |
| ***Pseudovibrio* sp. W64** | 49 | NZ_LMCI01000029 | 80,426 | *repB* (58,391-57,297) |
| ***Pseudovibrio* sp. W74** | 64 | NZ_LMCJ01000042 | 156,575 | - |
| ***Pseudovibrio* sp. WM33** | 159 | NZ_LMCK01000096 | 59,121 | - |
| ***Pseudovibrio* sp. Ad5** | 66 | NZ_LMCH01000003 | 190,885 | - |
| ***Pseudovibrio* sp. Ad13** | 36 | NZ_LMCC01000034 | 410,131 (120 bp overlapped ends) | *repC* (287,243-288,496) |
| ***Pseudovibrio* sp. Ad14** | 57 | NZ_LMCD01000015 | 169,556 | - |
| ***Pseudovibrio* sp. Ad26** | 159 | NZ_LMCE01000064 | 40,354 | - |
| ***Pseudovibrio* sp. Ad37** | 224 | NZ_LMCF01000190 | 45,042 | - |
| ***Pseudovibrio* sp. Ad46** | 85 | NZ_LMCFG01000068 | 269,303 | - |
| ***Pseudovibrio* sp. Tun.PSC04-5.I4** | 8 | NZ_FNLB01000004 | 183,229 (10,68 kb overlapped ends) | *repC* (5,472-6,749) |

**Table S21.** Nonribosomal peptide bacterial products containing ureido-linkages. GenBank accession codes for known BGCs are also shown.

| **Compound**[reference] **(gene cluster)** | **Organism** | **Ureido linkage** |
|---|---|---|
| **minosaminomycin**[1] | *Streptomyces* sp. | Leu-CO-Arg |
| **SP-chymostatin**[2] | *Streptomyces libani* | Phe-CO-Arg |
| **α-MAPI**[3] | *Streptomyces nigrescens* | Phe-CO-Arg |
| **strepin**[4] | *Streptomyces tanabeensis* | Arn-CO-Leu |
| **Mer-N5075A**[5] | *Streptomyces chromofuscus* | Phe-CO-Arg |
| **GE20372**[6] | *Streptomyces* sp. | Tyr-CO-Arg |
| **mureidomycins**[7] **(DS999644.1)** | *Streptomyces flavidovirens* | $aa_1$-CO- $aa_2$<br>$aa_1$: *m*-Tyr, Phe<br>$aa_2$: Met, $Met^{SO}$, Leu, Ile |
| **pacidamycins**[8] **(HM855229.1)** | *Streptomyces coeruleorubidus* | $aa_1$-CO-Ala<br>$aa_1$: *m*-Tyr, Trp, Phe |
| **napsamycins**[9] **(HQ287563.1)** | *Streptomyces* sp. | *m*-Tyr-CO-Met |
| **antipains**[10] **(LWBU01000384.1)** | *Streptomyces albulus* | $aa_1$-CO-Arg<br>$aa_1$: Phe, Tyr |
| **muraymycins**[11] | *Streptomyces* sp. | Leu-CO-Arg |
| **sansanmycins**[12] | *Streptomyces* sp. | $aa_1$-CO- $aa_2$<br>$aa_1$: *m*-Tyr, Trp<br>$aa_2$: Met, $Met^{SO}$, Phe, Leu |
| **ferintoic acids**[13] | *Microcystis aeruginosa* | Trp-CO-Lys |
| **shizopeptins**[14] | *Schizothrix* sp. | Ile-CO-Lys |
| **brunsvicamides**[15] | *Tychonema* sp.s | Ile-CO-Lys |
| **pompanopeptins**[16] | *Lyngbya confervoides* | Ile-CO-Lys |
| **anabaenopeptins**[17–23] **(GU174493.1, EF672686.1)** | *Anabaena* spp.<br>*Planktothrix* sp.<br>*Nostoc* sp.<br>*Mycrocystis* spp.<br>*Oscillatoria* spp.<br>*Aphanizomenon flos-aquae*<br>Unknown cianobacteria bloom | $aa_1$-CO-Lys<br>$aa_1$: Tyr, Ile, Phe, Arg, Trp |
| **oscillamides**[21,24,25] | *Mycrocystis* sp.<br>*Oscillatoria agardhii*<br>*Planktohrix* spp. | $aa_1$-CO-Lys<br>$aa_1$: Tyr, Arg |
| **syringolin**[26,27] **(AJ548826.1, AJVWM01000074.1)** | *Pseudomonas syringae*<br>*Rhizobium* sp. | $aa_1$-CO- $aa_1$<br>$aa_1$: Leu, Ile |

| | | |
|---|---|---|
| **namalides**[19,28] | *Nostoc* sp. | $aa_1$-CO-Lys<br>$aa_1$: Leu, Ile, Phe |
| **lyngbyaureidamide**[29] | *Lyngbya* sp. | Phe-CO-Lys |
| **nostamides**[30] | *Nostoc punctiforme* | Phe-CO-Lys |

**Table S22.** Bacterial products containing Dhb residues. GenBank accession codes for known BGCs are also shown.

| **Compound[reference] (gene cluster)** | **Organism** | **Position** |
|---|---|---|
| **microcystins**[31–34] **(AF183408.1, AJ441056.1, AY212249.1, KX891213.1)** | *Microcystis aeruginosa*<br>*Anabaena* sp.<br>*Planktothrix agardhii*<br>*Fischerella* sp. | $aa_1$- $aa_2$-$aa_3$<br><br>$aa_1$: D-Glu, D-Glu($OCH_3$)<br>$aa_2$: Dha, Ser, Dhb, Lan<br>$aa_3$: D-Ala, D-Ser, D-Leu |
| **FR901228**[35] **(EF210776.1)** | *Chromobacterium violaceum* | D-Cys-Dhb-L-Val |
| **nodularin**[36] **(MF668122.1)** | *Nostoc* sp. | D-Glu-Dhb-D-Asp |
| **bogorol**[37,38] **(KY810814.1)** | *Brevibacillus laterosporus* | Hmp-Dhb-$aa_3$<br><br>$aa_3$: L-Val, L-Met, L-$Met^{SO}$ |
| **pantomycin**[39] | *Streptomyces* spp. | D-Ala-Dhb-D-Thr |
| **pseudomycins**[40] | *Pseudomonas syringae* | L-aThr-Dhb-L-Asp |
| **fuscopeptins**[41] | *Pseudomonas fuscovaginae* | FA-Dhb-D-Pro<br>D-Val-Dhb-D-aThr<br>L-Ala-Dha-Dha-L-Phe |
| **syringomycins**[42] **(CP000075.1)** | *Pseudomonas* spp. | L-Phe-Dhb-L-Asp(OH) |
| **syringopeptins**[42,43] **(AF286216.2)** | *Pseudomonas* sp. | FA-Dhb-Pro<br>Ala-Dhb-aThr<br>Ala-Dhb-Ala |
| **syringostatins**[44] | *Pseudomonas* sp. | Thr-Dhb-Asp(OH) |
| **hassallidins**[45,46] **(KJ502174.1, LT546031.1)** | *Anabaena* sp.<br>*Planktothrix serta* | D-Tyr-Dhb-D-Gln |
| **loihichelins**[47] | *Halomonas* sp. | D-Orn(OH)-Dhb-L-Ser |
| **tolaasins**[48] **(HE967327.1)** | *Pseudomonas* spp. | FA-Dhb-Pro<br>Val-Dhb-Thr |
| **sessilin**[49] **(JQ309920.1)** | *Pseudomonas* sp. | Val-Dhb-Thr |
| **corpeptins**[50] **cormycins** | *Pseudomonas corrugata* | FA-Dhb-Pro<br>Val-Dhb-Hse<br>Ile-Dha-Ala<br>Val-Dhb-Thr |
| **jessenipeptin**[51] | *Pseudomonas* sp. | FA-Dhb-ᴅ-Pro<br>D-Leu-Dhb-D-aThr |
| **puwaianaphycins**[52] **(MH325199.1, MH325197.1)** | *Anabaena* sp.<br>*Cylindrospermum alatosporum* | Val-Dhb-$aa_1$-$aa_2$<br><br>$aa_1$: Asn, Gln, Thr<br>$aa_2$: Dhb, Thr, Val |

**Table S23.** Natural products containing an imidazolidinyl-dione ring.

| Compound class[reference] | Organism |
|---|---|
| **Midcapamides**[53] | *Agelas* spp. |
| **5-[(6-Bromo-1*H*-indol-3-yl)methylene]-2,4-imidazolidinedione**[54] | *Smespongia* sp.<br>*Aspergillus* sp. |
| **Plyandrocarpamide**[55] | *Polyandrocarpa* sp. |
| **Agesamides**[56] | *Agelas* sp. |
| **Mukanadins**[57]<br>**Nakamurines**<br>**Nemoechines** | *Agelas* sp.<br>*Didiscus* sp. |
| **Parazoanthines**[58] | *Parazoanthus* sp. |
| **Lechagodines**[59] | *Leucetta* sp. |
| **Aplysinopsins-like**[60] | *Fascaplysinopsis* sp. |
| **Spiroreticulatine**[61] | *Fascaplysinopsis* sp. |
| **Hydantocidin**[62] | *Streptomyces* sp.<br>*Actinomadura* sp. |
| **(*S*)-5-Isopropyl-3-methoxyimidazolidine-2,4-dione**[63] | *Phoma* sp. |
| **Hemimycalin**[64] | *Hemimycale* sp. |
| **Naamines**<br>**Naamidines**<br>**Chagosendines**[65] | *Leucetta* sp. |
| **Clathridine**[66] | *Clathrina* sp. |
| **Axinohydantoin**[67]<br>**Fuscin**<br>**Spongiadicidins** | *Axinella* sp.<br>*Phacellia* sp. |
| **Tubastrindoles**[68] | *Smespongia* sp. |
| **Exaguamine**[69] | *Neopetrosia* sp. |

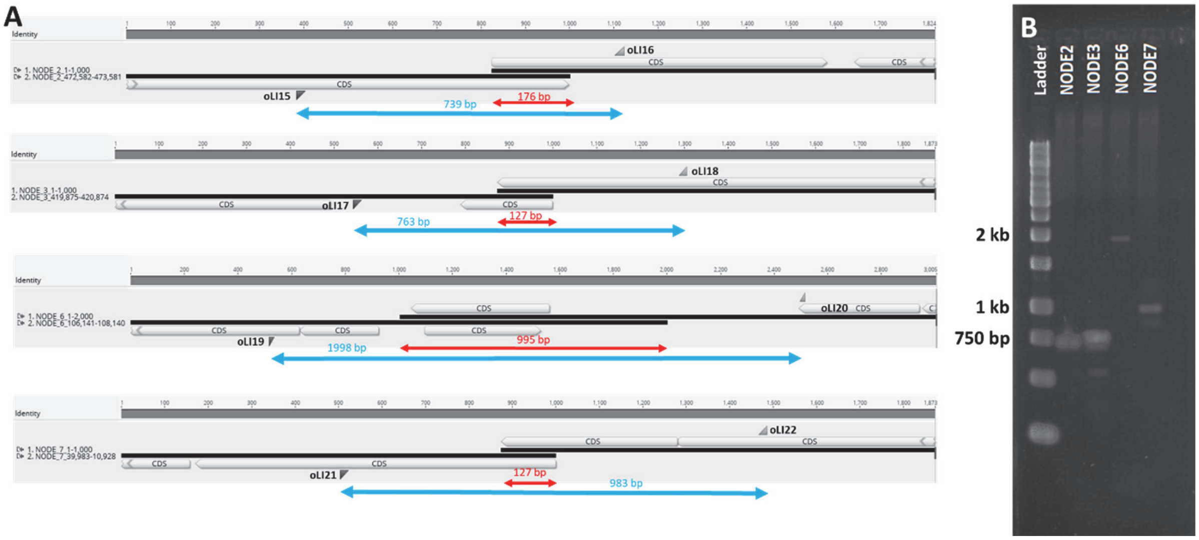

**Figure S1.** Multireplicon confirmation by PCR. A) Design of PCR to confirm that four contigs (NODES 2, 3, 6, and 7) represent circular plasmids which we named 1, 2, 4, and 5, respectively. Overlaps at the ends of each NODE are indicated with red double arrows; and expected fragments after PCR are indicated with blue double arrows. Primers used are indicated. B) Gel electrophoresis analysis of PCR products confirming that the nodes represent circular replicons. L, molecular weight DNA ladder (GeneRuler 1kb DNA Ladder - ThermoScientific).

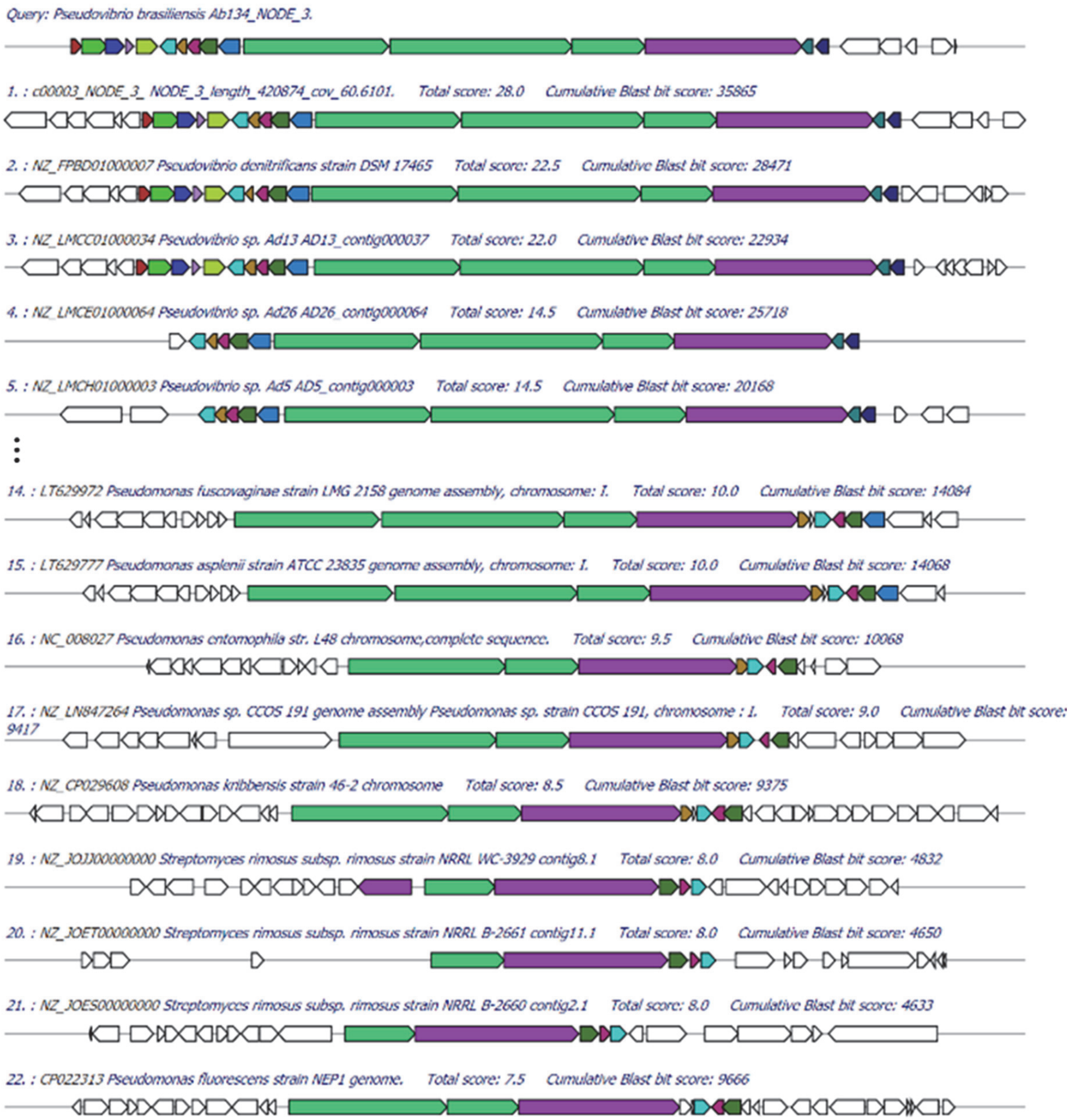

**Figure S2.** Representative Multigeneblast results. The *ppp* BGC from *P. brasiliensis* Ab134 was used as query to search for similar BGCs**.**

| Gene | *pppA* | | | | | *pppB* | | | | | | | *pppC* | | | *pppD* | | | | | |
|---|---|---|---|---|---|---|---|---|---|---|---|---|---|---|---|---|---|---|---|---|---|
| Module | 1 | 2 | | 3 | | 4 | | 5 | | 6 | | | 7 | | | 8 | 9 | | 10 | | |
| | A | C | A | $C_D$ | A | C | A | C | A | $C_y$ | A | Ox | C | A | TE | A | C | A | KS | KR | TE |
| ***Pseudovibrio brasiliensis* Ab134** | Y | ✓ | T | ✓ | X | ✓ | R | ✓ | P | ✓ | C | ✓ | ✓ | Q | ✓ | I | ✓ | P | ✓ | ✓ | ✓ |
| ***Pseudovibrio* sp. Alg231-02** | Y | ✓ | T | ✓ | G | ✓ | R | ✓ | P | ✓ | C | ✓ | ✓ | Q | ✓ | I | ✓ | P | ✓ | ✓ | ✓ |
| ***Pseudovibrio* sp. POLY-S9** | Y | ✓ | T | ✓ | G | ✓ | R | ✓ | P | ✓ | C | ✓ | ✓ | Q | ✓ | I | ✓ | P | ✓ | ✓ | ✓ |
| ***P. axinellae* Ad2** | Y | ✓ | T | ✓ | G | ✓ | R | ✓ | P | ✓ | C | ✓ | ✓ | Q | ✓ | I | ✓ | P | ✓ | ✓ | ✓ |
| ***P. axinellae* DSM24994** | Y | ✓ | T | ✓ | G | ✓ | R | ✓ | P | ✓ | C | ✓ | ✓ | Q | ✓ | I | ✓ | P | ✓ | ✓ | ✓ |
| ***Pseudovibrio* sp. W64** | Y | ✓ | T | ✓ | G | ✓ | R | ✓ | P | ✓ | C | ✓ | ✓ | Q | ✓ | I | ✓ | P | ✓ | ✓ | ✓ |
| ***Pseudovibrio* sp. W74** | Y | ✓ | T | ✓ | G | ✓ | R | ✓ | P | ✓ | C | ✓ | ✓ | Q | ✓ | I | ✓ | P | ✓ | ✓ | ✓ |
| ***Pseudovibrio* sp. WM33** | Y | ✓ | T | ✓ | G | ✓ | R | ✓ | P | ✓ | C | ✓ | ✓ | Q | ✓ | I | ✓ | P | ✓ | ✓ | ✓ |
| ***Pseudovibrio* sp. Ad5** | Y | ✓ | T | ✓ | G | ✓ | R | ✓ | P | ✓ | C | ✓ | ✓ | Q | ✓ | I | ✓ | P | ✓ | ✓ | ✓ |
| ***Pseudovibrio* sp. Ad13** | Y | ✓ | T | ✓ | G | ✓ | R | ✓ | P | ✓ | C | ✓ | ✓ | Q | ✓ | I | ✓ | P | ✓ | ✓ | ✓ |
| ***Pseudovibrio* sp. Ad14** | Y | ✓ | T | ✓ | G | ✓ | R | ✓ | P | ✓ | C | ✓ | ✓ | Q | ✓ | I | ✓ | P | ✓ | ✓ | ✓ |
| ***Pseudovibrio* sp. Ad26** | X | ✓ | T | ✓ | G | ✓ | R | ✓ | P | ✓ | C | ✓ | ✓ | Q | ✓ | I | ✓ | P | ✓ | ✓ | ✓ |
| ***Pseudovibrio* sp. Ad37** | Y | ✓ | T | ✓ | G | ✓ | R | ✓ | P | ✓ | C | ✓ | ✓ | Q | ✓ | I | ✓ | P | ✓ | ✓ | ✓ |
| ***Pseudovibrio* sp. Ad46** | Y | ✓ | T | ✓ | G | ✓ | R | ✓ | P | ✓ | C | ✓ | ✓ | Q | ✓ | I | ✓ | P | ✓ | ✓ | ✓ |
| ***Pseudovibrio* sp. Tun.PSC04-5.I4** | Y | ✓ | T | ✓ | G * | ✓ | R | ✓ | P | ✓ | C | ✓ | ✓ | Q | ✓ | I | ✓ | P | ✓ | ✓ | ✓ |
| ***P. denitrificans* DSM17465** | X | ✓ | T | ✓ | X | ✓ | R | ✓ | P | ✓ | C | ✓ | ✓ | Q | ✓ | I | ✓ | P | ✓ | ✓ | ✓ |
| ***P. denitrificans* JCM12308** | X | ✓ | T | ✓ | X | ✓ | R | ✓ | P | ✓ | C | ✓ | ✓ | Q | ✓ | I | ✓ | P | ✓ | ✓ | ✓ |
| ***Ps. fuscovaginae* LMG2158** | X | ✓ | T | ✓ | T | ✓ | R | ✓ | P | ✓ | C | ✓ | ✓ | X | ✓ | I | ✓ | P | ✓ | ✓ | ✓ |
| ***Ps. asplenii* ATCC 23835** | X | ✓ | T | ✓ | T | ✓ | R | ✓ | P | ✓ | C | ✓ | ✓ | X | ✓ | I | ✓ | P | ✓ | ✓ | ✓ |
| ***Ps. kribbensis* 46-2** | - | - | - | - | - | - | R | ✓ | P | ✓ | C | ✓ | ✓ | X | ✓ | I | ✓ | P | ✓ | C2 | ✓ |
| ***Pseudomonas* sp. CCOS 191** | - | - | - | - | - | - | R | ✓ | P | ✓ | C | ✓ | ✓ | X | ✓ | I | ✓ | P | ✓ | C2 | ✓ |
| ***Ps. entomophila* L48** | - | - | - | - | - | - | R | ✓ | P | ✓ | C | ✓ | ✓ | X | ✓ | I | ✓ | P | ✓ | C2 | ✓ |
| ***Ps. fluorescens* NEP1** | - | - | - | - | - | - | R | ✓ | P | ✓ | C | ✓ | ✓ | X | ✓ | I | ✓ | P | ✓ | C2 | ✓ |
| ***Pseudomonas* sp. DR 5-09** | - | - | - | - | - | - | R | ✓ | P | ✓ | C | ✓ | ✓ | X | ✓ | I | ✓ | P | ✓ | C2 | ✓ |
| ***Ps. moraviensis* BS3668** | - | - | - | - | - | - | R | ✓ | P | ✓ | C | ✓ | ✓ | X | ✓ | I | ✓ | P | ✓ | C2 | ✓ |
| ***Ps. reinekei* BS3776** | - | - | - | - | - | - | R | ✓ | P | ✓ | C | ✓ | ✓ | X | ✓ | I | ✓ | P | ✓ | C2 | ✓ |
| ***Ps. syringae pv. syringae* CFBP4215** | - | - | - | - | - | - | R | ✓ | P | ✓ | C | ✓ | ✓ | X | ✓ | X | ✓ | P | ✓ | C2 | ✓ |
| ***Ps. syringae* 31R1** | - | - | - | - | - | - | R | ✓ | P | ✓ | C | ✓ | ✓ | X | ✓ | X | ✓ | P | ✓ | C2 | ✓ |
| ***Ps. viridiflava* CFBP1590** | - | - | - | - | - | - | R | ✓ | P | ✓ | C | ✓ | ✓ | X | ✓ | X | ✓ | P | ✓ | C2 | ✓ |
| ***Ps. savastanoi pv. phaseolicola* 1448A** | - | - | - | - | - | - | R | ✓ | P | ✓ | C | ✓ | ✓ | X | ✓ | X | ✓ | P | ✓ | C2 | ✓ |
| ***Ps. amygdali pv. lachrymans* NM002** | - | - | - | - | - | - | R | ✓ | P | ✓ | C | ✓ | ✓ | X | ✓ | X | ✓ | P | ✓ | C2 | ✓ |
| ***Ps. savastanoi pv. savastanoi* NCPPB3335** | - | - | - | - | - | - | R | ✓ | P | ✓ | C | ✓ | ✓ | X | ✓ | X | ✓ | P | ✓ | C2 | ✓ |
| ***Ps. syringae pv. syringae* SM** | - | - | - | - | - | - | R | ✓ | P | ✓ | C | ✓ | ✓ | X | ✓ | X | ✓ | P | ✓ | C2 | ✓ |
| ***Ps. syringae pv. syringae* CFB2118** | - | - | - | - | - | - | R | ✓ | P | ✓ | C | ✓ | ✓ | X | ✓ | X | ✓ | P | ✓ | C2 | ✓ |
| ***Ps. syringae pv. cerasicola* CFB6109** | - | - | - | - | - | - | R | ✓ | P | ✓ | C | ✓ | ✓ | X | ✓ | X | ✓ | P | ✓ | C2 | ✓ |
| ***Ps. syringae pv. lapsa* ATCC 10859** | - | - | - | - | - | - | R | ✓ | P | ✓ | C | ✓ | ✓ | X | ✓ | X | ✓ | P | ✓ | C2 | ✓ |
| ***Ps. synringae pv. syringae* B728a** | - | - | - | - | - | - | R | ✓ | P | ✓ | C | ✓ | ✓ | X | ✓ | X | ✓ | P | ✓ | C2 | ✓ |
| ***Ps. syringae pv. atrofacien* LMG5095** | - | - | - | - | - | - | R | ✓ | P | ✓ | C | ✓ | ✓ | X | ✓ | X | ✓ | P | ✓ | C2 | ✓ |
| ***Ps. syringae pv. syringae* B64** | - | - | - | - | - | - | R | ✓ | P | ✓ | C | ✓ | ✓ | X | ✓ | X | ✓ | P | ✓ | C2 | ✓ |
| ***Ps. syringae pv. syringae* HS191** | - | - | - | - | - | - | R | ✓ | P | ✓ | C | ✓ | ✓ | X | ✓ | X | ✓ | P | ✓ | ✓ | ✓ |
| ***Ps. syringar pv. syringae* B301D** | - | - | - | - | - | - | R | ✓ | P | - | X | - | - | X | ✓ | X | ✓ | P | ✓ | C2 | ✓ |
| ***Ps. cerasi* PL963** | - | - | - | - | - | - | R | ✓ | P | - | - | - | - | - | ✓ | X | ✓ | P | ✓ | C2 | ✓ |
| ***Ps. syringae pv. syringae* Pss9097** | - | - | - | - | - | - | R | ✓ | P | ✓ | X | ✓ | ✓ | X | ✓ | X | ✓ | P | ✓ | C2 | ✓ |
| ***Ps. amygdali pv. morsprunorum* R15244** | - | - | - | - | - | - | R | ✓ | P | ✓ | - | ✓ | ✓ | X | ✓ | X | ✓ | P | ✓ | C2 | ✓ |

**Figure S3.** Domain organization of various *ppp* BGCs belonging to *Pseudovibrio* spp. and *Pseudomonas* spp.. Summary of antiSMASH results**.** The single letter amino acid code is indicated below each A domain. If the amino acid could not be predicted, it is indicated with “X”. Domain key as depicted on Fig. 1. C2, inactive KR domain. *, additional NRPS module composed of C-A-PCP coding for G.

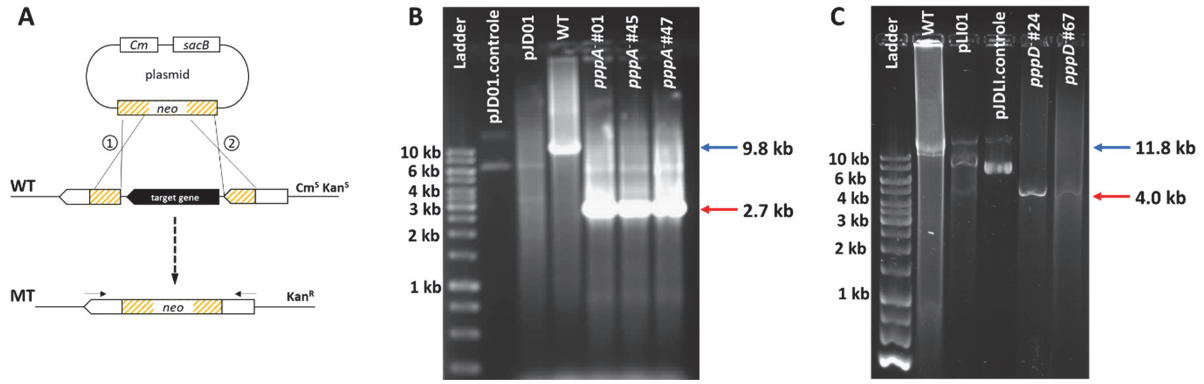

**Figure S4.** *pppA* and *pppD* gene knockout. A) Design of gene replacement experiments. *neo*, neomycin/kanamycin resistance marker. Cm, chloramphenicol resistance gene. $Kan^{R/S}$, kanamycin resistance / sensitivity. $Cm^{R/S}$, chloramphenicol resistance / sensitivity. Numbers 1 and 2 indicate crossover events. Black arrows indicate primer location. WT, wild-type. MT, mutant. B) Analysis of *pppA* double crossover mutants by PCR. Expected fragment length for MTs, 2.7 kb. Expected fragment length for the WT, 9.8 kb. C) Analysis of *pppD* double crossover mutants by PCR. Expected fragment length of 11.8 kb (WT) and 4.0 kb (MT). Ladder, molecular weight DNA ladder (GeneRuler 1kb DNA Ladder - ThermoScientific).

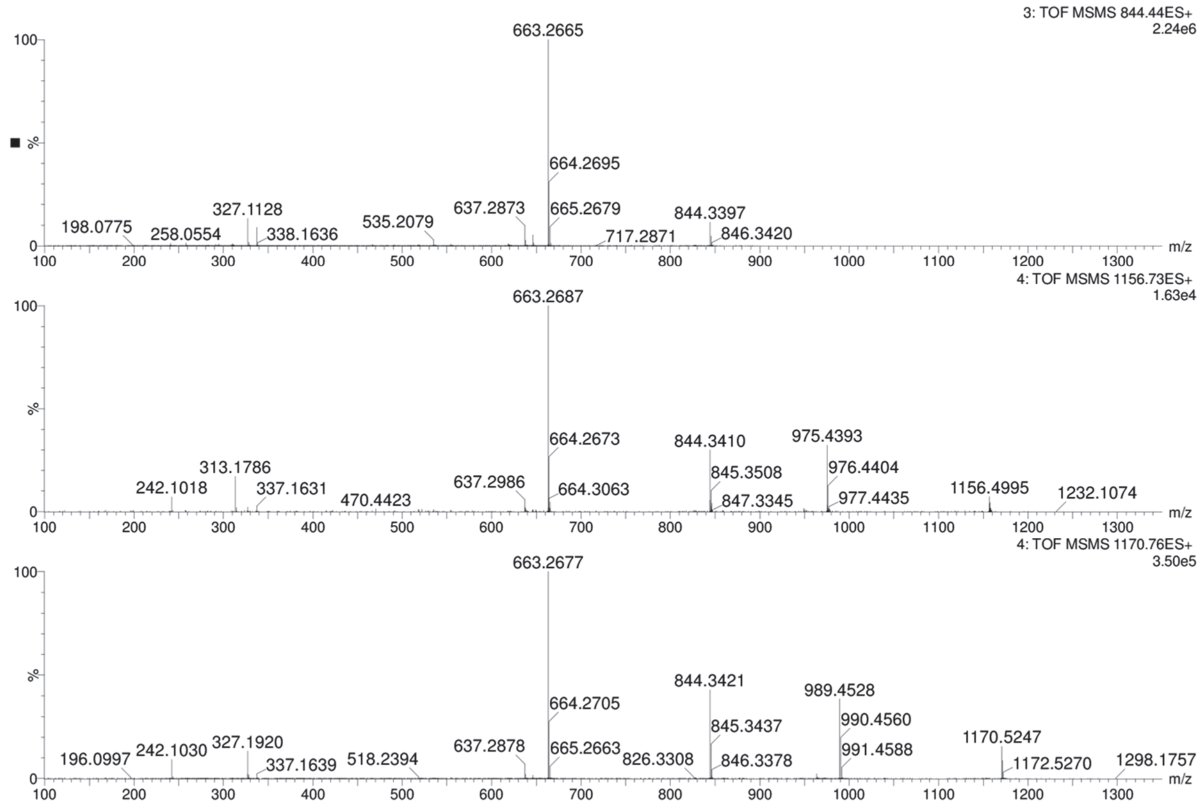

**Figure S5.** Mass spectra MS$^2$ of compounds encoded by the *ppp* BGC observed on QTOF-MS analysis of the crude extract of *P. brasiliensis* Ab134: 844.3 (top), 1156.5 (middle) and 1170.5 (bottom).

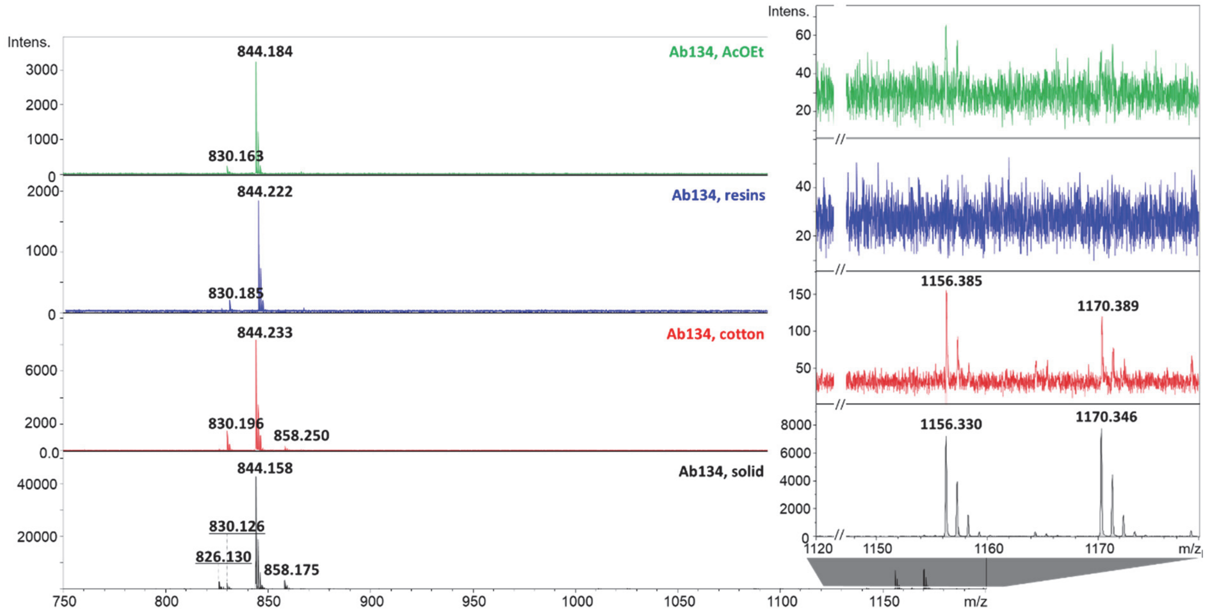

**Figure S6.** Comparative metabolite analysis of different culture extracts of *P. brasiliensis* Ab134 using MALDI-TOF mass spectrometry: liquid cultures extracted with AcOEt (green, top), resins added to liquid cultures and extracted with methanol (blue), cotton balls placed in liquid cultures and later extracted with methanol (red), methanol extract of cultures grown on solid media, i.e. swarming assay conditions (black, bottom).

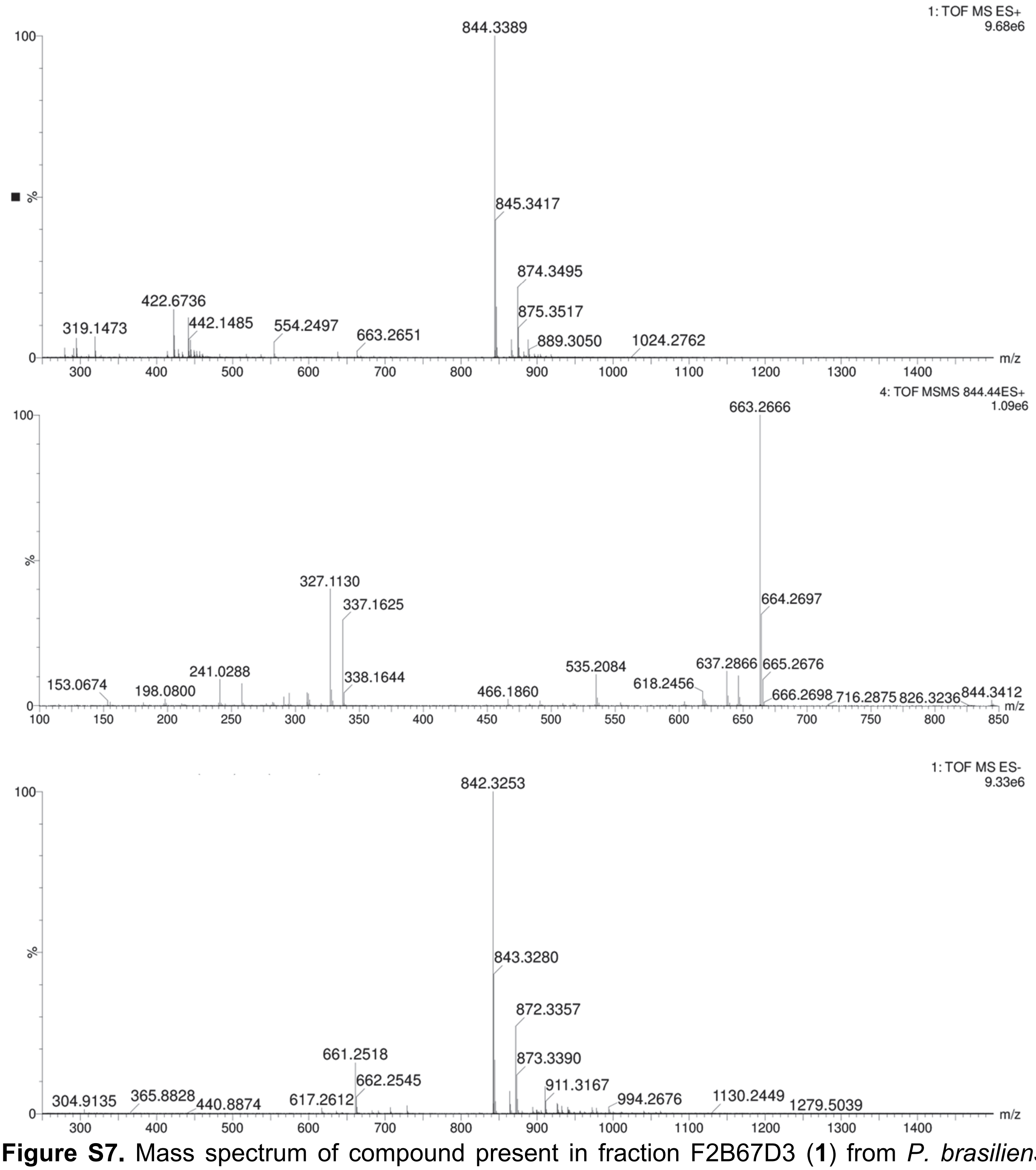

**Figure S7.** Mass spectrum of compound present in fraction F2B67D3 (**1**) from *P. brasiliensis* Ab134. QToF-MS in positive mode (top), $MS^2$ in positive mode (middle) and MS in negative mode (bottom).

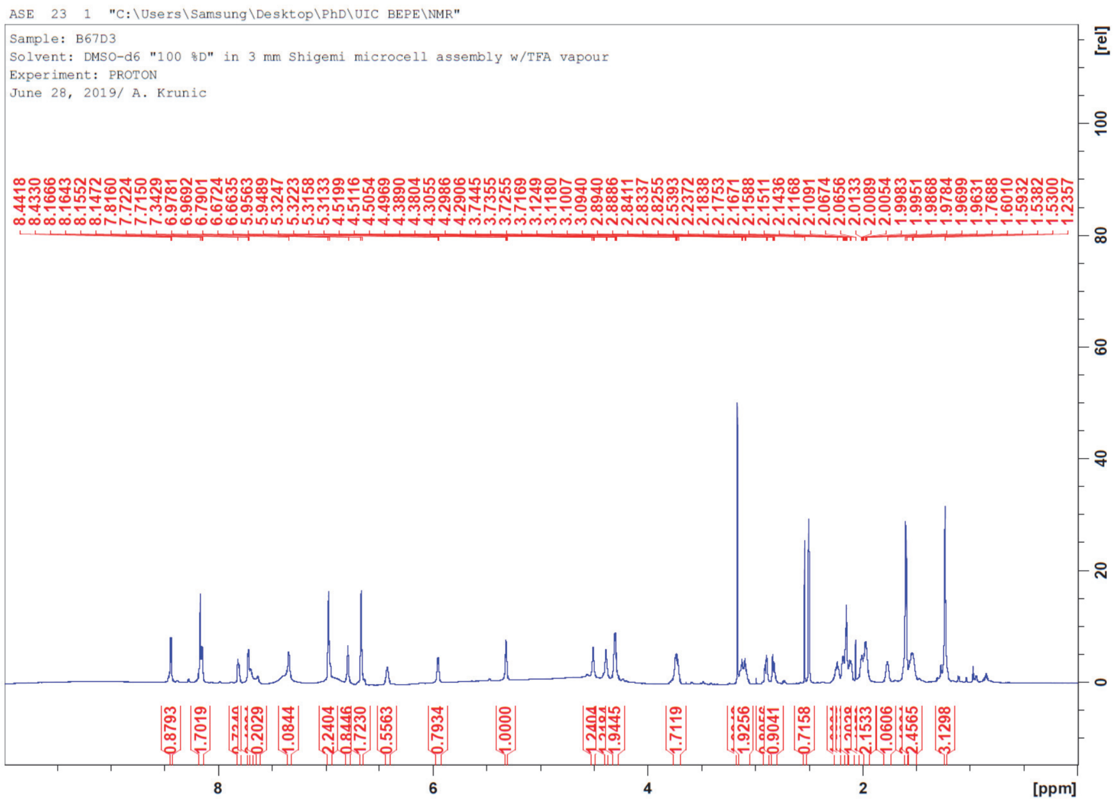

**Figure S8.** $^{1}$H NMR spectrum of **1**, fraction F2B67D3 (900 MHz, DMSO-$d_6$ + TFA vapor).

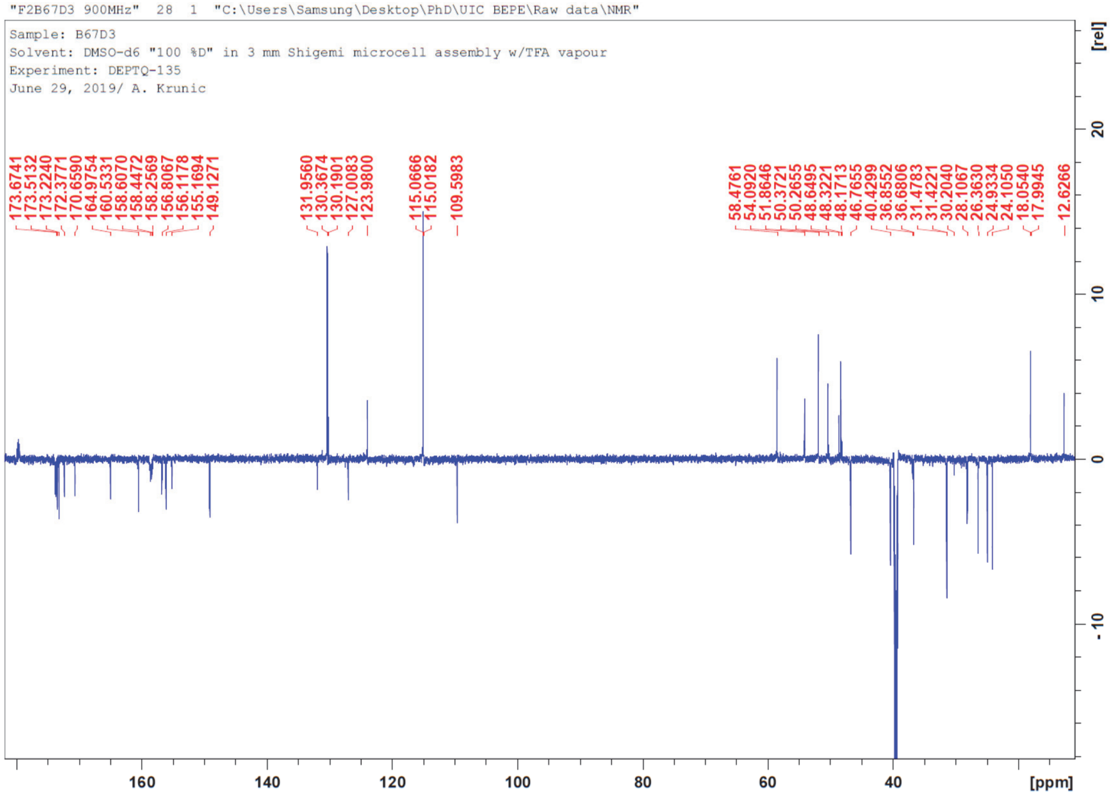

**Figure S9.** $^{13}$C DEPTQ NMR spectrum of **1**, fraction F2B67D3 (226 MHz, DMSO-$d_6$ + TFA vapor).

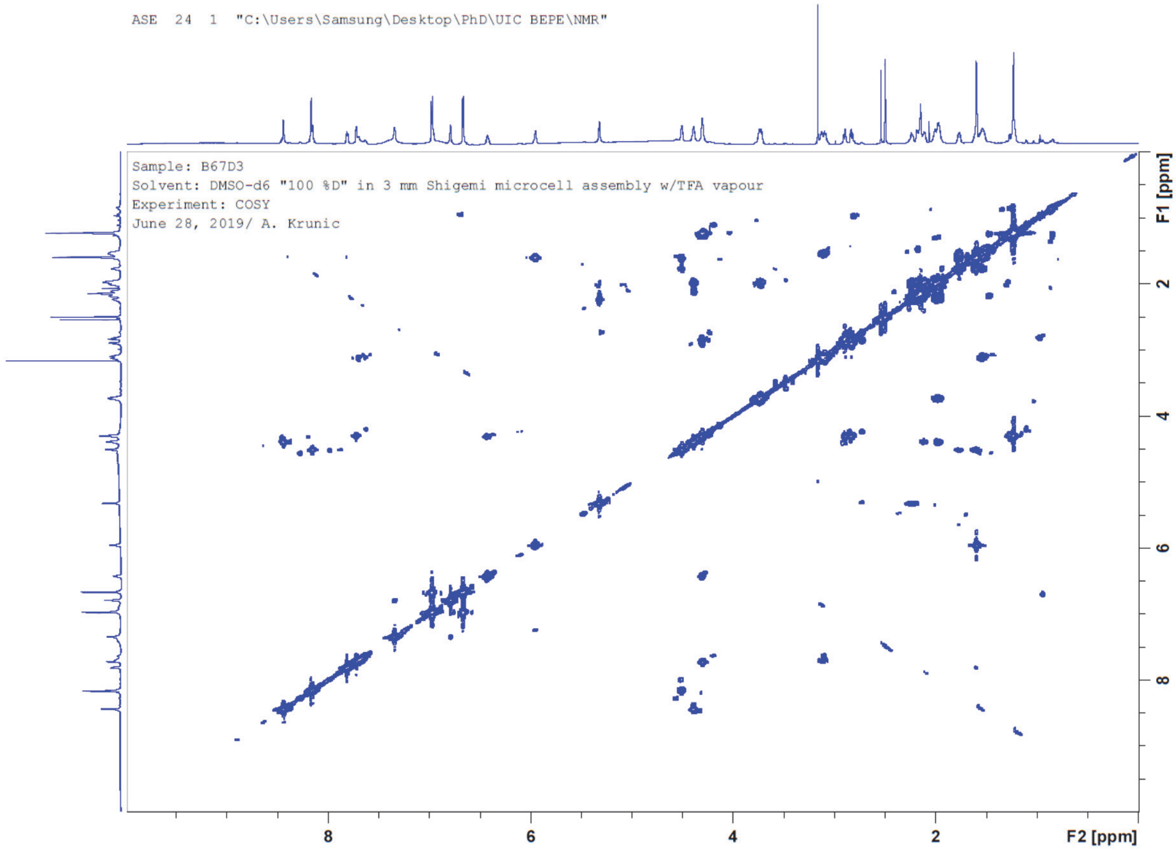

**Figure S10.** COSY NMR spectrum of **1**, fraction F2B67D3 (900 MHz, DMSO-$d_6$ + TFA vapor).

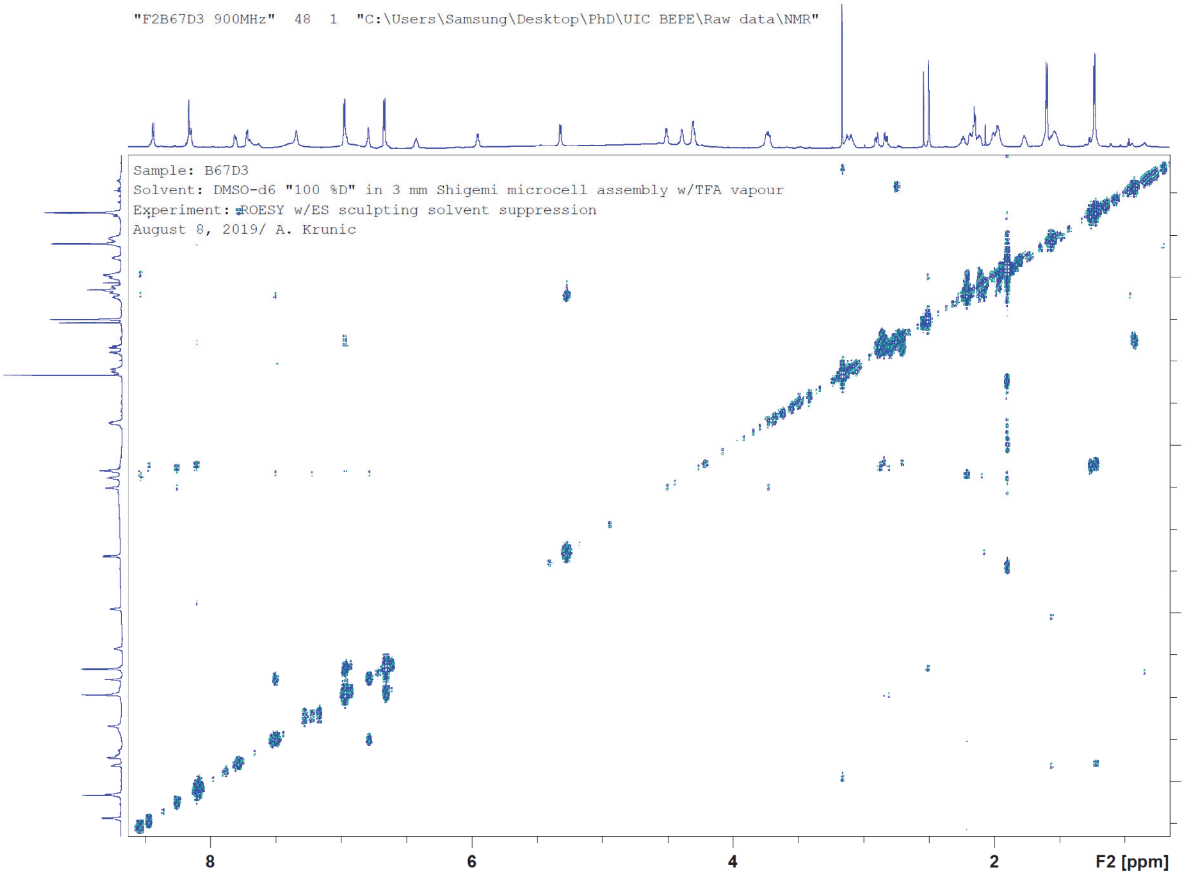

**Figure S11.** ROESY NMR spectrum of **1**, fraction F2B67D3 (900 MHz, DMSO-$d_6$ + TFA vapor).

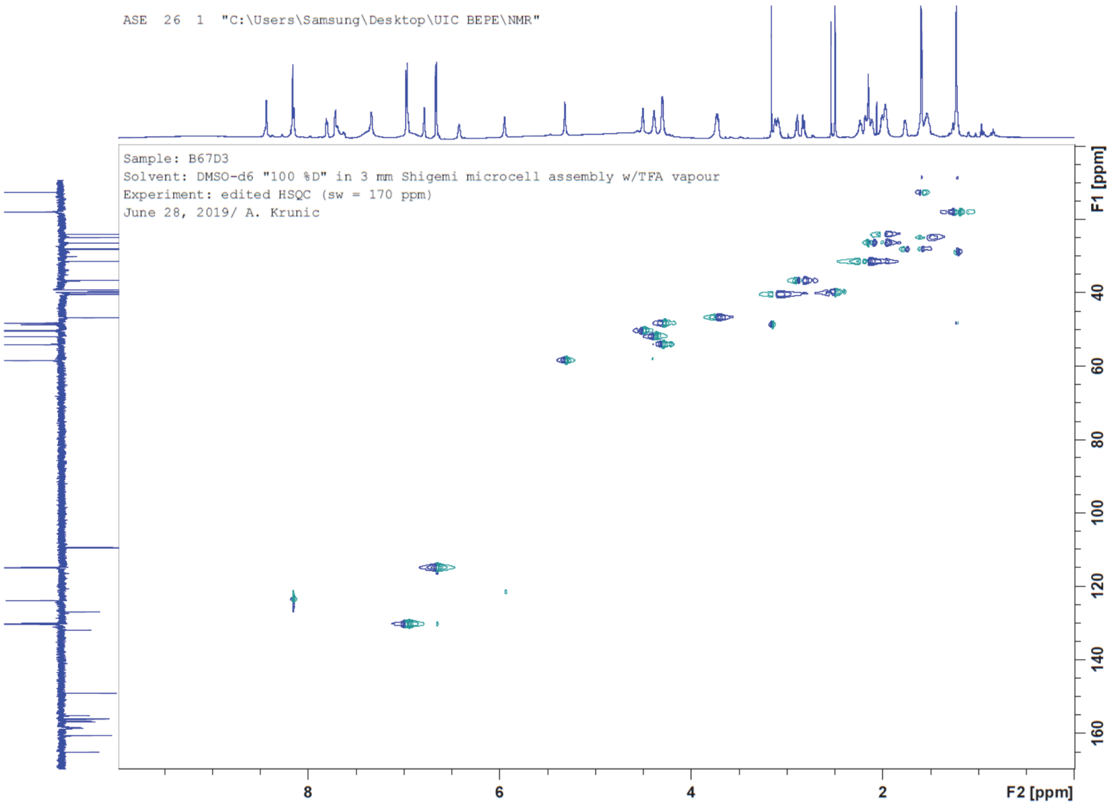

**Figure S12.** HSQC NMR spectrum of **1**, fraction F2B67D3 (226 MHz:900 MHz, DMSO-$d_6$ + TFA vapor).

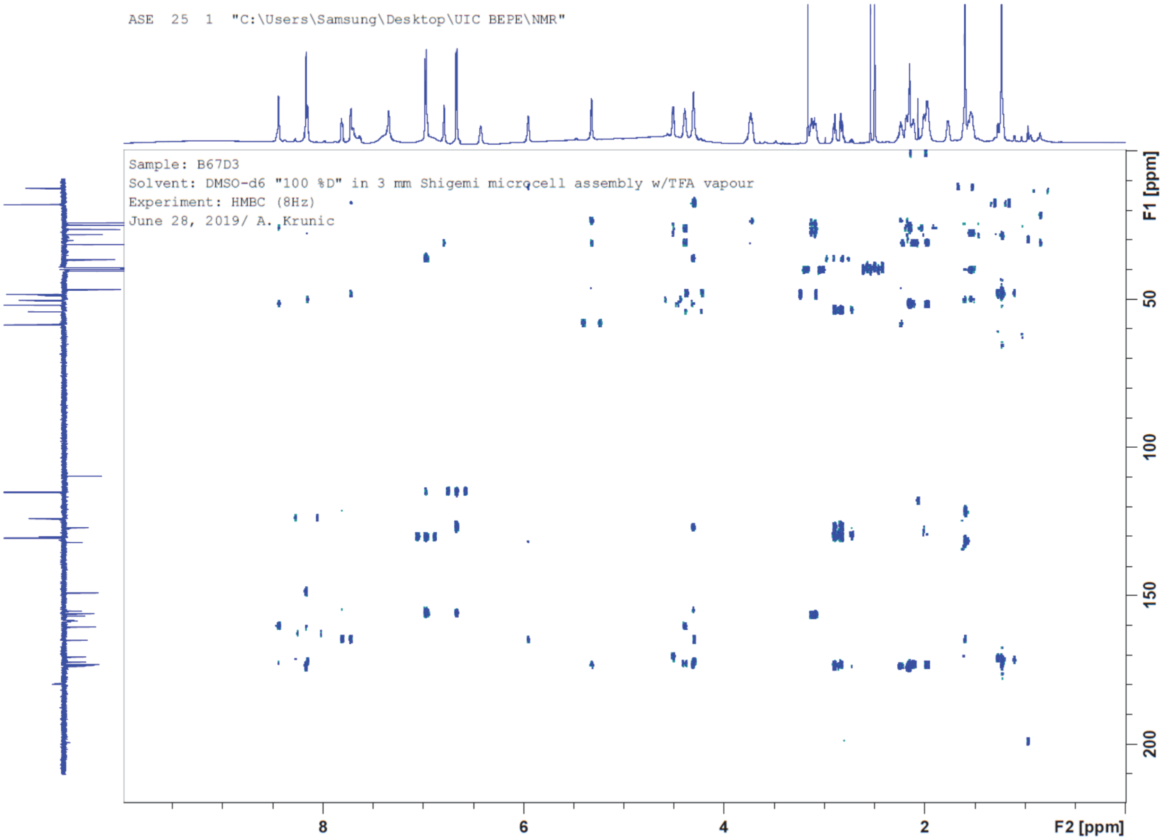

**Figure S13.** HMBC NMR spectrum of **1**, fraction F2B67D3 (226 MHz:900 MHz, DMSO-$d_6$ + TFA vapor).

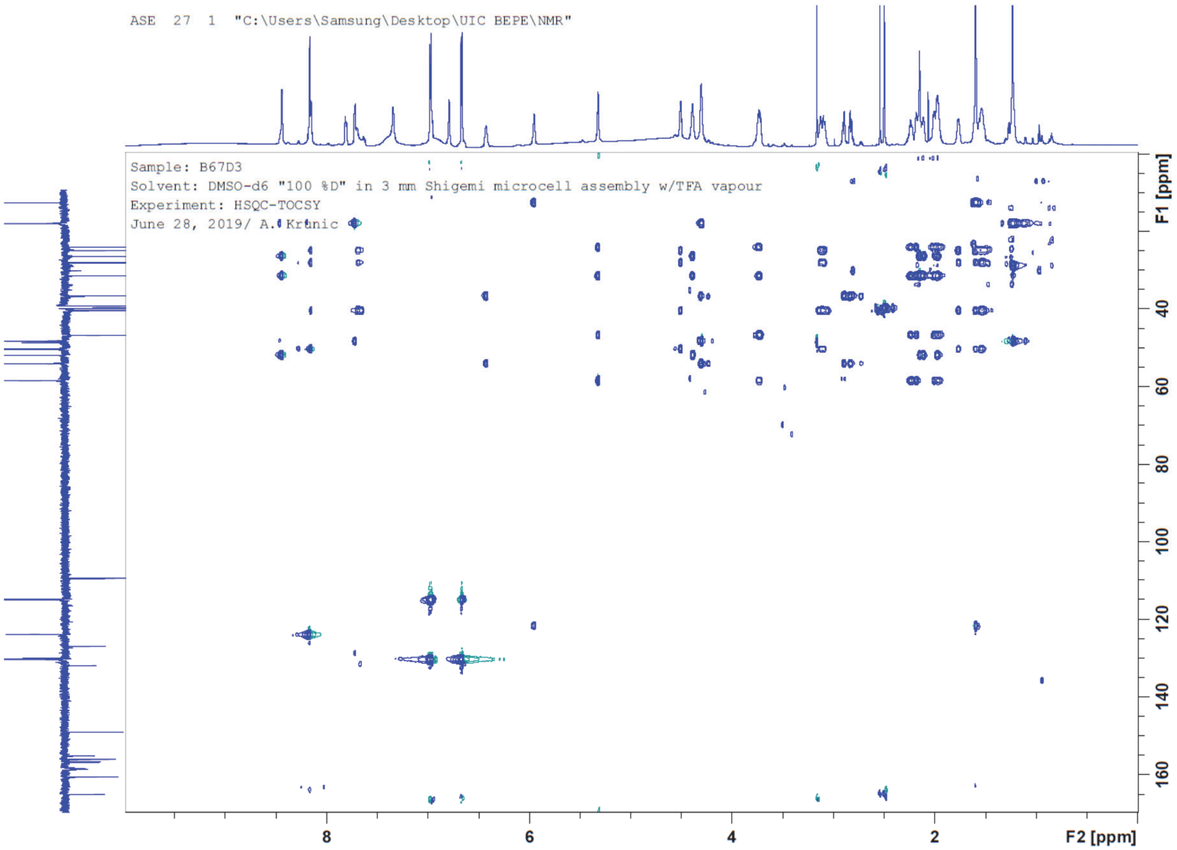

**Figure S14.** HSCQ-TOCSY NMR spectrum of **1**, fraction F2B67D3 (226 MHz:900 MHz, DMSO-$d_6$ + TFA vapor).

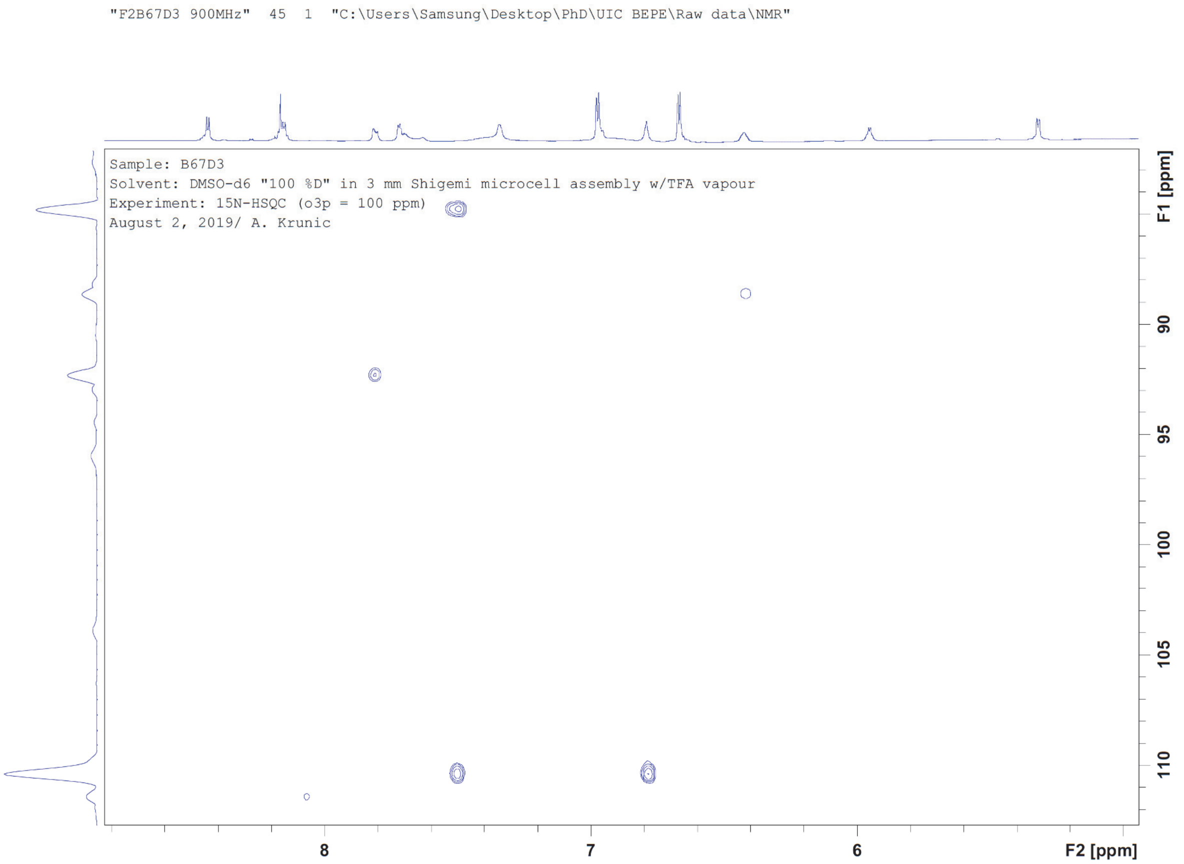

**Figure S15.** $^{15}$N NMR spectrum of **1**, fraction F2B67D3 (91 MHz:900 MHz, DMSO-$d_6$ + TFA vapor).

**Figure S16**. Key $^{1}$H, $^{13}$C and $^{15}$N NMR chemical shifts and HMBC, COSY and NOE/ROE correlations observed for compound **1**.

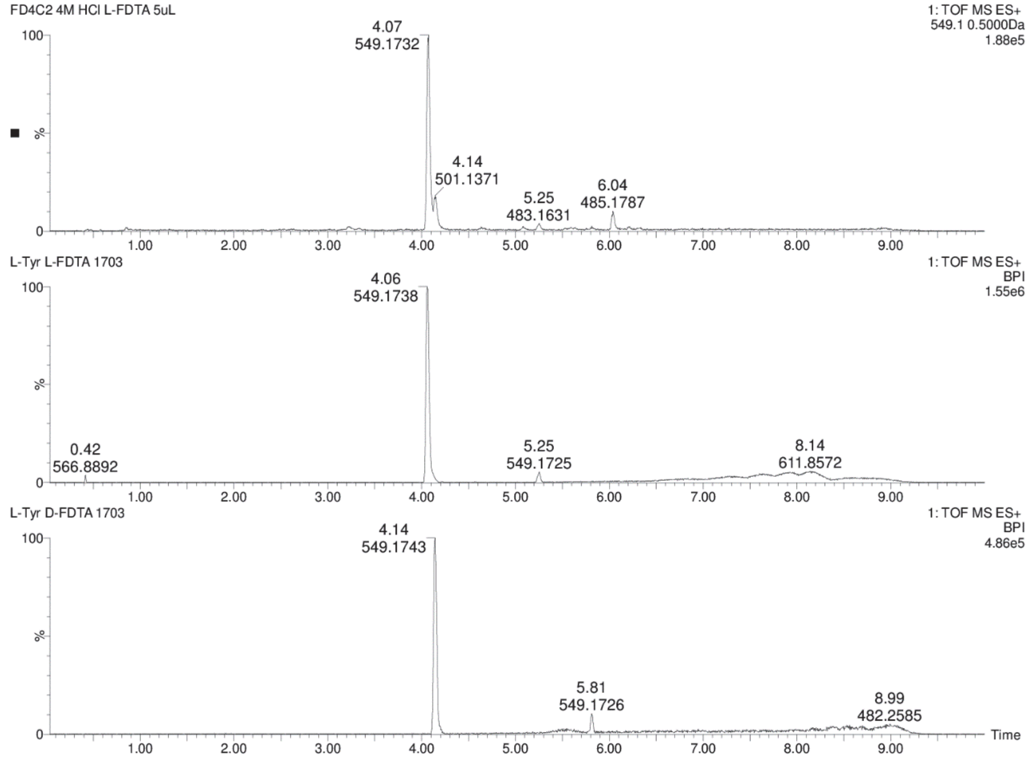

**Figure S17.** Chromatograms for comparison of **1** + L-FDTA (top) and L-Tyr + L-FDTA (middle) and L-Tyr + D-FDTA (bottom) in UPLC-MS analysis.

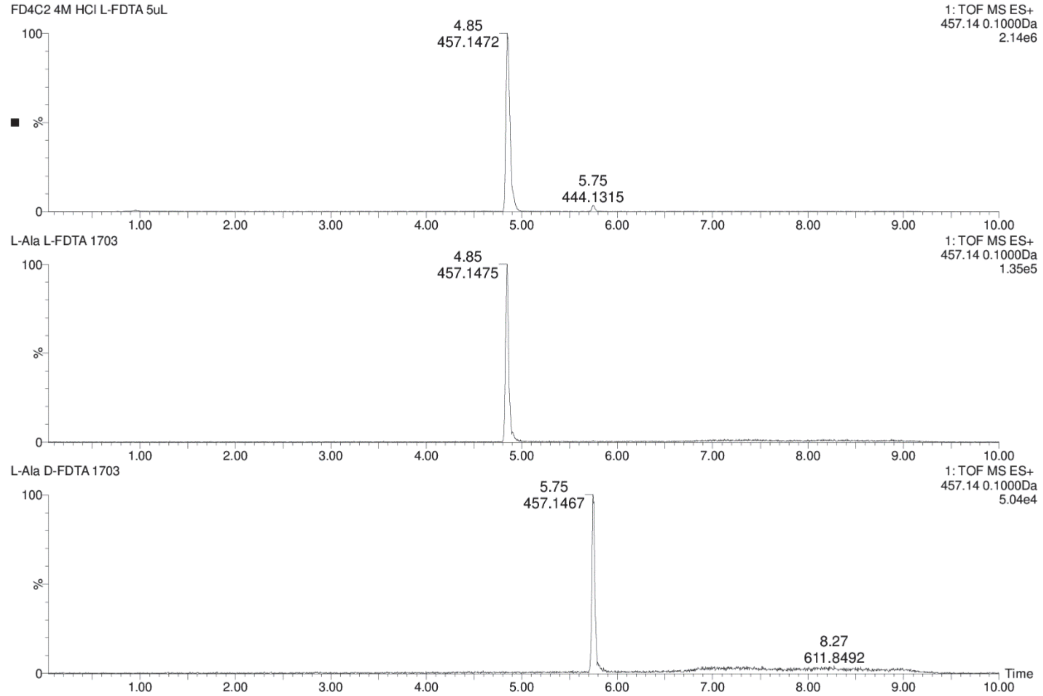

**Figure S18.** Chromatograms for comparison of **1** + L-FDTA (top) and L-Ala + L-FDTA (middle) and L-Ala + D-FDTA (bottom) in UPLC-MS analysis.

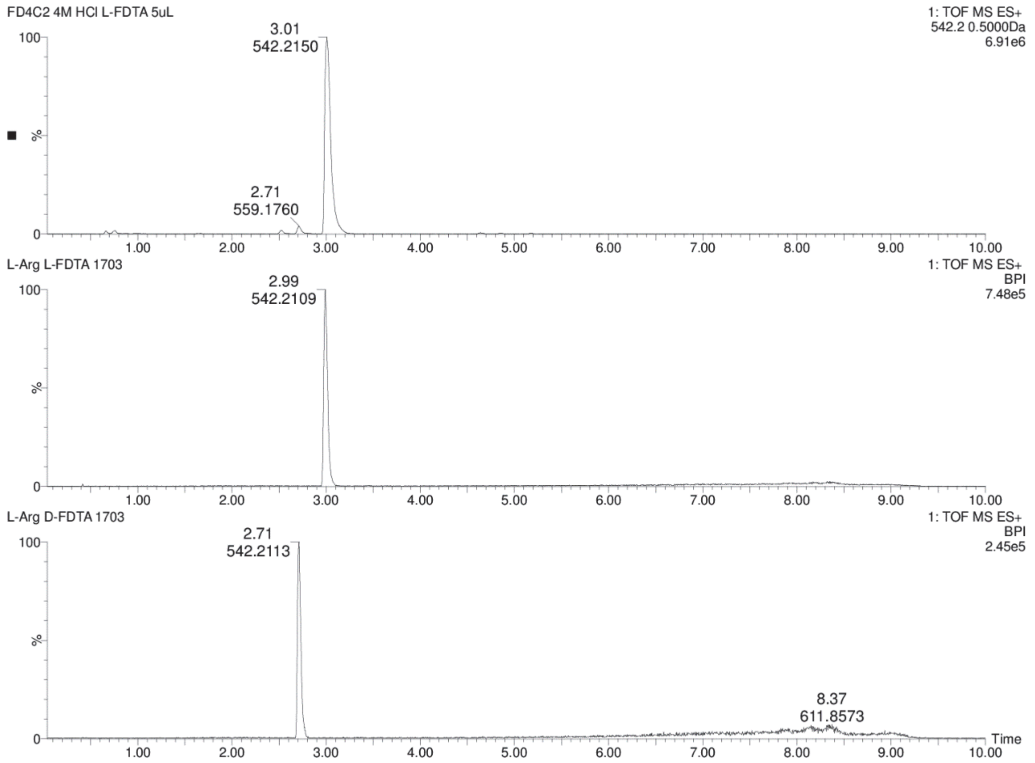

**Figure S19.** Chromatograms for comparison of **1** + L-FDTA (top) and L-Arg + L-FDTA (middle) and L-Arg + D-FDTA (bottom) in UPLC-MS analysis.

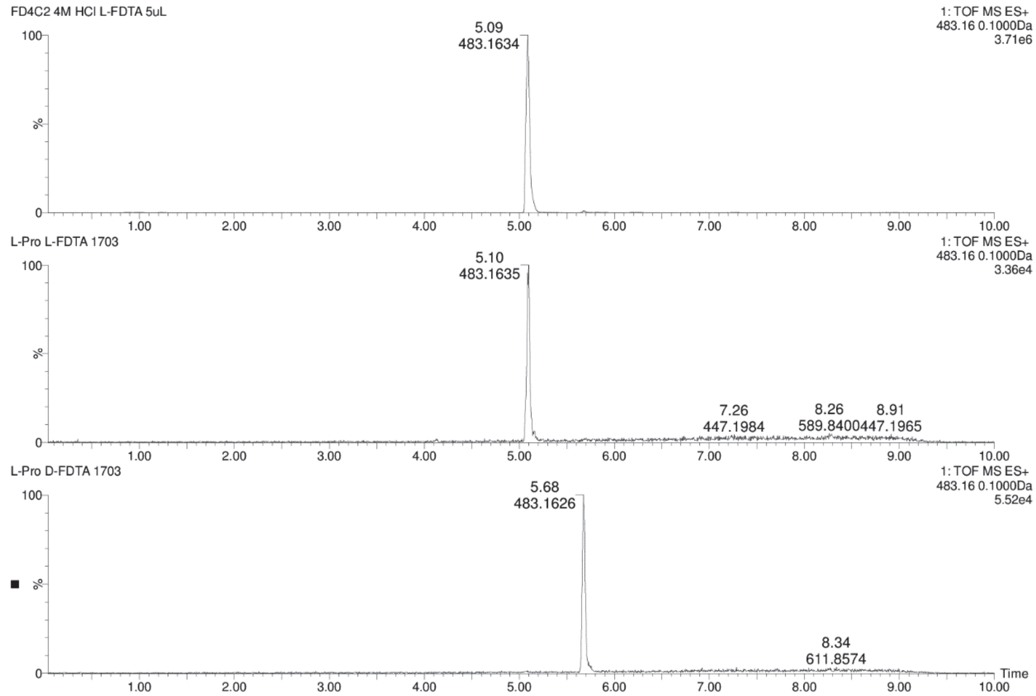

**Figure S20.** Chromatograms for comparison of **1** + L-FDTA (top) and L-Pro + L-FDTA (middle) and L-Pro + D-FDTA (bottom) in UPLC-MS analysis.

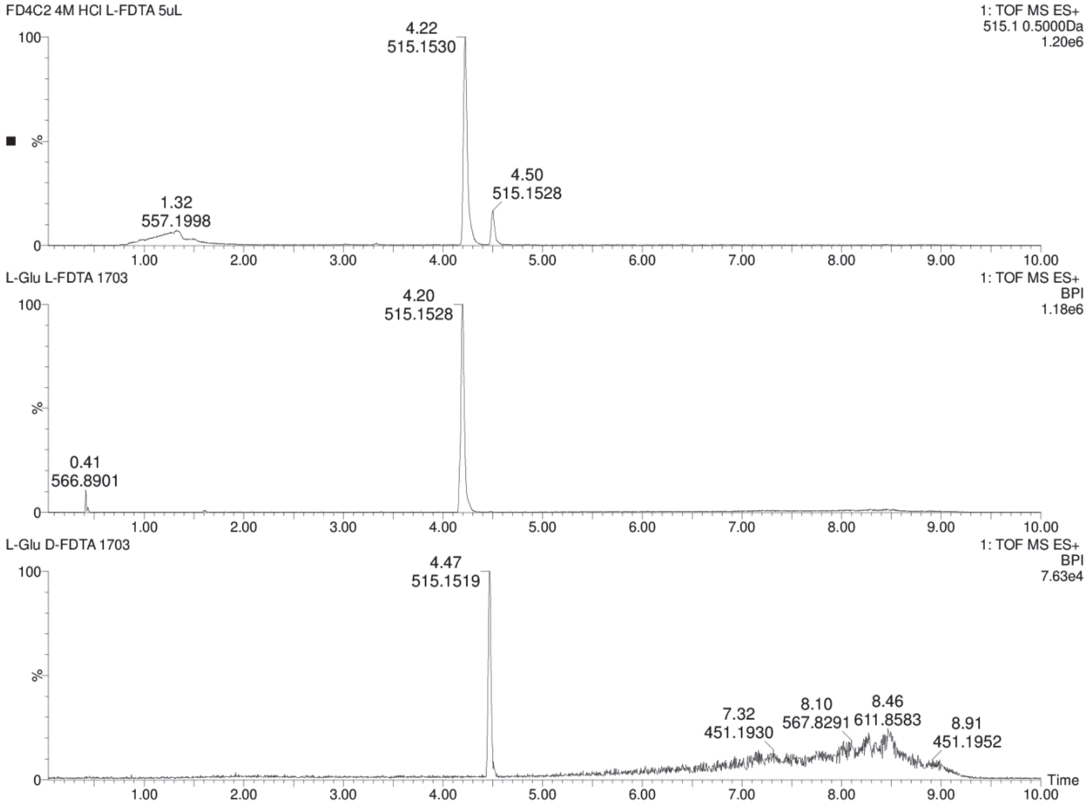

**Figure S21.** Chromatograms for comparison of **1** + L-FDTA (top) and L-Glu + L-FDTA (middle) and L-Glu + D-FDTA (bottom) in UPLC-MS analysis.

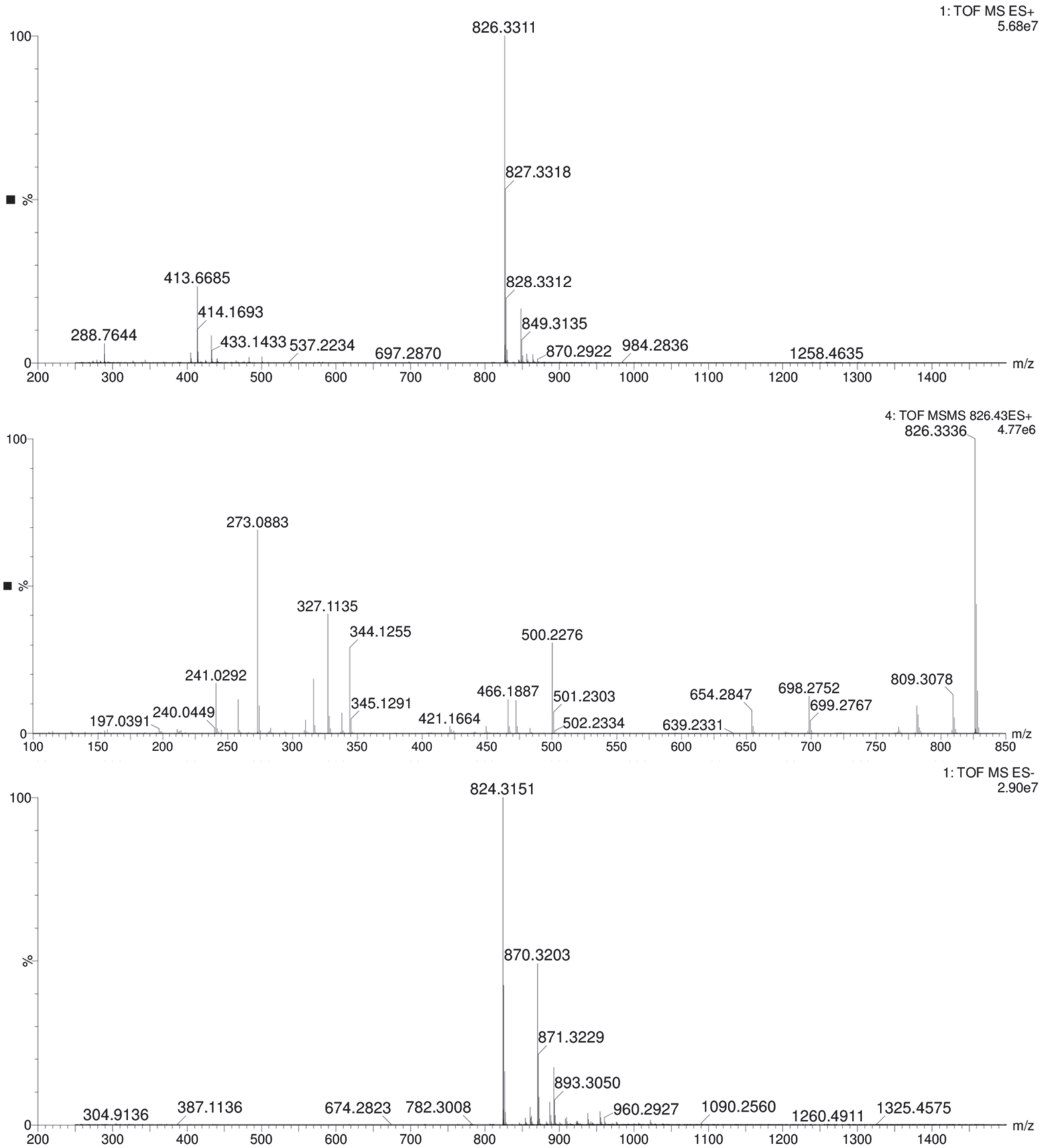

**Figure S22.** Mass spectrum of compound present in fraction F2B67D1/FD2B3D1 (**2**) from *P. brasiliensis* Ab134. QToF-MS in positive mode (top), $MS^2$ in positive mode (middle) and MS in negative mode (bottom).

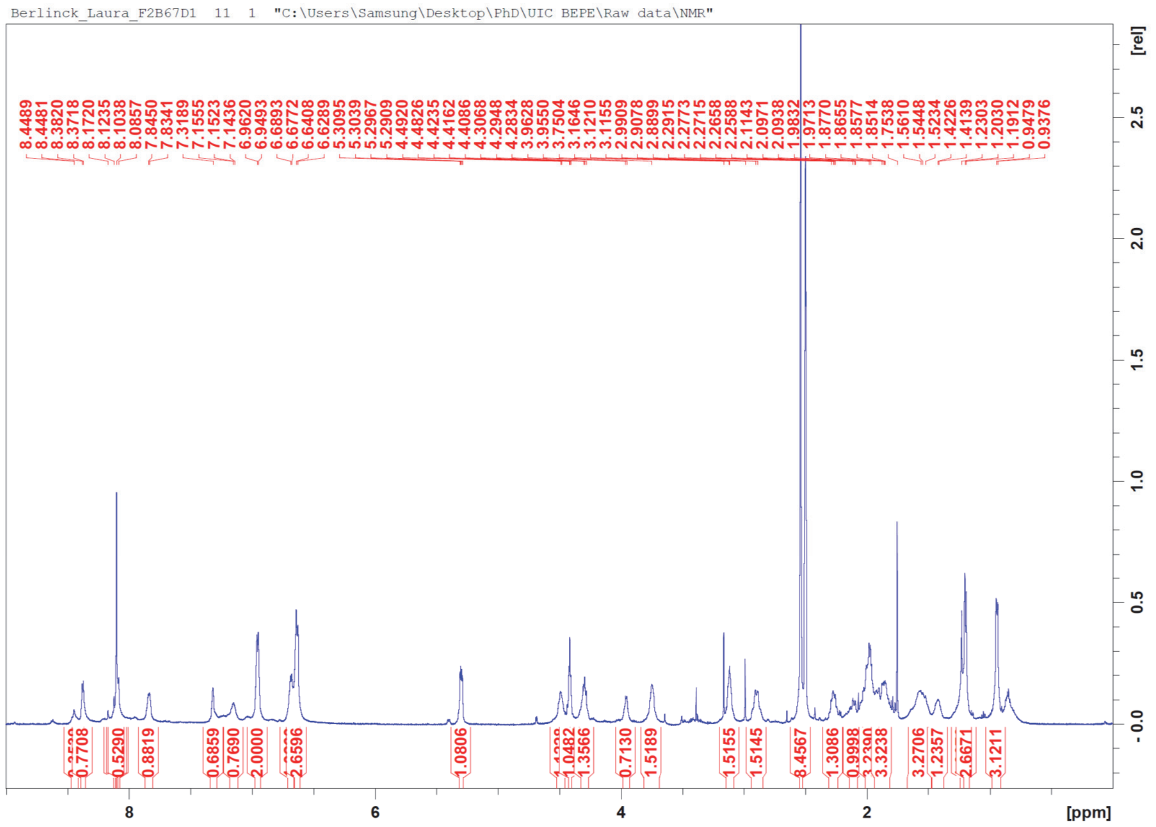

**Figure S23.** [1]H NMR spectrum of **2**, fraction F2B67D1/FD2B3D1 (600 MHz, DMSO-$d_6$ + TFA vapor).

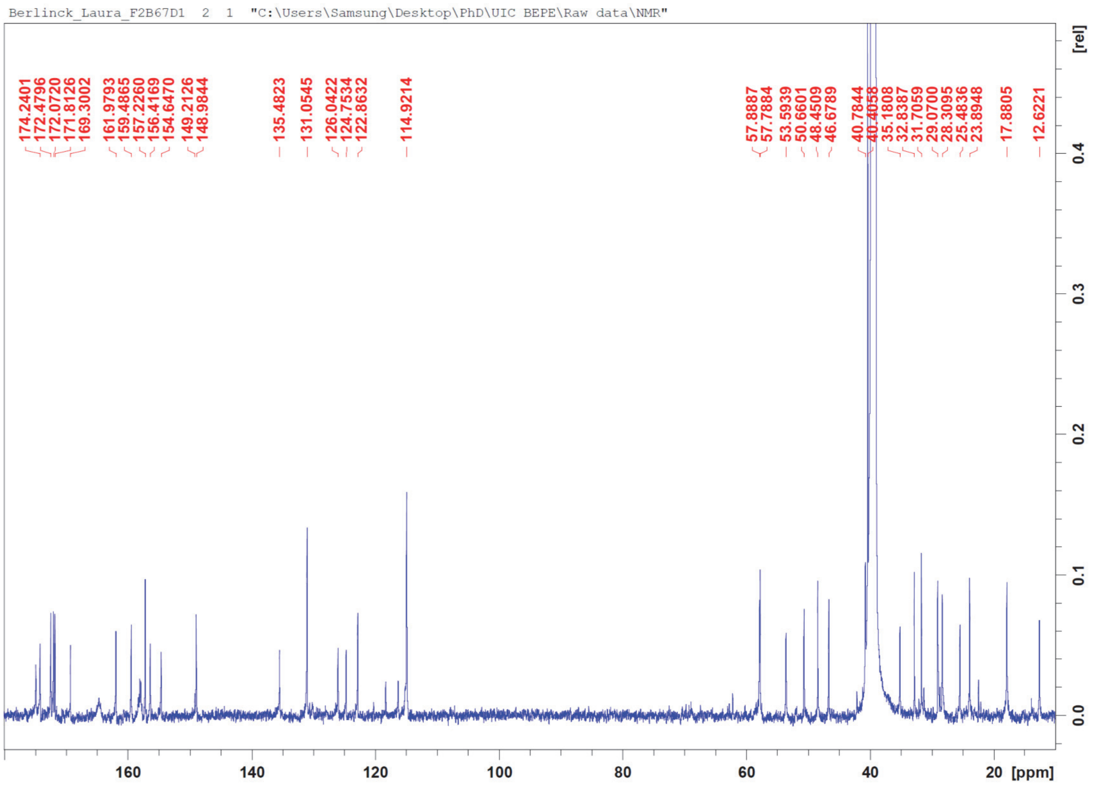

**Figure S24.** $^{13}$C NMR spectrum of **2**, fraction F2B67D1/FD2B3D1 (150 MHz, DMSO-$d_6$ + TFA vapor).

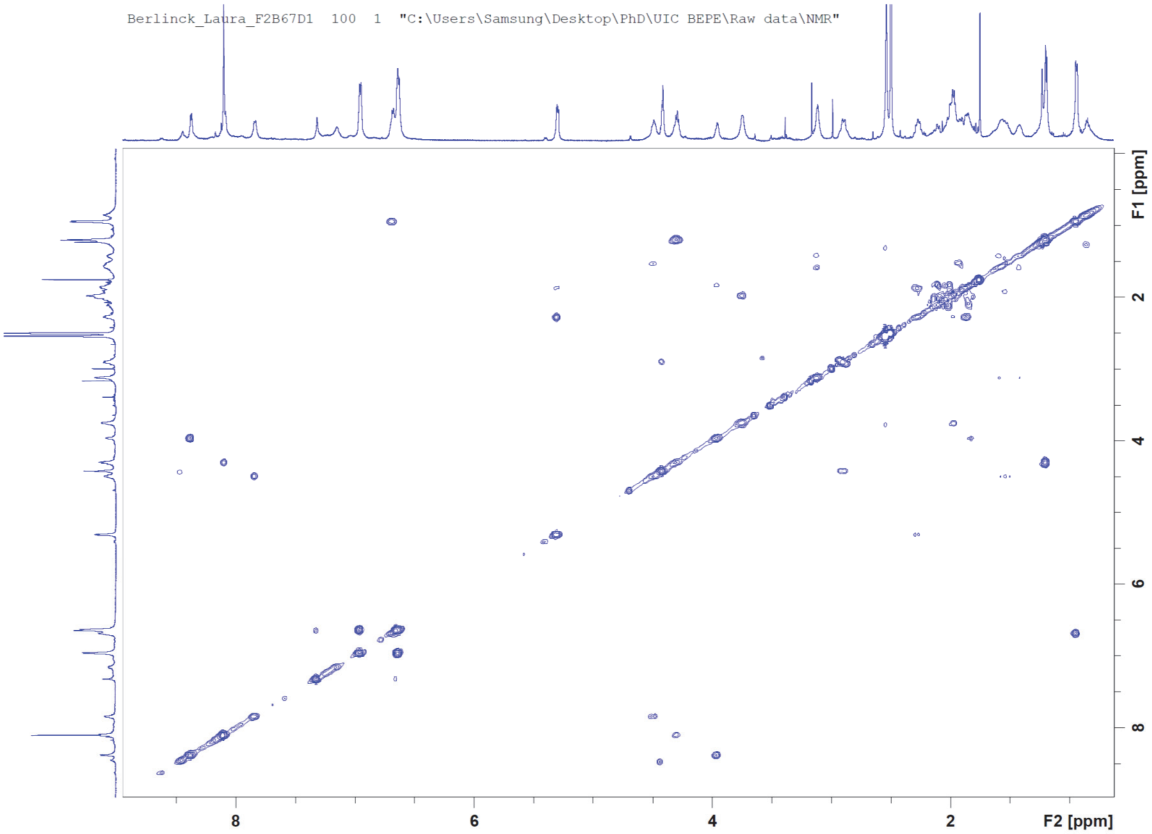

**Figure S25.** COSY NMR spectrum of **2**, fraction F2B67D1/FD2B3D1 (600 MHz, DMSO-$d_6$ + TFA vapor).

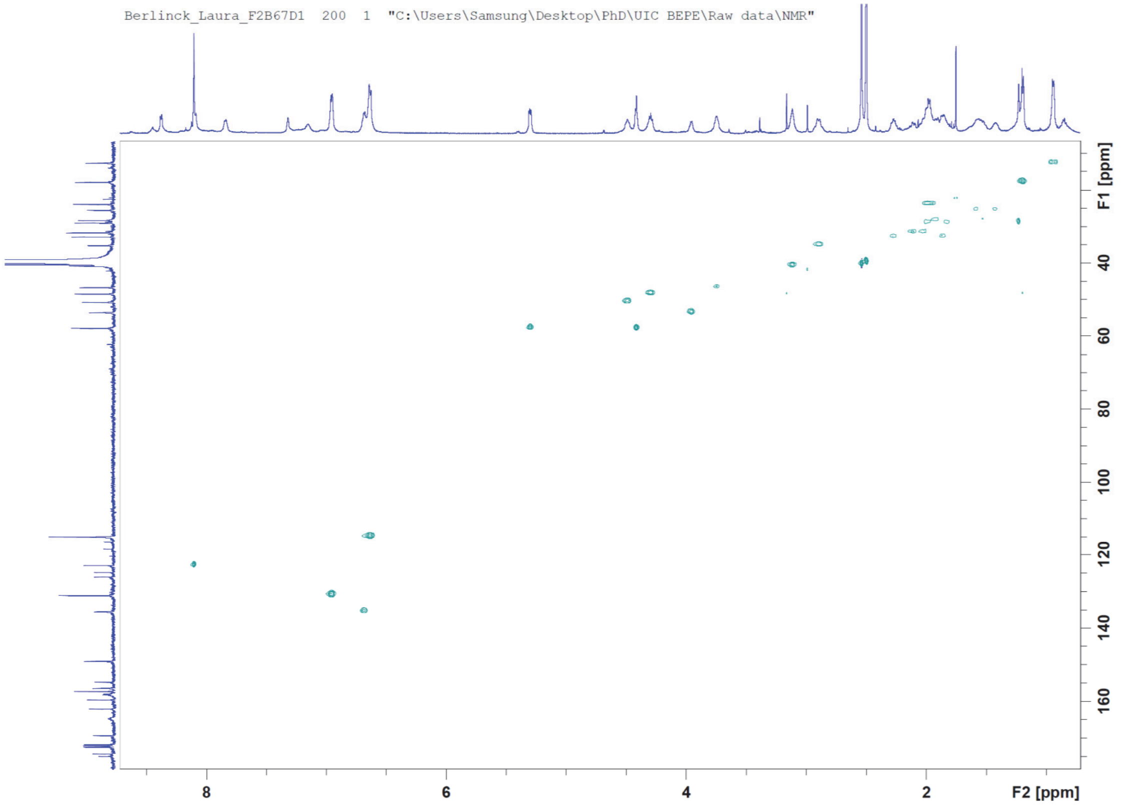

**Figure S26.** HSQC NMR spectrum of **2**, fraction F2B67D1/FD2B3D1 (150 MHz:600 MHz, DMSO-$d_6$ + TFA vapor).

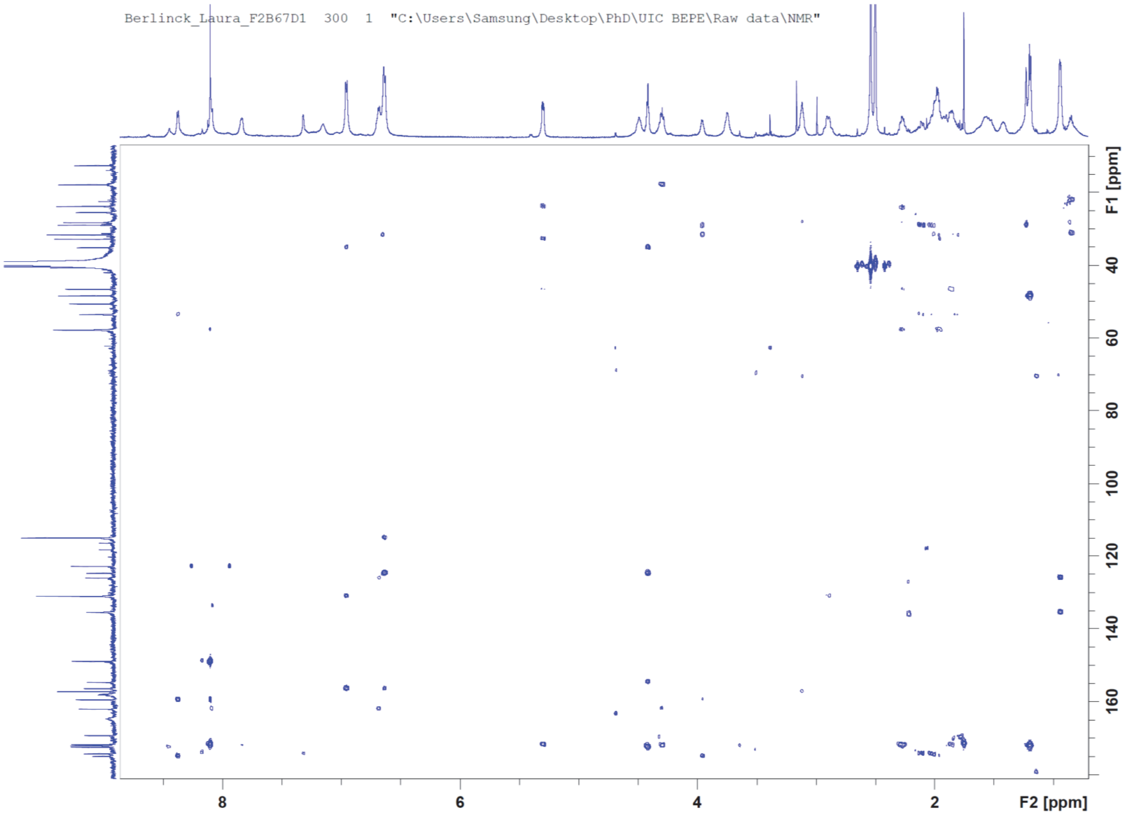

**Figure S27.** HMBC NMR spectrum of **2**, fraction F2B67D1/FD2B3D1 (150 MHz:600 MHz, DMSO-$d_6$ + TFA vapor).

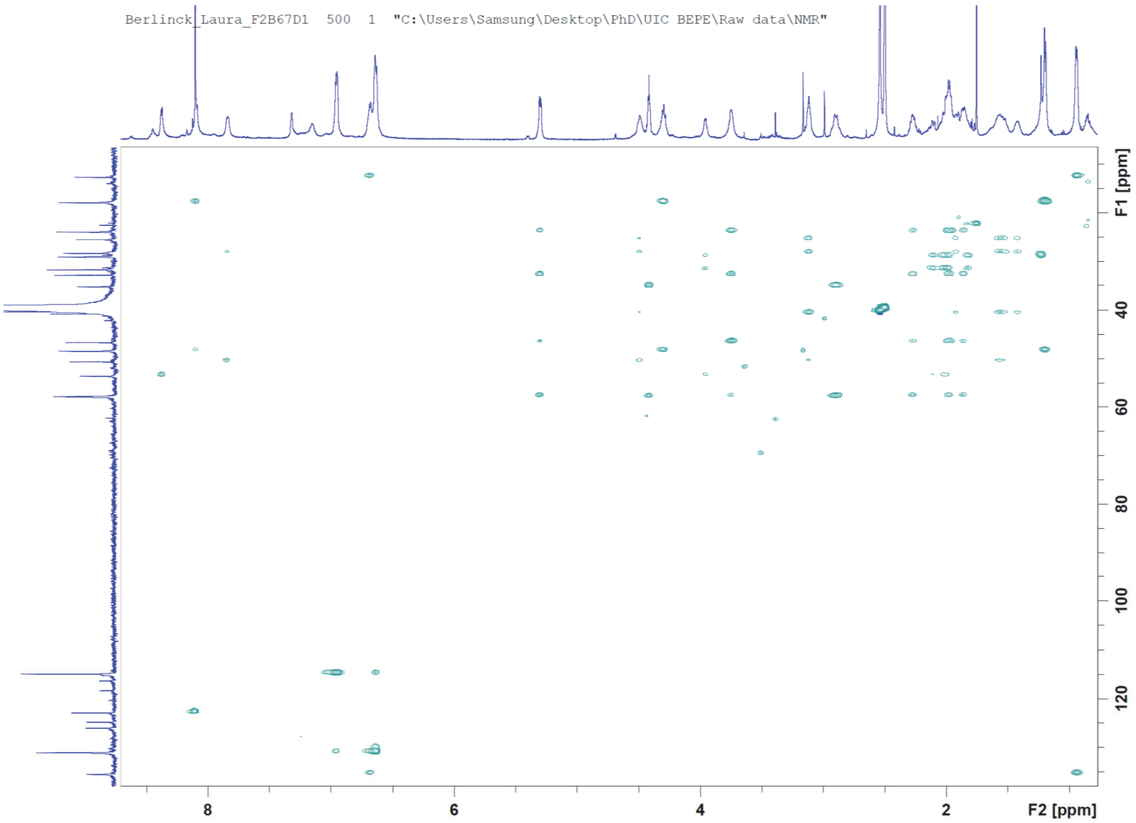

**Figure S28.** HSQC-TOCSY NMR spectrum of **2**, fraction F2B67D1/FD2B3D1 (150 MHz:600 MHz, DMSO-$d_6$ + TFA vapor).

**Figure S29**. Key $^1$H and $^{13}$C NMR chemical shifts and HMBC, COSY and NOE/ROE correlations observed for compound 2.

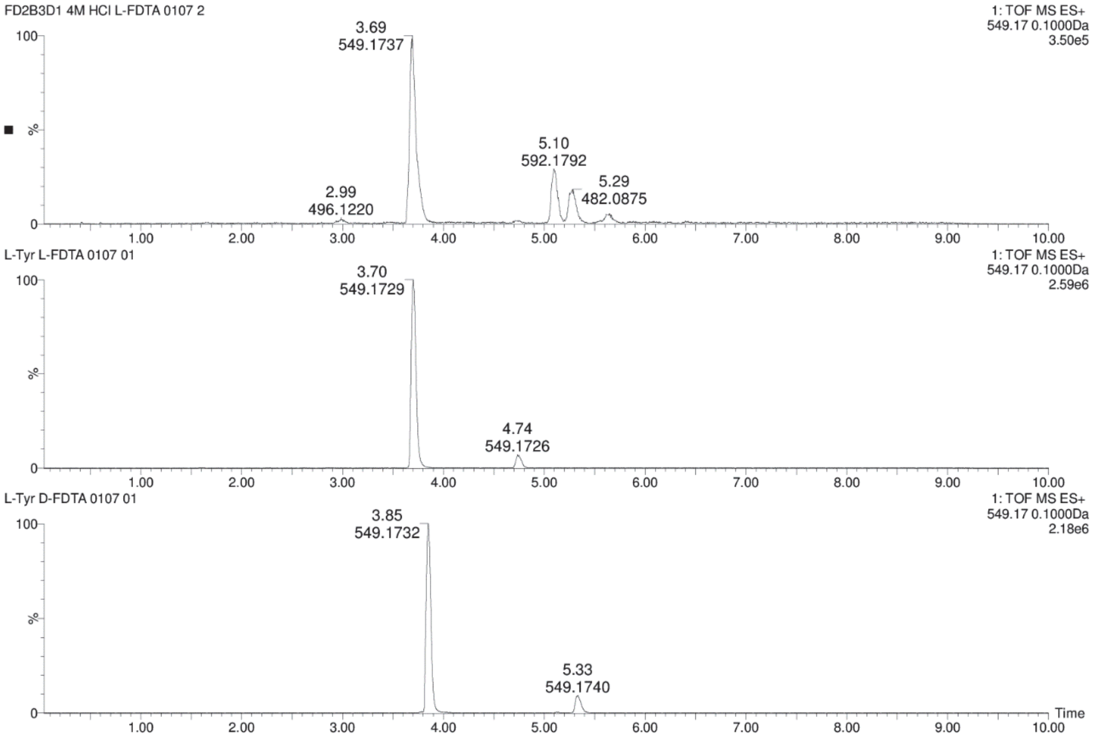

**Figure S30.** Chromatograms for comparison of **2** + L-FDTA (top) and L-Tyr + L-FDTA (middle) and L-Tyr + D-FDTA (bottom) in UPLC-MS analysis.

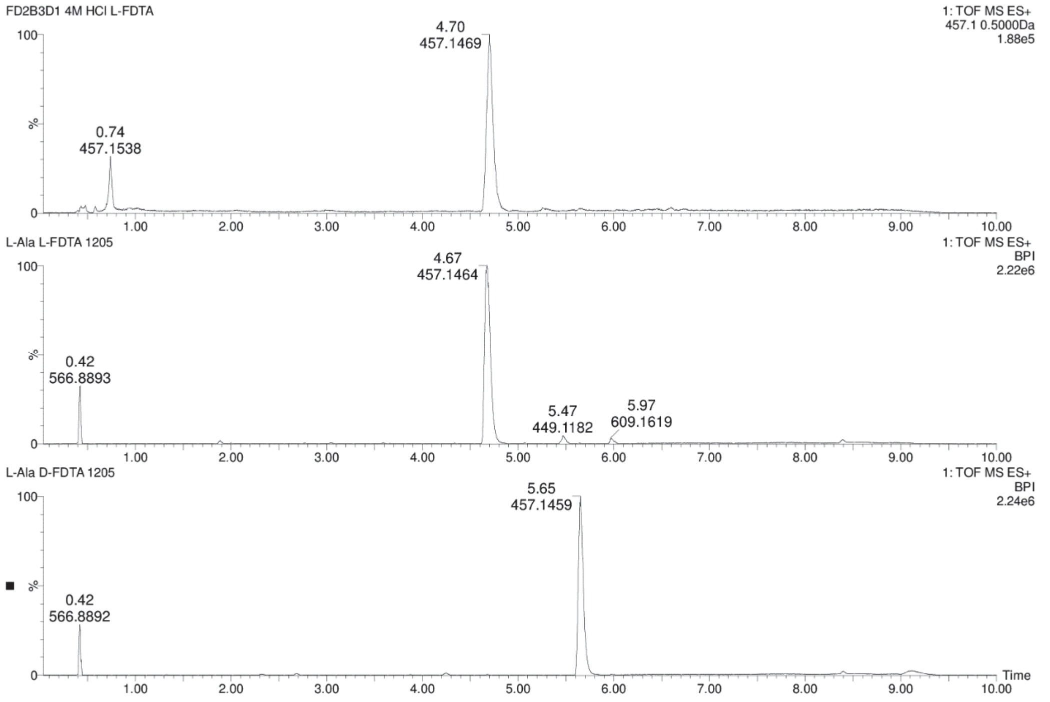

**Figure S31.** Chromatograms for comparison of **2** + L-FDTA (top) and L-Ala + L-FDTA (middle) and L-Ala + D-FDTA (bottom) in UPLC-MS analysis.

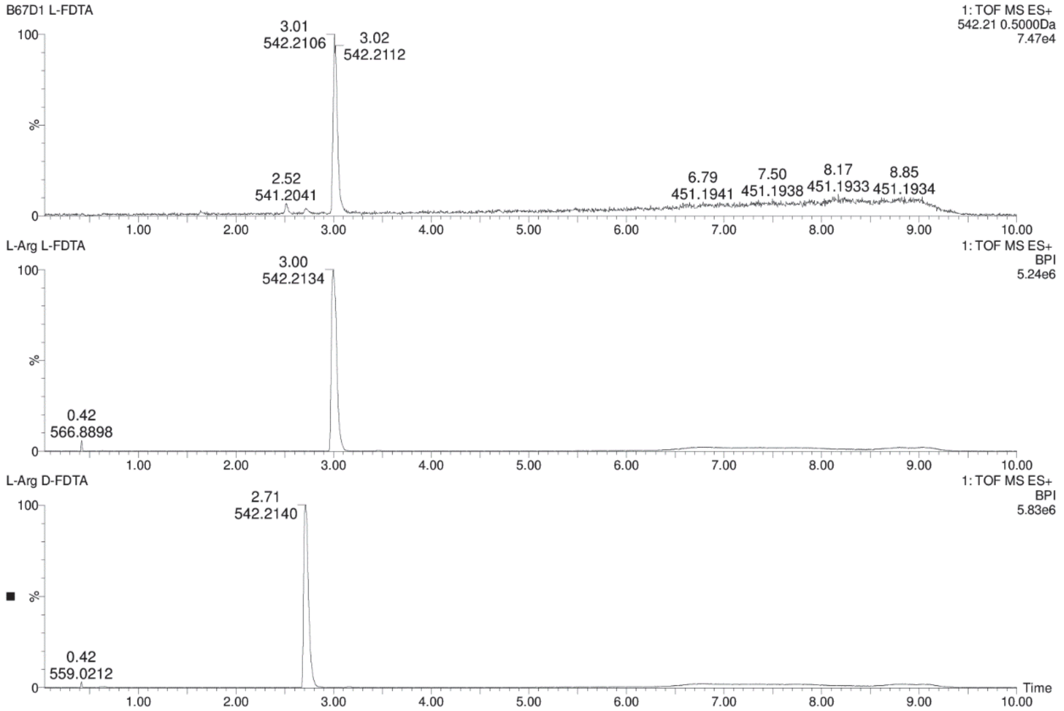

**Figure S32.** Chromatograms for comparison of **2** + L-FDTA (top) and L-Arg + L-FDTA (middle) and L-Arg + D-FDTA (bottom) in UPLC-MS analysis.

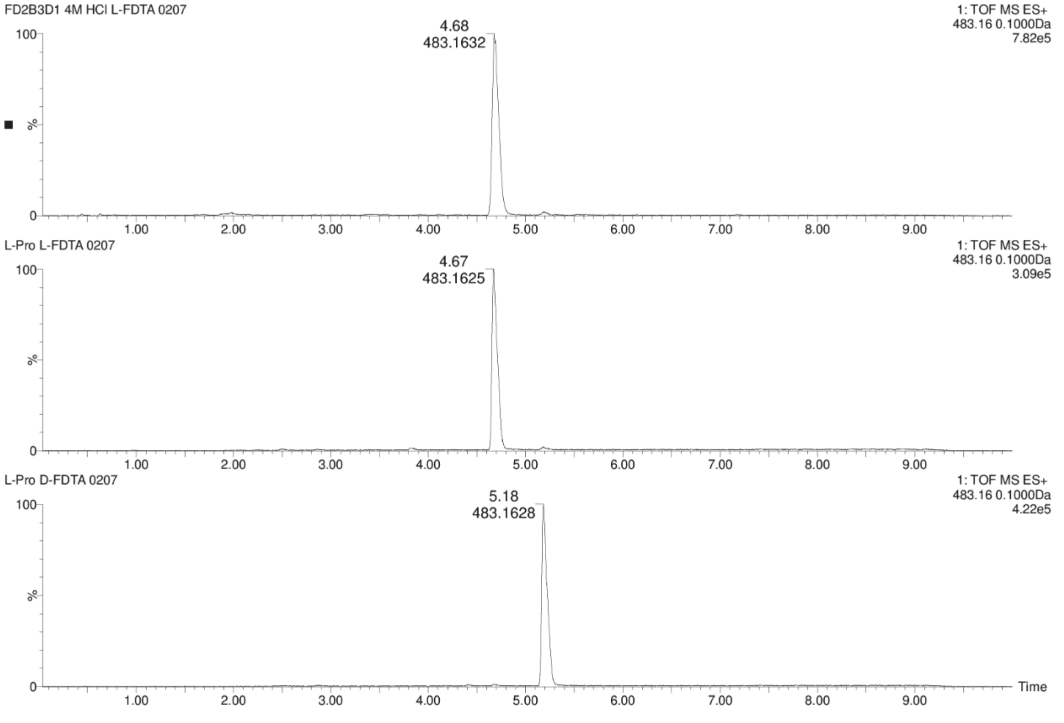

**Figure S33.** Chromatograms for comparison of **2** + L-FDTA (top) and L-Pro + L-FDTA (middle) and L-Pro + D-FDTA (bottom) in UPLC-MS analysis.

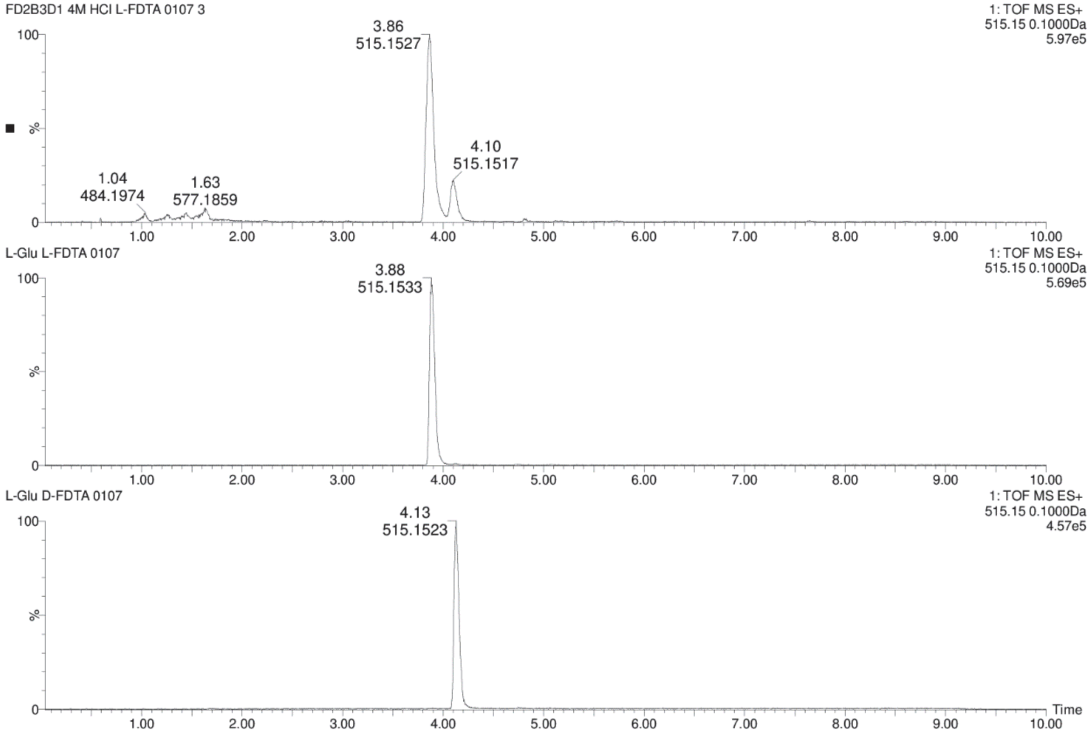

**Figure S34.** Chromatograms for comparison of **2** + L-FDTA (top) and L-Glu + L-FDTA (middle) and L-Glu + D-FDTA (bottom) in UPLC-MS analysis.

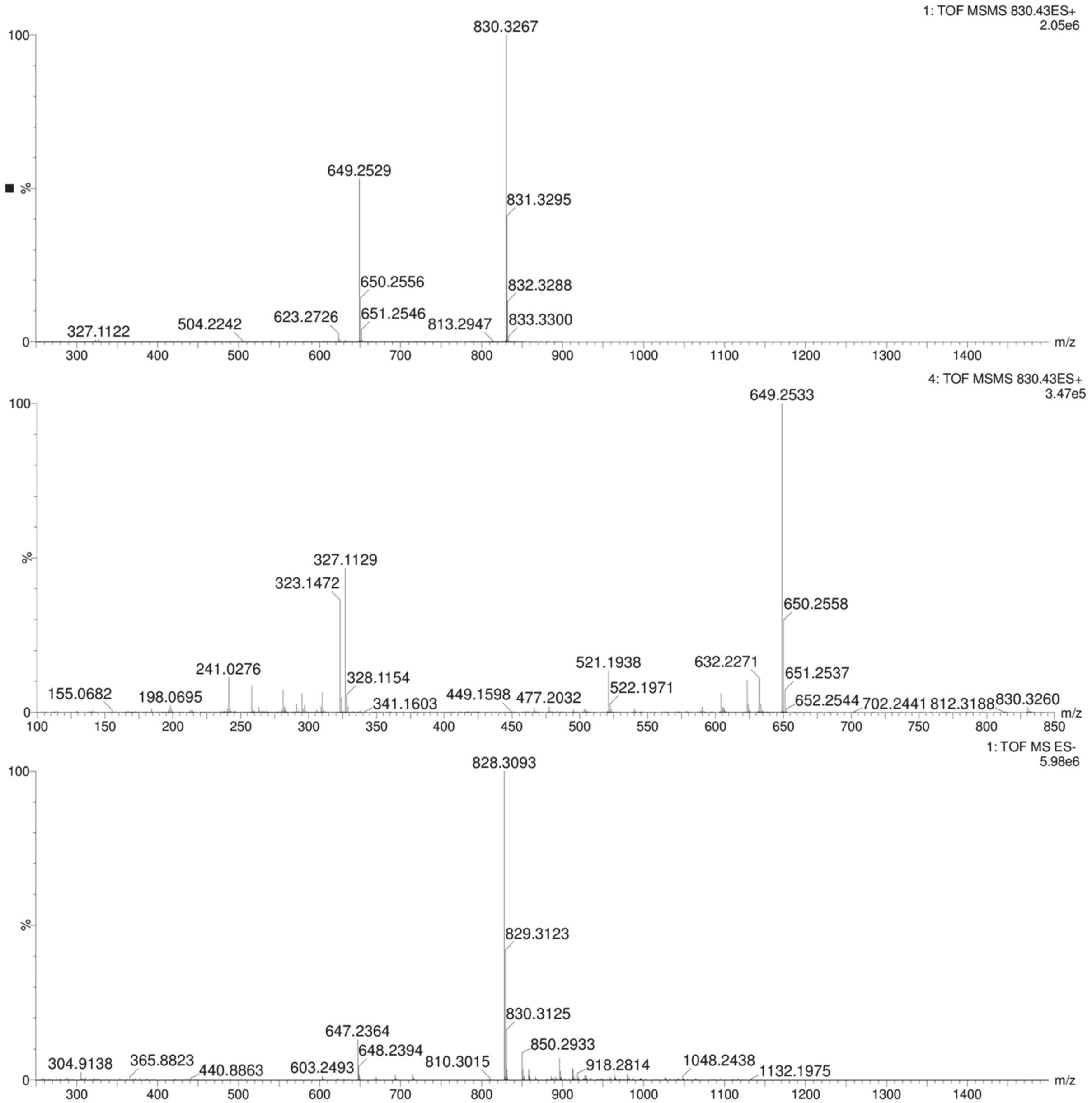

**Figure S35.** Mass spectrum of compound present in fraction FD2B3A1 (**3**) from *P. brasiliensis* Ab134. QToF-MS in positive mode (top), $MS^2$ in positive mode (middle) and MS in negative mode (bottom).

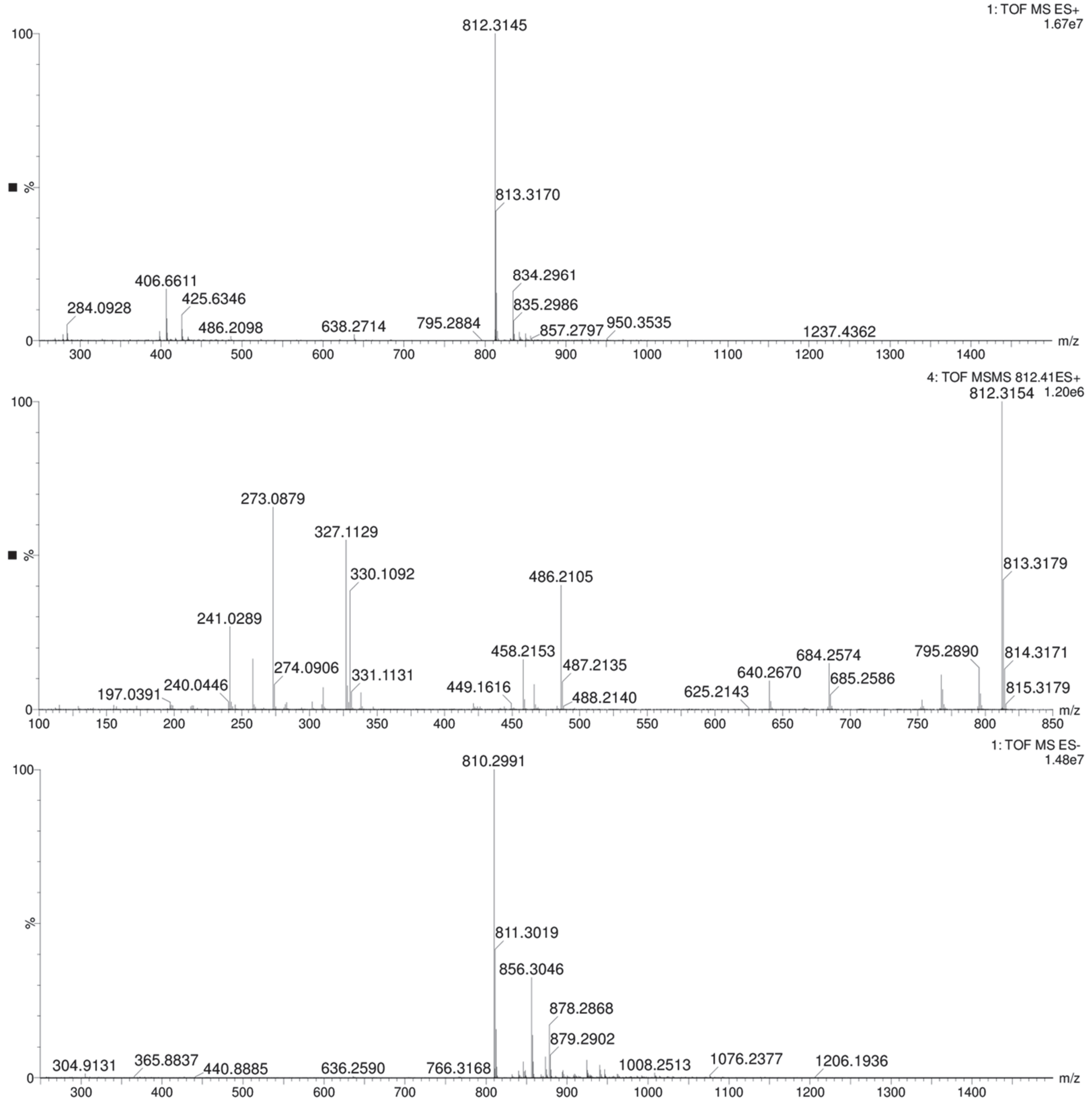

**Figure S36.** Mass spectrum of compound present in fraction FD2B3C1 (**4**) from *P. brasiliensis* Ab134. QToF-MS in positive mode (top), $MS^2$ in positive mode (middle) and MS in negative mode (bottom).

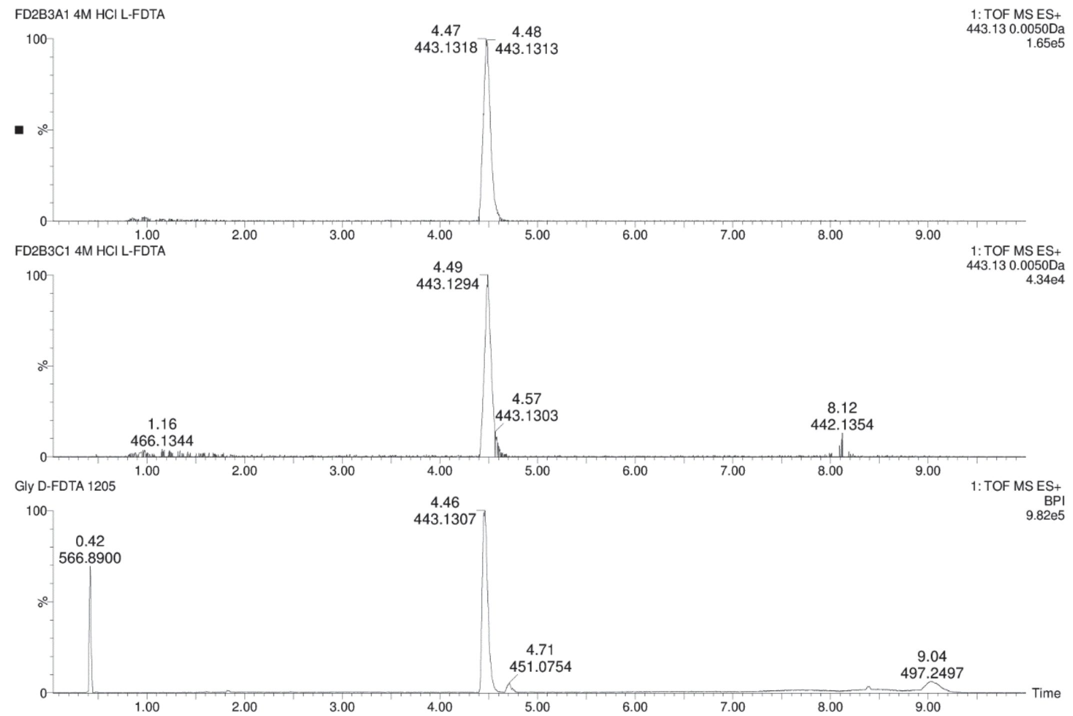

**Figure S37.** Chromatograms for comparison of **3** + L-FDTA (top) and **4** + L-FDTA (middle) and Gly + D-FDTA (bottom) in UPLC-MS analysis.

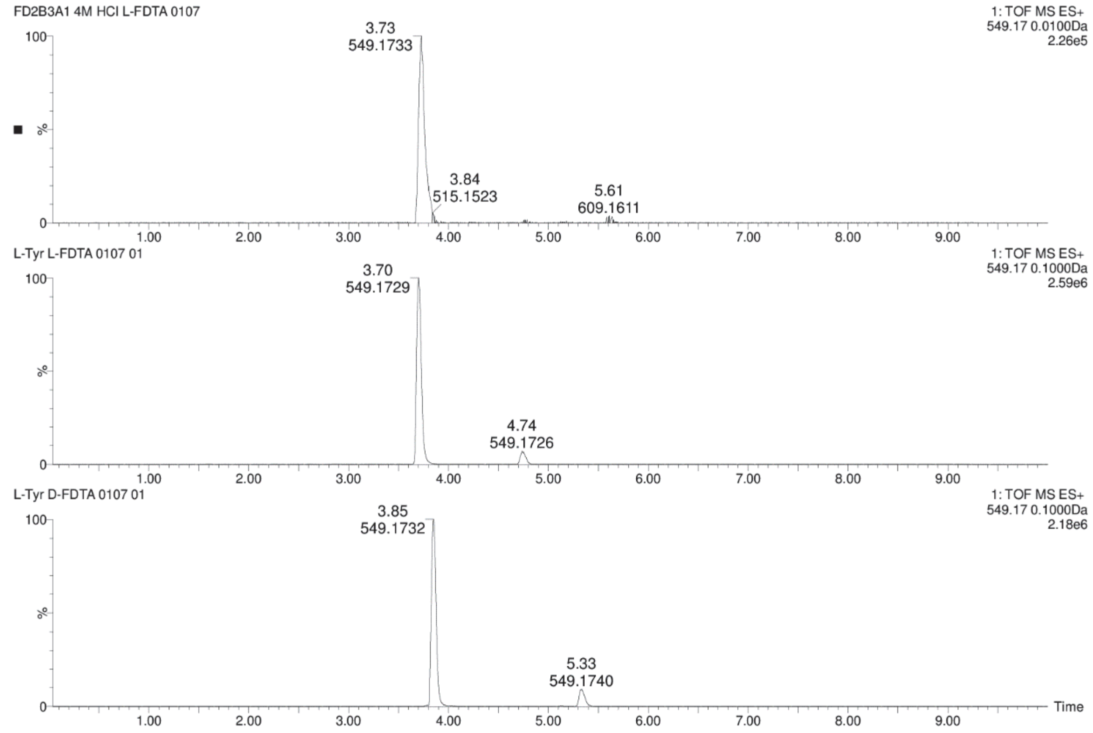

**Figure S38.** Chromatograms for comparison of **3** + L-FDTA (top) and L-Tyr + L-FDTA (middle) and L-Tyr + D-FDTA (bottom) in UPLC-MS analysis.

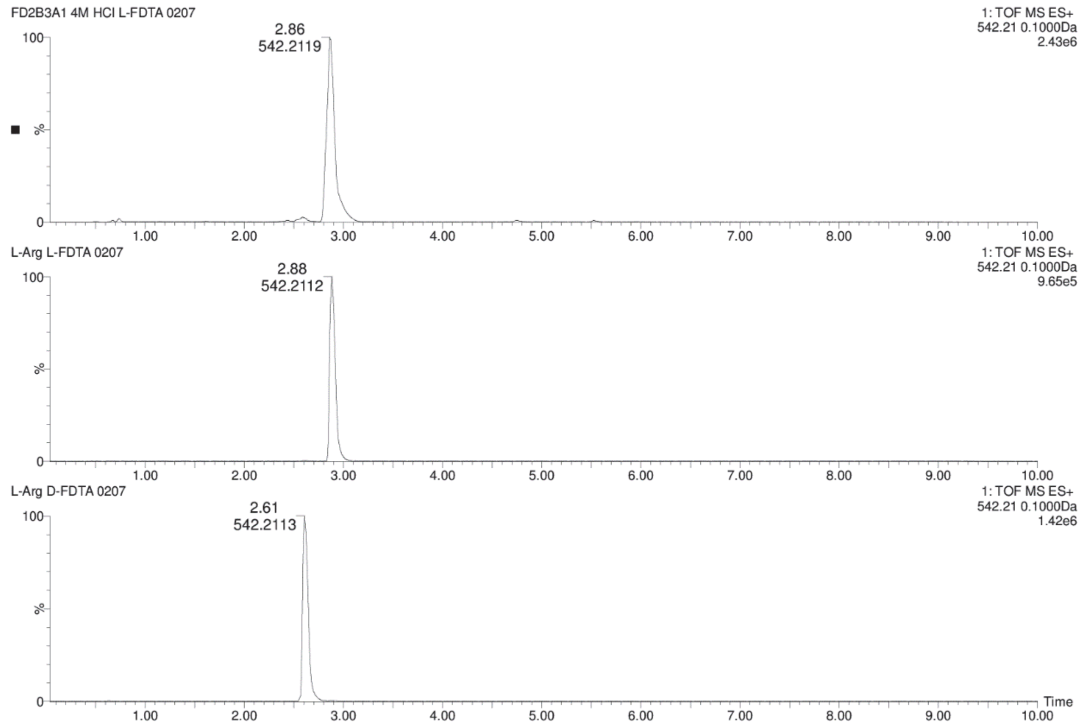

**Figure S39.** Chromatograms for comparison of **3** + L-FDTA (top) and L-Arg + L-FDTA (middle) and L-Arg + D-FDTA (bottom) in UPLC-MS analysis.

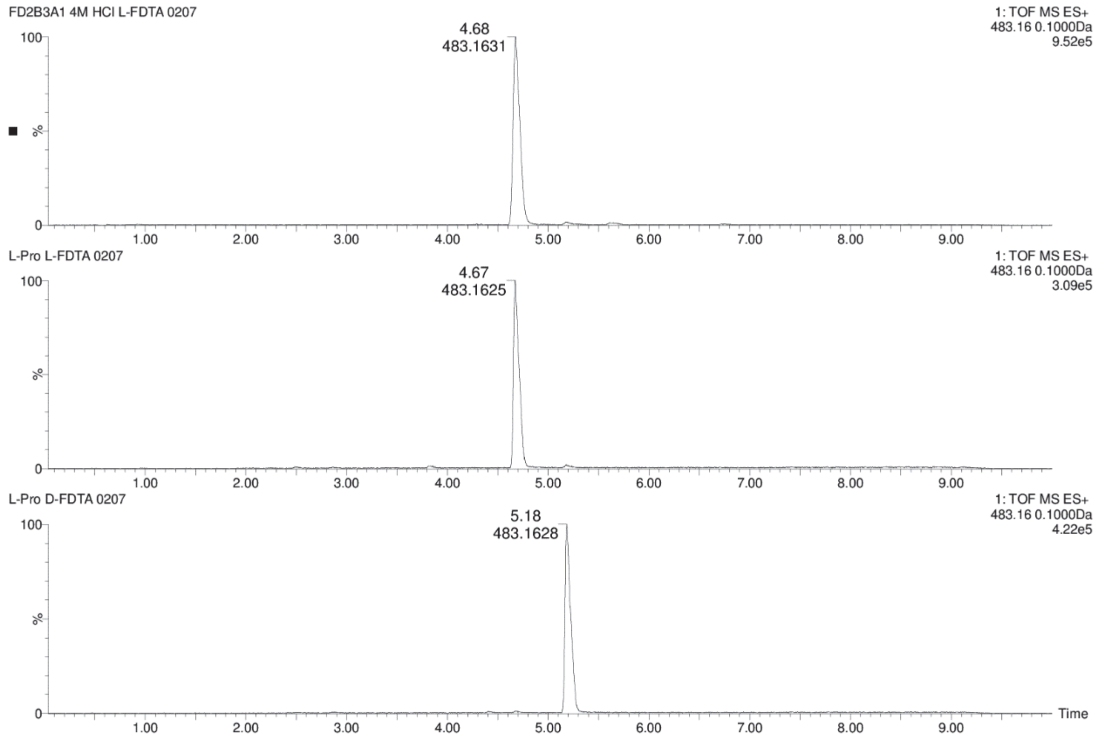

**Figure S40.** Chromatograms for comparison of **3** + L-FDTA (top) and L-Pro + L-FDTA (middle) and L-Pro + D-FDTA (bottom) in UPLC-MS analysis.

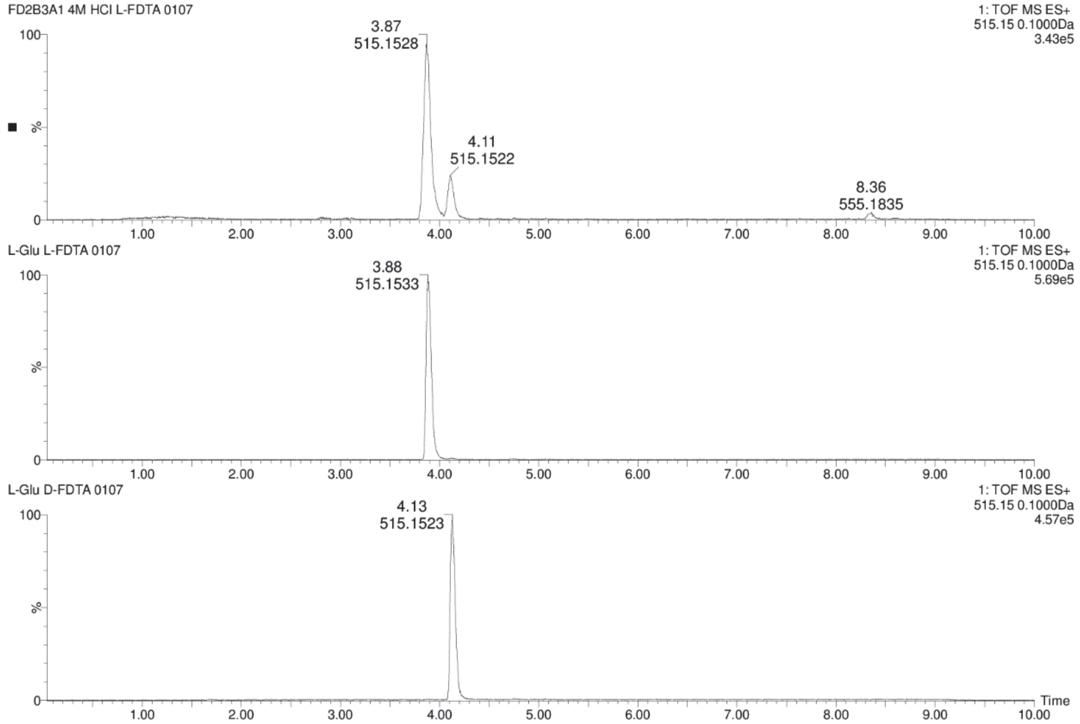

**Figure S41.** Chromatograms for comparison of **3** + L-FDTA (top) and L-Glu + L-FDTA (middle) and L-Glu + D-FDTA (bottom) in UPLC-MS analysis.

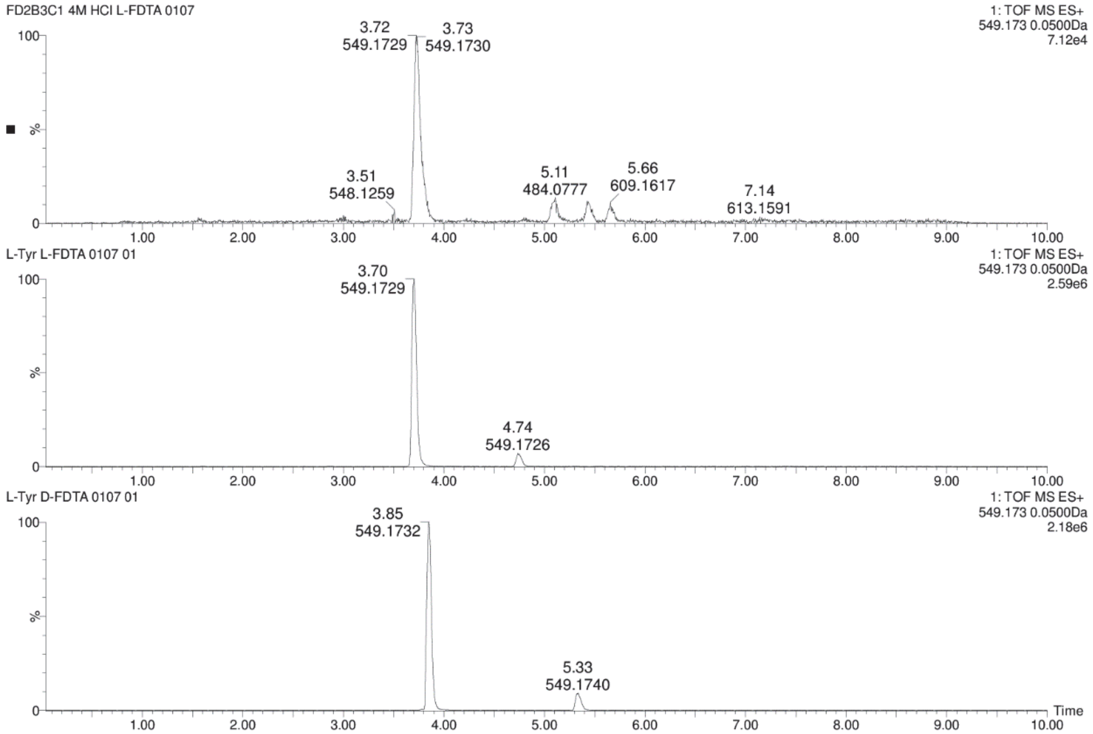

**Figure S42.** Chromatograms for comparison of **4** + L-FDTA (top) and L-Tyr + L-FDTA (middle) and L-Tyr + D-FDTA (bottom) in UPLC-MS analysis.

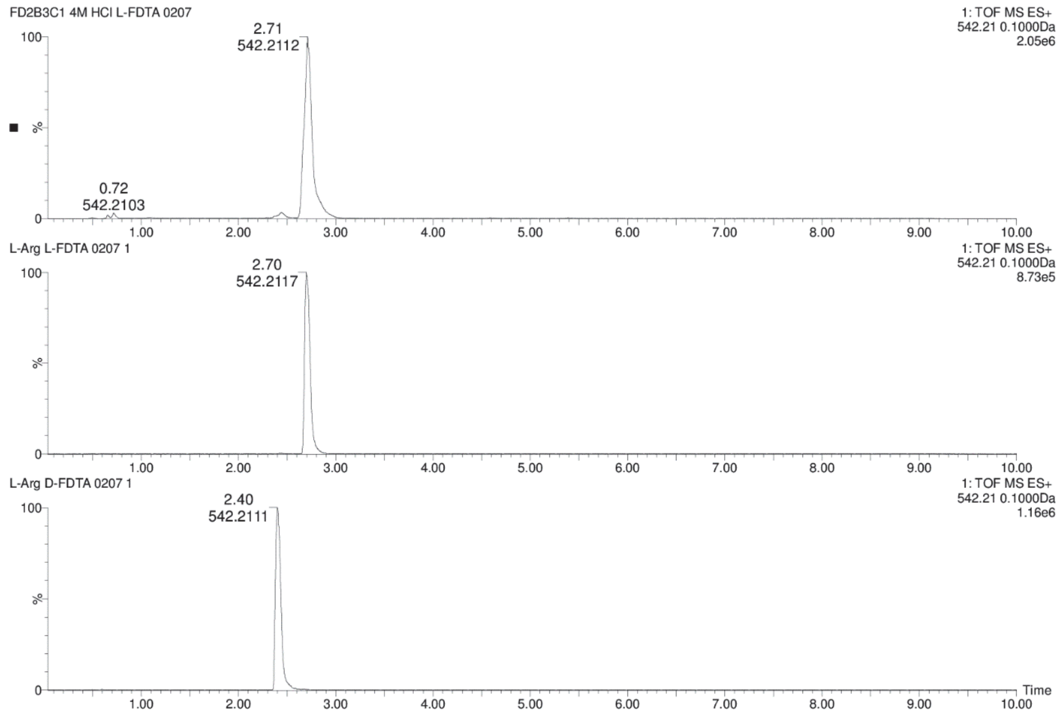

**Figure S43.** Chromatograms for comparison of **4** + L-FDTA (top) and L-Arg + L-FDTA (middle) and L-Arg + D-FDTA (bottom) in UPLC-MS analysis.

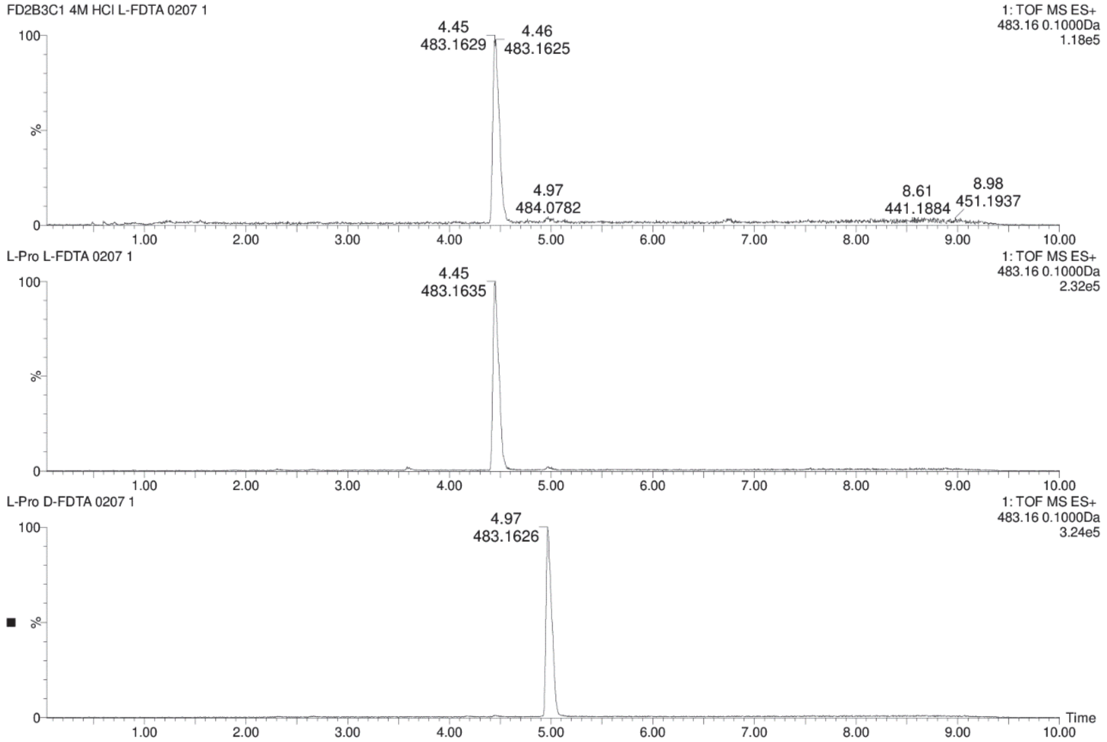

**Figure S44.** Chromatograms for comparison of **4** + L-FDTA (top) and L-Pro + L-FDTA (middle) and L-Pro + D-FDTA (bottom) in UPLC-MS analysis.

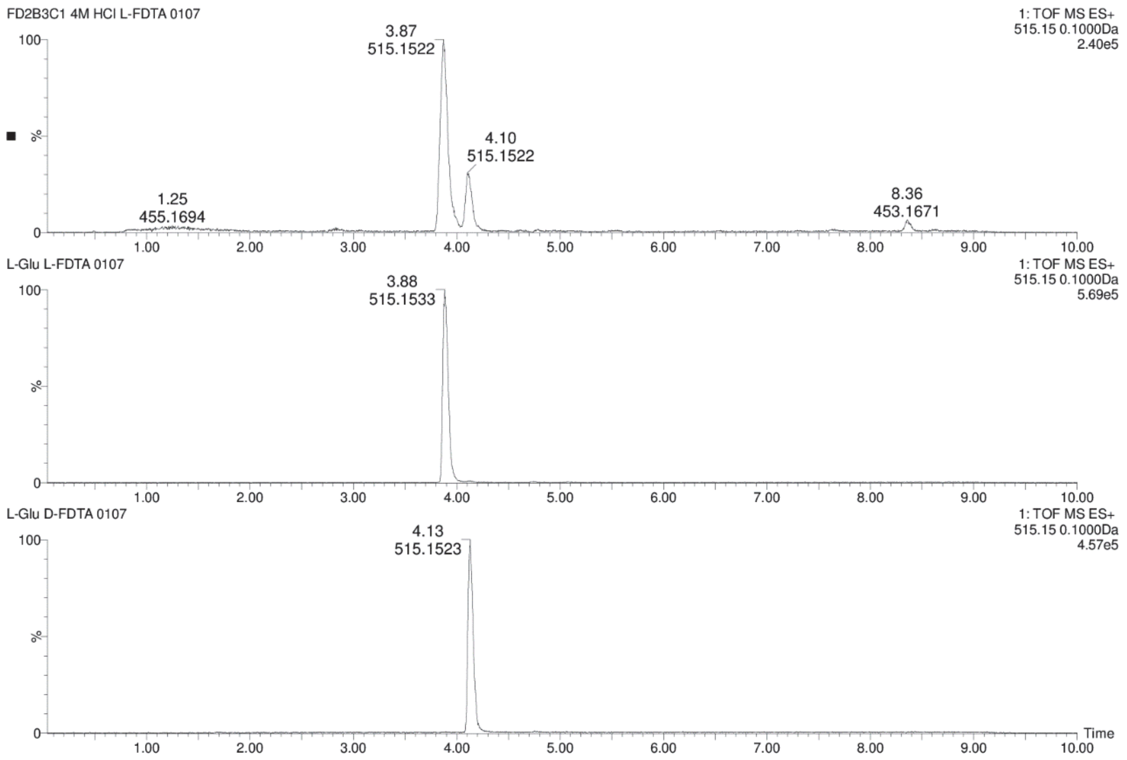

**Figure S45.** Chromatograms for comparison of **4** + L-FDTA (top) and L-Glu + L-FDTA (middle) and L-Glu + D-FDTA (bottom) in UPLC-MS analysis.

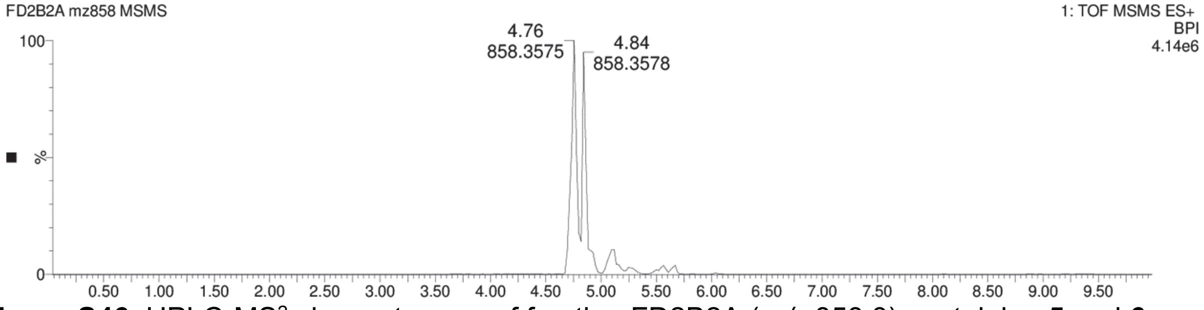

**Figure S46.** UPLC-MS$^2$ chromatogram of fraction FD2B2A ($m/z$ 858.3) containing **5** and **6**.

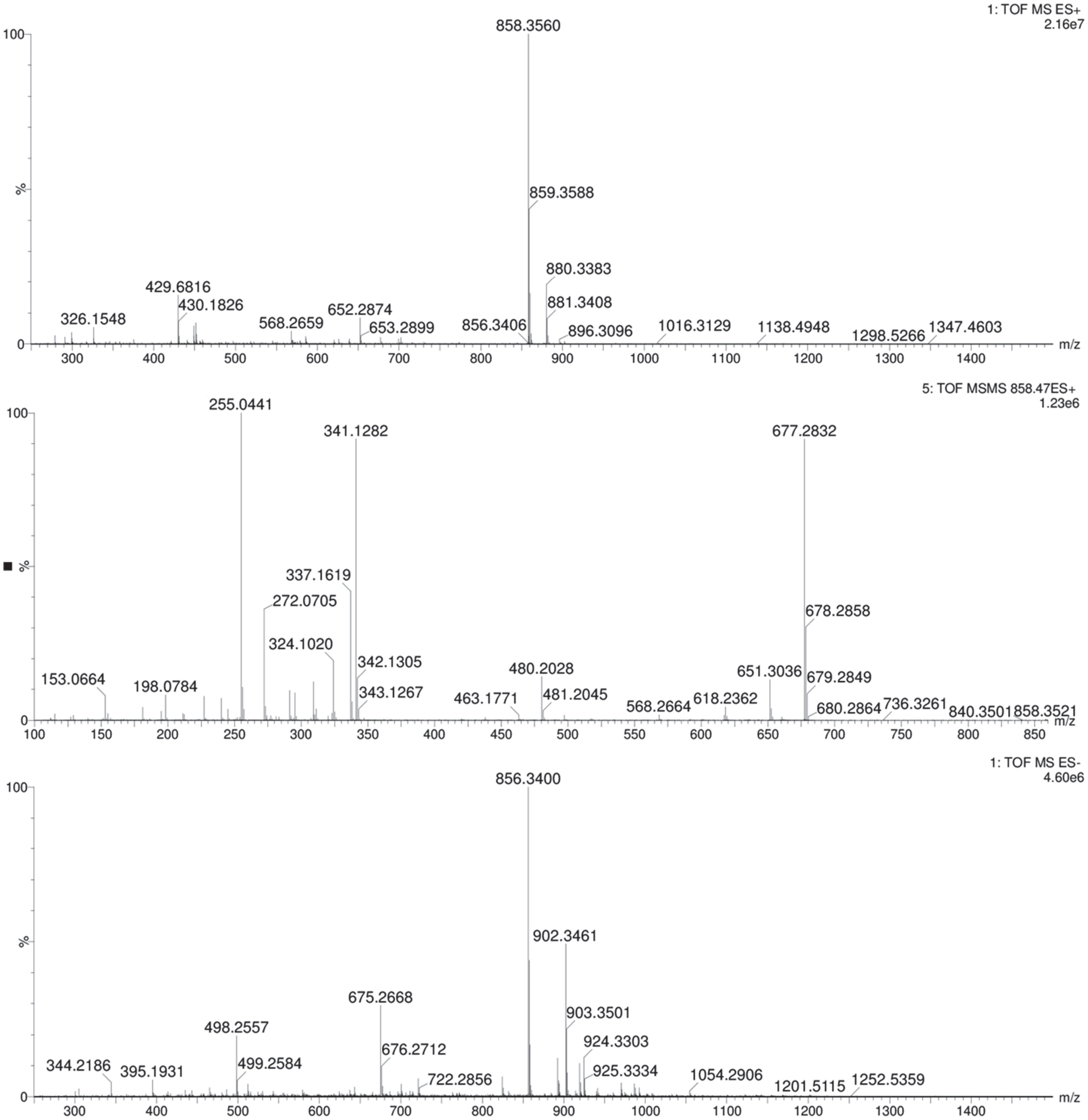

**Figure S47.** Mass spectrum of compound present in fraction FD2B2A (**5**) from *P. brasiliensis* Ab134. QToF-MS in positive mode (top), MS$^2$ in positive mode (middle) and MS in negative mode (bottom).

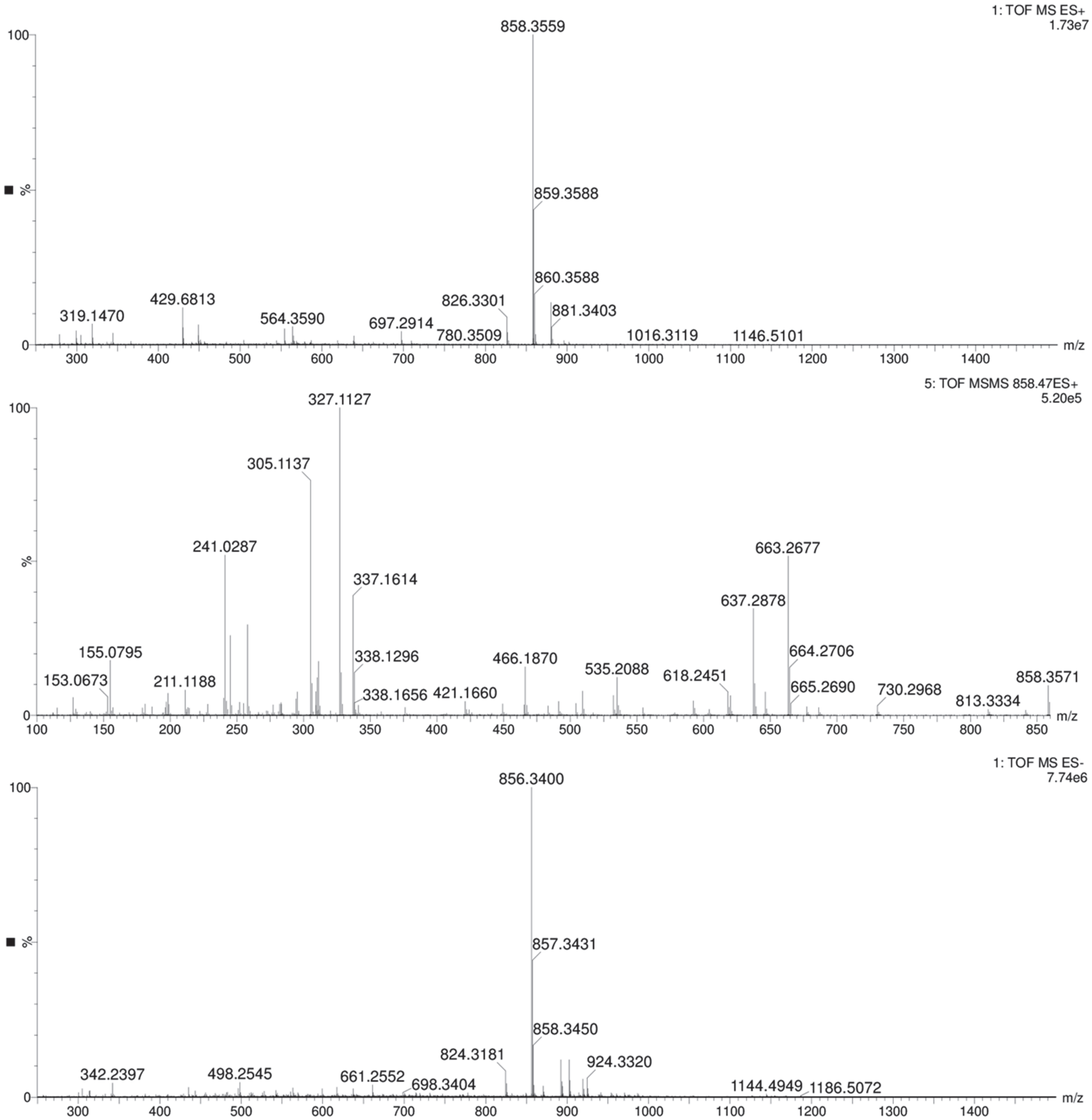

**Figure S48.** Mass spectrum of compound present in fraction FD2B2A (**6**) from *P. brasiliensis* Ab134. QToF-MS in positive mode (top), $MS^2$ in positive mode (middle) and MS in negative mode (bottom).

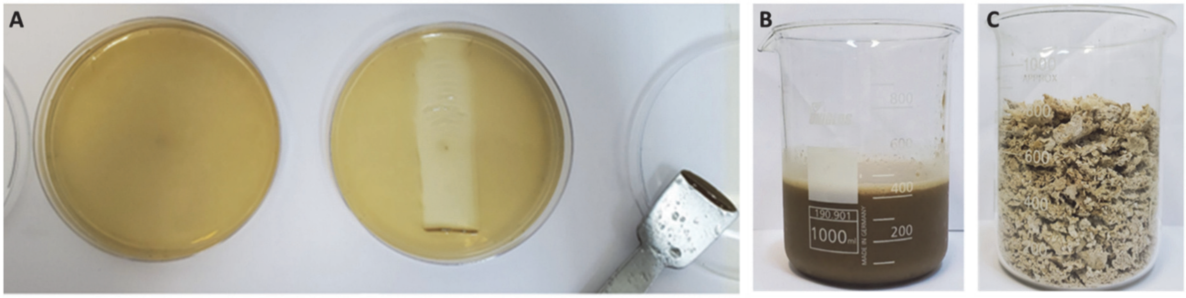

**Figure S49.** Pictures of swarming plates extraction. A) Picture of a swarming assay plate before (left) and during cell recovery (right) to generate extract C. The spatula with cell matt is shown on the left. B) Picture of the volume of cells obtained from 500 swarming plates after freezing and thawing, and before methanol extraction. C) Representative picture of lyophilized solid culture (cells and media) used to generate extract FD.

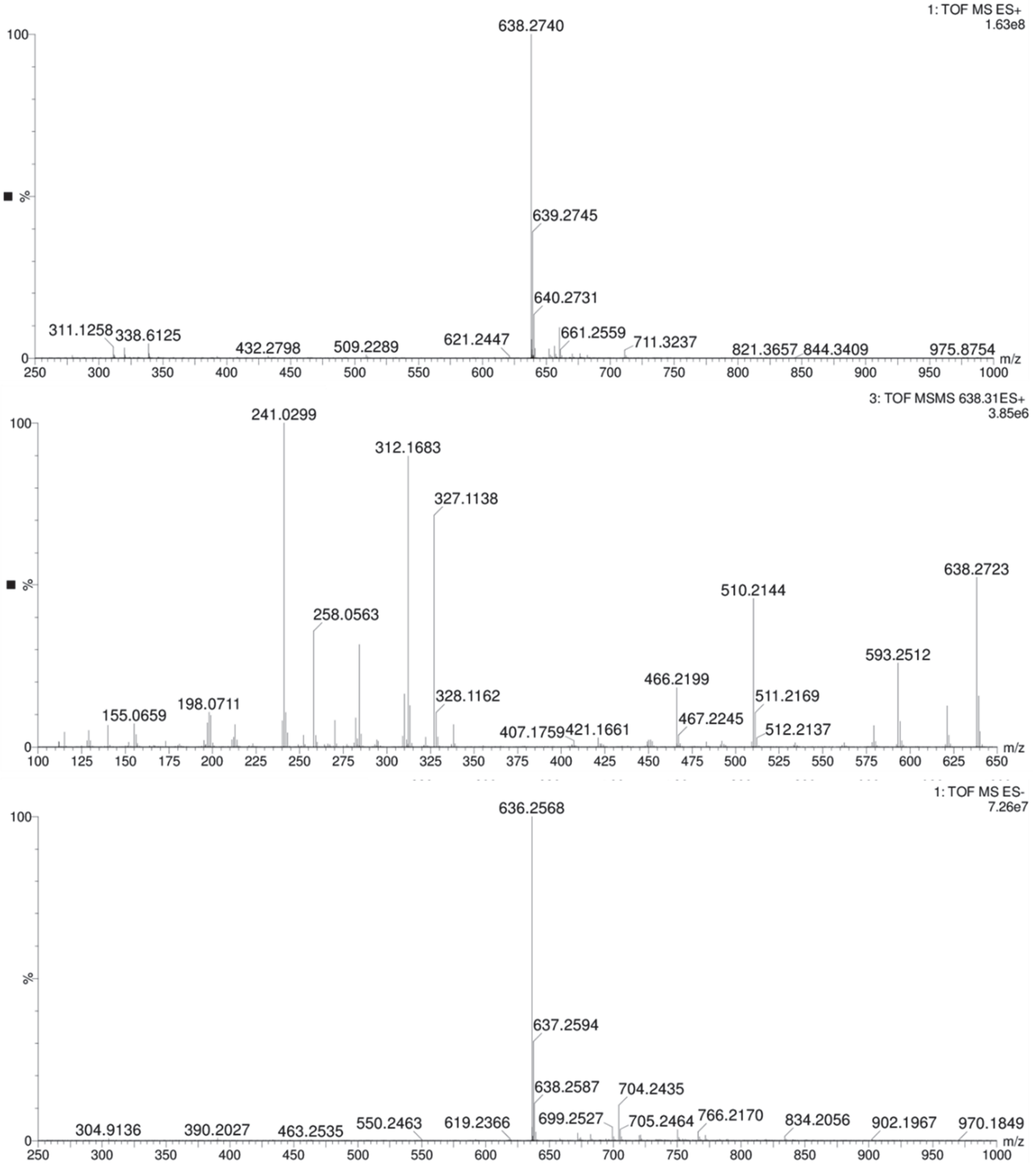

**Figure S50.** Mass spectrum of compound present in fraction FD2B3A, degradation product of pseudovibriamide A1 (**1**). QToF-MS in positive mode (top), MS$^2$ in positive mode, collision energy of 35V (middle) and MS in negative mode (bottom). See Fig. S51 for predicted degradation product structure and Table S13 for expected fragment ions.

A

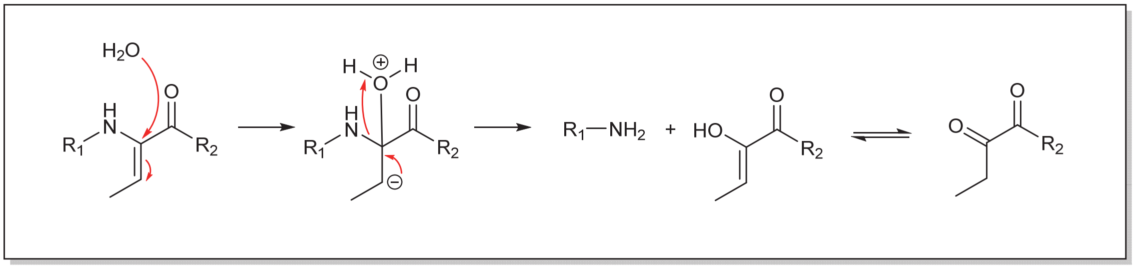

B

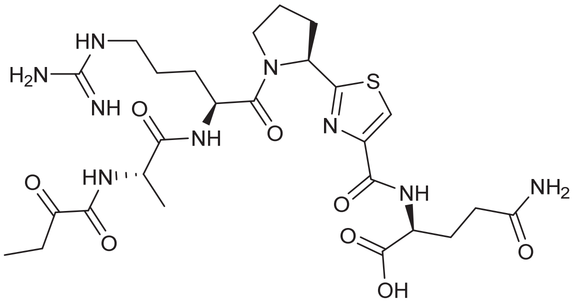

**Figure S51**. Pseudovibriamide A1 degradation. **A**) Hypothetical mechanism of pseudovibriamides degradation. **B**) Pseudovibriamide A1 degradation product with $[M + H]^+$ at *m/z* 638.

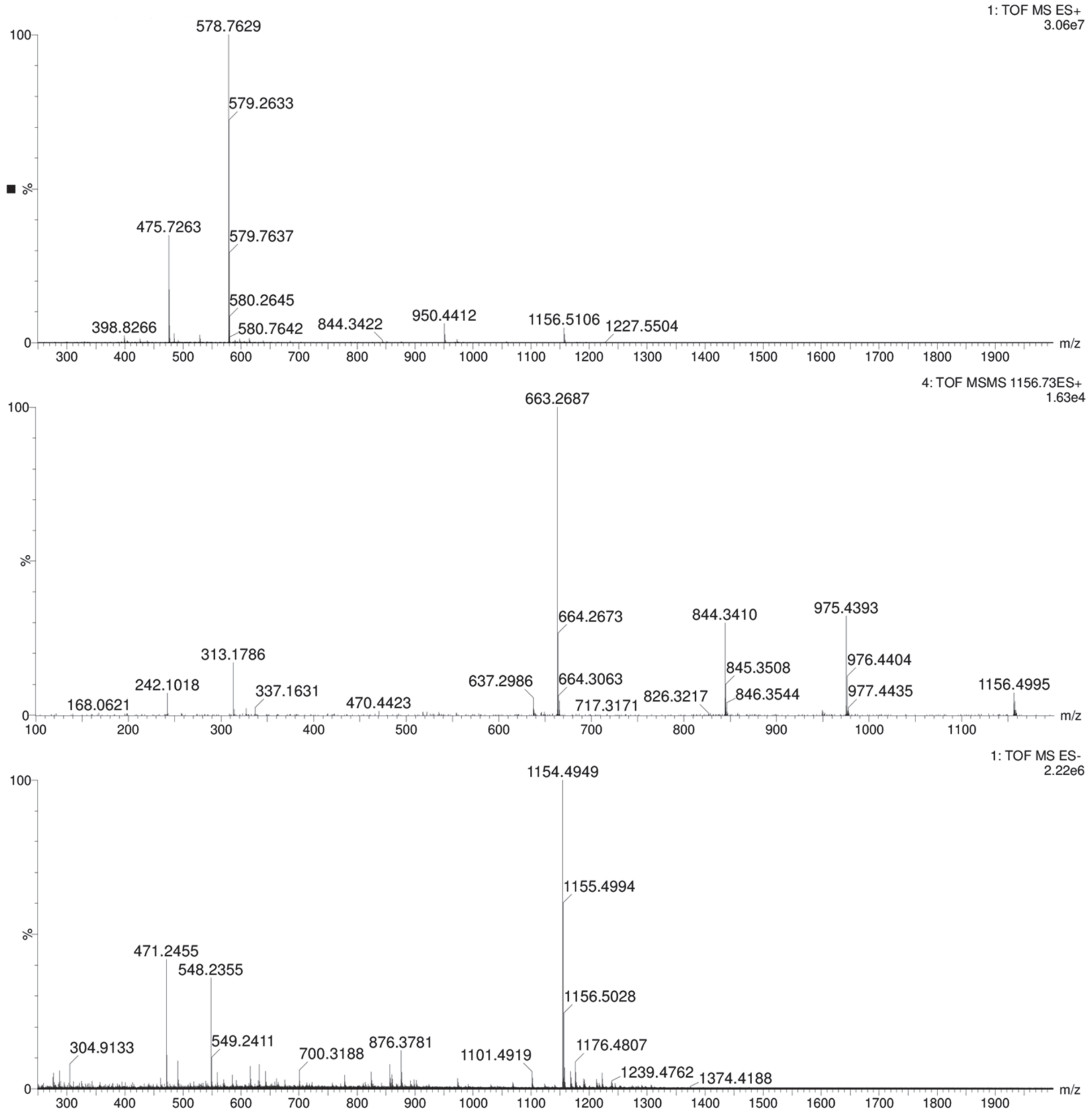

**Figure S52.** Mass spectrum of compound present in fraction C23C1B (**7**) from *P. brasiliensis* Ab134. QToF-MS in positive mode (top), $MS^2$ in positive mode (middle) and MS in negative mode (bottom). See also Table S15.

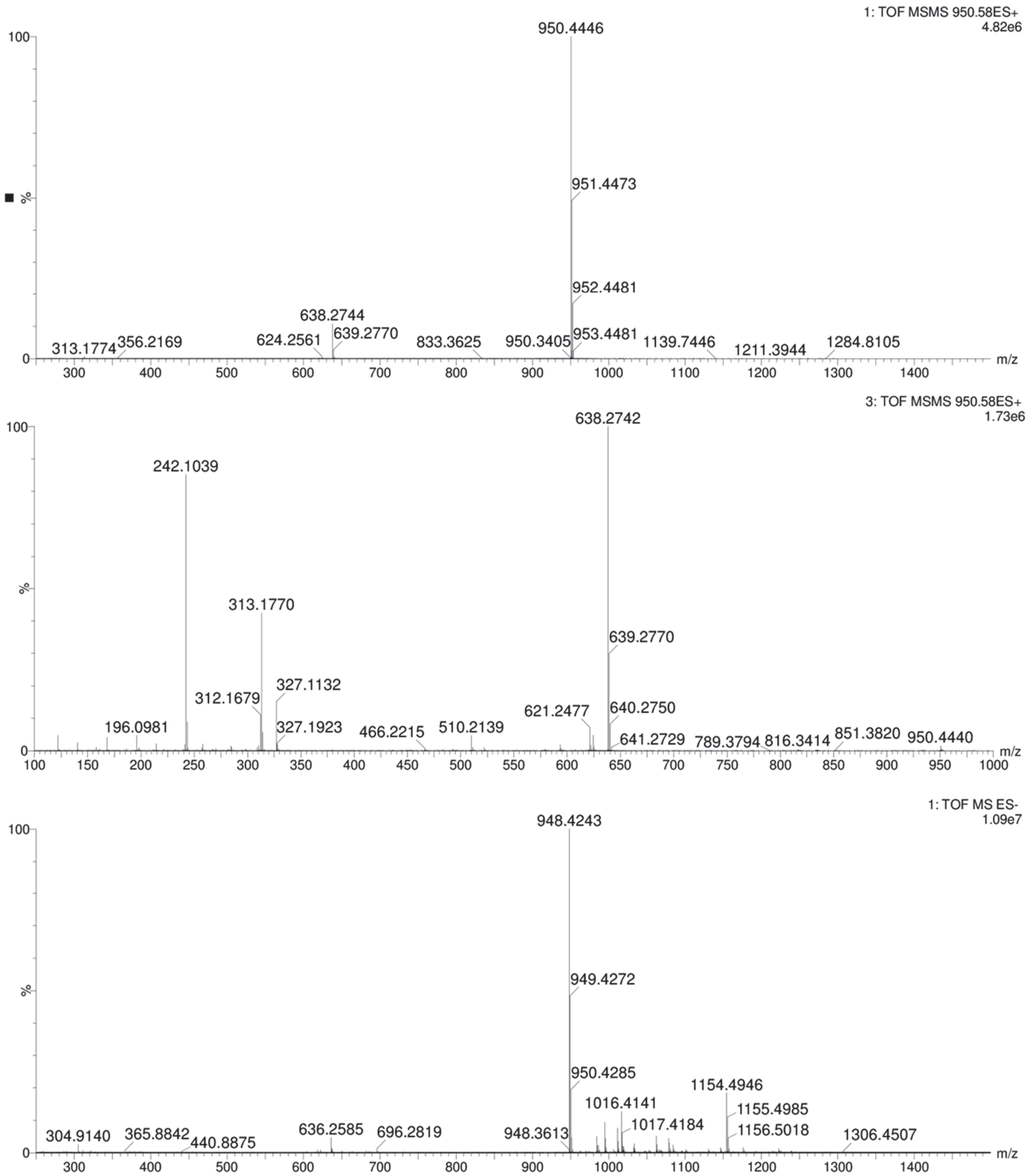

**Figure S53.** Mass spectrum of compound present in fraction C23C1B, degradation product of pseudovibriamide B1 (**7**). QToF-MS in positive mode (top), MS$^2$ in positive mode, collision energy of 45V (middle) and MS in negative mode (bottom).

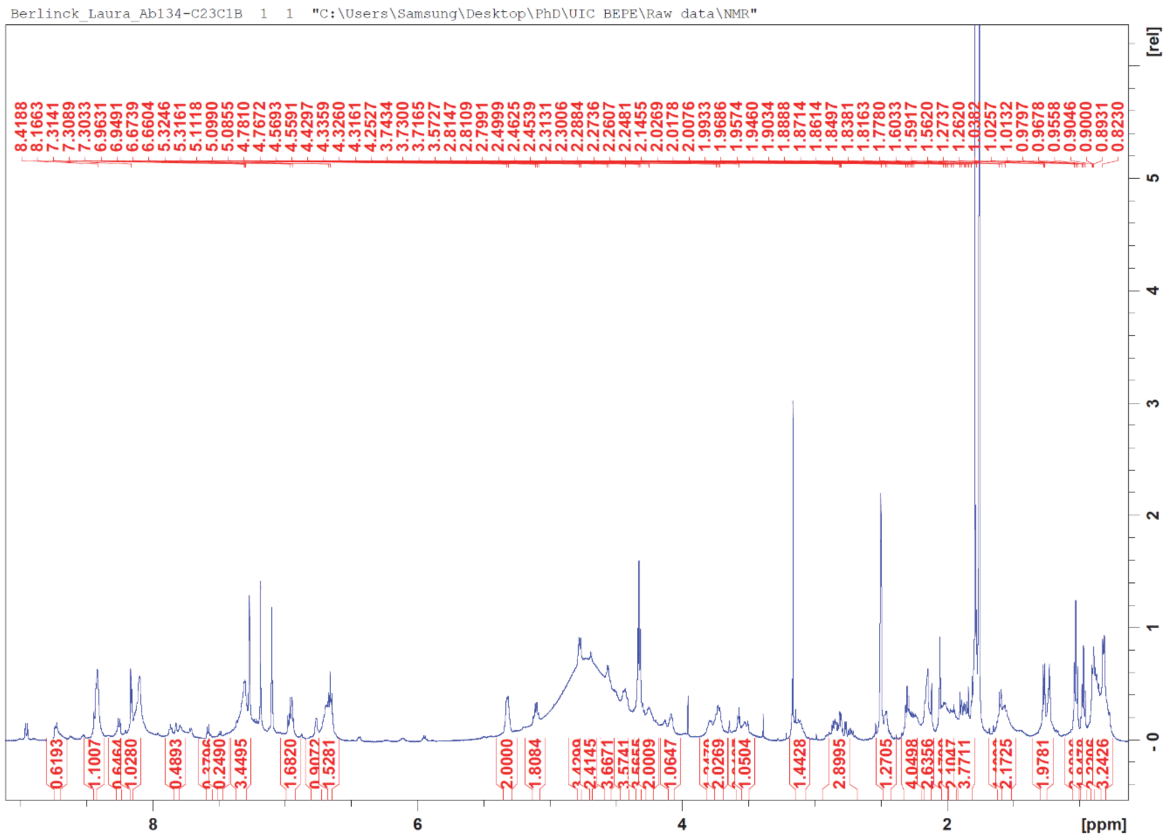

**Figure S54.** $^1$H NMR spectrum of C23C1B (**7**) at 600 MHz, DMSO-$d_6$ + TFA vapor.

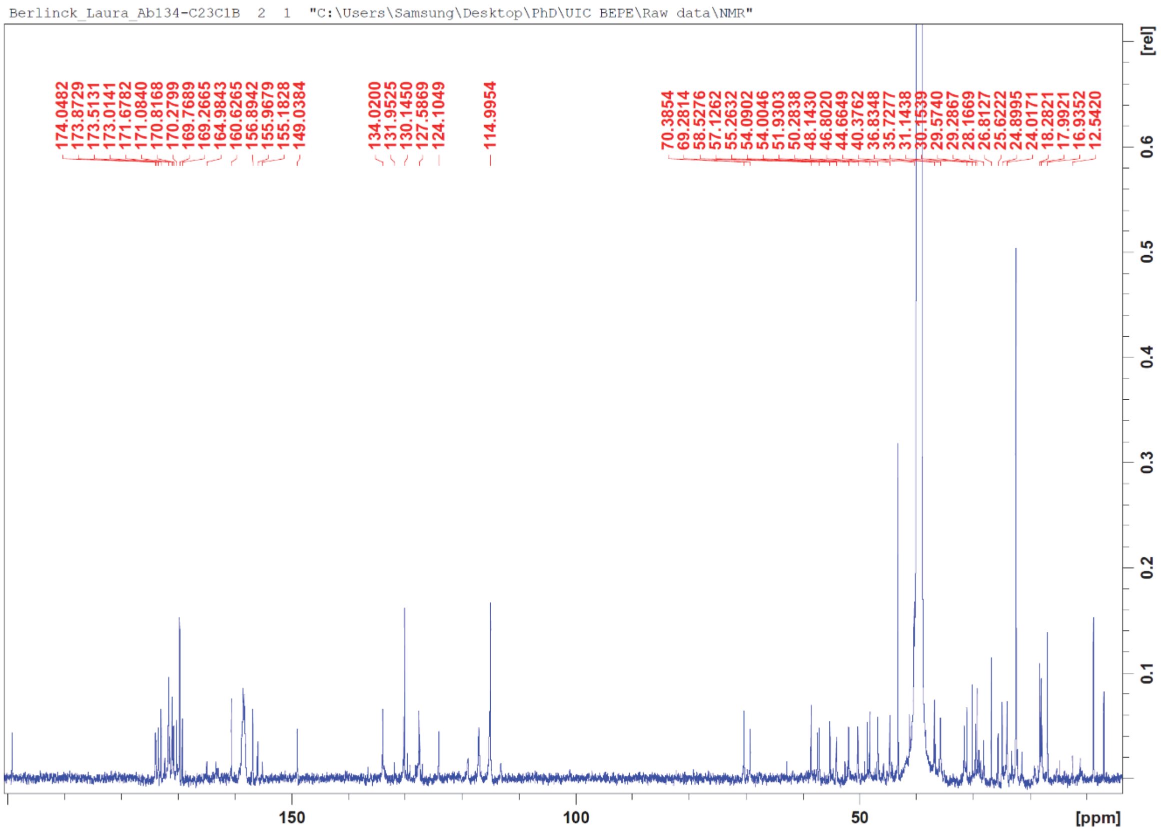

**Figure S55.** $^{13}C$ NMR spectrum of C23C1B (**7**) at 150 MHz, DMSO-$d_6$ + TFA vapor.

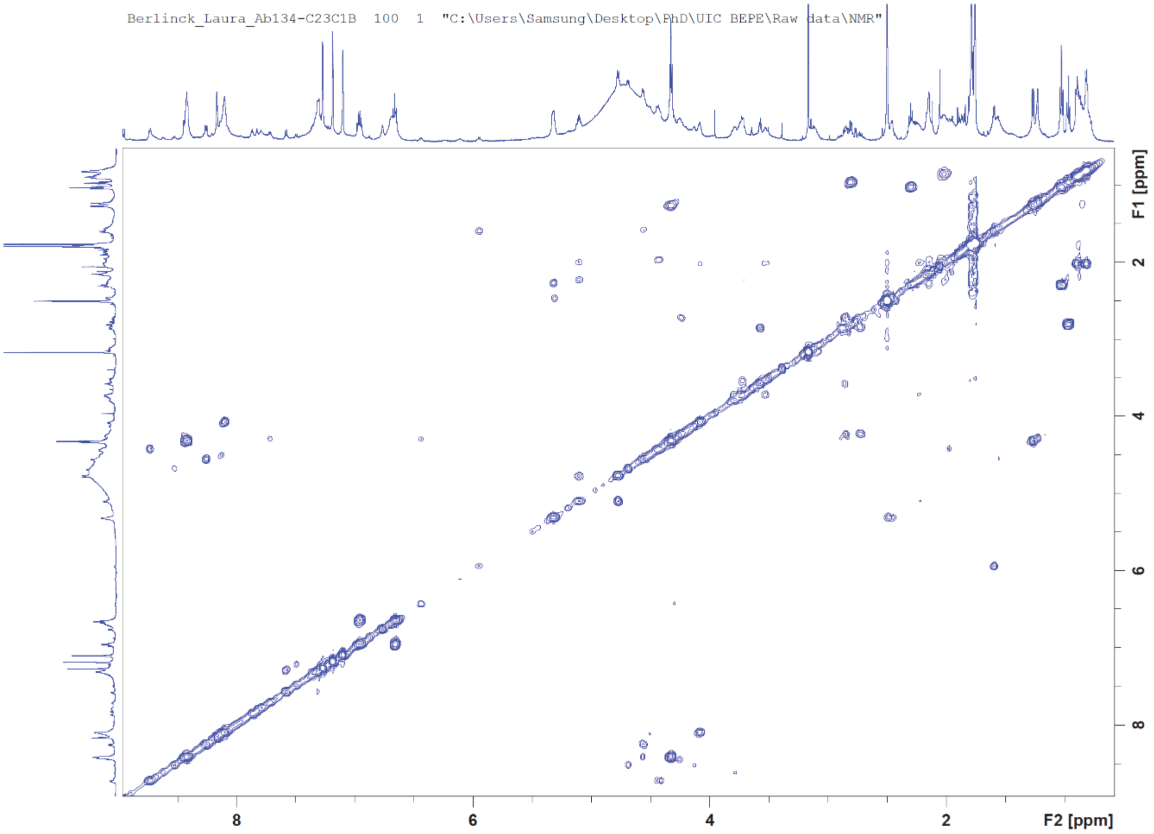

**Figure S56.** COSY NMR spectrum of C23C1B (**7**) at 600 MHz, DMSO-$d_6$ + TFA vapor.

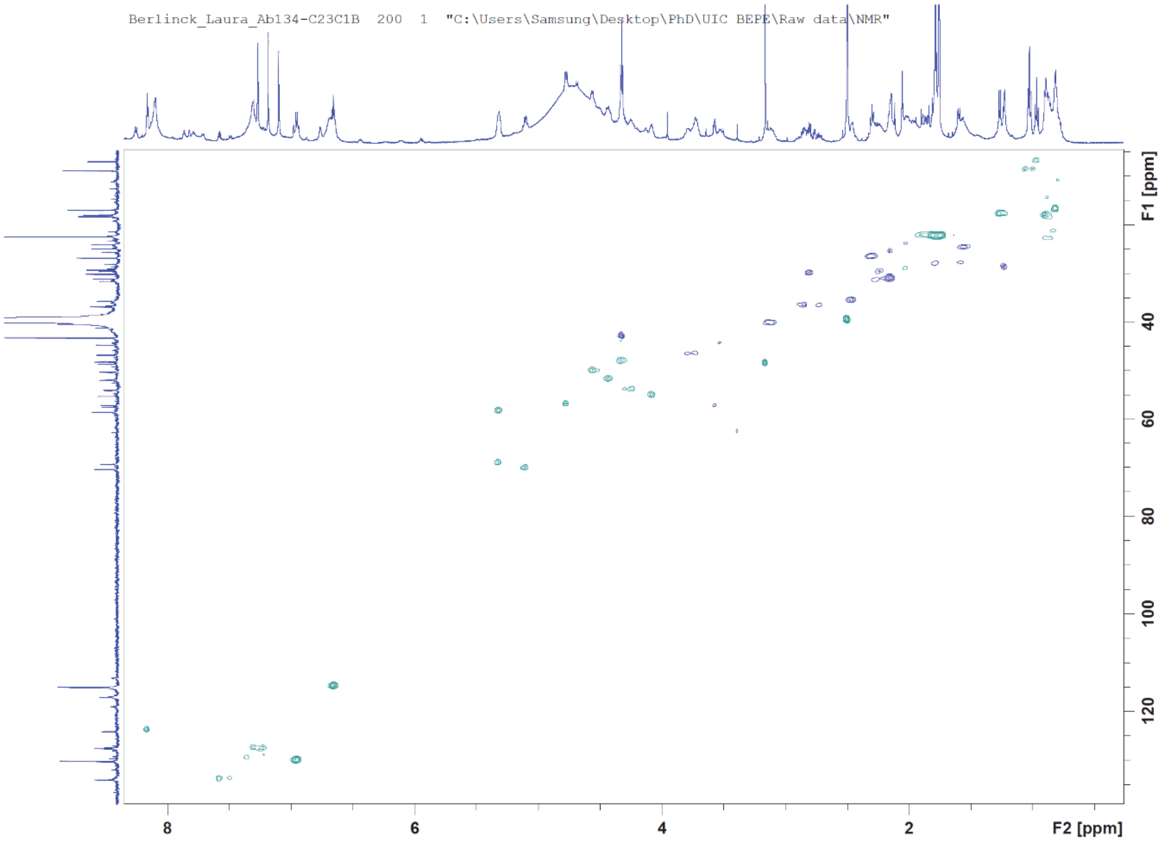

**Figure S57.** HSQC NMR spectrum of C23C1B (**7**) at 150 MHz:600 MHz, DMSO-$d_6$ + TFA vapor.

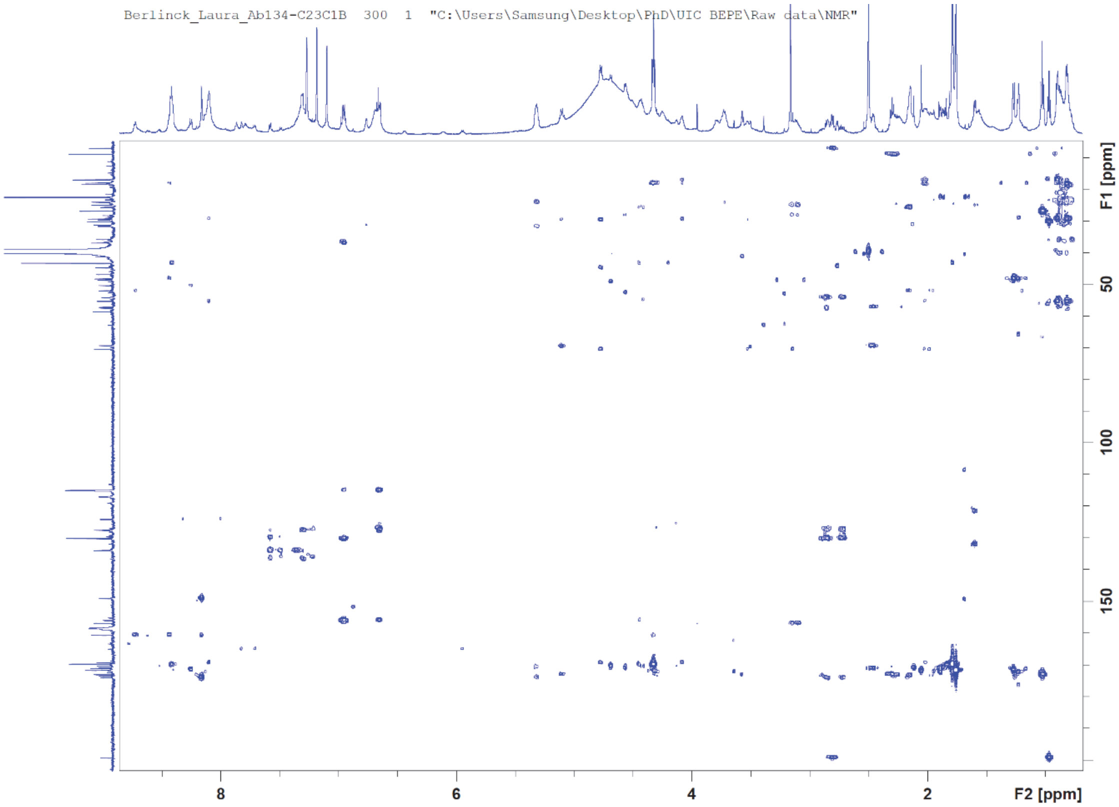

**Figure S58**. HMBC NMR spectrum of C23C1B (**7**) at 150 MHz:600 MHz, DMSO-$d_6$ + TFA vapor.

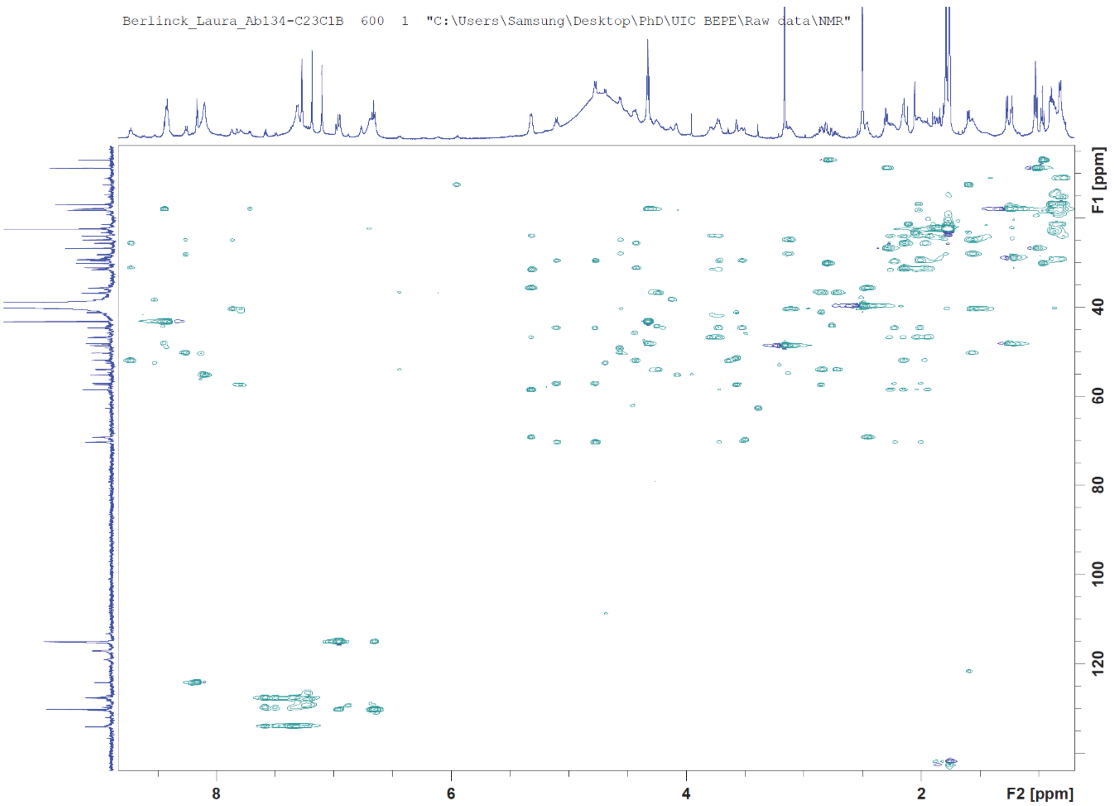

**Figure S59.** HSQC-TOCSY NMR spectrum of C23C1B (**7**) at 150 MHz:600 MHz, DMSO-$d_6$ + TFA vapor.

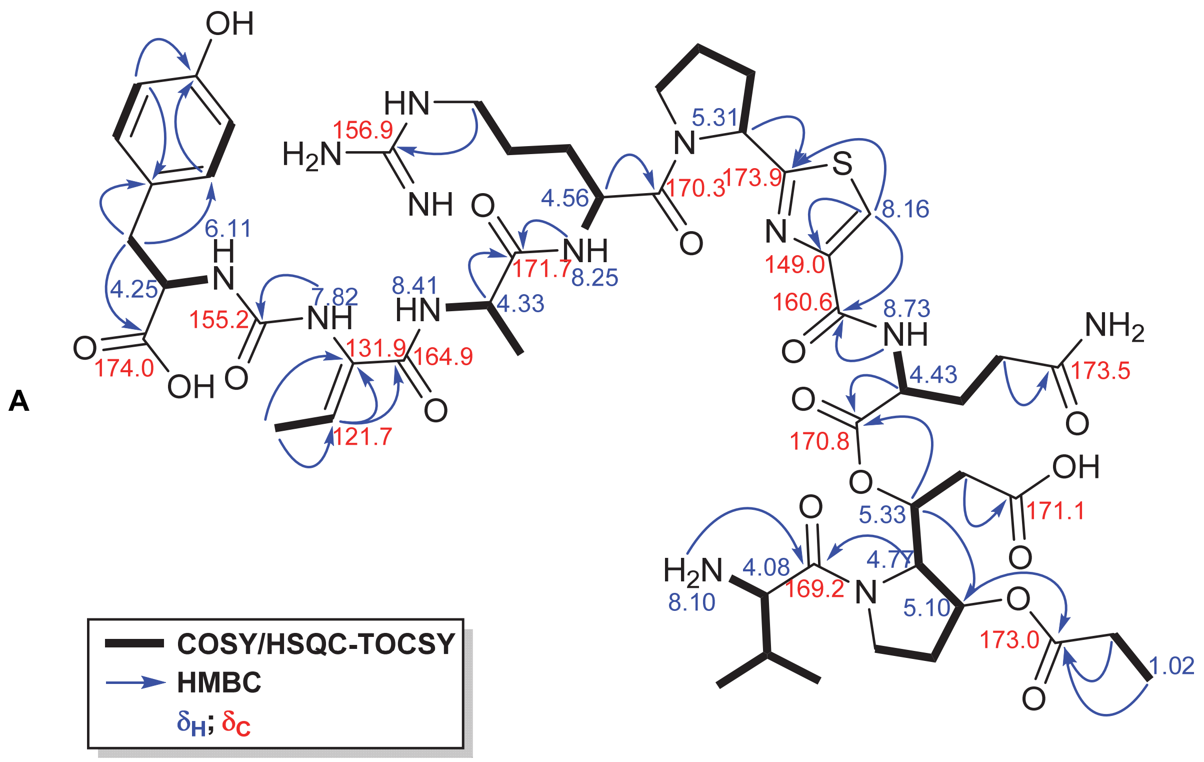

**Figure S60.** Key $^1$H and $^{13}$C NMR chemical shifts and HMBC, COSY and NOE/ROE correlations observed for compound **7**.

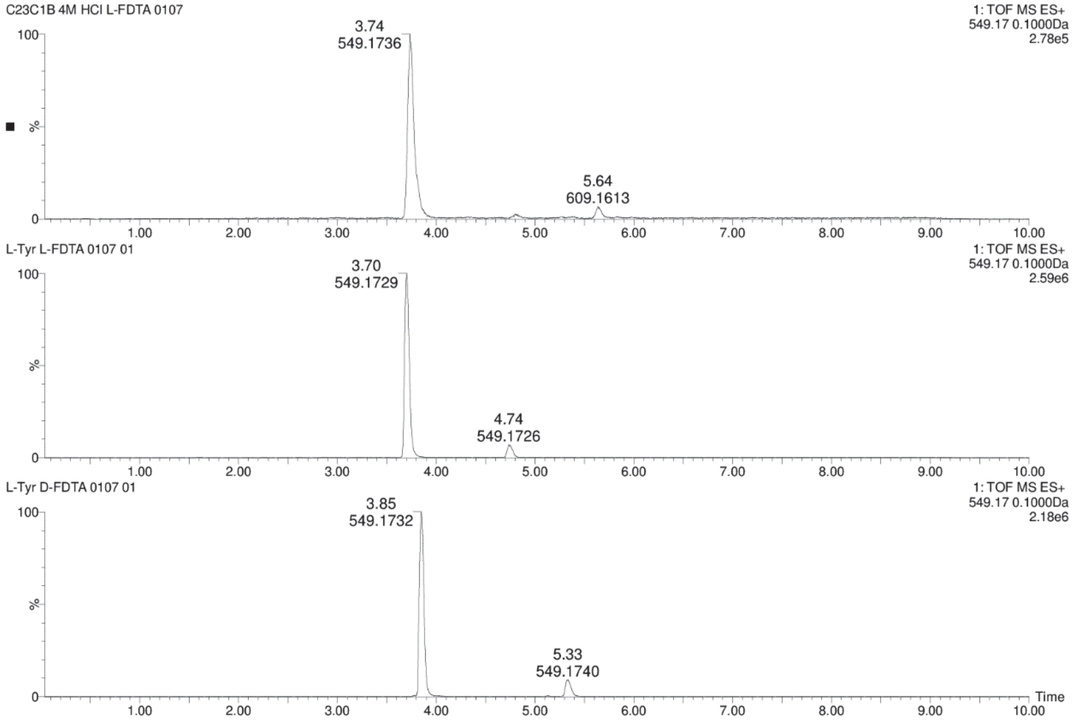

**Figure S61.** Chromatograms for comparison of **7** + L-FDTA (top) and L-Tyr + L-FDTA (middle) and L-Tyr + D-FDTA (bottom) in UPLC-MS analysis.

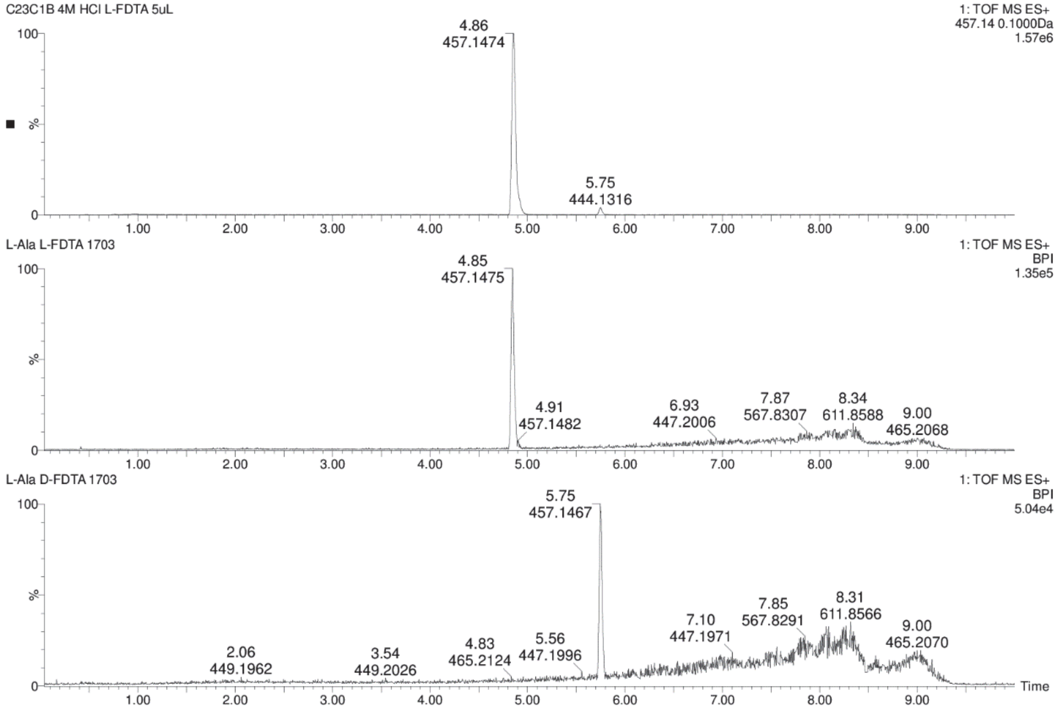

**Figure S62.** Chromatograms for comparison of **7** + L-FDTA (top) and L-Ala + L-FDTA (middle) and L-Ala + D-FDTA (bottom) in UPLC-MS analysis.

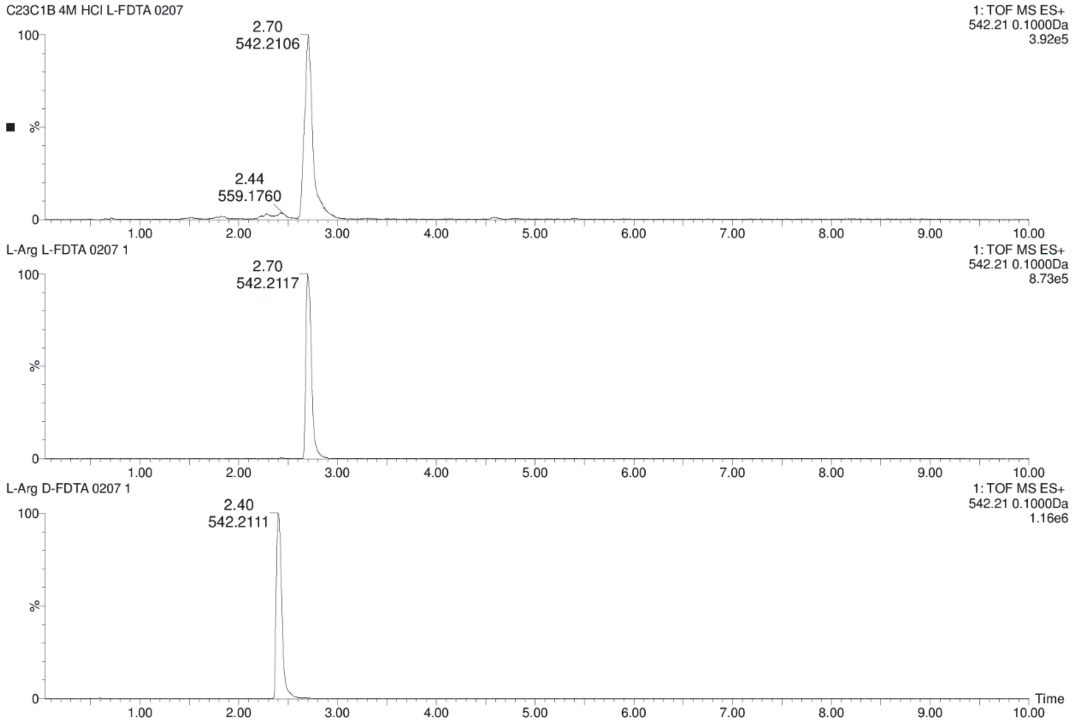

**Figure S63.** Chromatograms for comparison of **7** + L-FDTA (top) and L-Arg + L-FDTA (middle) and L-Arg + D-FDTA (bottom) in UPLC-MS analysis.

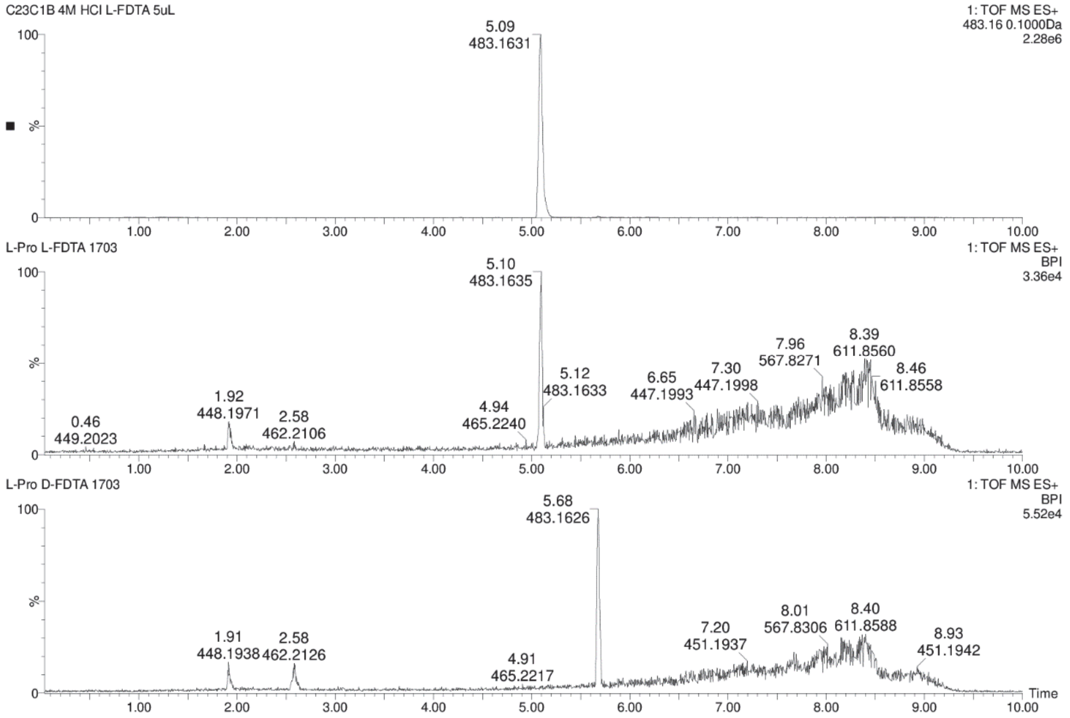

**Figure S64.** Chromatograms for comparison of **7** + L-FDTA (top) and L-Pro + L-FDTA (middle) and L-Pro + D-FDTA (bottom) in UPLC-MS analysis.

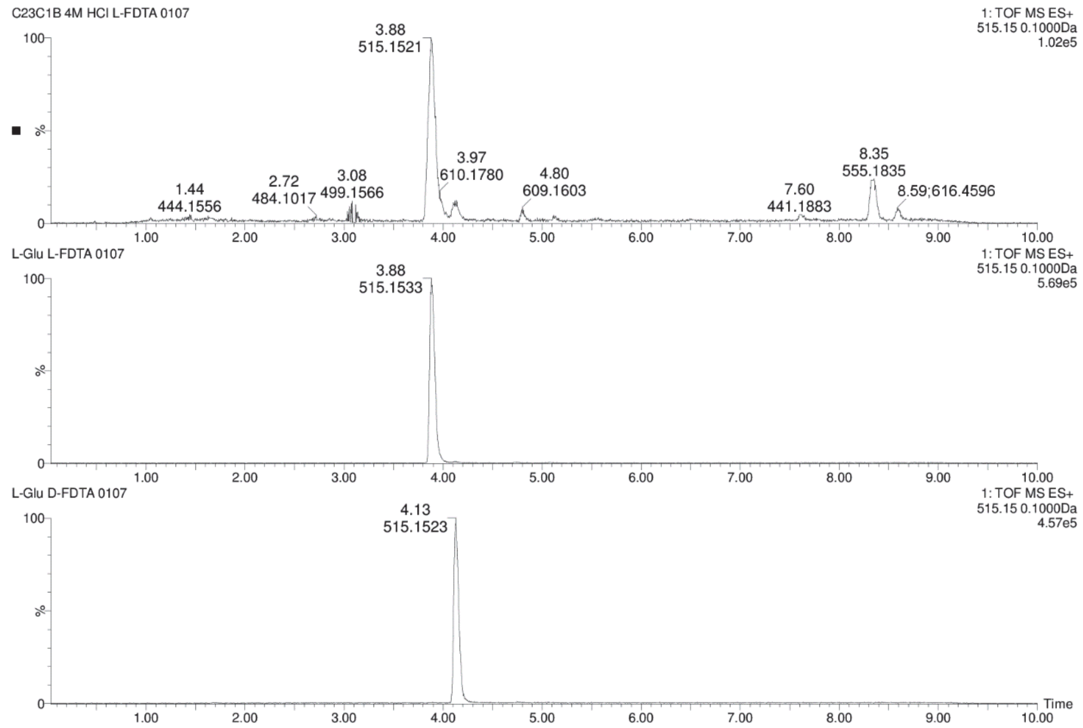

**Figure S65.** Chromatograms for comparison of **7** + L-FDTA (top) and L-Glu + L-FDTA (middle) and L-Glu + D-FDTA (bottom) in UPLC-MS analysis.

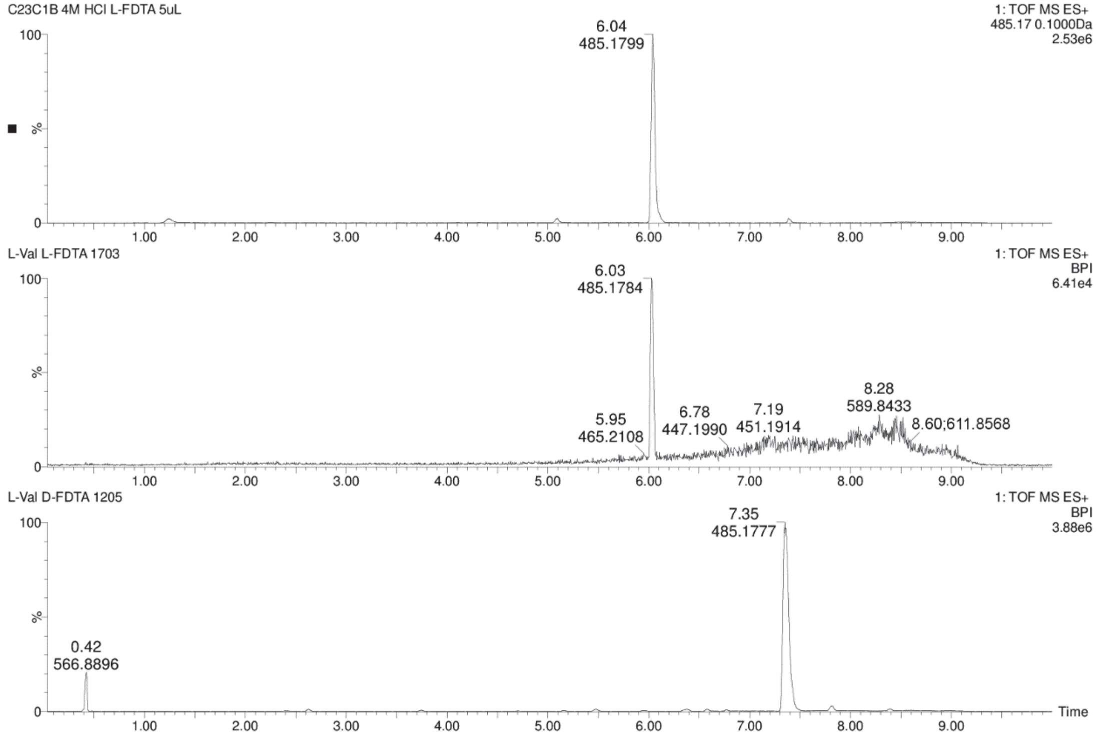

**Figure S66.** Chromatograms for comparison of **7** + L-FDTA (top) and L-Val + L-FDTA (middle) and L-Val + D-FDTA (bottom) in UPLC-MS analysis.

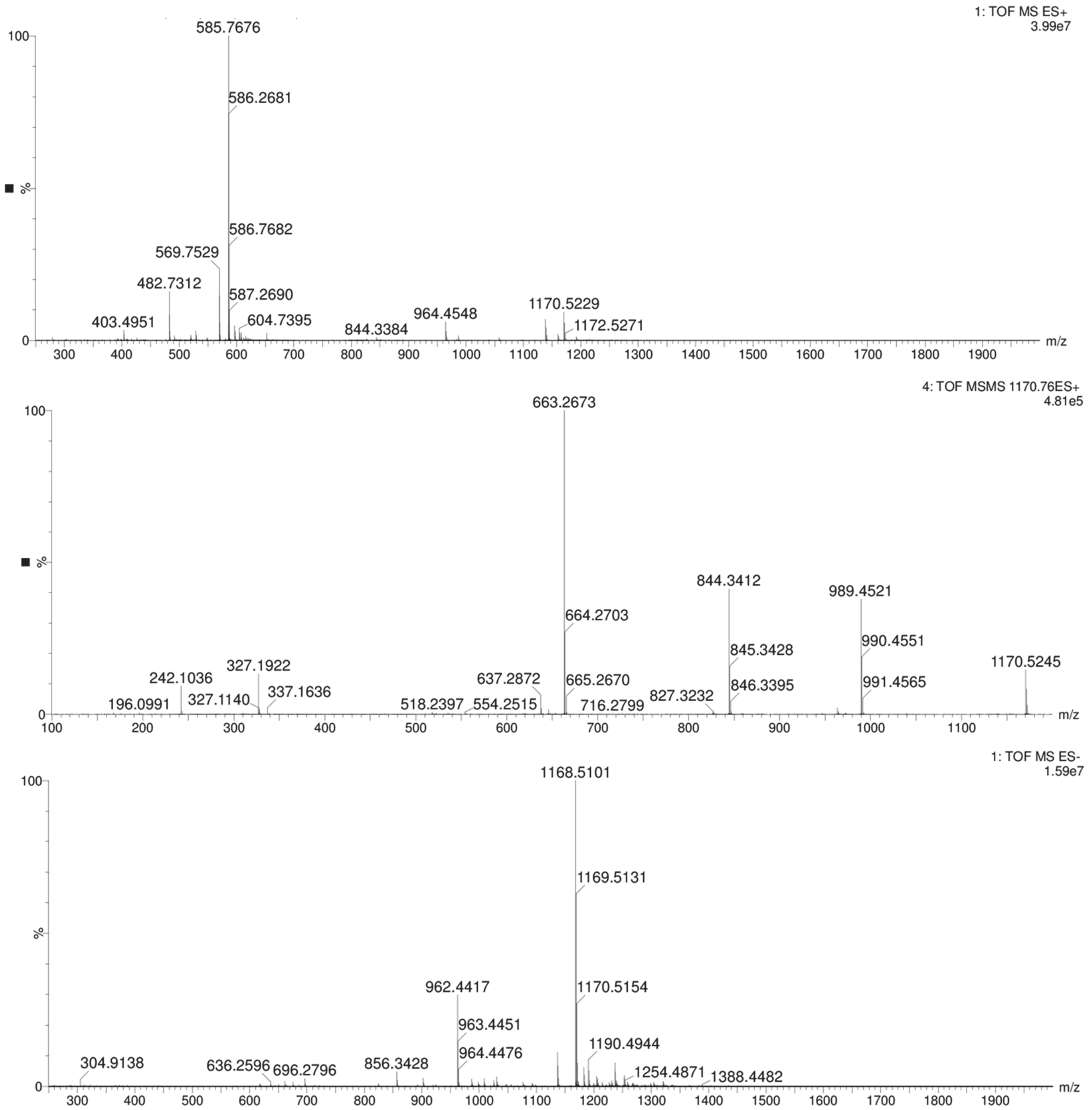

**Figure S67.** Mass spectrum of compound present in fraction C23C1C (**8-9**). QToF-MS in positive mode (top), $MS^2$ in positive mode, collision energy of 45V (middle) and MS in negative mode (bottom).

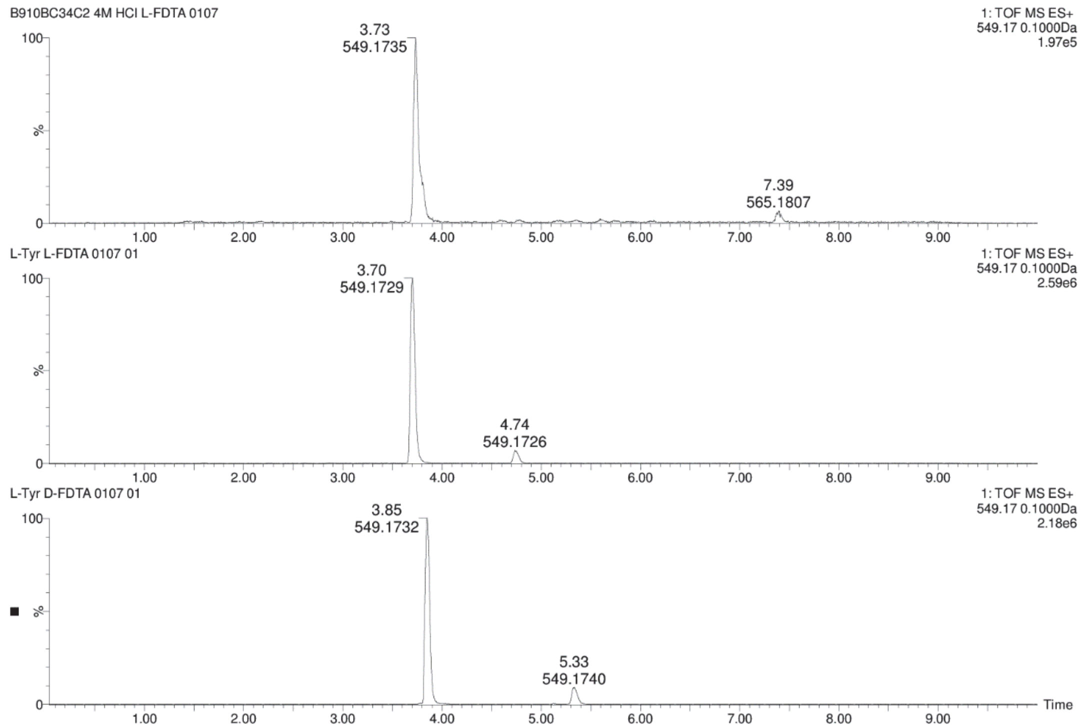

**Figure S68.** Chromatograms for comparison of **8-9** + L-FDTA (top) and L-Tyr + L-FDTA (middle) and L-Tyr + D-FDTA (bottom) in UPLC-MS analysis.

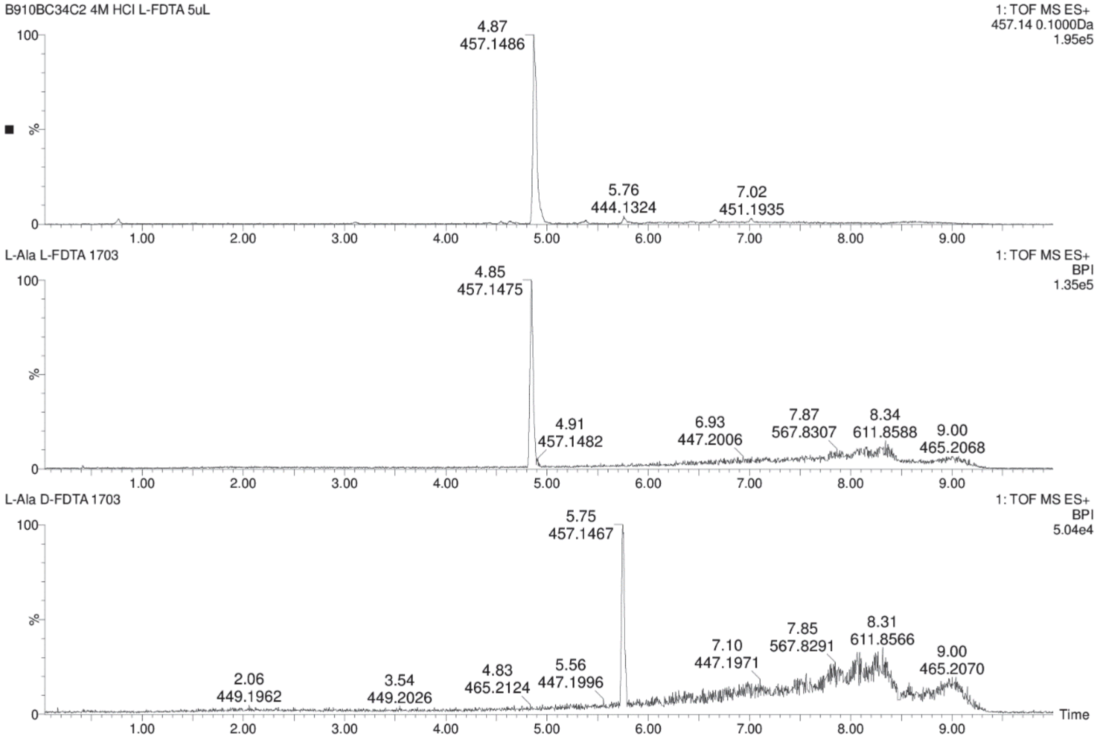

**Figure S69.** Chromatograms for comparison of **8-9** + L-FDTA (top) and L-Ala + L-FDTA (middle) and L-Ala + D-FDTA (bottom) in UPLC-MS analysis.

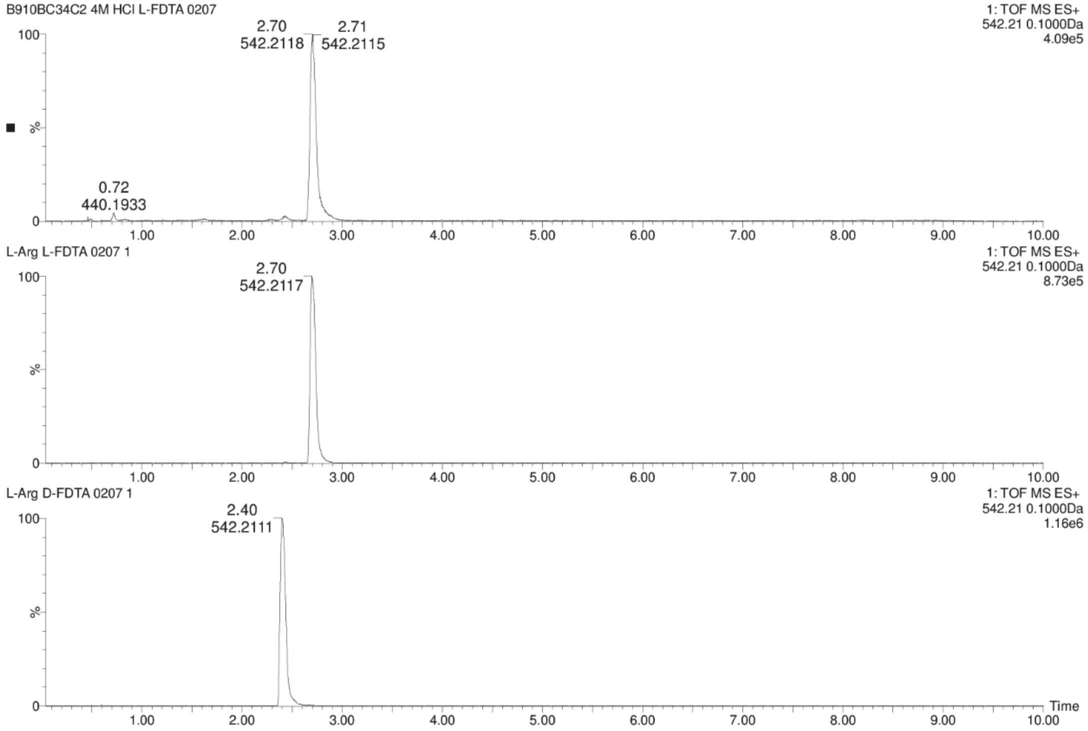

**Figure S70.** Chromatograms for comparison of **8-9** + L-FDTA (top) and L-Arg + L-FDTA (middle) and L-Arg + D-FDTA (bottom) in UPLC-MS analysis.

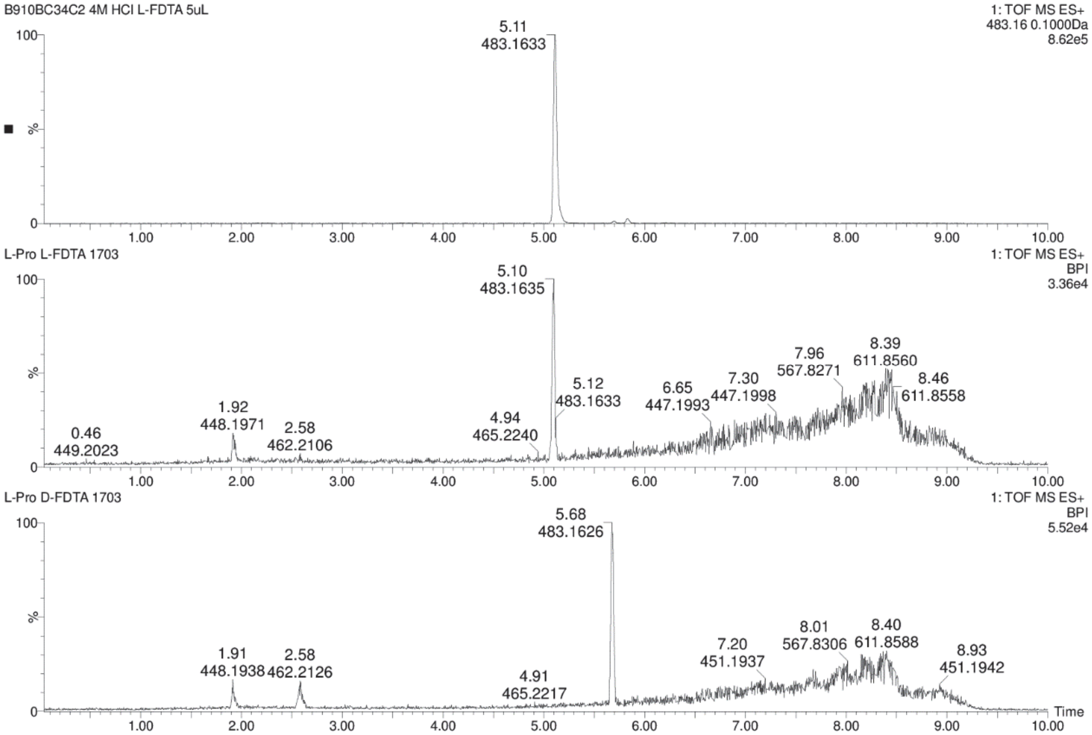

**Figure S71.** Chromatograms for comparison of **8-9** + L-FDTA (top) and L-Pro + L-FDTA (middle) and L-Pro + D-FDTA (bottom) in UPLC-MS analysis.

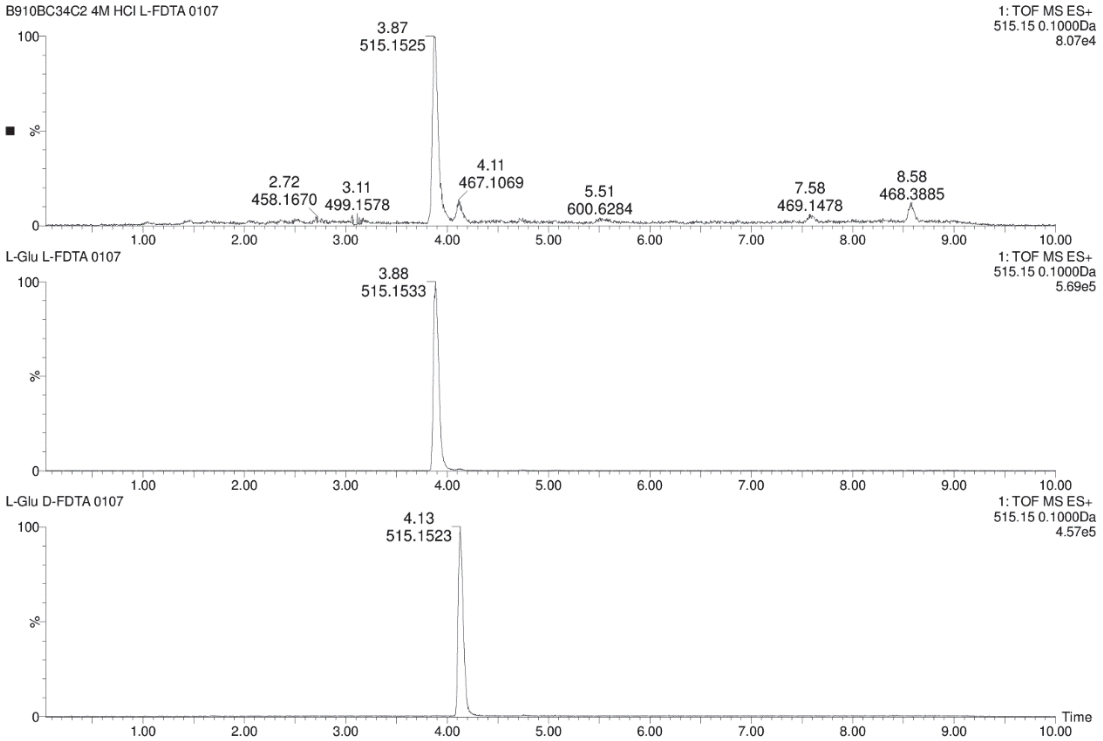

**Figure S72.** Chromatograms for comparison of **8-9** + L-FDTA (top) and L-Glu + L-FDTA (middle) and L-Glu + D-FDTA (bottom) in UPLC-MS analysis.

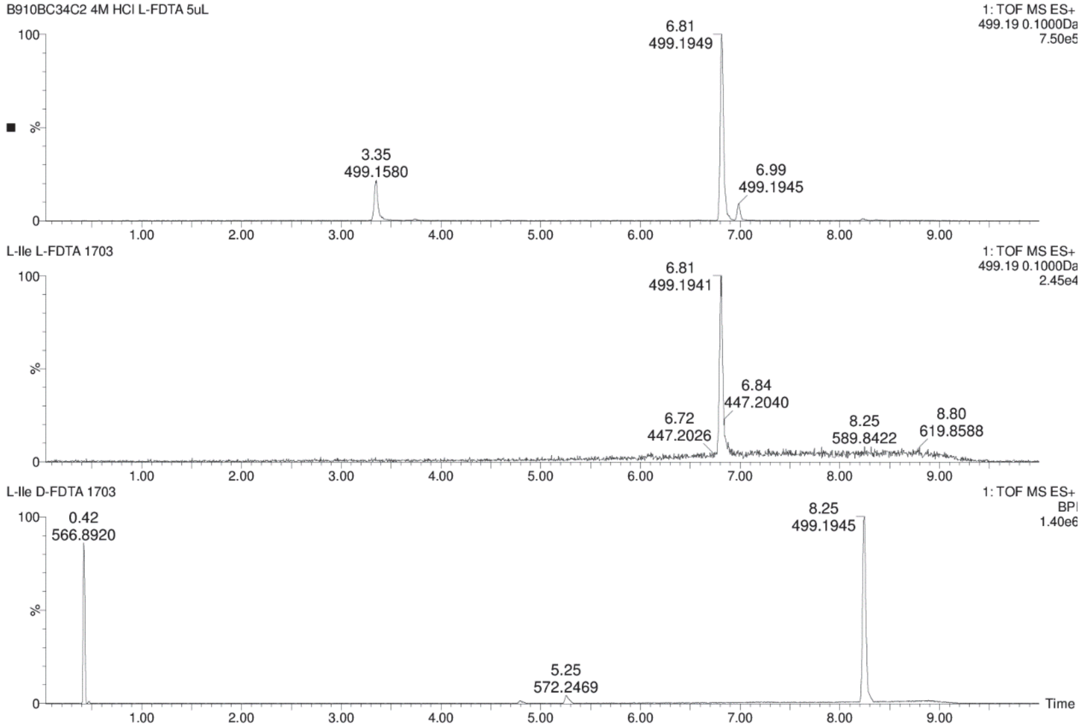

**Figure S73.** Chromatograms for comparison of **8-9** + L-FDTA (top) and L-Ile + L-FDTA (middle) and L-Ile + D-FDTA (bottom) in UPLC-MS analysis.

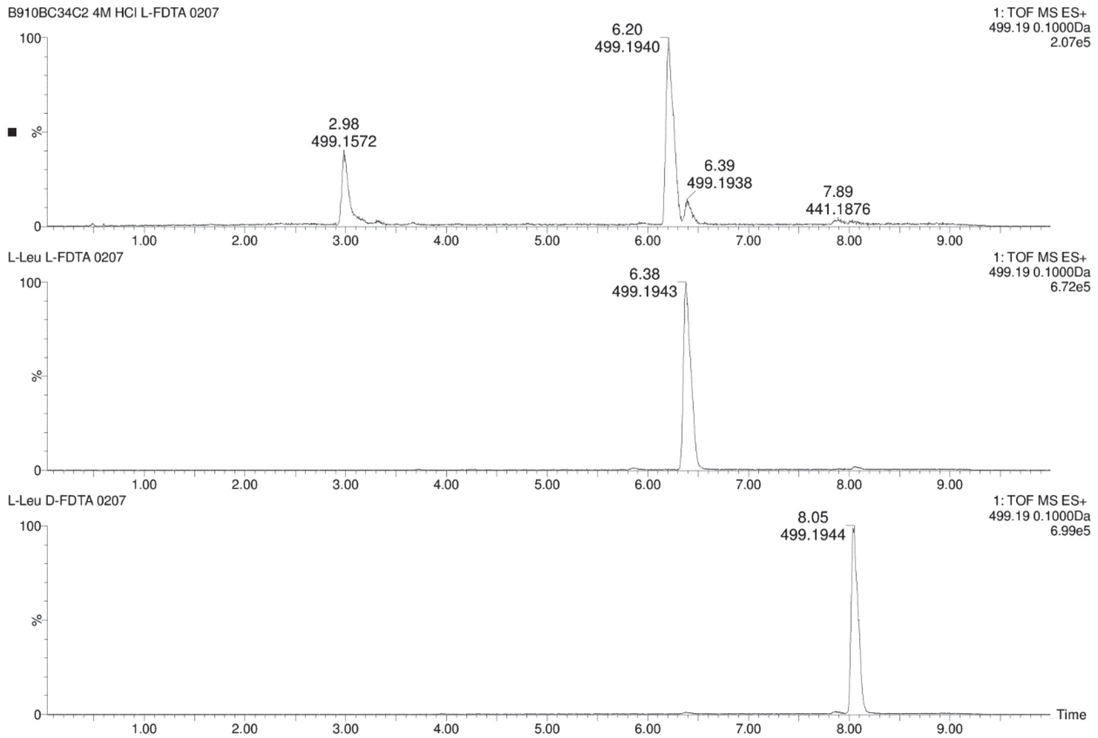

**Figure S74.** Chromatograms for comparison of **8-9** + L-FDTA (top) and L-Leu + L-FDTA (middle) and L-Leu + D-FDTA (bottom) in UPLC-MS analysis.

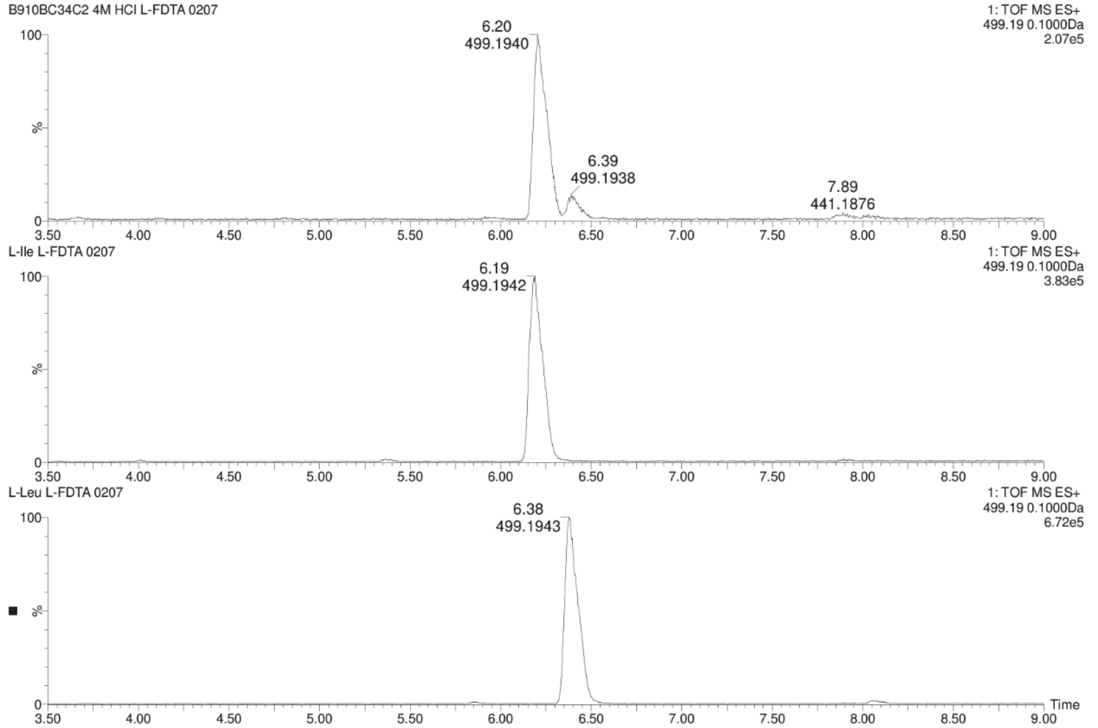

**Figure S75.** Chromatograms for comparison of **8-9** + L-FDTA (top) and L-Ile + L-FDTA (middle) and L-Leu + L-FDTA (bottom) in UPLC-MS analysis.

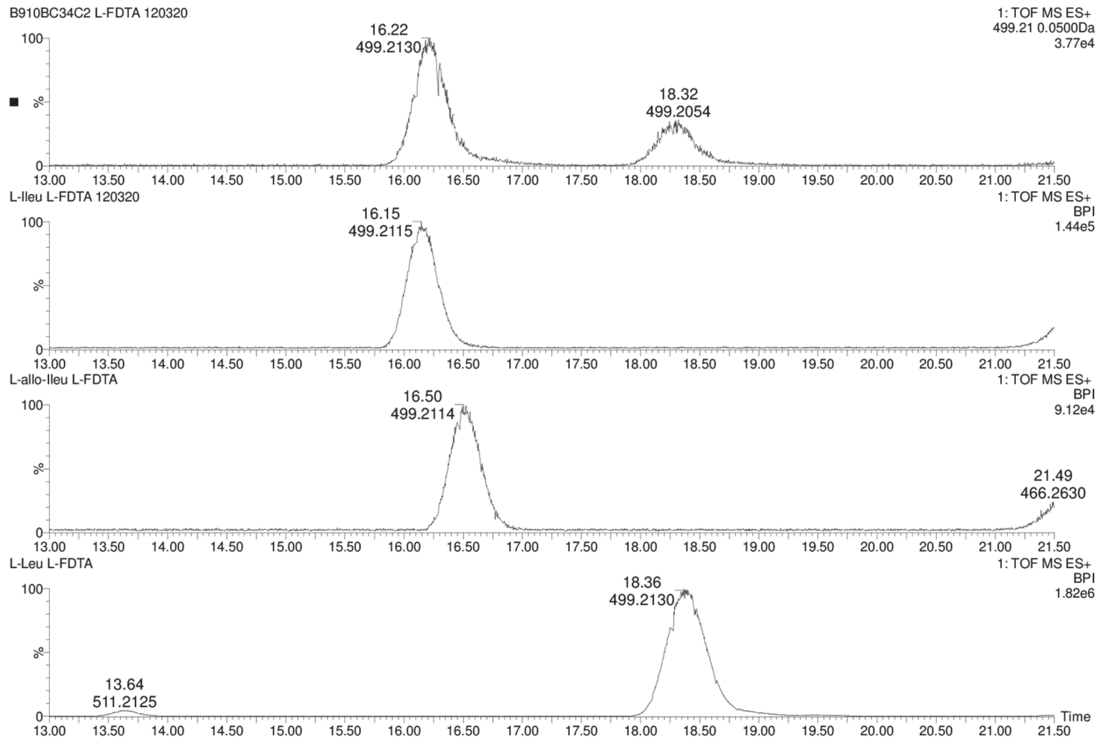

**Figure S76.** Chromatograms for comparison of **8-9** + L-FDTA (top), L-Ile + L-FDTA (2nd), L-*allo*-Ile + L-FDTA (3rd) and L-Leu + L-FDTA (bottom) in UPLC-MS analysis using analysis conditions as descried before[70].

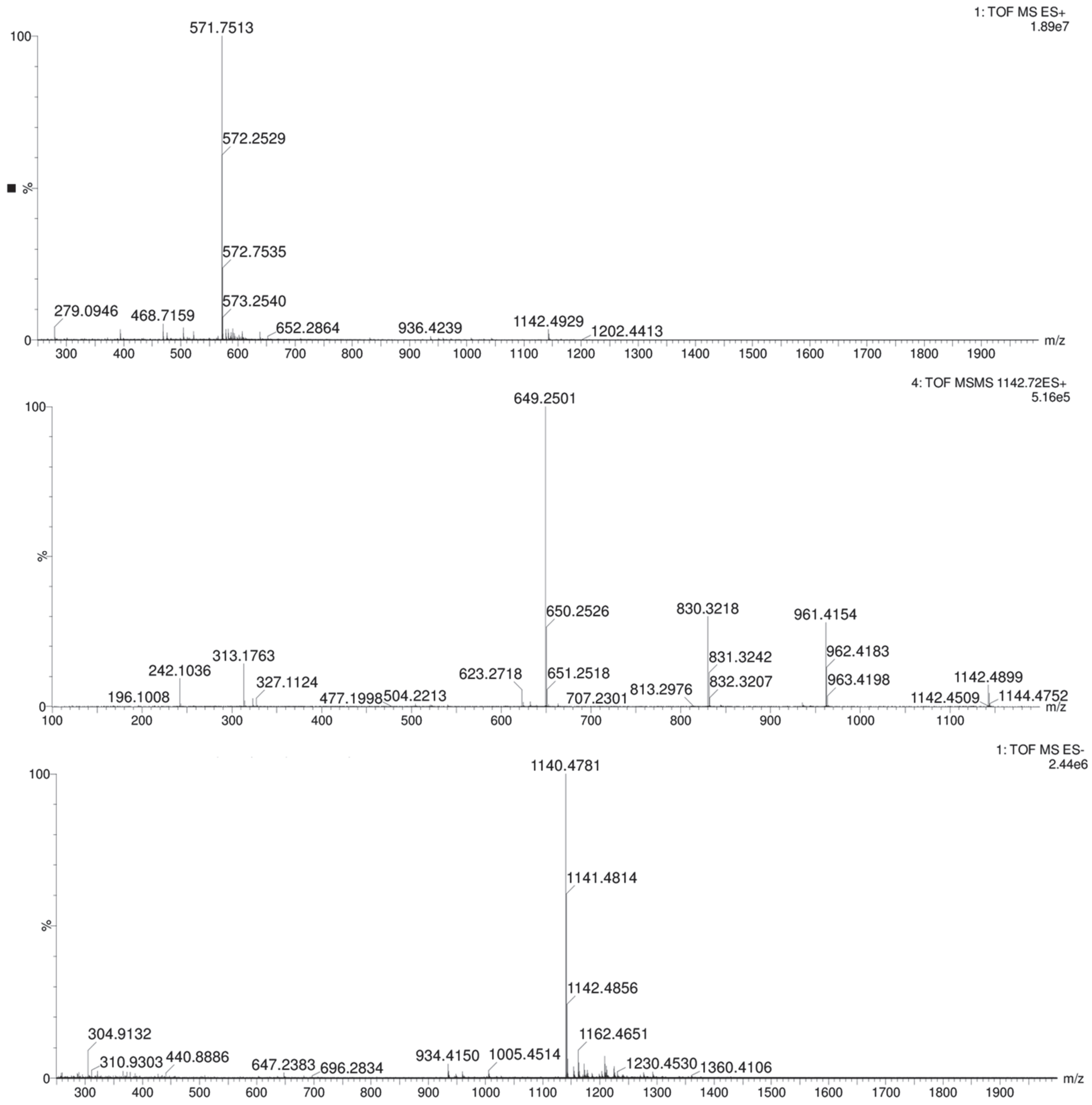

**Figure S77.** Mass spectrum of compound present in fraction C23C1A (**10**). QToF-MS in positive mode (top), $MS^2$ in positive mode, collision energy of 45V (middle) and MS in negative mode (bottom).

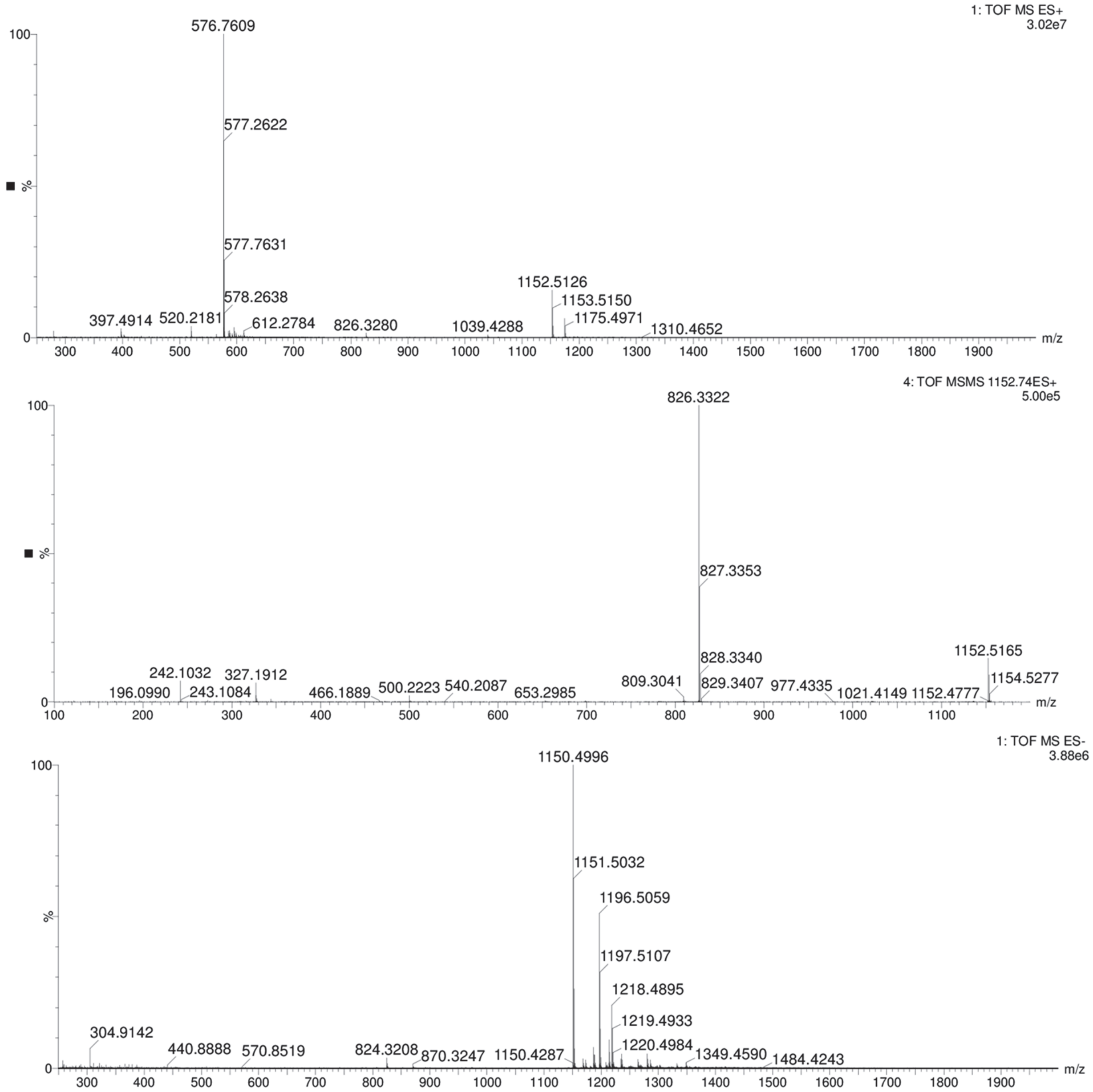

**Figure S78.** Mass spectrum of compound present in fraction C23C1D (**11-12**). QToF-MS in positive mode (top), MS$^2$ in positive mode, collision energy of 45V (middle) and MS in negative mode (bottom).

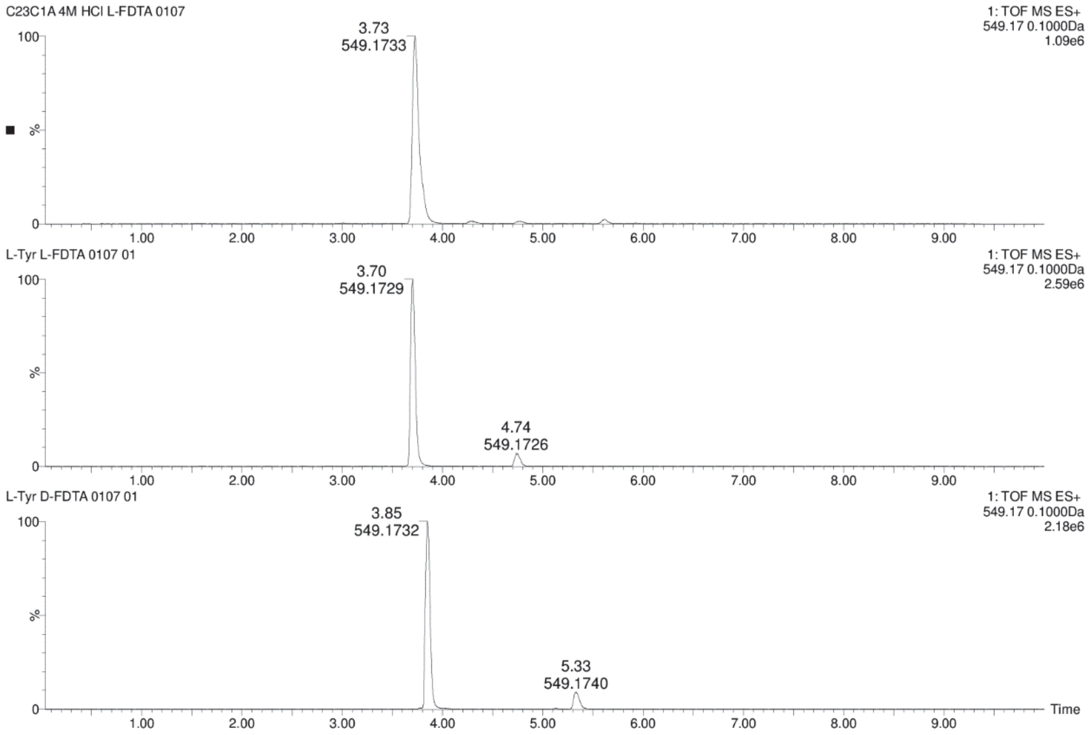

**Figure S79.** Chromatograms for comparison of **10** + L-FDTA (top) and L-Tyr + L-FDTA (middle) and L-Tyr + D-FDTA (bottom) in UPLC-MS analysis.

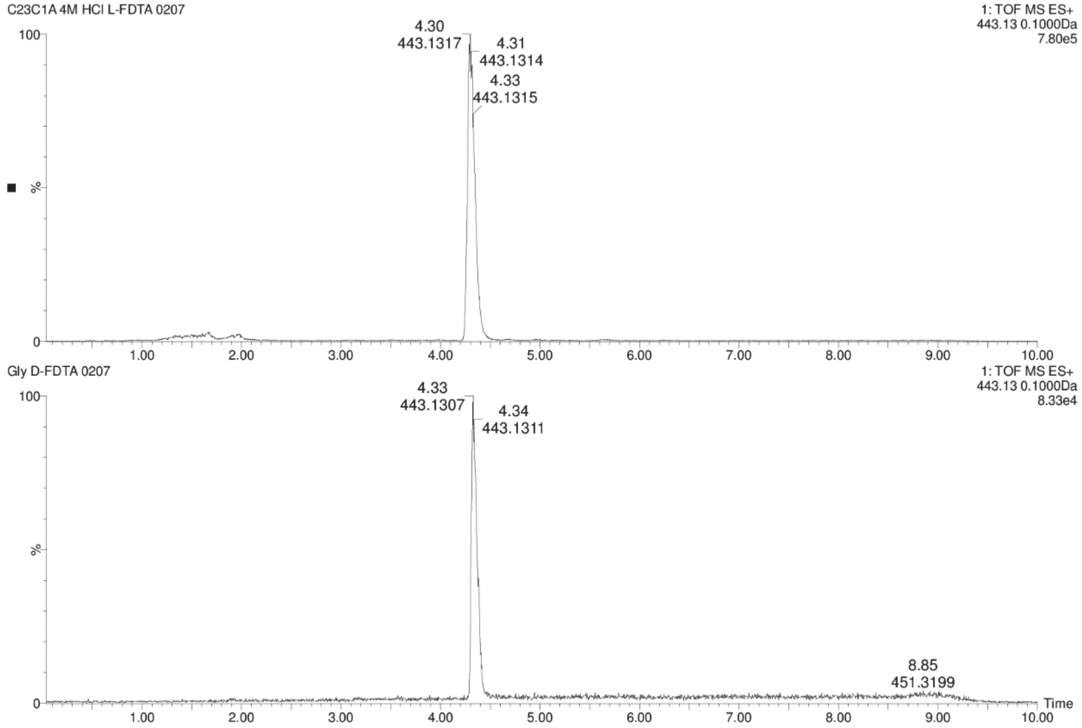

**Figure S80.** Chromatograms for comparison of **10** + L-FDTA (top) and Gly + D-FDTA (bottom) in UPLC-MS analysis.

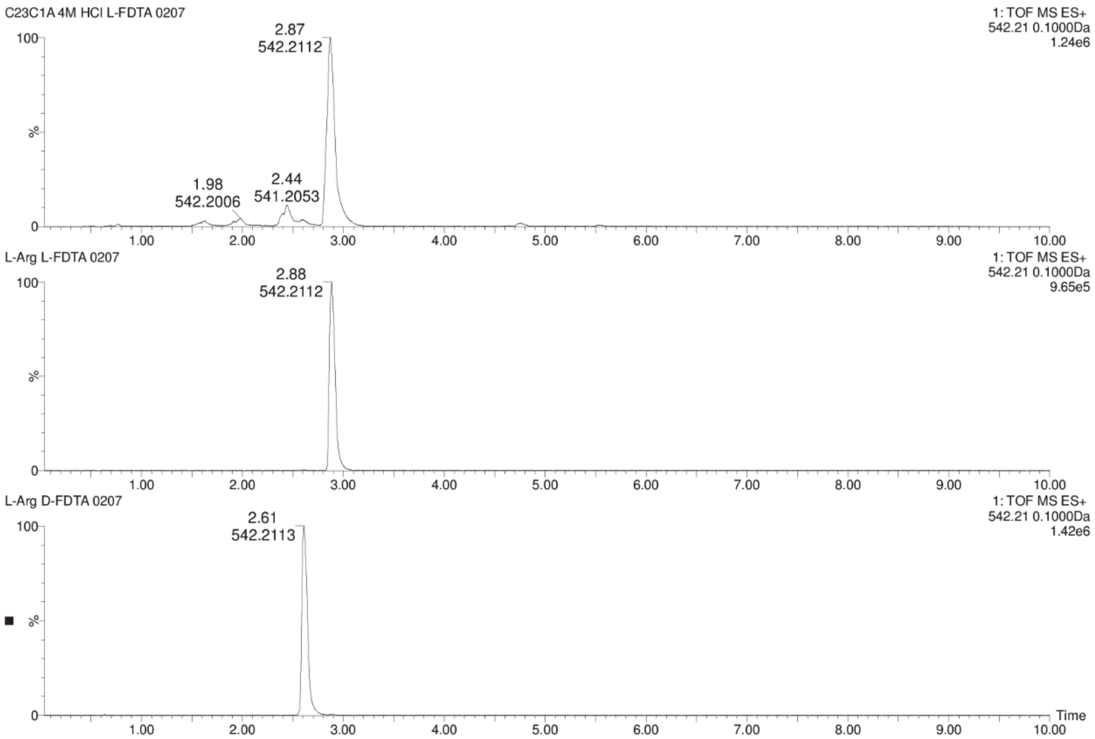

**Figure S81.** Chromatograms for comparison of **10** + L-FDTA (top) and L-Arg + L-FDTA (middle) and L-Arg + D-FDTA (bottom) in UPLC-MS analysis.

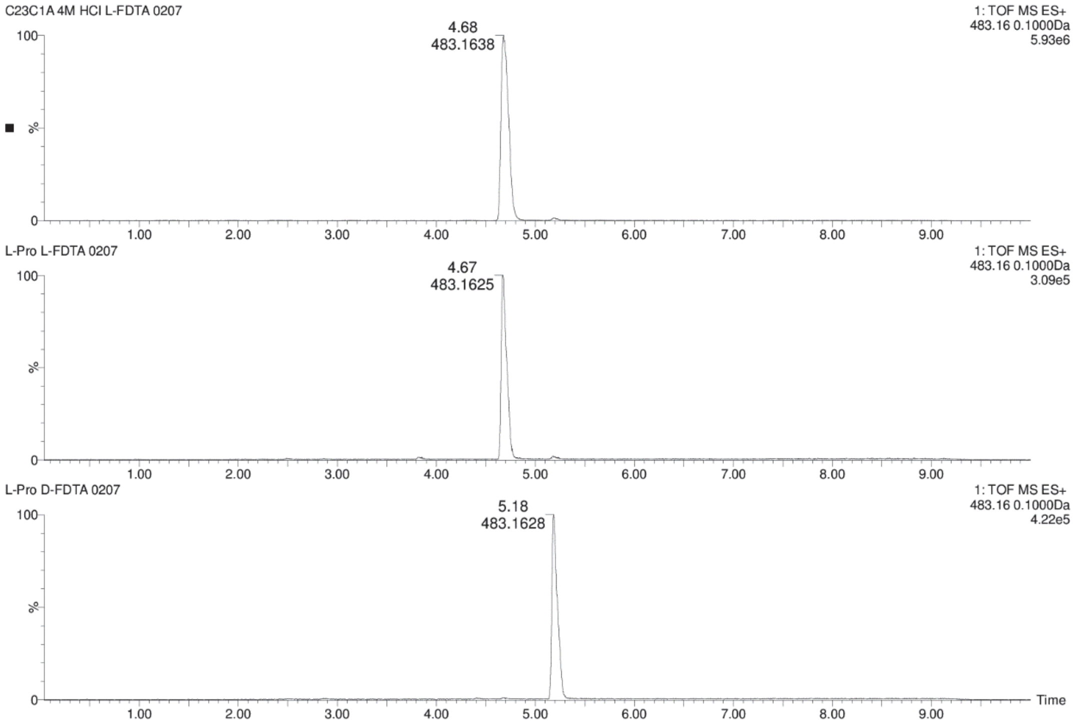

**Figure S82.** Chromatograms for comparison of **10** + L-FDTA (top) and L-Pro + L-FDTA (middle) and L-Pro + D-FDTA (bottom) in UPLC-MS analysis.

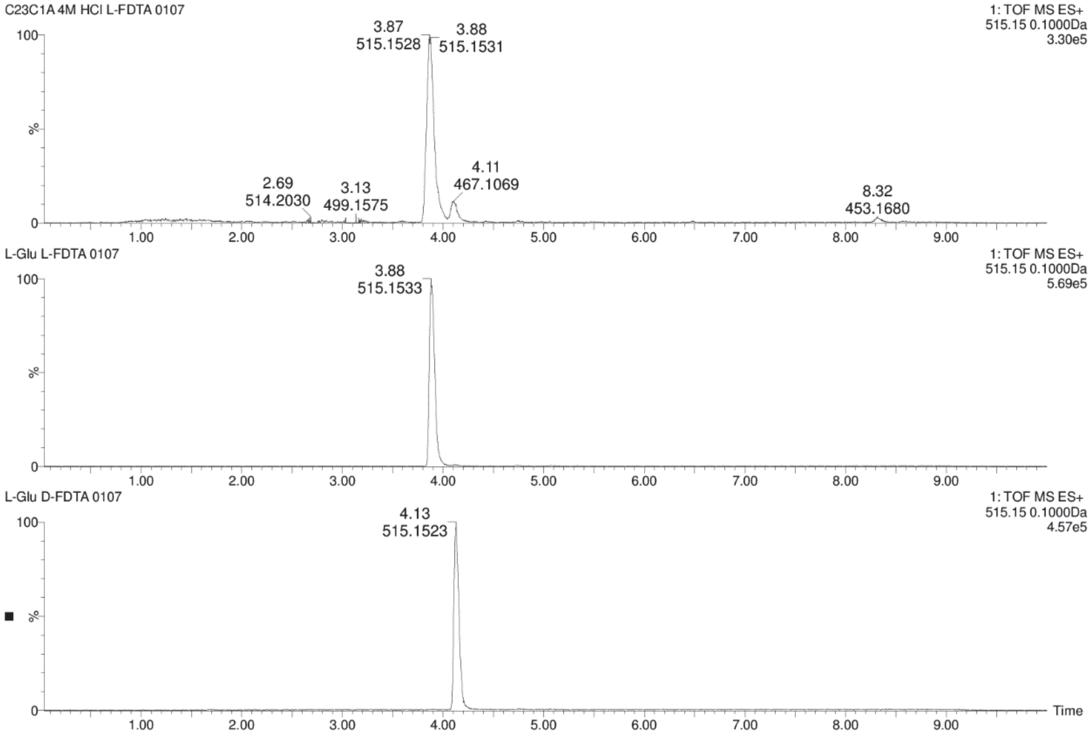

**Figure S83.** Chromatograms for comparison of **10** + L-FDTA (top) and L-Glu + L-FDTA (middle) and L-Glu + D-FDTA (bottom) in UPLC-MS analysis.

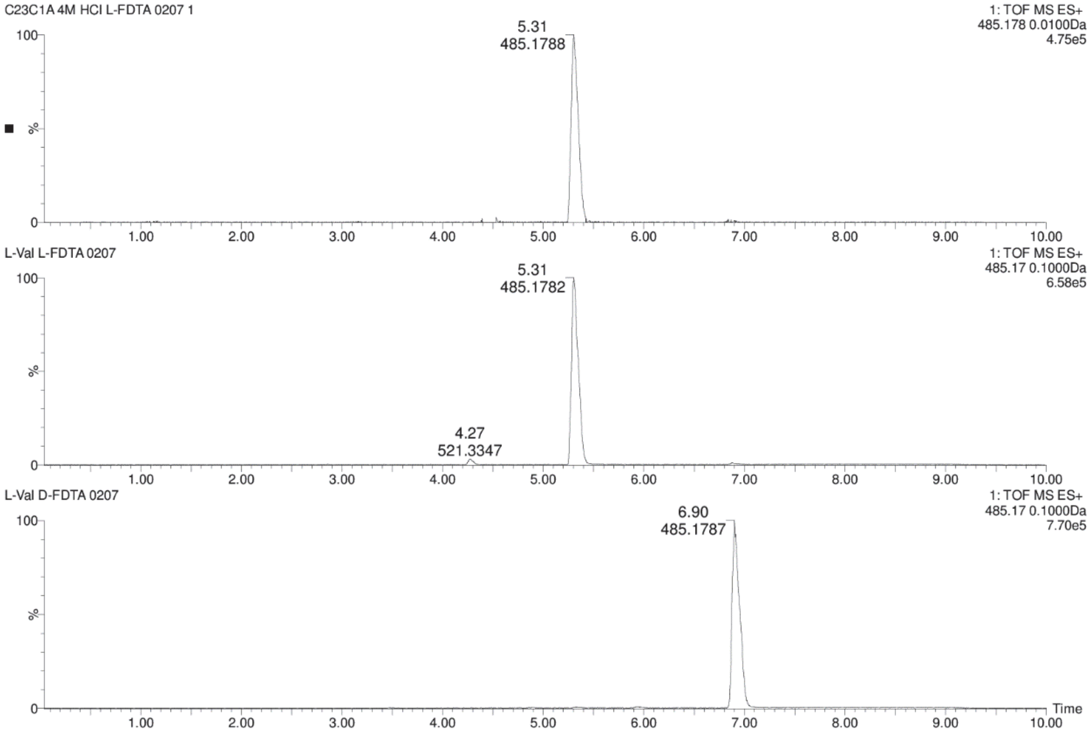

**Figure S84.** Chromatograms for comparison of **10** + L-FDTA (top) and L-Val + L-FDTA (middle) and L-Val + D-FDTA (bottom) in UPLC-MS analysis.

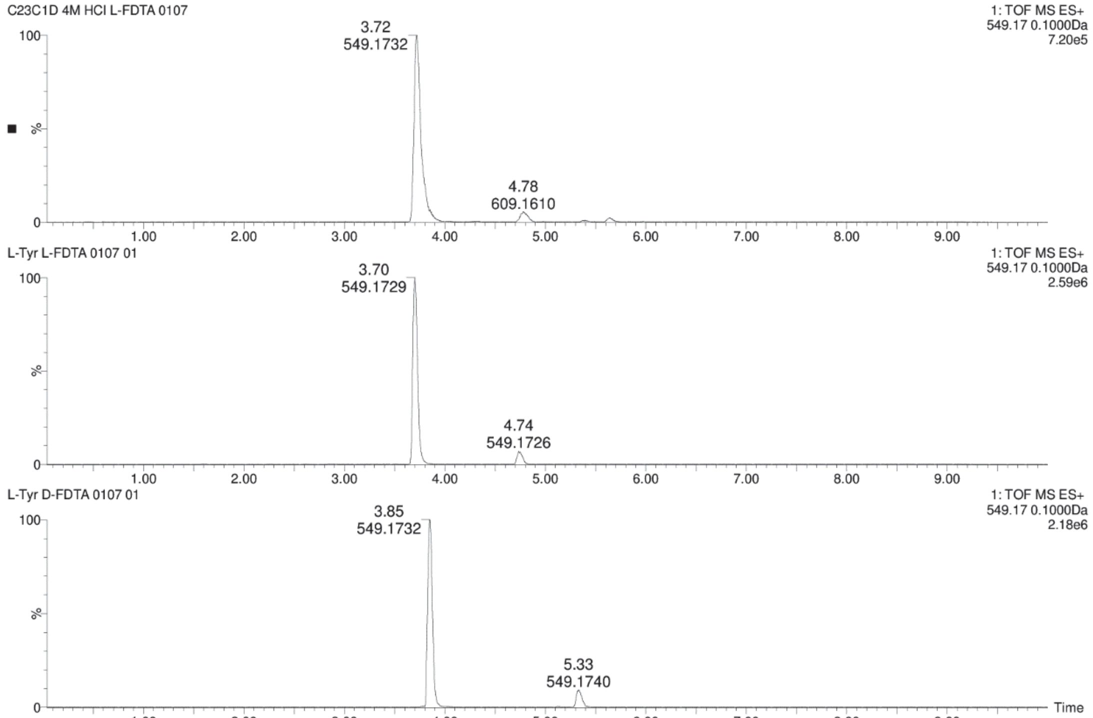

**Figure S85.** Chromatograms for comparison of **11-12** + L-FDTA (top) and L-Tyr + L-FDTA (middle) and L-Tyr + D-FDTA (bottom) in UPLC-MS analysis.

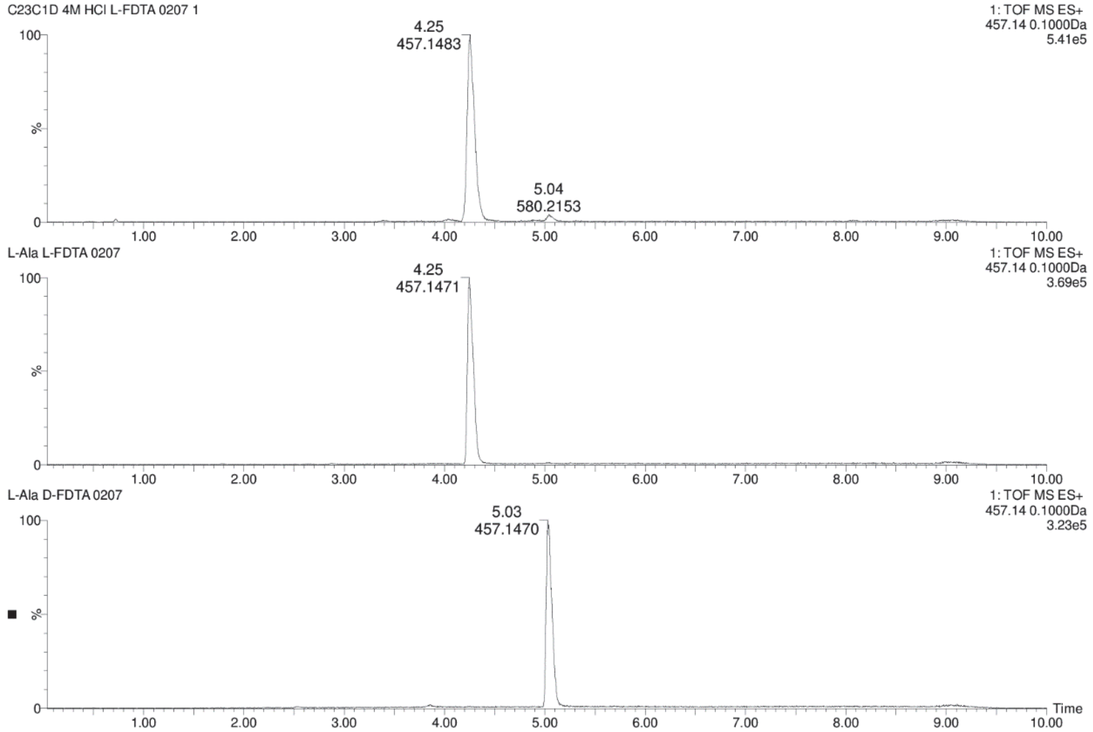

**Figure S86.** Chromatograms for comparison of **11-12** + L-FDTA (top) and L-Ala + L-FDTA (middle) and L-Ala + D-FDTA (bottom) in UPLC-MS analysis.

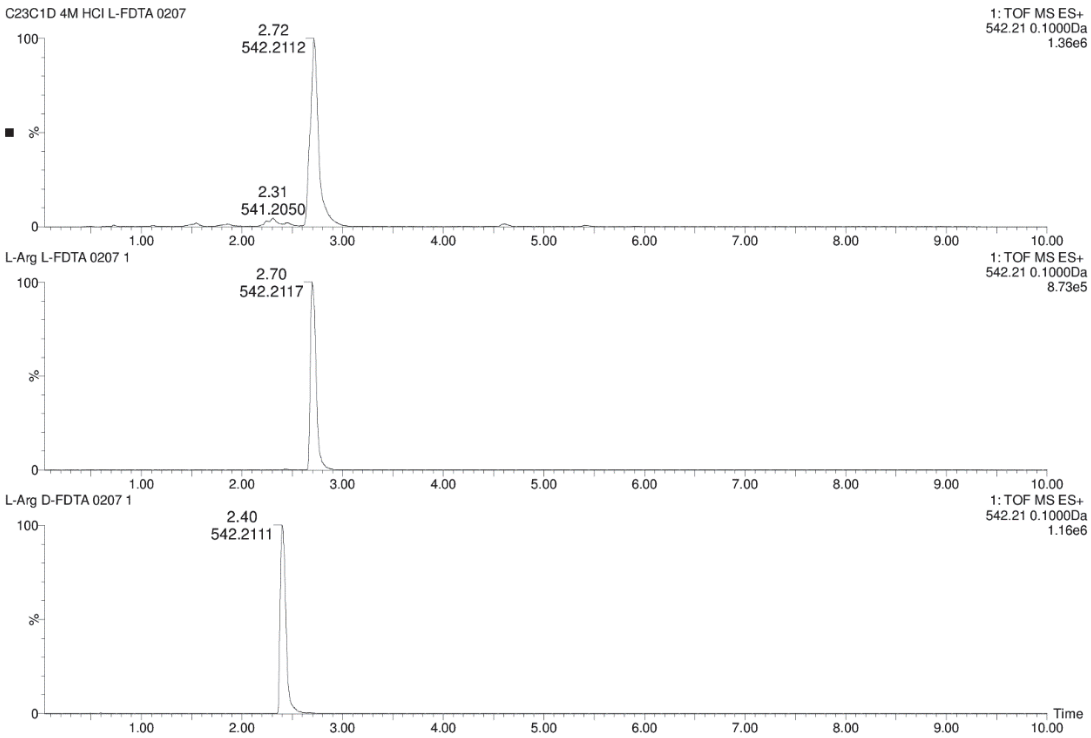

**Figure S87.** Chromatograms for comparison of **11-12** + L-FDTA (top) and L-Arg + L-FDTA (middle) and L-Arg + D-FDTA (bottom) in UPLC-MS analysis.

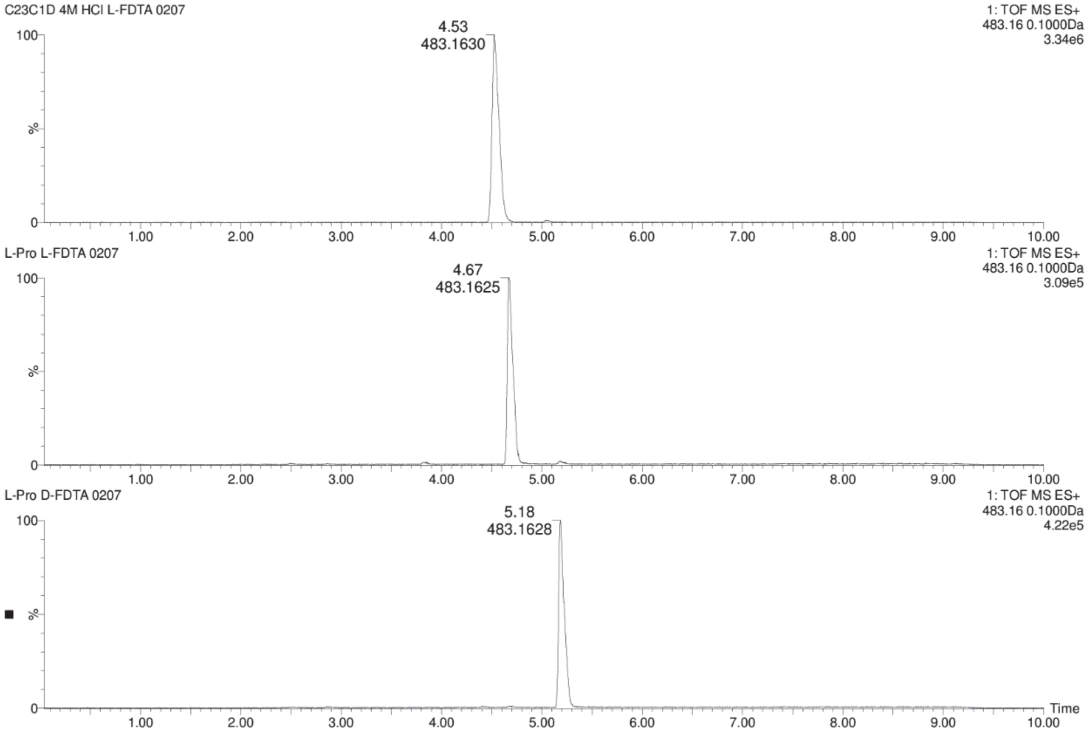

**Figure S88.** Chromatograms for comparison of **11-12** + L-FDTA (top) and L-Pro + L-FDTA (middle) and L-Pro + D-FDTA (bottom) in UPLC-MS analysis.

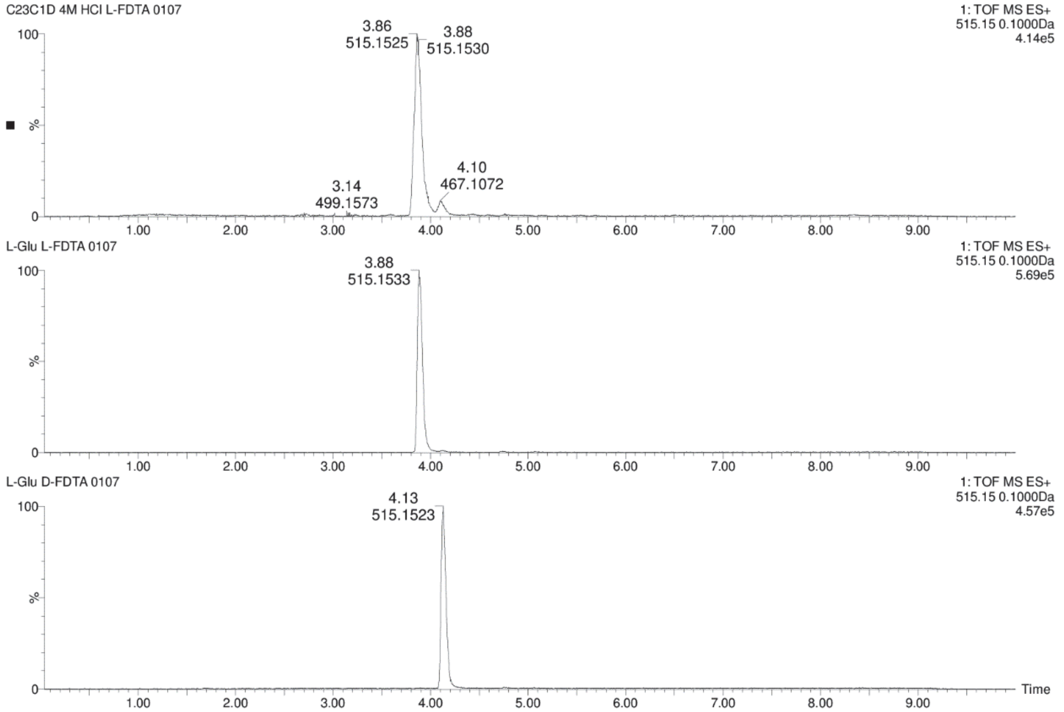

**Figure S89.** Chromatograms for comparison of **11-12** + L-FDTA (top) and L-Glu + L-FDTA (middle) and L-Glu + D-FDTA (bottom) in UPLC-MS analysis.

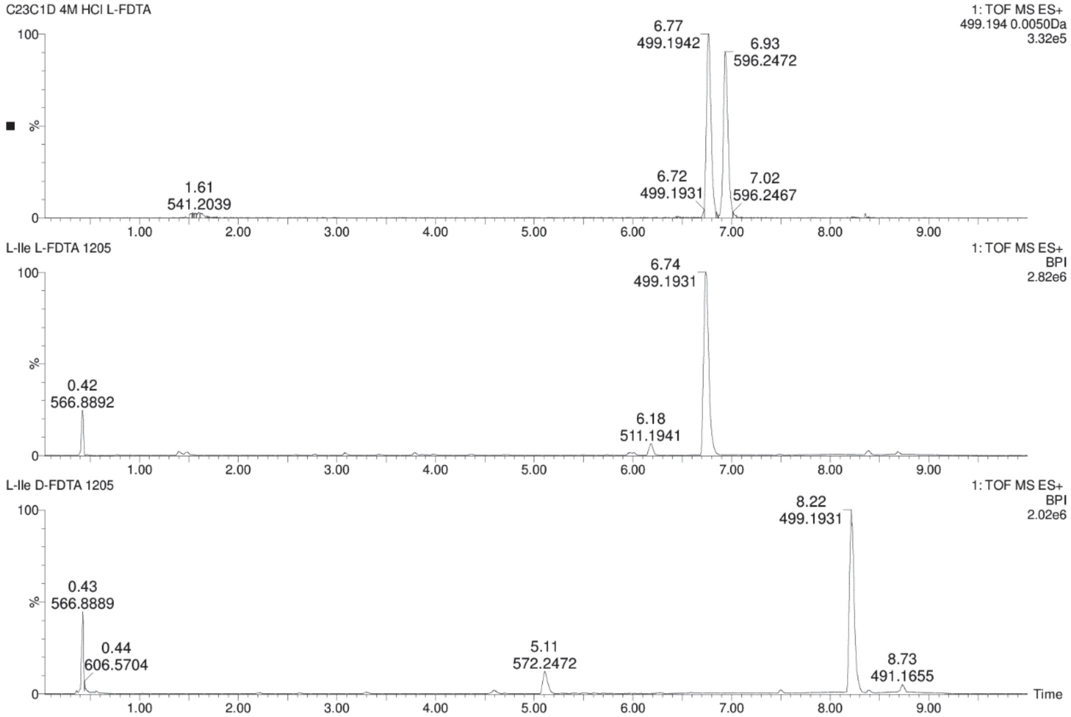

**Figure S90.** Chromatograms for comparison of **11-12** + L-FDTA (top) and L-Ile + L-FDTA (middle) and L-Ile + D-FDTA (bottom) in UPLC-MS analysis.

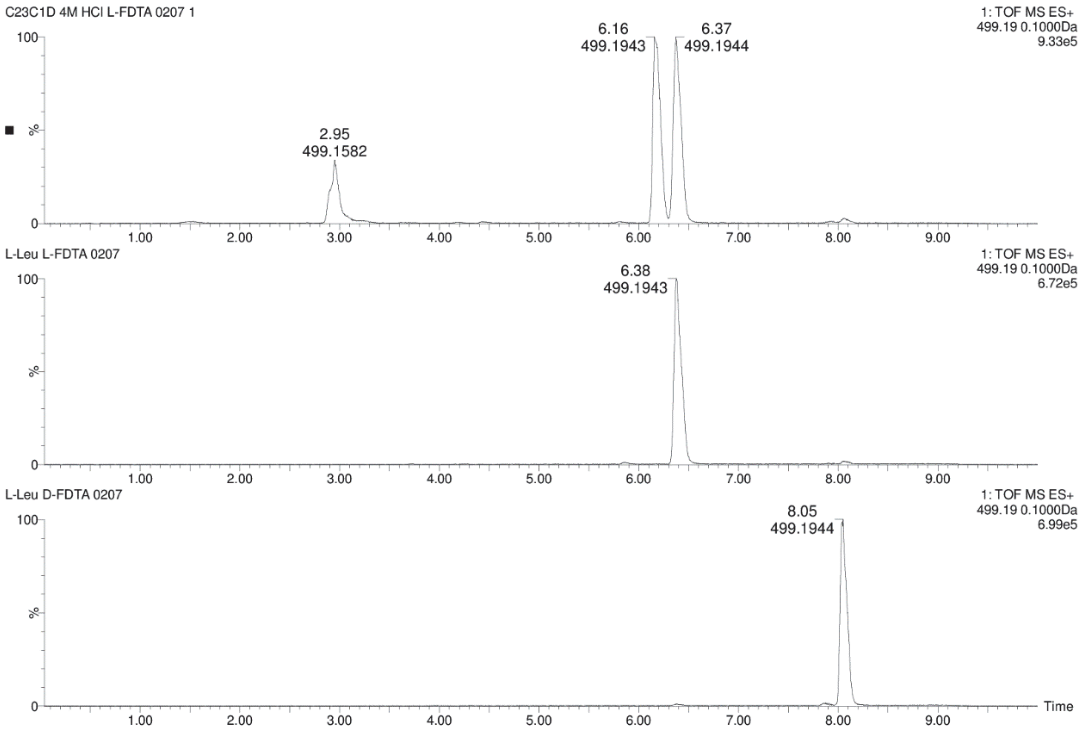

**Figure S91.** Chromatograms for comparison of **11-12** + L-FDTA (top) and L-Leu + L-FDTA (middle) and L-Leu + D-FDTA (bottom) in UPLC-MS analysis.

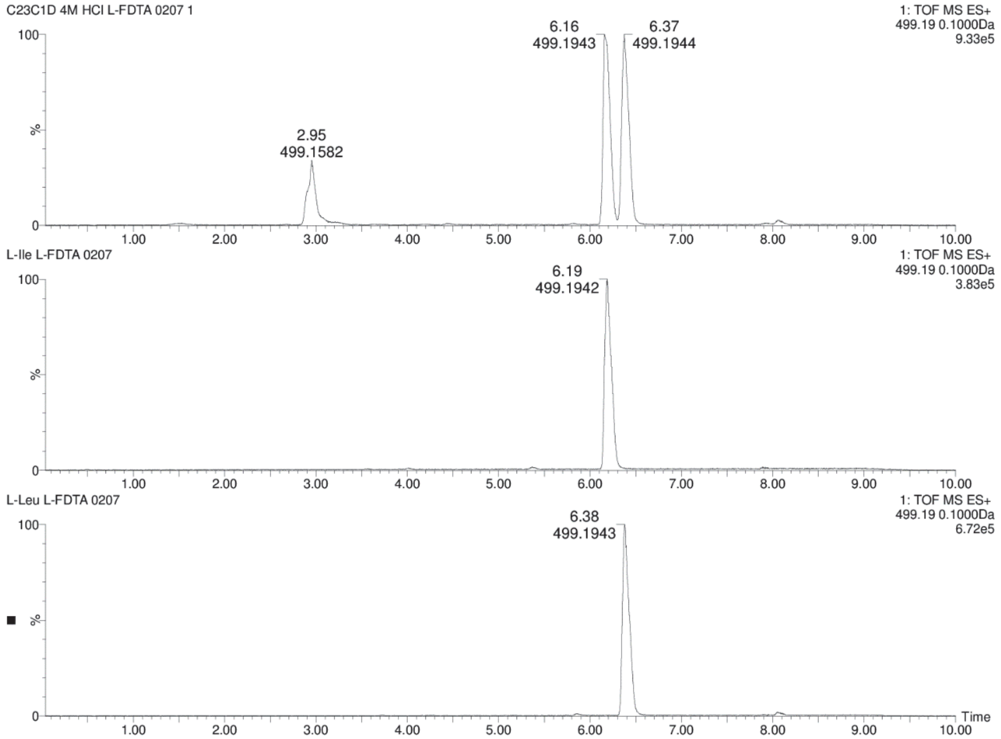

**Figure S92.** Chromatograms for comparison of **11-12** + L-FDTA (top) and L-Ile + L-FDTA (middle) and L-Leu + L-FDTA (bottom) in UPLC-MS analysis.

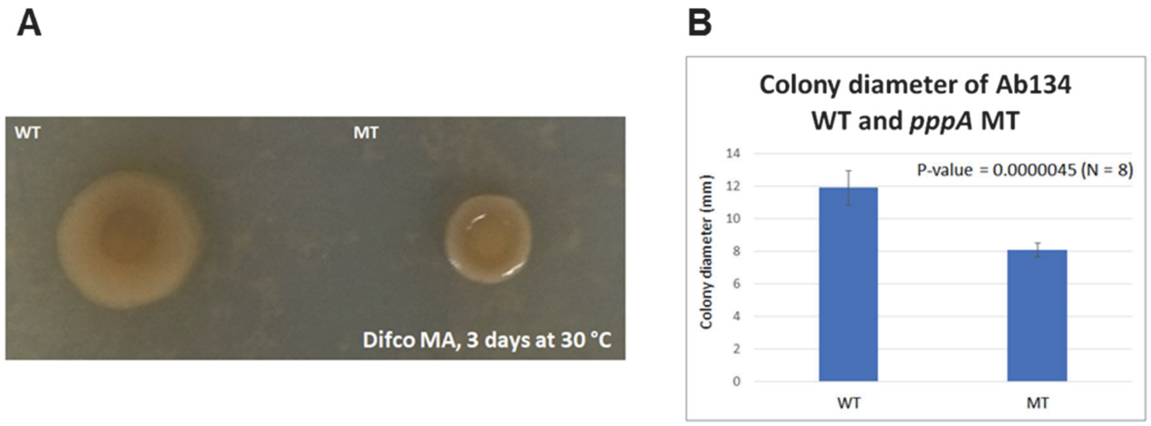

**Figure S93.** Colony diameter comparison. A) Difference in colony size between *P. brasiliensis* Ab134 WT and *pppA* mutant after spotting a 10-µl drop of liquid culture on marine agar and incubating for 3 days. B) Quantification of colony diameter of *P. brasiliensis* Ab134 WT and *pppA* mutant (N=8 each).

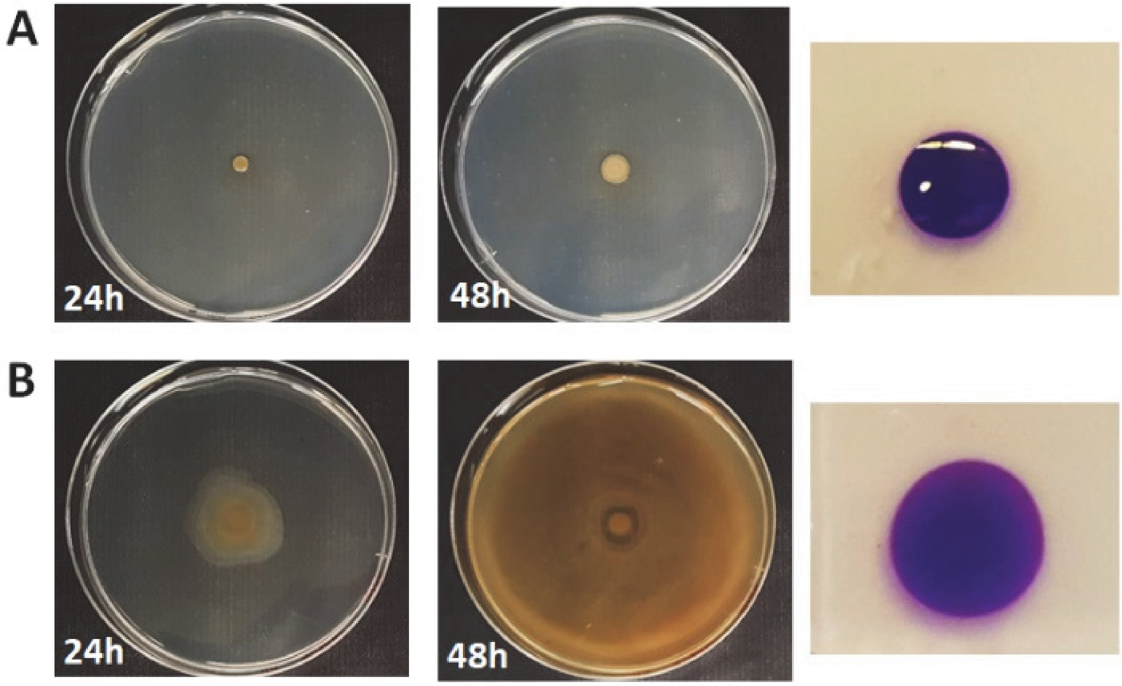

**Figure S94.** Establishment of swarming assay for *P. brasiliensis* Ab134. A) From left to right: pictures of *P. brasiliensis* Ab134 WT inoculated on the center of MB + 0.5 % Bacto agar at 24h and 48h after inoculation, respectively; picture of water + crystal violet droplet on MA (1.2 % Bacto agar). B) From left to right: pictures of *P. brasiliensis* Ab134 WT inoculated on the center of MB + 0.5 % Eiken agar at 24h and 48h after inoculation, respectively; picture of water + crystal violet droplet at MB + 1.2 % Eiken agar.

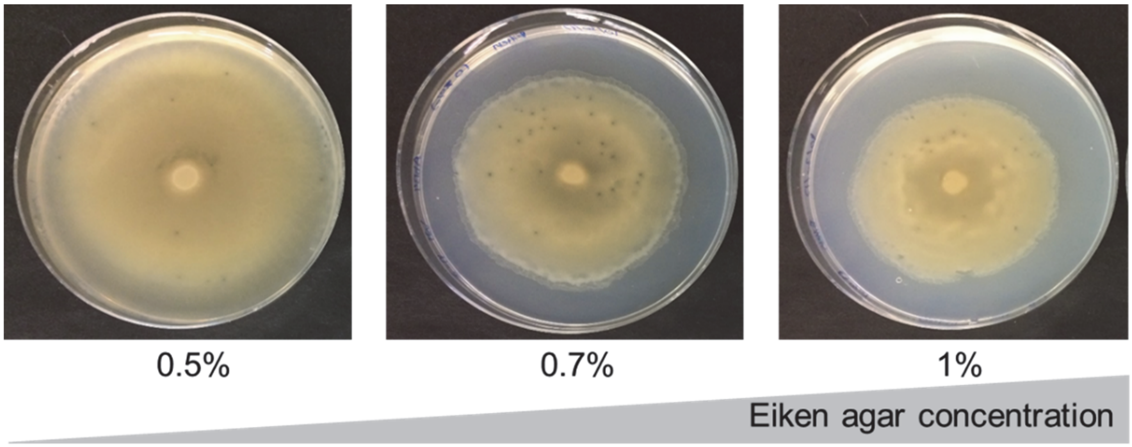

**Figure S95.** Effect of agar concentration on the swarming phenotype of *P. brasiliensis* Ab134 WT. A 10-µl drop of cryopreserved culture was inoculated on the center of MB containing Eiken agar at the indicated concentration and incubated at 30 °C. Pictures were taken 48h after inoculation.

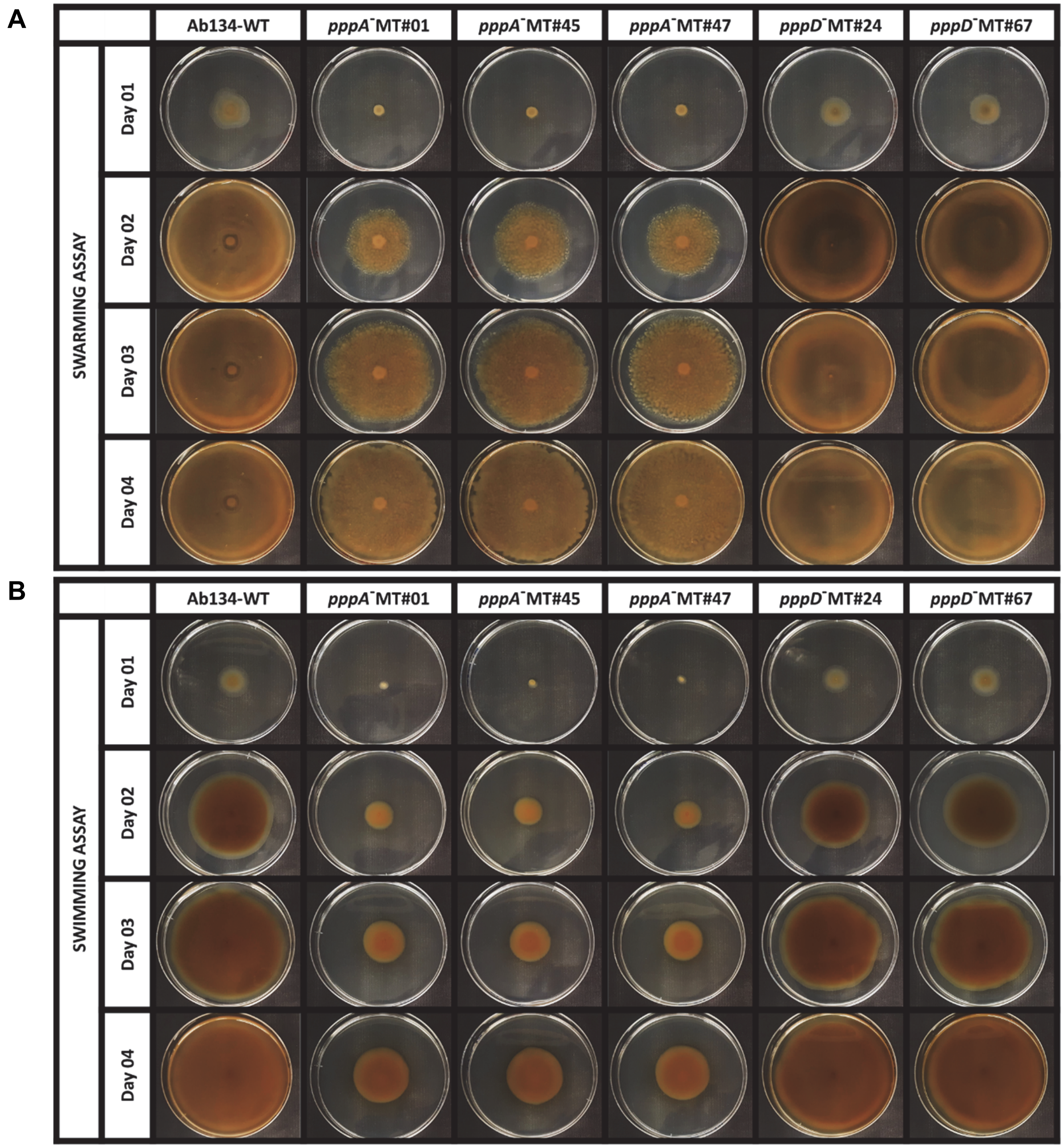

**Figure S96.** Effect of *pppA* and *pppD* deletion on flagellar motility of independent mutant clones. (A) swarming assay and (B) swimming assay. Assays were performed on marine agar containing either 0.5% or 0.3% Eiken agar, respectively. Plates were inoculated with 5 µL of cryo-preserved cultures of Ab134 wild-type (WT) or *pppA*$^-$/*pppD*$^-$ mutant strains normalized to $OD_{600}$ of 1.0. Day 1 corresponds to 24 h incubation.

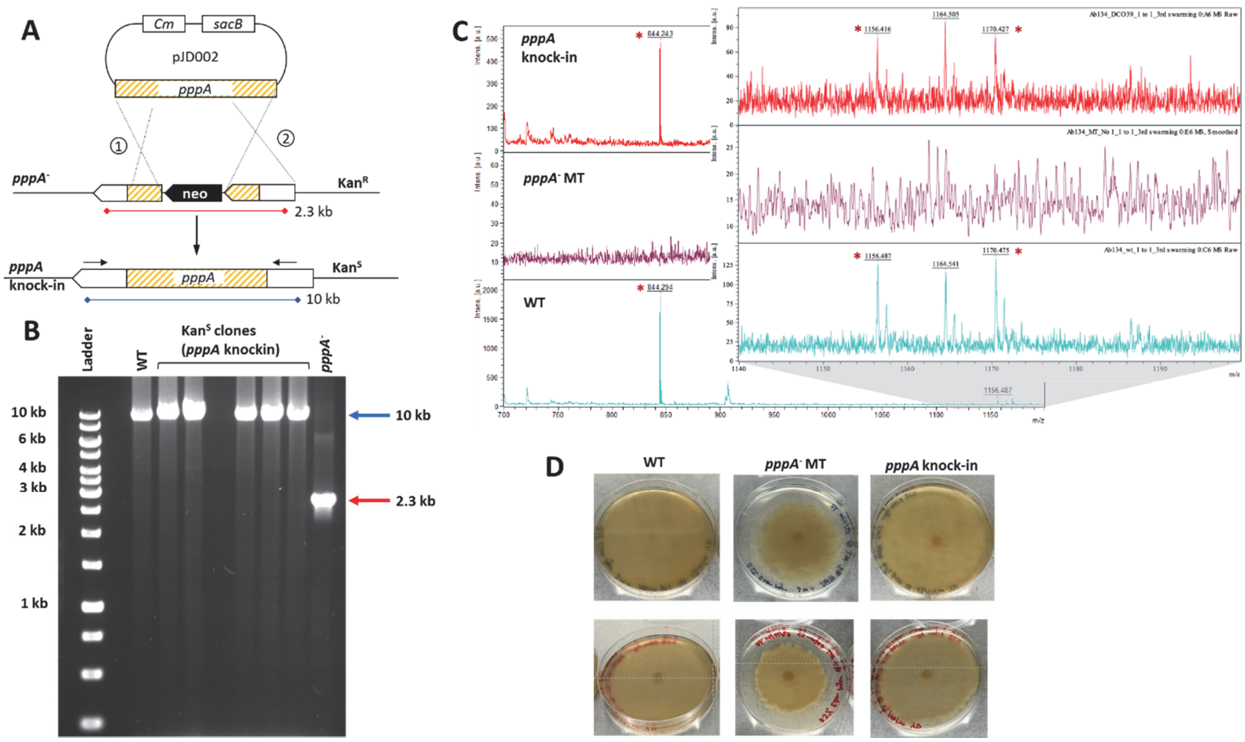

**Figure S97.** *pppA* genetic complementation. A) Design of gene replacement experiments. *neo*, neomycin/kanamycin resistance marker. Cm, chloramphenicol resistance gene. Kan$^{R/S}$, kanamycin resistance / sensitivity. Numbers 1 and 2 indicate crossover events. Black arrows indicate primer location. B) PCR analysis of Kan$^{S}$, *pppA* double crossover mutants. Expected fragment length for *pppA* knock-in mutants (i.e. reversion to the wild-type genotype), 10 kb. Wild-type (WT) genomic DNA was used as positive control. Expected fragment length for the *pppA*$^{-}$ parent, 2.3 kb. Ladder, GeneRuler 1kb DNA Ladder from ThermoScientific. C) MALDI-TOF dried-droplet mass spectrometry analysis of a representative *pppA* knock-in mutant (top) in comparison to the *pppA*$^{-}$ mutant (middle) and the wild-type (bottom) strains. D) Swarming assay. Assays were performed on marine agar containing 0.5% Eiken agar. Plates were inoculated with 5 µL of $OD_{600}$-normalized cultures from Ab134 wild-type (WT), *pppA*$^{-}$ mutant, or *pppA* knock-in mutant strains, respectively. Pictures were taken after three days at 30 °C. The assay was run twice, each time in triplicates. Top and bottom images show representative plates from each independent experiment that used either fresh seed cultures (top) or cryopreserved cultures (bottom).

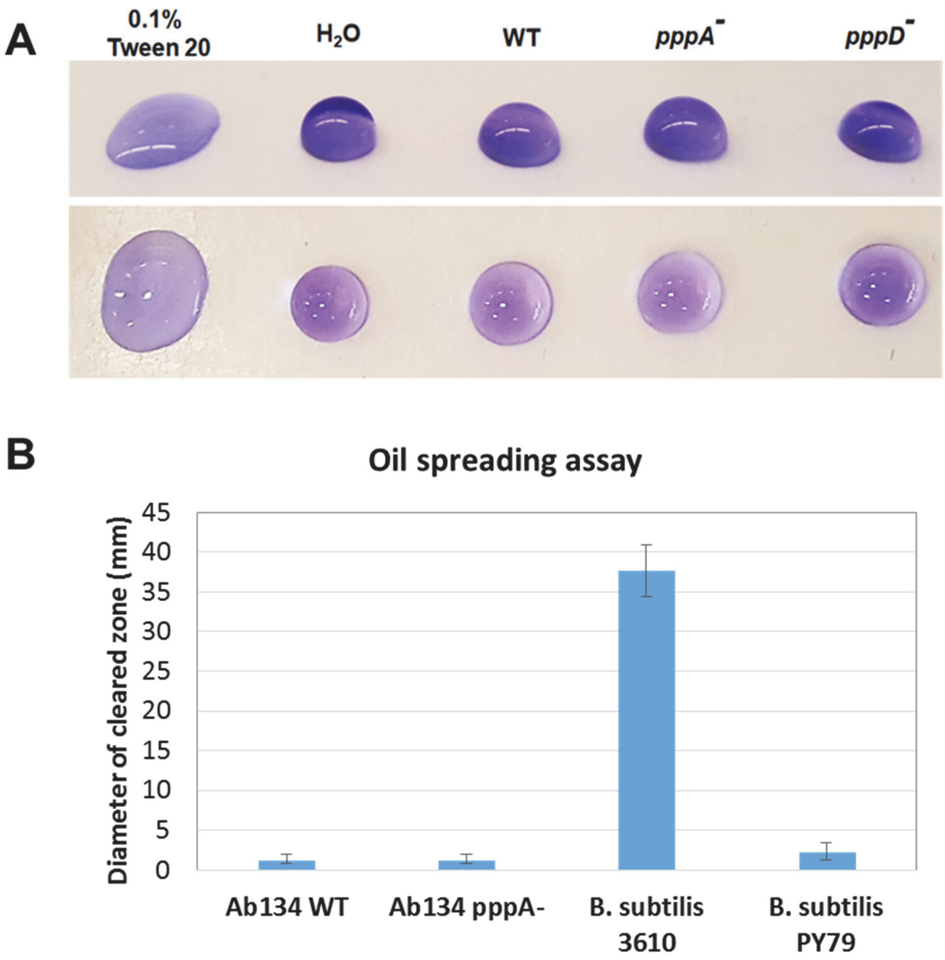

**Figure S98.** Surfactant activity assays. (**A**) Drop collapse assay for *P. brasiliensis* Ab134 WT and *pppA* and *pppD* mutants. A solution of 0.1% of Tween 20 was used as positive control and water as negative control. Crystal violet at 0.002% was added to aid visualization. (**B**) Oil spreading assay. A film of mineral oil was placed on top of water on a Petri dish. A drop of supernatants from overnight liquid cultures of either *P. brasiliensis* Ab134 WT or the *pppA* mutant was dropped on top of the oil and the diameter of the cleared zone was measured. *Bacillus subtilis* 3610 (a producer of surfactin, a lipopeptide biosurfactant) was used as positive control, and the surfactin-defective strain PY79 was used as negative control. The experiment was performed in triplicates. Error bars represent standard deviation.

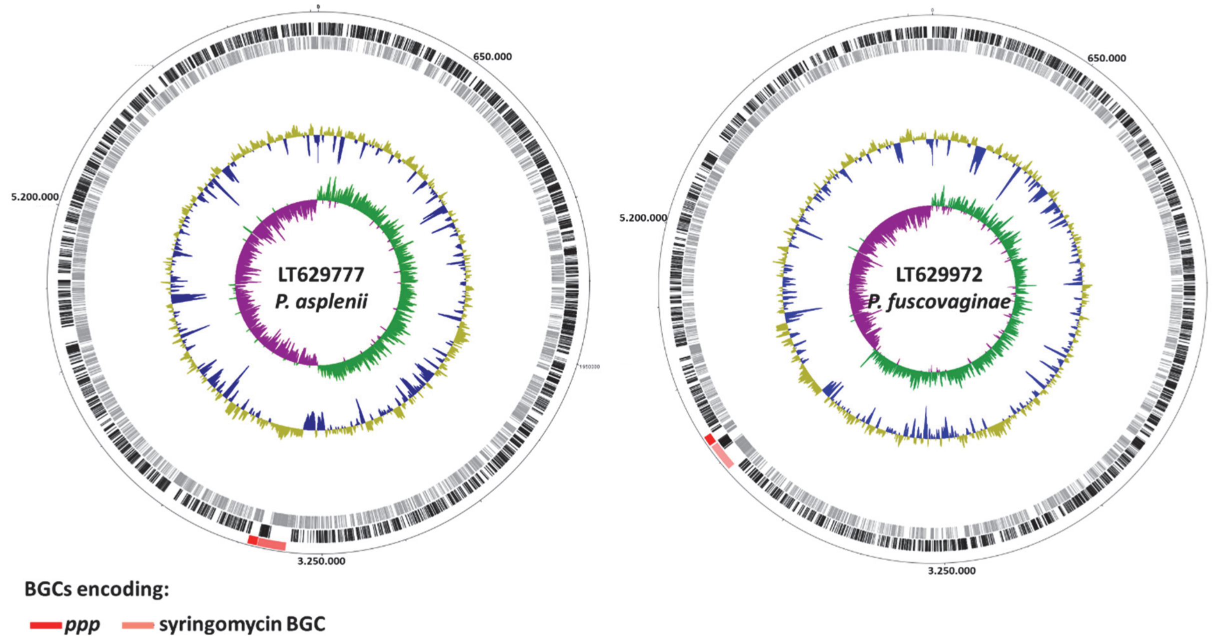

**Figure S99.** Location of the ppp gene cluster in the **c**hromosomes of *Pseudomonas asplenii* (LT62977) and *P. fuscvaginae* (LT629972). The outer circle depicts the size in base pairs. Lane 1 (from the outside in) shows the predicted *ppp* BGC (red) and syringomycin BGC (light red). Lanes 2 and 3 show predicted open reading frames (ORFs) on the leading (black) and lagging (gray) strands, respectively. Lanes 4 and 5 depict normalized plot of GC content (yellow/blue) and normalized plot of GC skew (purple/green), respectively. The chromosome is oriented to *dnaA*.

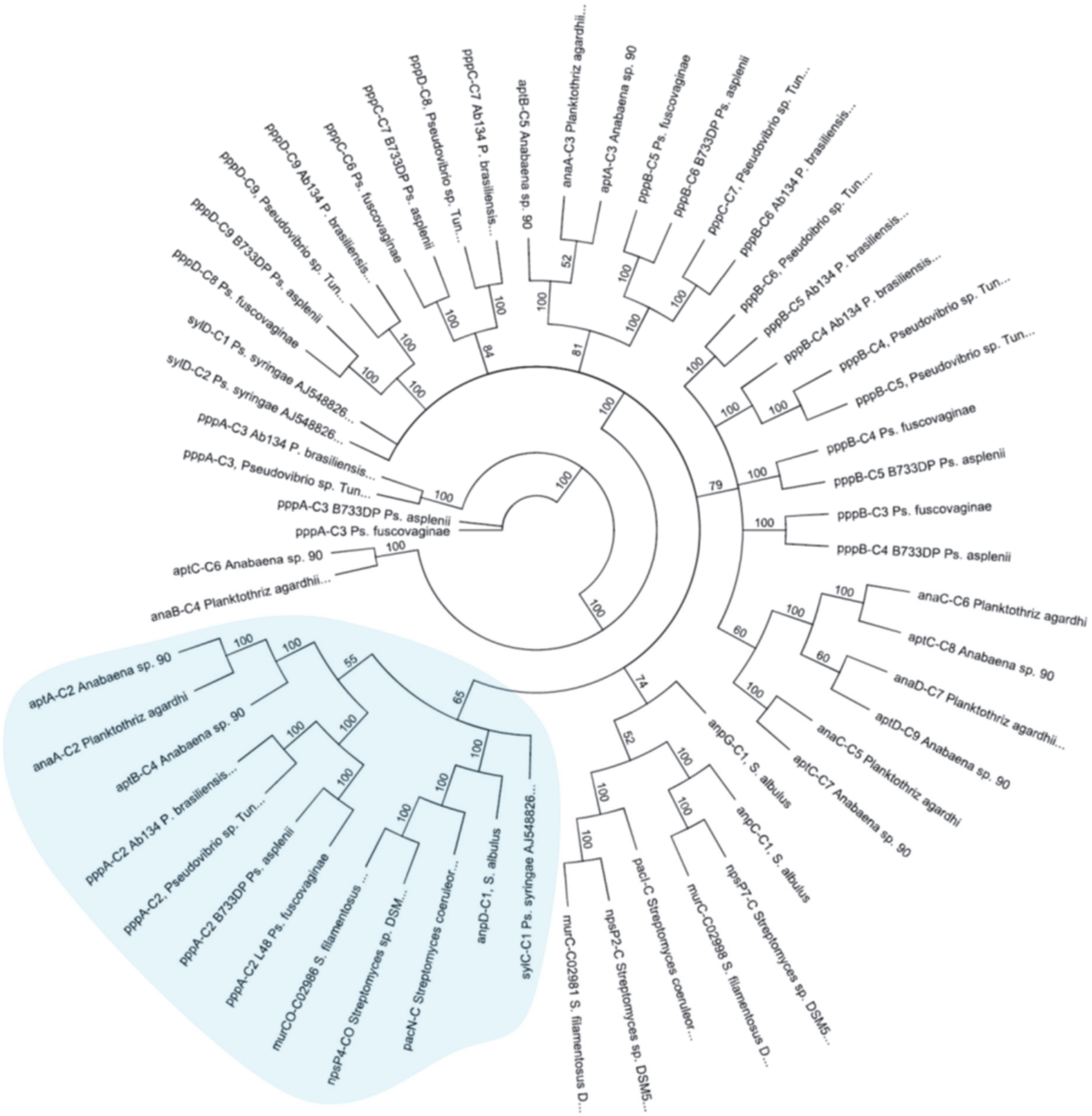

**Figure S100.** Phylogenetic analysis of condensation domains of ureido-containing natural product BGCs. The ureido C domain branch is highlighted in blue. C domains were aligned with MUSCLE and the tree was obtained using neighbor-joining algorithm implemented in the Geneious Prime software. GenBank accession codes are depicted on Table S19 of encoding ureido-linkage natural products.

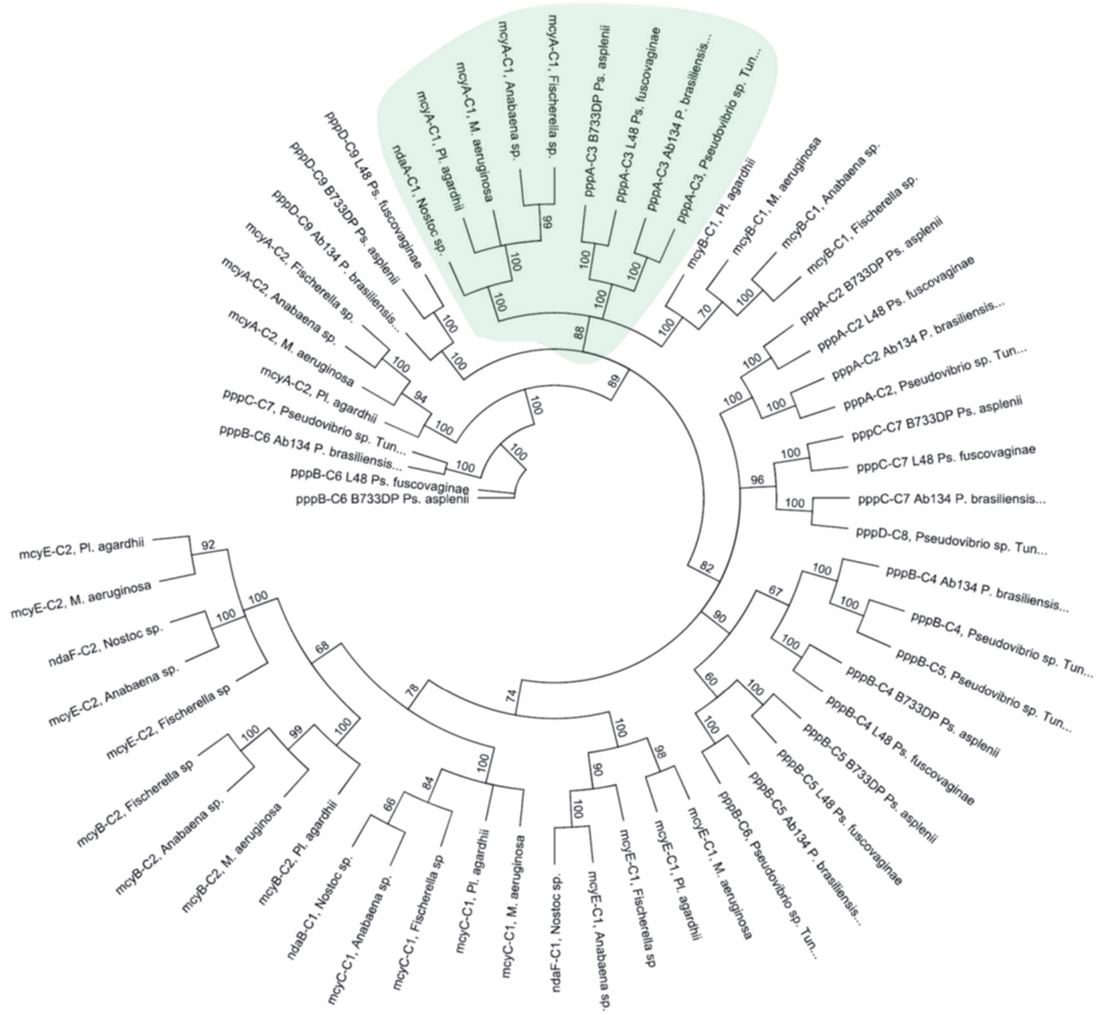

**Figure S101.** Phylogenetic analysis of condensation domains of Dhb-containing natural product BGCs. The Dhb/Dha C domain branch is highlighted. The third branch in the same node incorporates D-Ala (mycB-C1) as there is an epimerazation domain in the preceeding module (located in mycA). C domains of *ppp*, *mcy* and *nod* BGC were aligned with MUSCLE and the tree was obtained using neighbor-joining algorithm implemented in the Geneious Prime software. GenBank accession codes are depicted on Table S20.

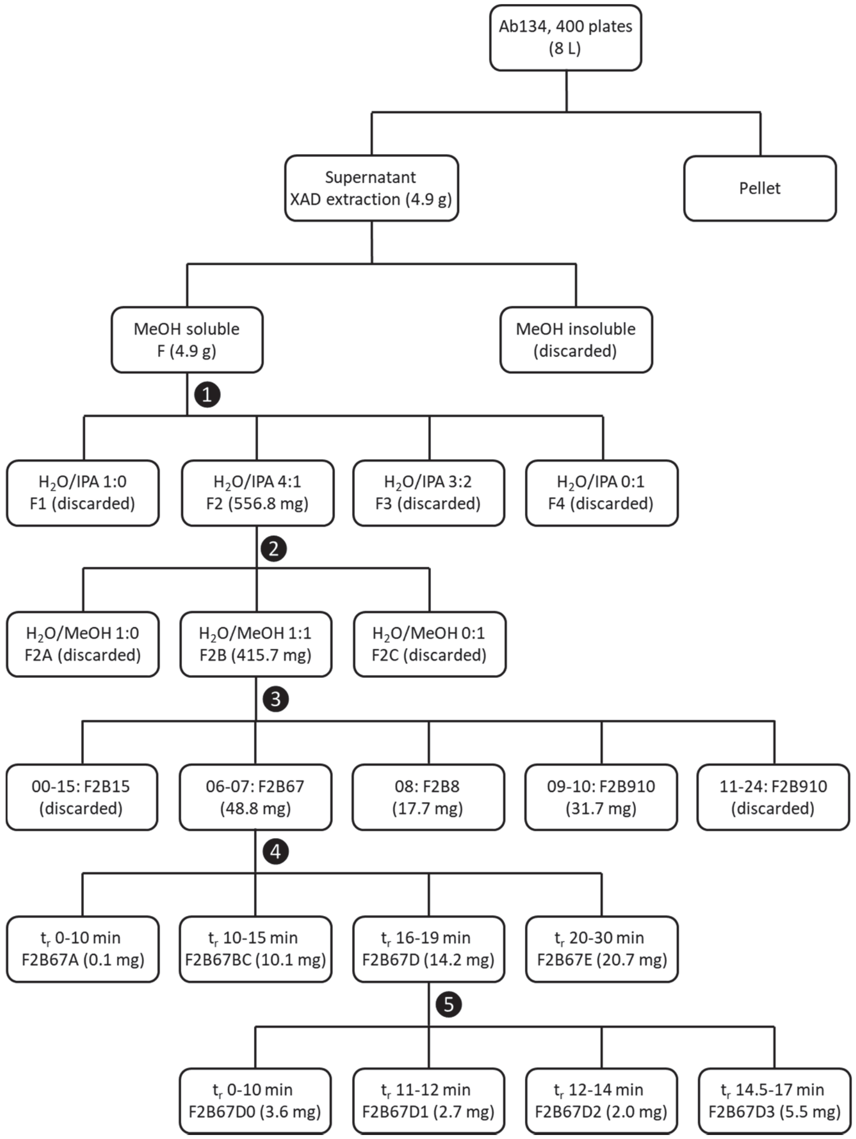

**Figure S102.** Fractionation scheme of extract F obtained from 400 swarming plates of *P. brasiliensis* Ab134. Summary of steps: ❶ Diaion HP-20SS cartridge, gradient of IPA in $H_2O$. ❷ C18 cartridge, gradient of MeOH in $H_2O$. ❸ Semi-preparative HPLC, C18 column, gradient of MeOH/MeCN in $H_2O$. ❹ Semi-preparative HPLC, C18 column, isocratic. ❺ Analytical HPLC, HILIC column, isocratic. Details of the fractionation are described on the Methods section.

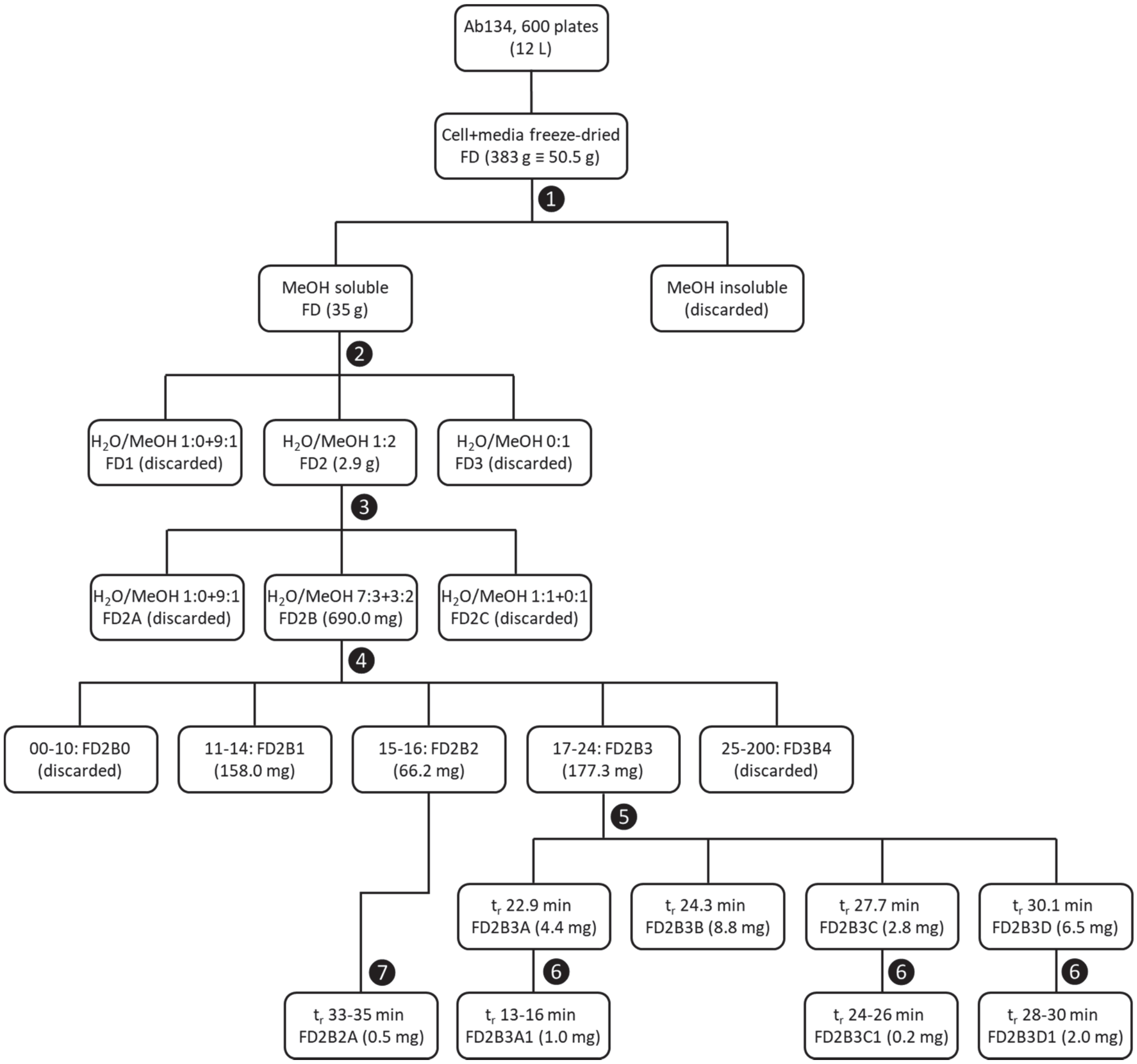

**Figure S103.** Fractionation scheme of extract FD obtained from 600 swarming plates of *P. brasiliensis* Ab134. Summary of steps: ❶ Clean-up by MeOH solubilization. ❷ C18 cartridge, gradient of MeOH in $H_2O$. ❸ Phenyl cartridge, gradient of MeOH in $H_2O$. ❹ Size-exclusion Biogel-P2 column, isocratic. ❺ Analytical HPLC, C18 PFP column, isocratic. ❻ and ❼ Analytical HPLC, Phenyl column, isocratic. Details of the fractionation are described on the Methods section.

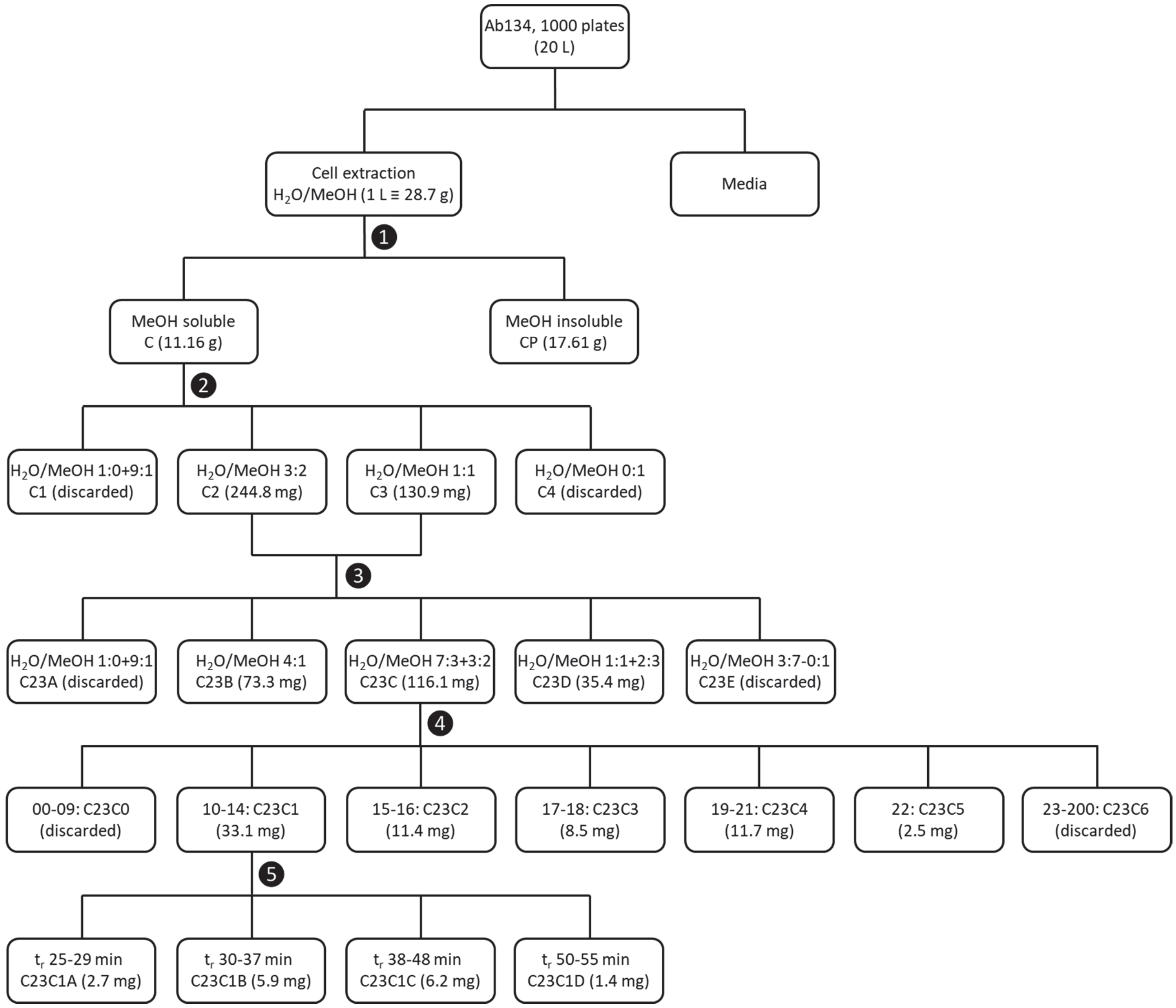

**Figure S104.** Fractionation scheme of extract C obtained from 1000 swarming plates of *P. brasiliensis* Ab134. Summary of steps: ❶ Clean-up by MeOH solubilization. ❷ C18 cartridge, gradient of MeOH in $H_2O$. ❸ Phenyl cartridge, gradient of MeOH in $H_2O$. ❹ Size-exclusion Biogel-P2 column, isocratic. ❺ Analytical HPLC, Phenyl column, isocratic. Details of the fractionation are described on the Methods section.
